# Dynamic Causal White Matter Atlas of Auditory and Visual Speech Networks at Millisecond Resolution: Intracranial Evidence from 125 Patients

**DOI:** 10.1101/2025.07.15.664981

**Authors:** Ryuzaburo Kochi, Aya Kanno, Hiroshi Uda, Keisuke Hatano, Hidenori Endo, Michael Cools, Robert Rothermel, Aimee F. Luat, Eishi Asano

## Abstract

**Background and Objectives:** Since the era of Penfield, invasive neurophysiology has laid a lasting foundation for functional neuroscience by elucidating brain regions necessary for speech. However, whole-brain investigations have yet to distinguish the millisecond-scale dynamics and specific white matter pathways that support rapid naming in the auditory and visual domains.

**Methods:** In this observational study, we constructed a whole-brain Dynamic Causal Tractography atlas using intracranial neurophysiological data from 125 neurosurgical patients. The resulting video atlas captured local cortical high-gamma activity and cortico-cortical coactivation via white matter tracts during rapid and delayed auditory and picture naming. Direct electrical stimulation was employed to assess the causal significance of the observed neural dynamics.

**Results:** The atlas revealed white matter coactivation intensity patterns at specific 5-millisecond time windows that best aligned with sensorimotor and language symptoms elicited by electrical stimulation (Spearman’s ρ = 0.58–0.91; p = 4.8 × 10cc to 1.6 × 10c²c). Rapid auditory naming was associated with deactivation of the right rostral middle frontal gyrus and increased coactivation along the left arcuate fasciculus, linked to stimulation-induced receptive and expressive aphasia. In contrast, delayed auditory naming correlated with a late surge in bifrontal coactivation. Rapid picture naming involved early coactivation between cortices connected via the bilateral inferior longitudinal fasciculi—associated with stimulation-induced visual distortions—coinciding with transient co-inactivation of Broca’s area.

**Discussion:** These findings delineate dissociable white matter–mediated mechanisms supporting rapid naming in the auditory and visual domains. Reduced inhibitory monitoring by the right dorsolateral prefrontal cortex may facilitate efficient lexical retrieval via left perisylvian pathways during auditory naming. In contrast, excessive bifrontal interaction may underlie delayed auditory naming. Rapid visual object recognition appears to rely on early occipitotemporal coactivation with minimal involvement of Broca’s area. The resulting atlas—accompanied by a publicly available dataset (61.2 GB) and analysis code—serves as a valuable resource for students and trainees studying the network dynamics underlying speech, as well as for presurgical language mapping in patients undergoing cortical or subcortical intervention.

## Introduction

Cortical regions and white matter pathways involved in naming have been extensively studied, with the left arcuate fasciculus recognized as a critical structure.^1^ However, the timing and specific white matter pathways supporting rapid naming in the auditory and visual domains remain to be distinguished. Competing hypotheses propose that faster naming results from enhanced or suppressed neural interactions. Some suggest that Broca’s area and temporal regions activate before the motor cortex,^2,3^ supporting the notion that early, strong coactivation—particularly via the arcuate fasciculus—facilitates lexical retrieval. Other studies link faster naming to reduced activation in the left prefrontal and posterior temporal cortices,^4,5^ suggesting that suppression minimizes interference. Slower responses may reflect domain-general cognitive processes.^6^ Here, we analyzed the dynamics of local neural modulation and inter-regional communication through white matter pathways during auditory and picture naming. We tested these competing hypotheses by examining neural activity during faster and slower trials within the same individuals.

We developed a Dynamic Causal Tractography atlas to visualize patterns of local neural engagement and functional coactivation during rapid and delayed naming in the auditory and visual domains. Intracranial EEG (iEEG) recordings from non-epileptic regions enabled millisecond-resolution measurement of task-related high-gamma activity,^7^ which correlates with neuronal firing and hemodynamic responses.^8,9^ Prior iEEG studies reported that high-gamma activation best predicted postoperative language outcomes.^10,11^ White matter tracts connecting regions with concurrent high-frequency responses are suggested to facilitate coordinated neural interactions.^12^ Using a validated approach, we defined functional coactivation as a significant, sustained increase in high-gamma amplitude between structurally connected regions.^7,12^ Direct electrical stimulation was used to determine the causal role of the observed neural dynamics. By examining how patient demographics and epilepsy severity influenced task performance and high-gamma activity,^13^ we improved the clinical relevance and generalizability of the resulting Dynamic Causal Tractography atlas.

## Materials and methods

### Participants

Between January 2007 and December 2023, we enrolled patients with focal seizures who met the following criteria: native English speakers aged ≥4 years completing auditory or picture naming tasks during extraoperative iEEG at Detroit Medical Center (**eFigure 1; eTables 1-2**). Exclusion criteria included hearing or visual impairments, major brain malformations, prior epilepsy surgery, or right-hemispheric language dominance, as determined by Wada test or by left-handedness combined with congenital left-hemispheric neocortical lesions. These criteria ensured that the left hemisphere included essential language functions.^10,12^ The study was approved by the Wayne State University IRB, with written informed consent from guardians and assent from participants aged ≥13 years.

### Intracranial electrodes

Platinum disk or depth electrodes were placed intracranially. A cortical surface model was created for each patient, and electrode positions were localized by coregistering preoperative MRI with post-implant CT using FreeSurfer (http://surfer.nmr.mgh.harvard.edu).^14^ For group analysis, electrode coordinates were aligned to the FreeSurfer standard brain and assigned to regions of interest (ROIs) (**eFigure 2**).^13,15^ Compared to our previous study,^7^ we subdivided temporal and Rolandic regions into anterior/posterior and superior/inferior portions, based on evidence of their distinct functional roles.^16,17^

### iEEG

After electrode placement, patients underwent video-iEEG monitoring using the Nihon Kohden system (Nihon Kohden America, Foothill Ranch, CA). Signals were digitized at 1,000 Hz. Only artifact-free, nonepileptic sites^7^ —unrelated to seizure onset, interictal spikes, or MRI lesions—were included in the analysis.

### Tasks

Patients performed auditory and picture naming tasks.^7,10^ In the auditory task, up to 100 spoken prompts (e.g., “What flies in the sky?”) elicited brief noun responses. Stimulus and response timings were identified using a photosensor and microphone; response time was defined from stimulus offset to response onset. In the picture task, up to 60 grayscale images (e.g., “cat”) were shown on a monitor, and patients named each aloud. Response time was measured from stimulus to response onset. Trials without appropriate responses were excluded.^7^

### Time-frequency analysis

We analyzed high-gamma (70–110 Hz) amplitude using a bipolar montage and complex demodulation (5-ms/10-Hz bins^6^) in BESA Software (BESA, Gräfelfing, Germany), which is mathematically equivalent to a Gabor transform.^18^ Amplitude (square root of power) was calculated relative to a 400-ms baseline (200–600 ms before stimulus onset). iEEG traces were aligned to stimulus onset, stimulus offset, and response onset in the auditory task, and to stimulus and response onsets in the picture task. We quantified percent changes in high-gamma amplitude from −200 to +500 ms around stimulus onset, −500 to +500 ms around stimulus offset, and −500 to +500 ms around response onset. These changes were visualized on a FreeSurfer brain with 10-mm interpolation using MATLAB R2023 (MathWorks, Natick, MA).

### ROI analysis

We analyzed iEEG data across 66 ROIs, each with ≥5 artifact-free, nonepileptic electrode sites from ≥5 patients. Significant high-gamma augmentation was defined as a lower 99.99% CI bound >0 for five consecutive 5-ms bins; attenuation required the upper CI bound <0 for the same duration. Given high temporal autocorrelation (100-fold increase in conditional probability), the chance of observing significant augmentation in an ROI was estimated at 5.0×10c¹c.^7^

### Structural connectivity analysis

We provide new structural connectivity templates for 13 white matter pathways (**eFigures 3-5**). The white matter courses in children with drug-resistant epilepsy closely resemble those in healthy controls (r: 0.8),^12^ supporting the use of Human Connectome Project data (http://brain.labsolver.org/diffusion-mri-templates/hcp-842-hcp-1021)^19^ for template generation. Using DSI Studio (http://dsi-studio.labsolver.org/) in Montreal Neurological Institute space, we visualized the shortest streamlines between cortical ROIs under defined parameters.

### Functional coactivation and co-inactivation networks

We determined functional coactivation using the virtual brain framework^7,20,21^, and previously discussed its validity.^7^ Here, we identified time intervals and white matter pathways exhibiting functional coactivation or co-inactivation. ROI pairs met criteria if (i) both showed significant, simultaneous high-gamma changes across ≥five consecutive 5-ms bins, and (ii) were directly connected by a tractography-defined streamline. The estimated chance of such high-gamma co-augmentation (or co-attenuation) in at least two ROIs was 1.1 × 10c¹c.^7^ We visualized these interactions in 5-ms sliding steps (**Figures 1 and 2**). Functional coactivation and co-inactivation intensity between ROI pairs was defined as the square root of the product of their high-gamma percent changes.

**Figure 1.**
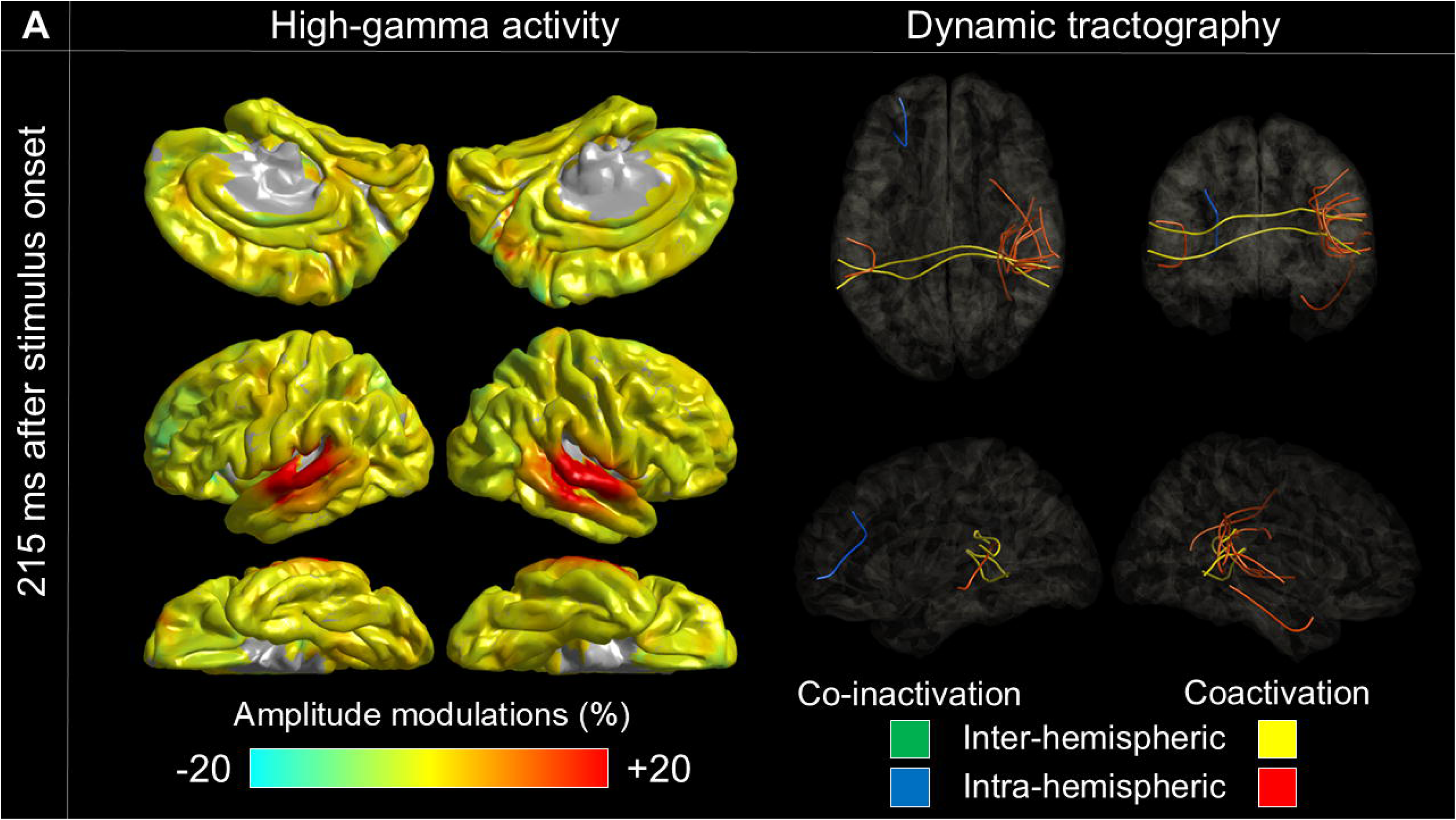
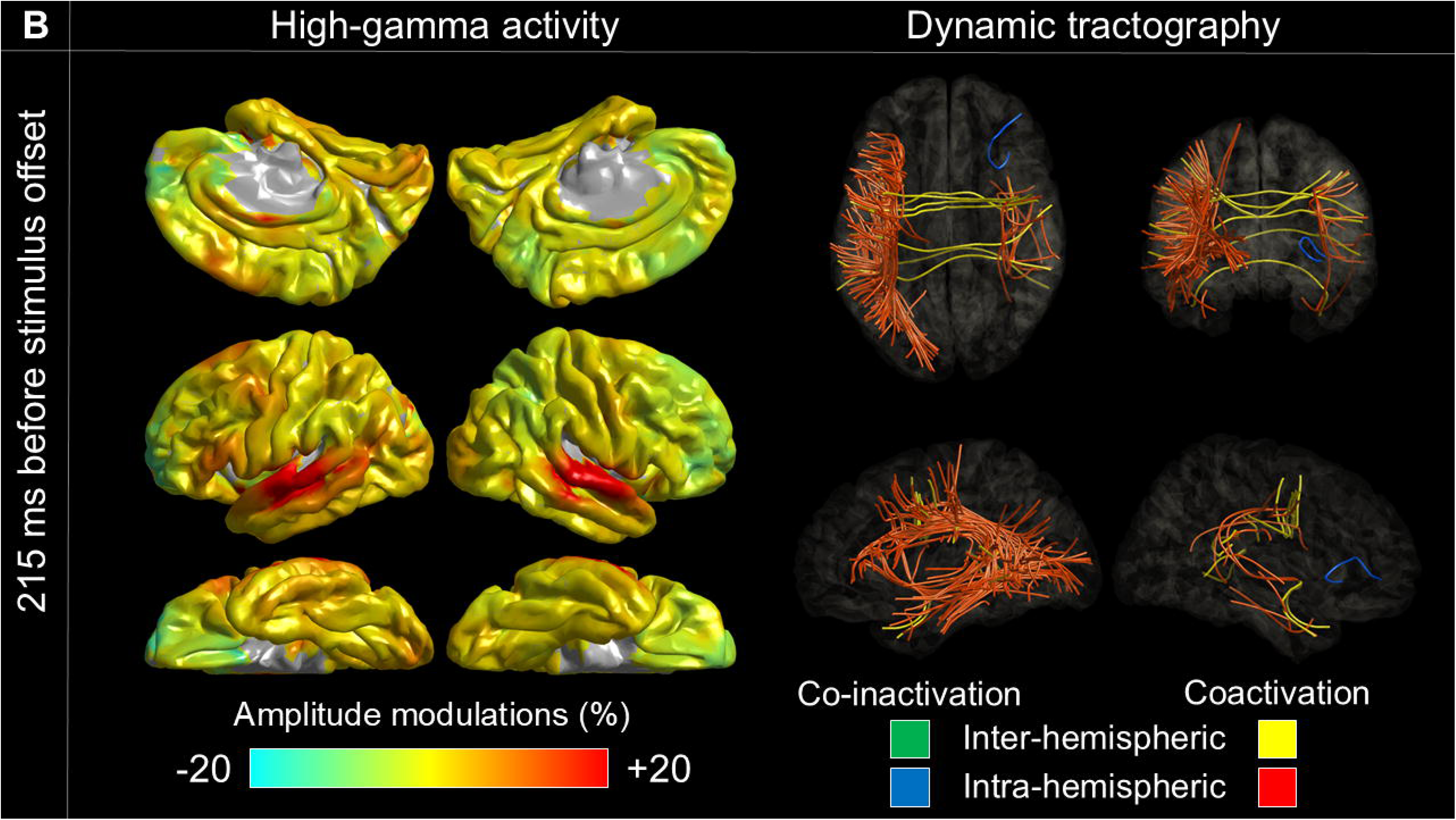
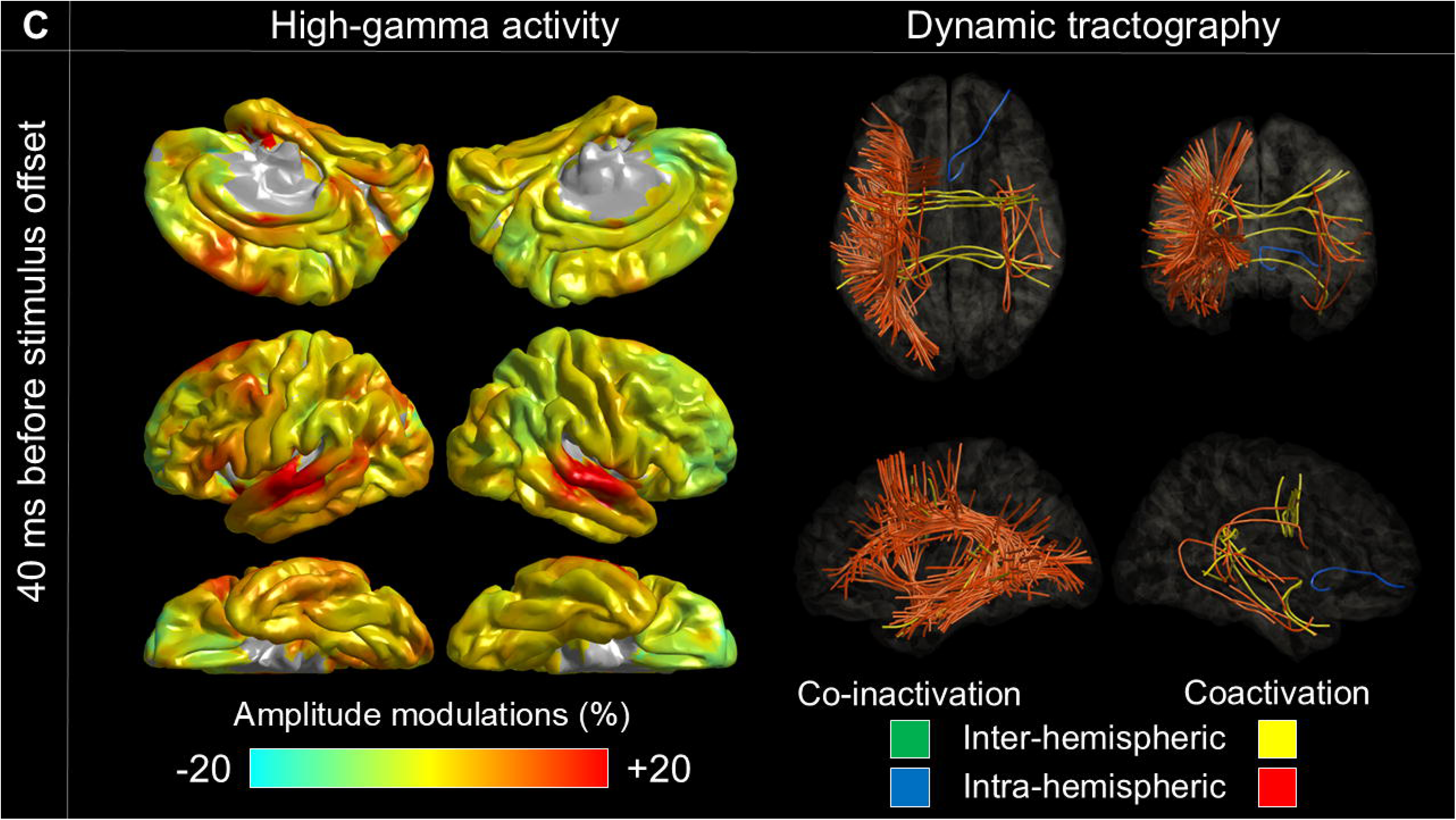
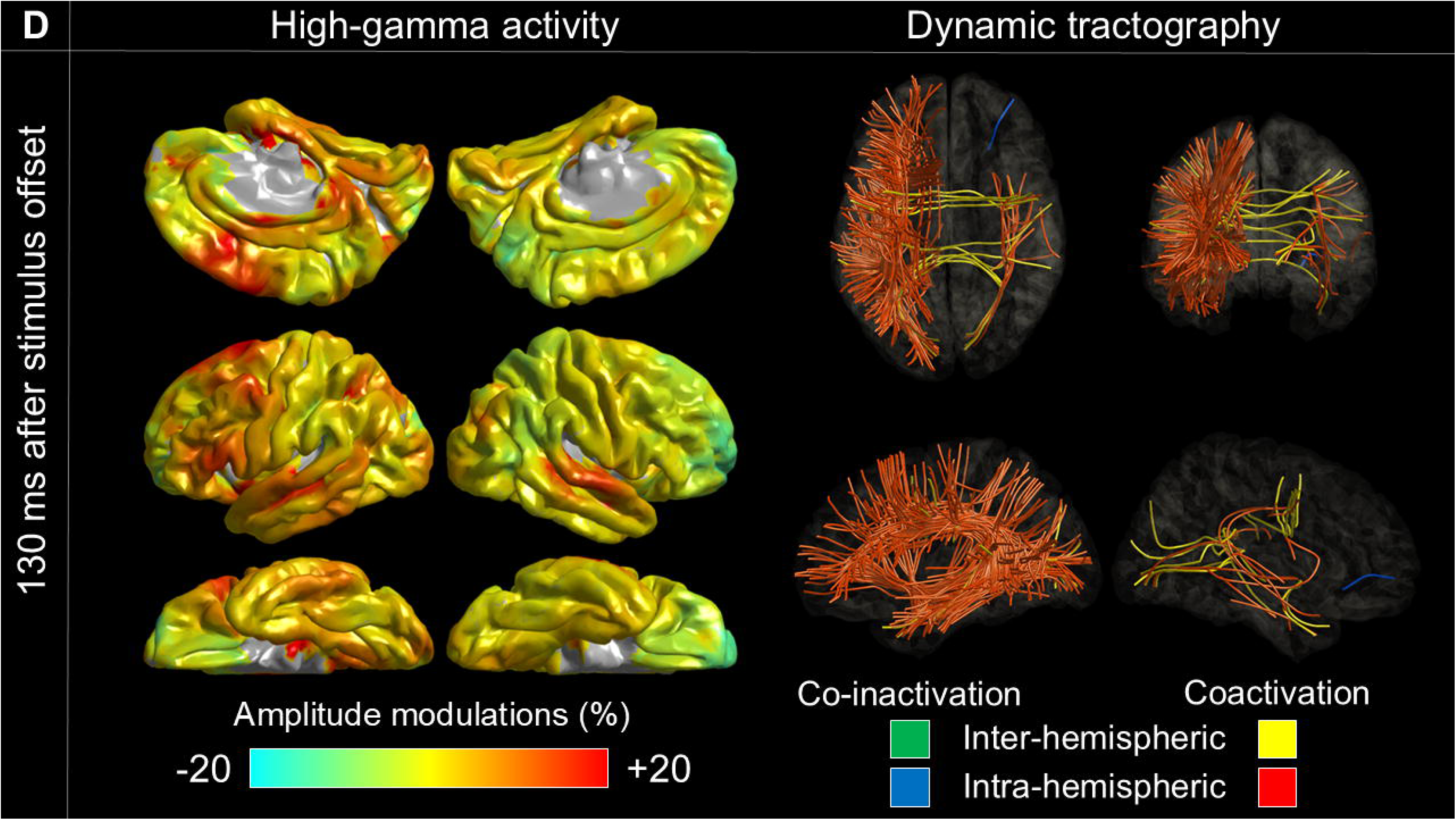
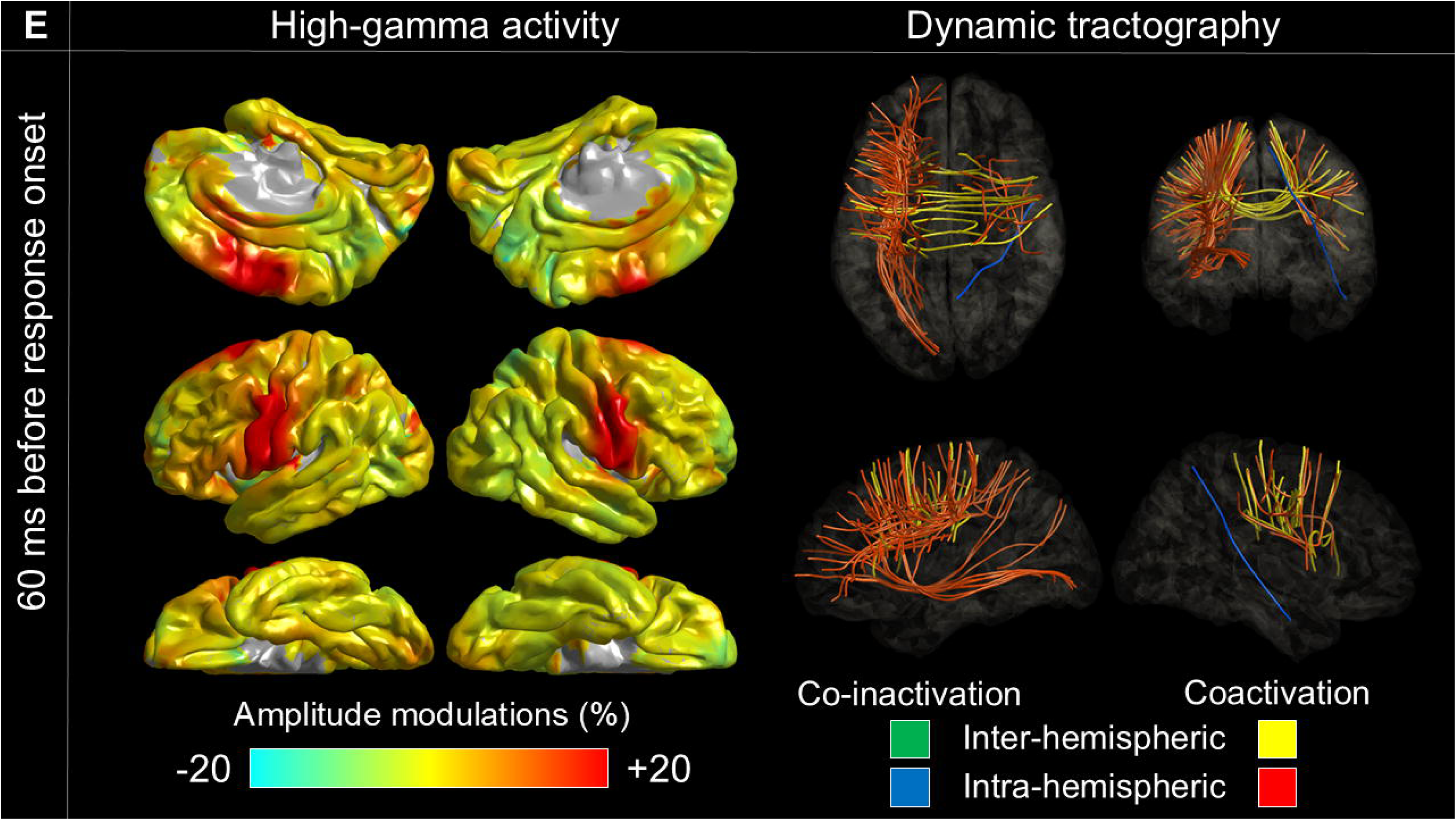
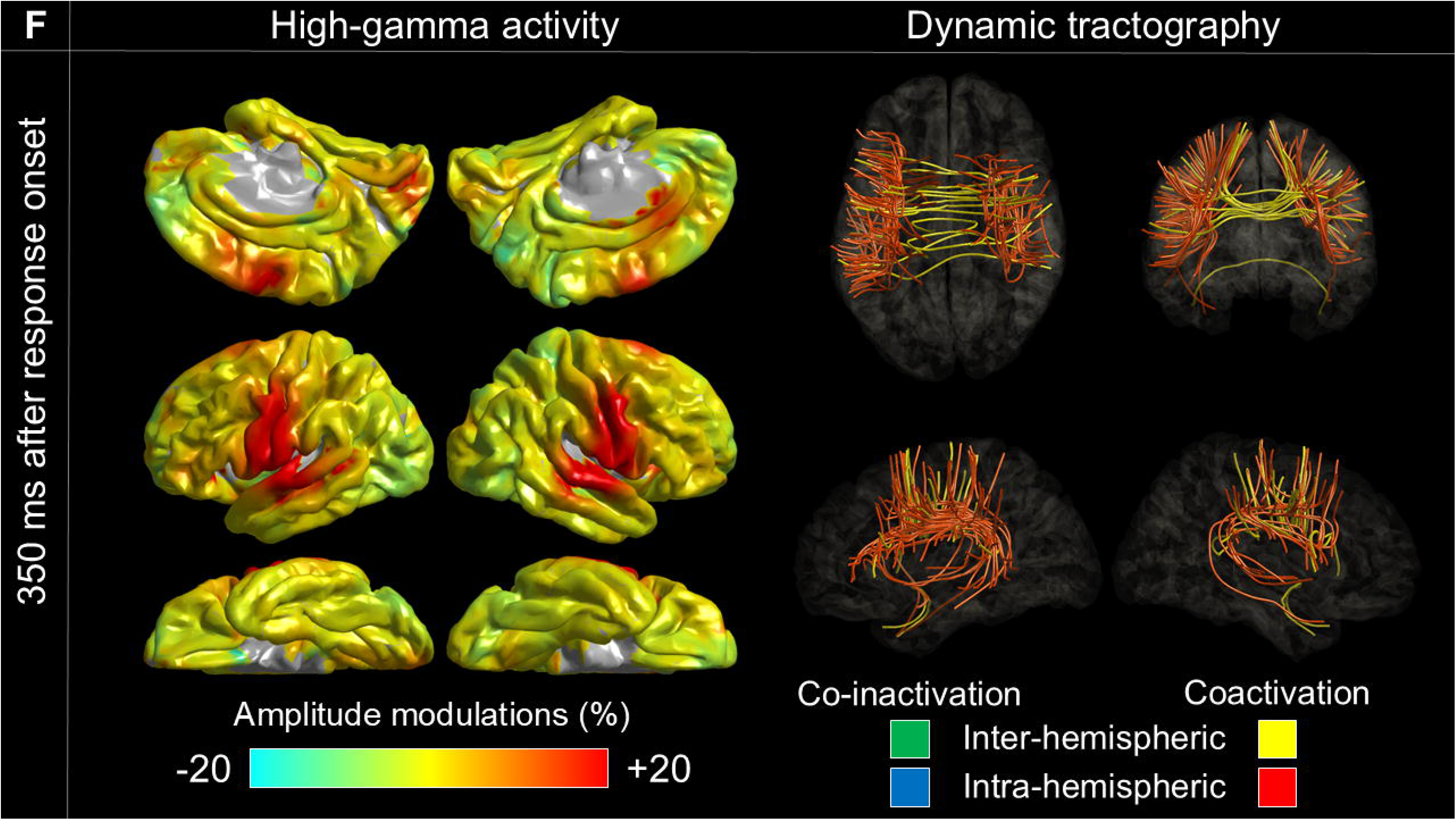
Cortical high-gamma amplitude and functional coactivation/co-inactivation during auditory naming. Left: task-related high-gamma changes. Right: coactivation (red/yellow) and co-inactivation (blue/green) streamlines. (A) 215 ms after stimulus onset; peak correlation with stimulation-induced auditory hallucination. (B) 215 ms before stimulus offset; correlation with faster responses through right rostral middle frontal high-gamma attenuation and left perisylvian coactivation. (C) 40 ms before stimulus offset; peak correlation with receptive aphasia. (D) 130 ms after stimulus offset; peak correlation with expressive aphasia. (E) 60 ms before response onset; peak correlation with speech arrest. (F) 350 ms after response onset; peak correlation with facial sensorimotor symptoms. See **Video 1** for full network dynamics. L: left; R: right.

**Figure 2.**
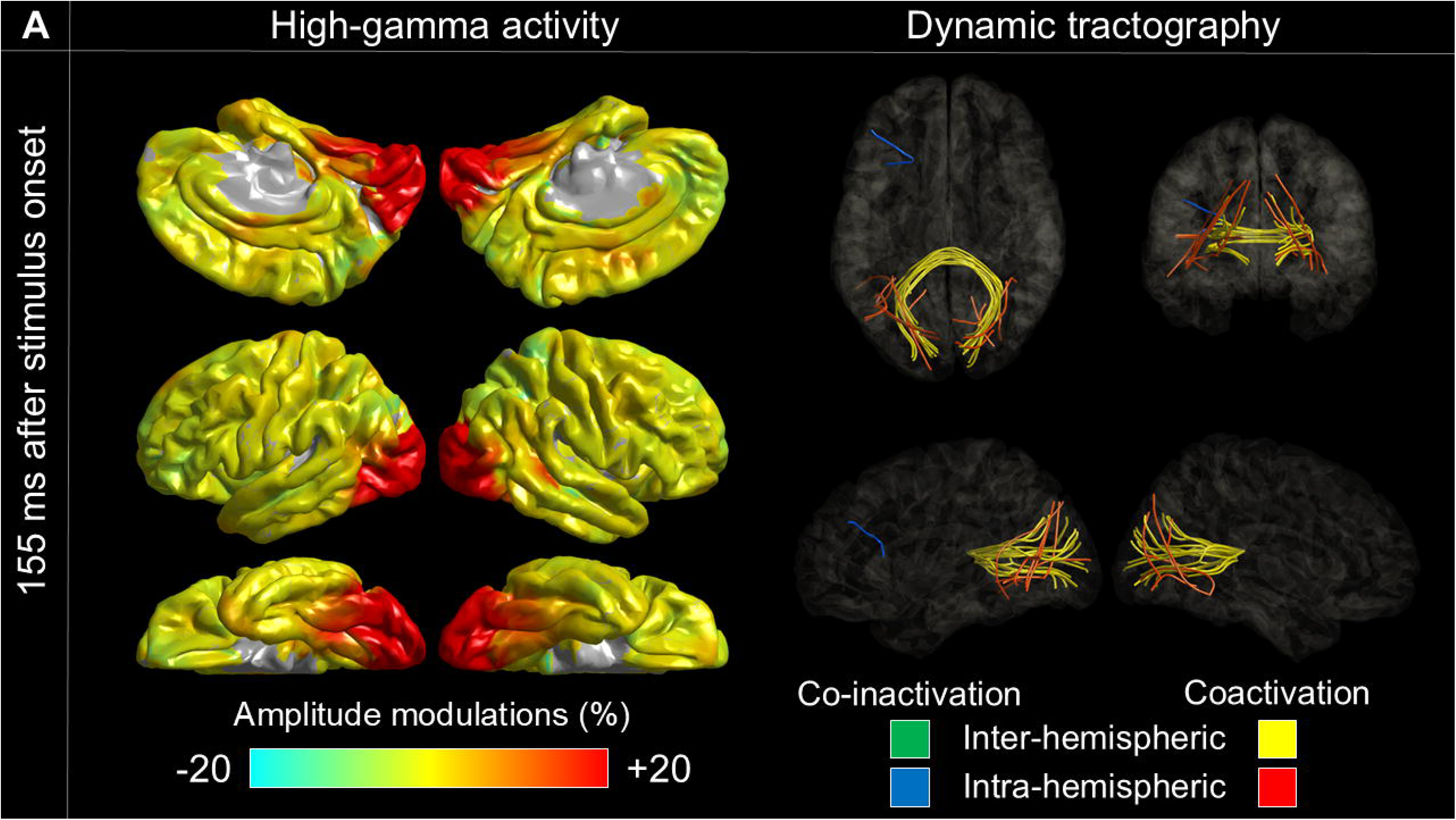
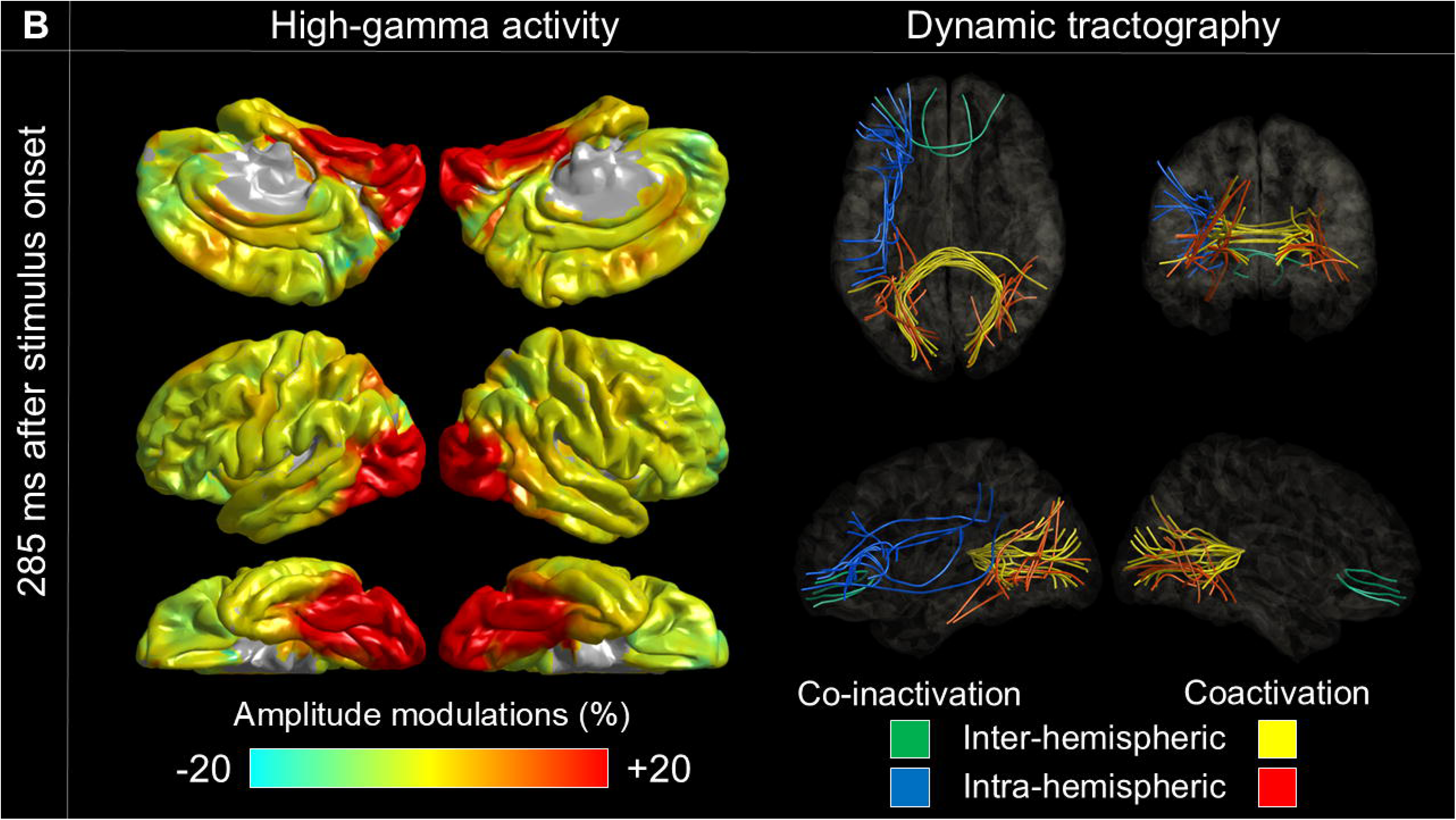
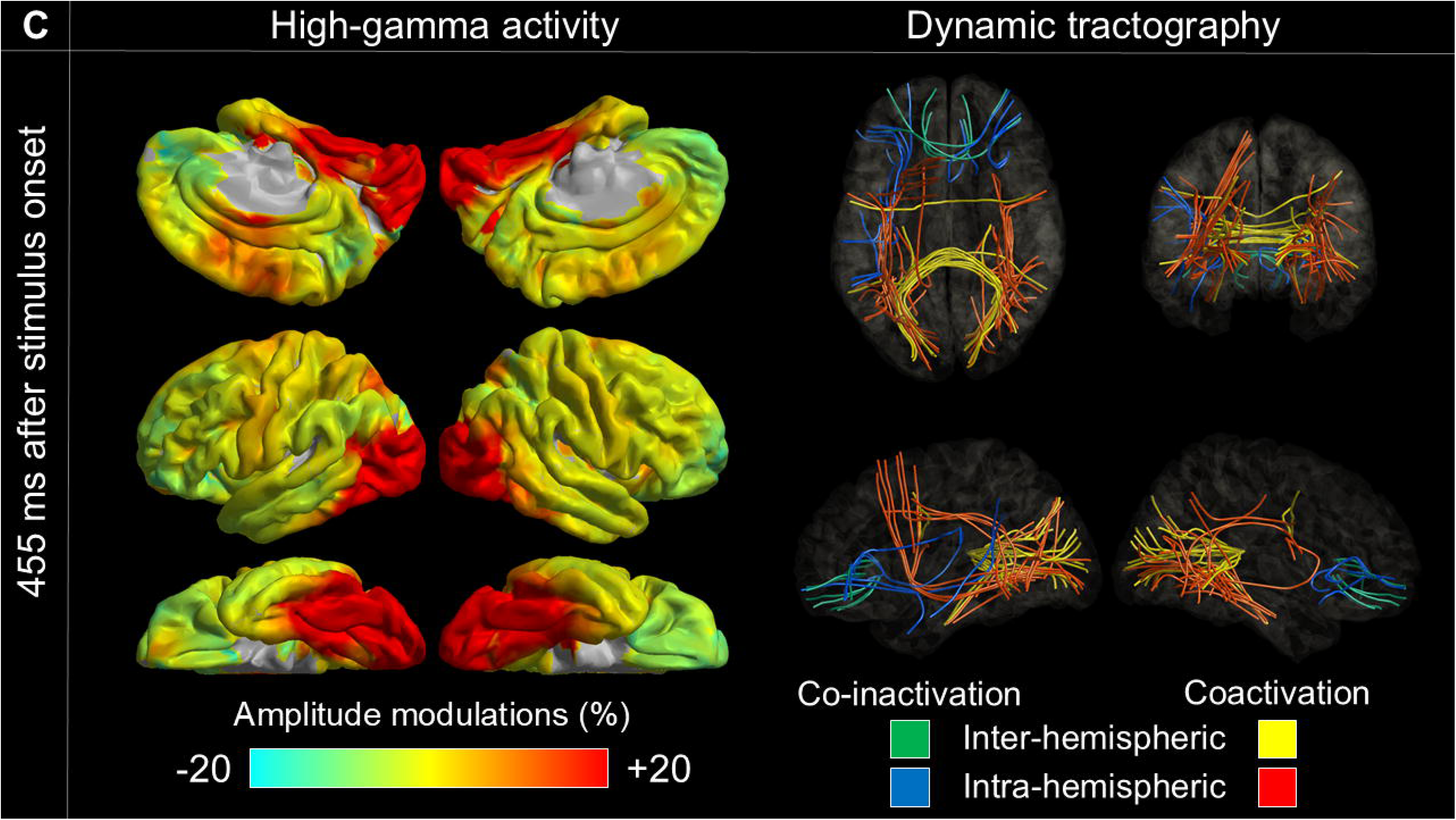
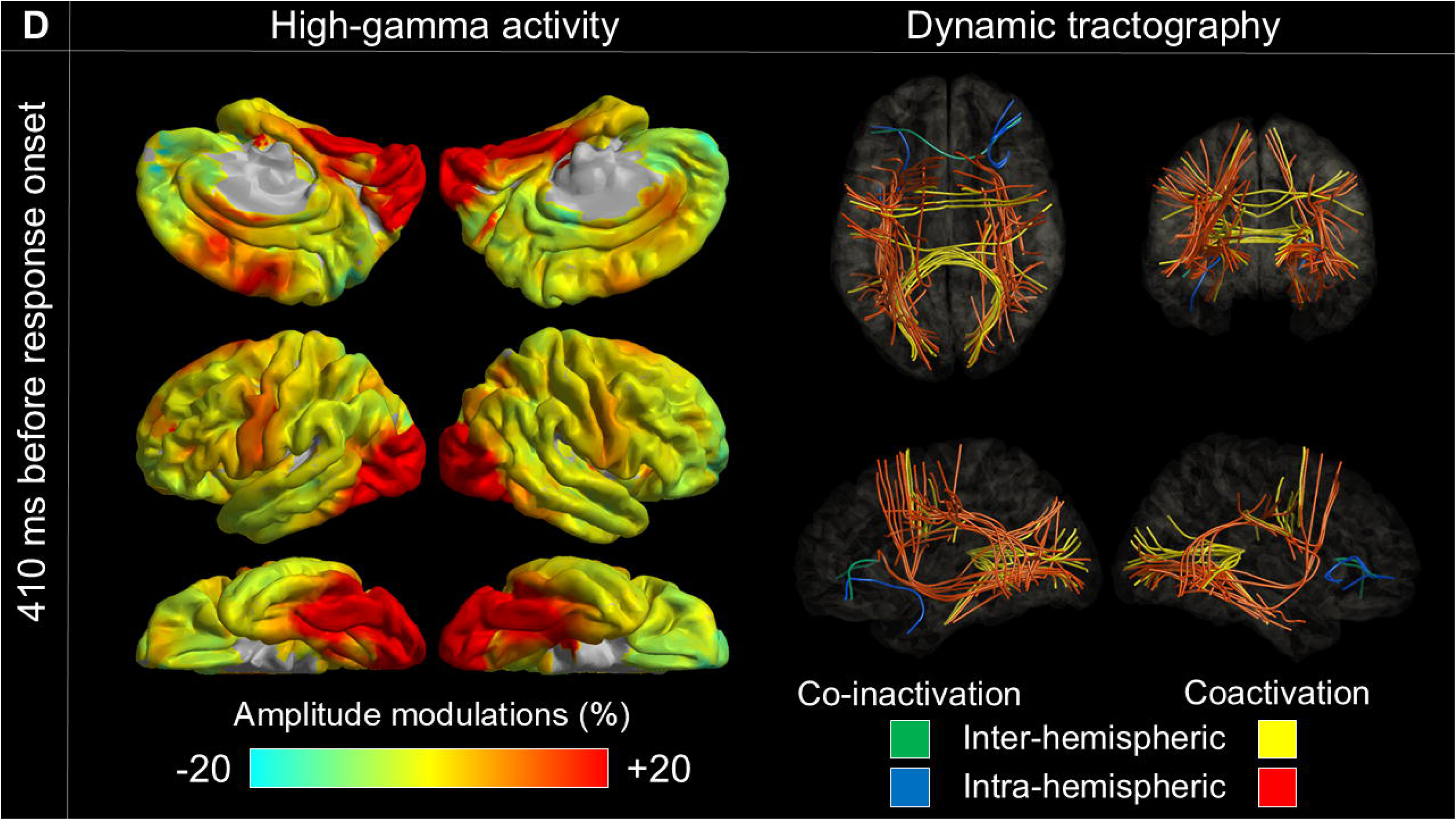
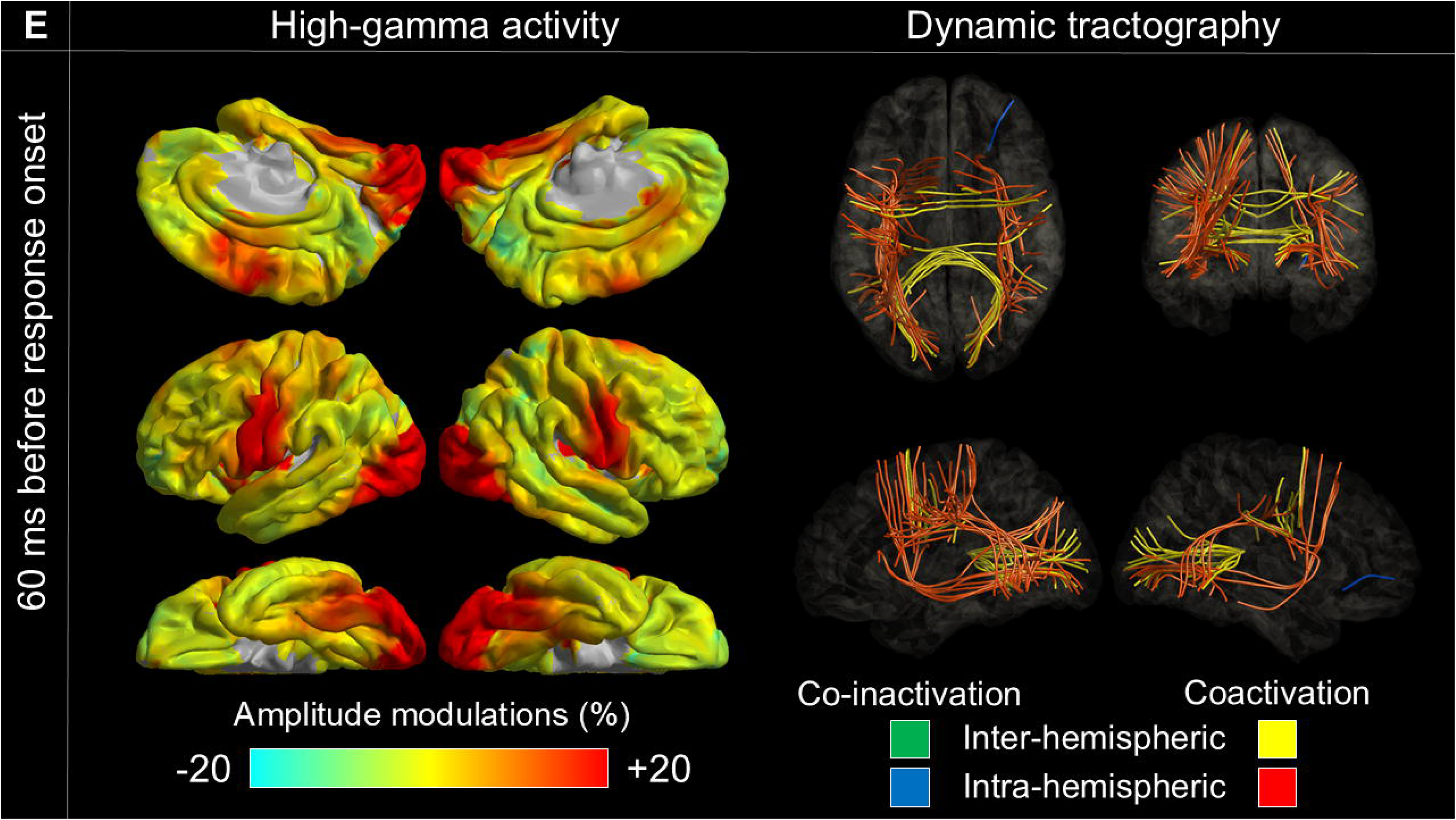
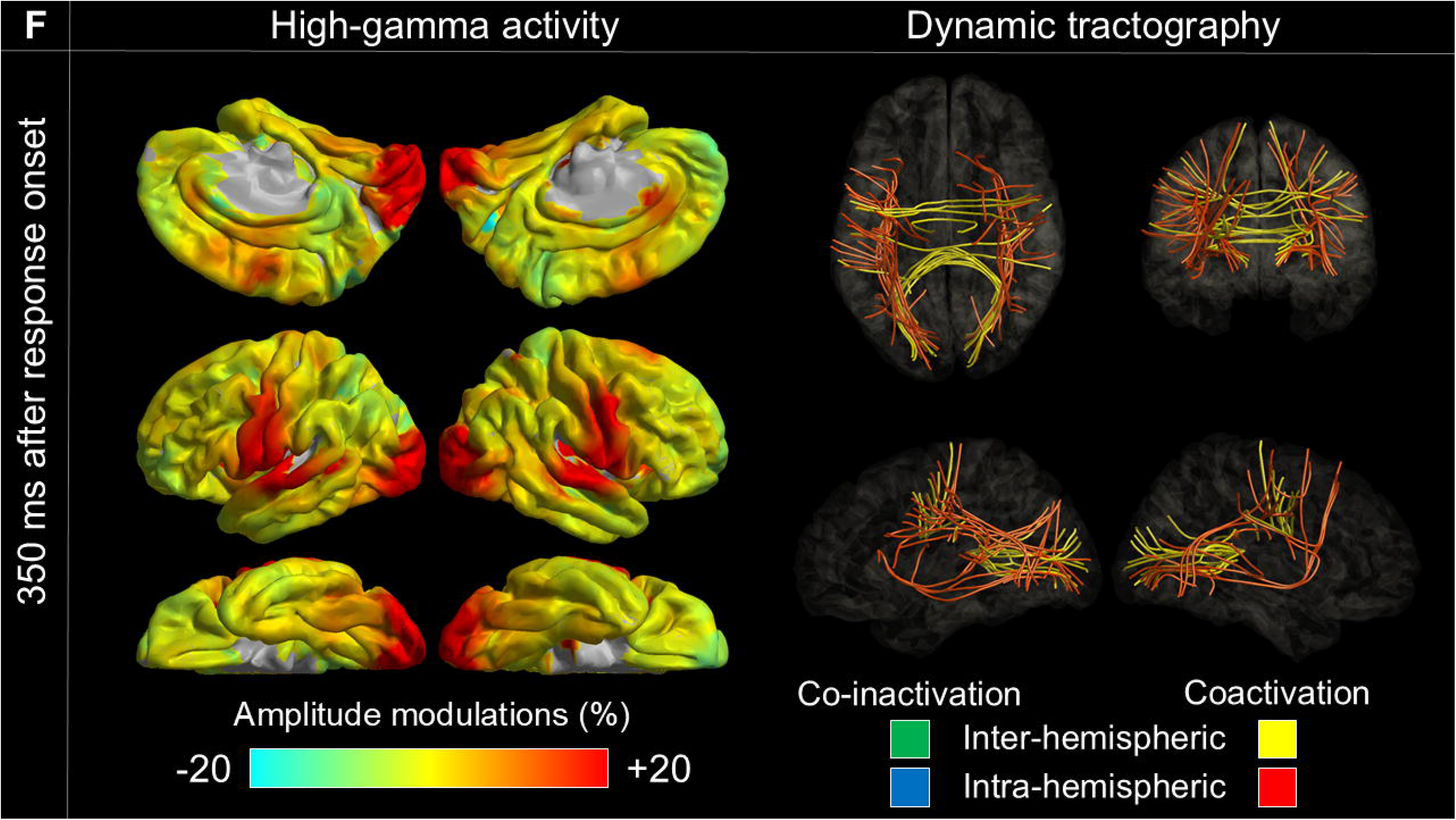
Cortical high-gamma amplitude and functional coactivation/co-inactivation during picture naming. (A) 155 ms after stimulus onset; peak correlation with stimulation-induced phosphene. (B) 285 ms after stimulus onset; peak correlation with stimulation-induced visual distortion. (C) 455 ms after stimulus onset; stronger left posterior inferior temporal coactivation linked to faster responses. (D) 410 ms before response onset. (E) 60 ms before response onset. (F) 350 ms after response onset. See Video 2 for full network dynamics.

### Stimulation

Comprehensive details are available in our publications.^7,10^ We performed stimulation mapping using 50-Hz pulse trains through adjacent electrode pairs. During stimulation, patients answered *wh-*questions or named objects, and blinded investigators verified reproducible effects. Supplementary tasks clarified site-specific functions. For each electrode, we calculated and averaged the probability of induced manifestations across patients, visualized on the FreeSurfer standard brain. Manifestations analyzed included auditory hallucinations, receptive/expressive aphasia, speech arrest, facial sensorimotor symptoms, phosphenes, visual distortions, and picture naming errors.

### Impact of patient profiles on response time and neural measures

We used mixed-effects models to assess how the following factors influence response time and high-gamma activity. Fixed factors included: square root of age (√years), handedness, left-hemisphere epileptogenicity, presence of an MRI-visible lesion, and number of antiseizure medications.^7^ Polytherapy was viewed as a proxy for greater cognitive burden.^22^ For response time, we used median values as the outcome, with intercept and patient as random effects. Analyses were performed in SPSS 28 (IBM, Chicago, IL), with significance set at two-sided p-value<0.05. To assess neural effects, we applied the same model to median high-gamma amplitude per electrode, calculated in 100-ms bins from stimulus onset to 500 ms post-response over a 2,500-ms window. Significance was defined using a Bonferroni-corrected p-value<0.05 across 66 ROIs and 25 bins.

### Functional coactivation and stimulation-induced manifestations

We examined whether functional coactivation at specific time bins was associated with stimulation-induced symptoms.^7^ Coactivation intensity was calculated every 5 ms for each ROI by averaging across all tract-connected ROI pairs. Using Spearman’s rank correlation, we tested whether higher coactivation (%) at an ROI predicted greater probability (%) of specific symptoms. For each symptom, the peak association was defined by the highest Spearman’s rho across 500 time bins. A Bonferroni-corrected two-sided p-value<0.05 was used.

### Neural correlates of within-individual variability in response times

We assessed how high-gamma activity and functional coactivation relate to response times within individuals. Each patient’s trials were sorted by response time into six categories to control for inter-patient variability. For each response time category, we calculated mean high-gamma amplitude and coactivation intensity across tract-connected ROI pairs. Spearman’s rank correlation assessed associations between neural measures (%) and response times.

## Results

### Patient profiles on response time

We found 125 eligible patients (age 5–49 years; 61 females; **eFigure 1; eTables 1-2**). Of these, 119 completed the auditory naming task, 108 the picture naming task, and 102 completed both. In total, 9,526 artifact-free nonepileptic sites were analyzed (9,109 for auditory naming, 8,515 for picture naming, and 8,098 for both; **eFigure 2**).

Increased √age was associated with shorter median response times in both auditory (mixed model estimate: –0.191; 95%CI: –0.314 to –0.068; p-value: 0.003) and picture naming tasks (estimate: –0.197; 95%CI: –0.323 to –0.071; p-value: 0.002). These effects were independent of other fixed variables, none of which showed significant associations (p-values: 0.083–0.970).

### Patient profiles on neural dynamics

Left-handedness was associated with a 15.1% increase in auditory naming-related high-gamma amplitude at the right entorhinal cortex (300–400 ms post-stimulus offset; 95%CI: 9.9–20.3%; p-value: 7.8×10cc). Left-hemispheric epileptogenicity was associated with a 29.6% increase in picture naming-related high-gamma at the right posterior STG (0–100 ms before response onset; 95%CI: 17.7–41.5%; p-value: 6.6×10cc). No other fixed variables showed significant effects.

### Cortical high-gamma modulations and functional coactivation during auditory naming

**eVideo 1** shows dynamic changes in high-gamma activity and functional coactivation/co-inactivation during auditory naming, with snapshots in **Figure 1**. Upon hearing stimuli, high-gamma increased bilaterally in the STG, with coactivation between bilateral STG via the corpus callosum and between STG and precentral gyrus via the arcuate fasciculus. As stimulus offset neared, high-gamma decreased in the right rostral MFG, while left-hemispheric coactivation intensified. After stimulus offset, coactivation expanded within the left hemisphere, then localized to the left frontal region before response onset. During the response, inter-hemispheric coactivation was observed between the bilateral inferior Rolandic cortices, along with intra-hemispheric coactivation between the STG and Rolandic cortices.

As part of the atlas, we illustrate the proportion of specific white matter pathways that exhibited functional coactivation (**eFigure 6**) and co-inactivation (**eFigure 7**) across time bins. **eFigure 8** displays the temporal dynamics of mean high-gamma modulations with confidence interval bars. **eFigure 9** presents high-gamma modulations analyzed using a leave-one-out approach, demonstrating consistency in temporal dynamics and confirming that the results were not driven by single outliers. **eFigure 10** illustrates response time–sorted neural dynamics at each ROI.

### Auditory naming-related functional coactivation and stimulation-induced manifestations

**Figure 3** shows probability maps of stimulation-induced symptoms, while **Figure 4** illustrates how correlations between functional coactivation and symptom probability evolve over time. Peak correlations occurred at distinct time points:

**Figure 3.**
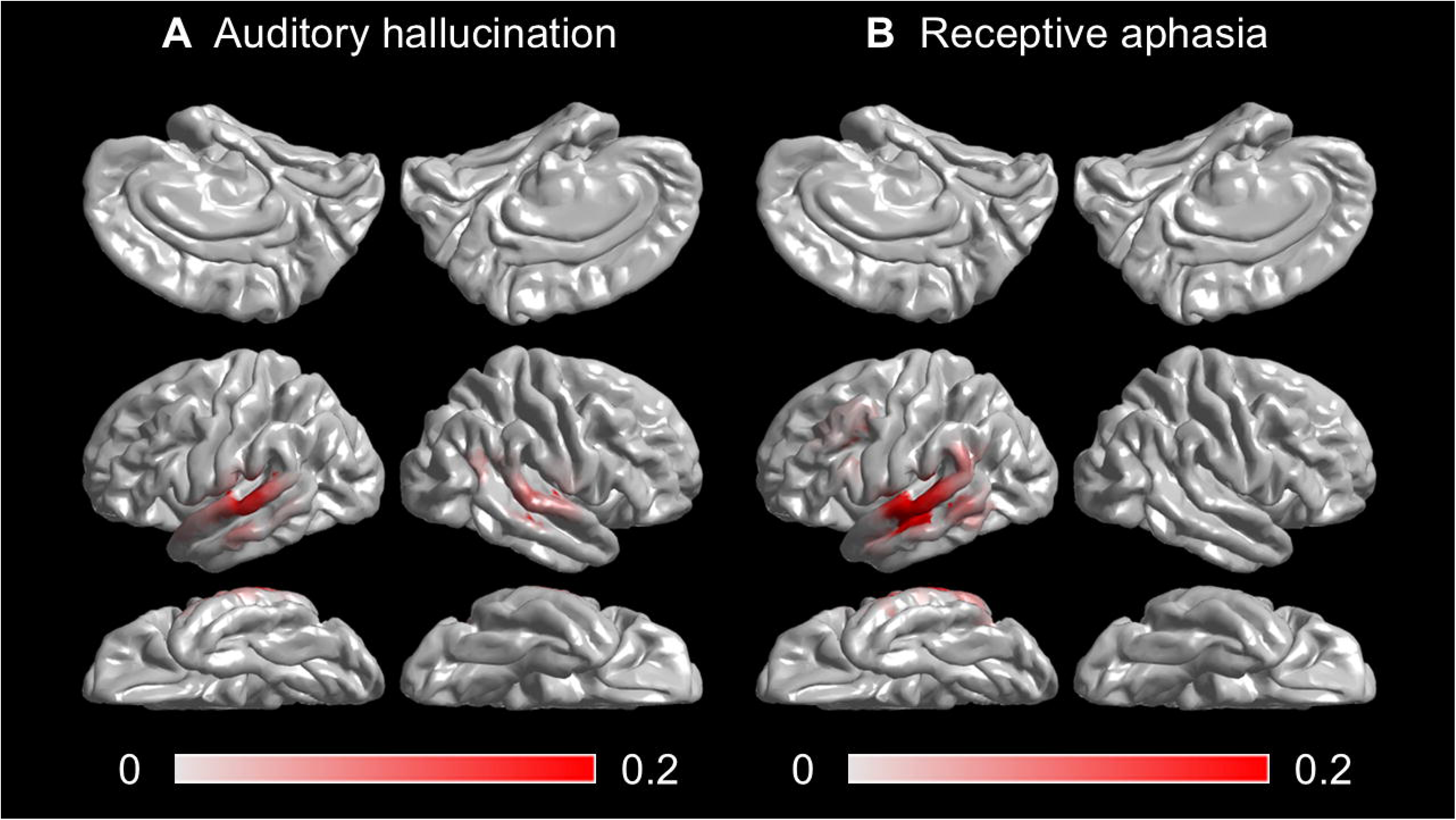
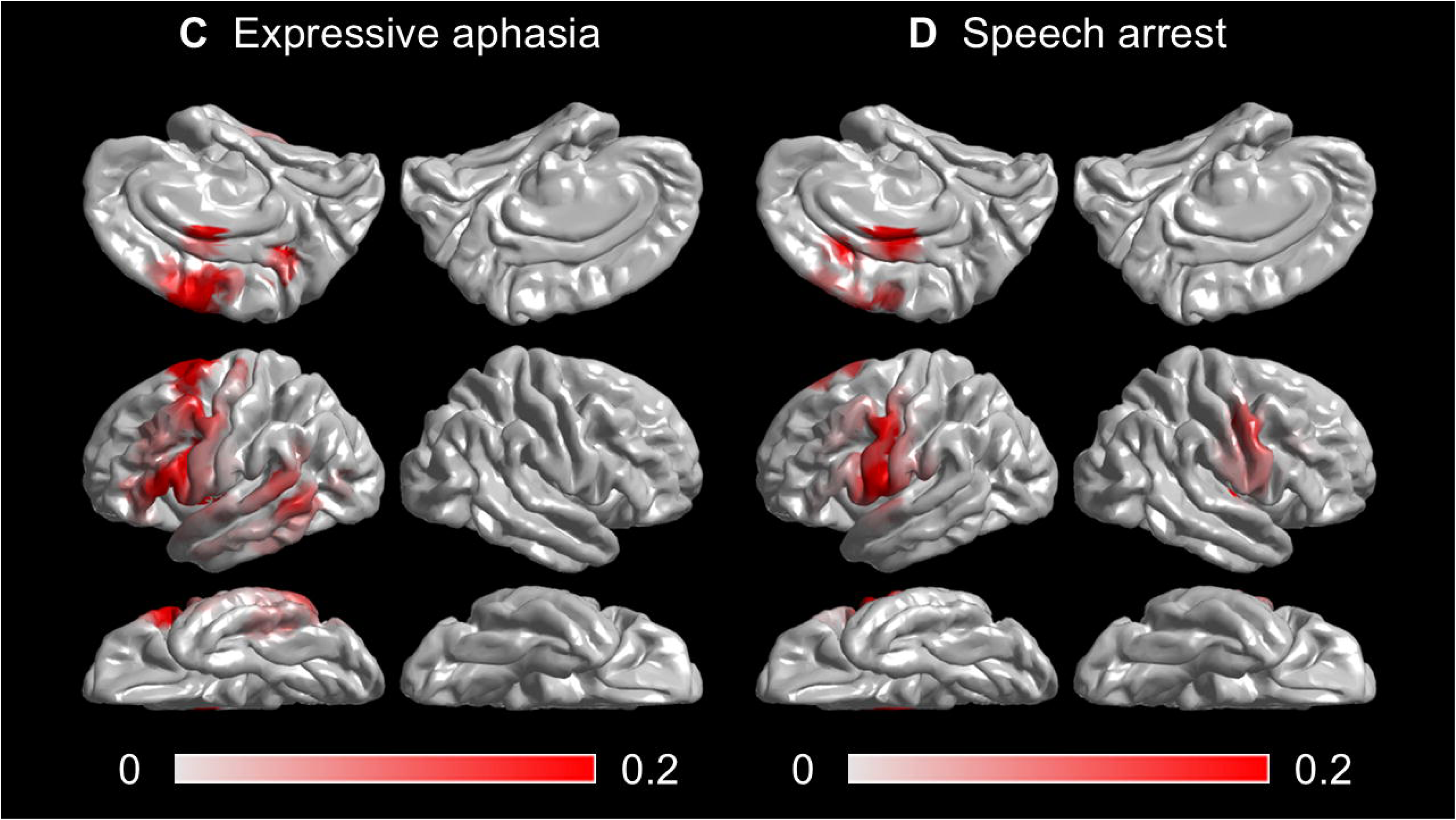
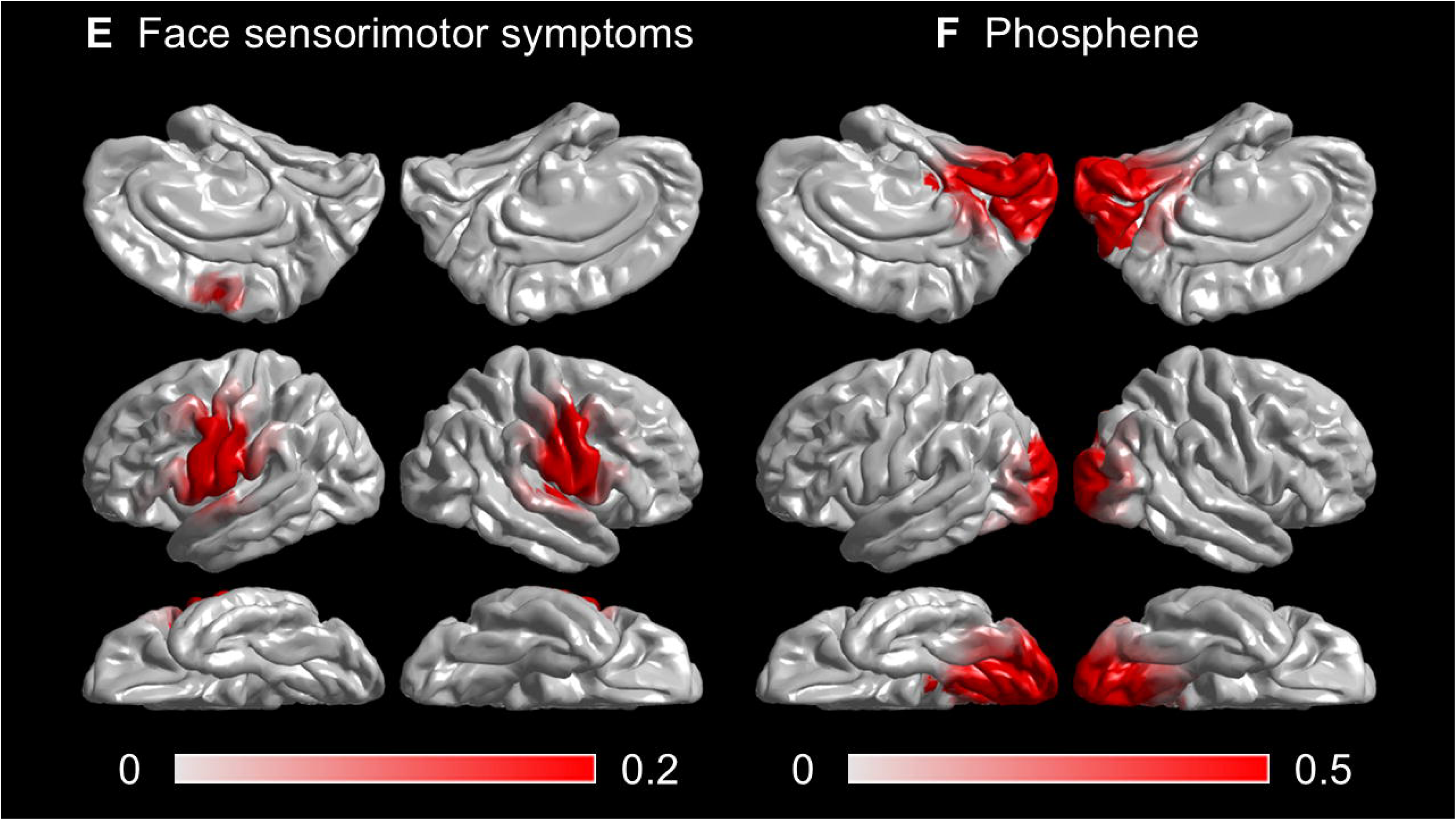
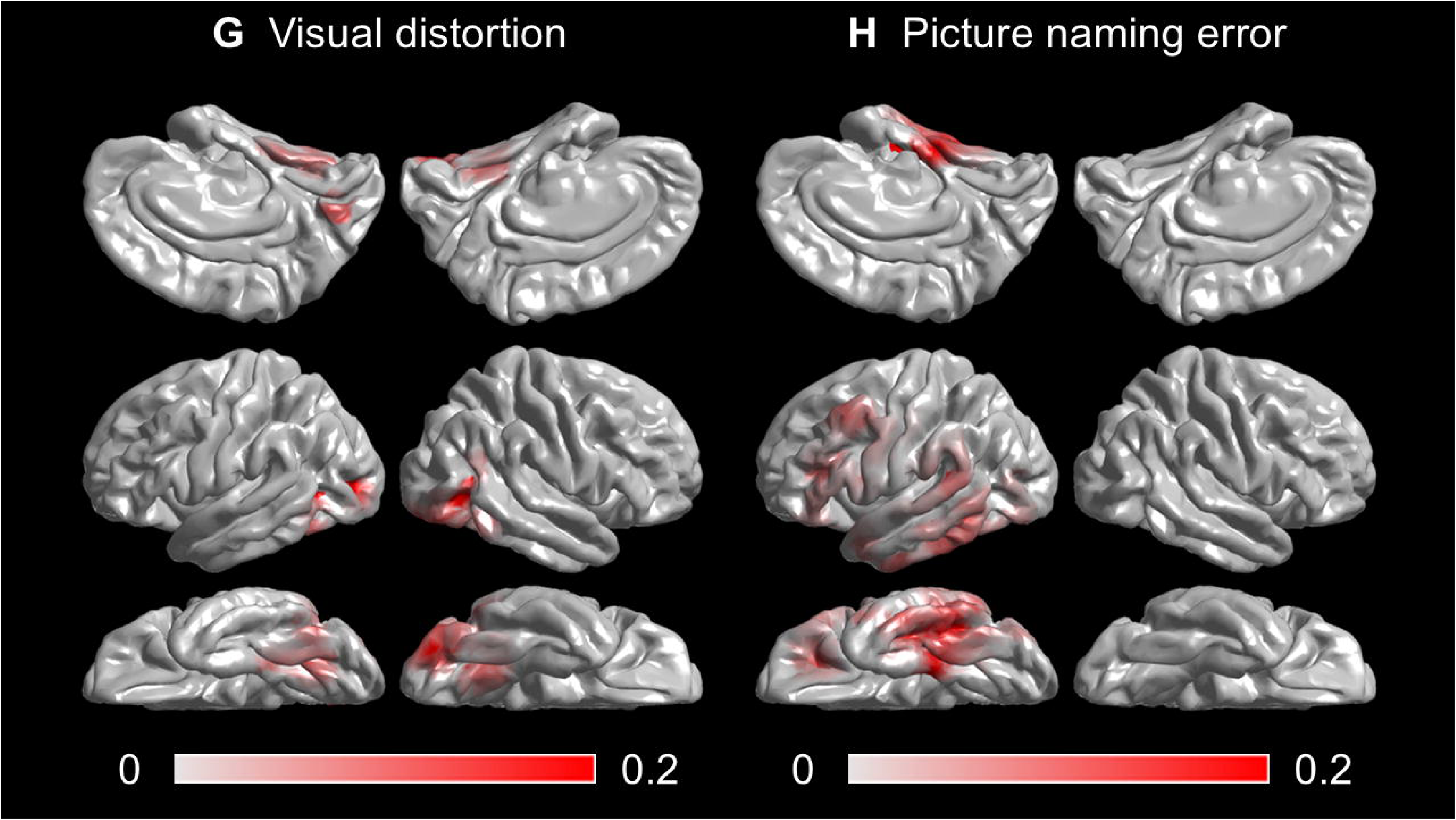
Probability map of stimulation-induced manifestations. (A) Auditory hallucinations. (B) Receptive aphasia. (C) Expressive aphasia. (D) Speech arrest. (E) Face sensorimotor symptoms. (F) Phosphene. (G) Visual distortion. (H) Picture naming error.

**Figure 4.**
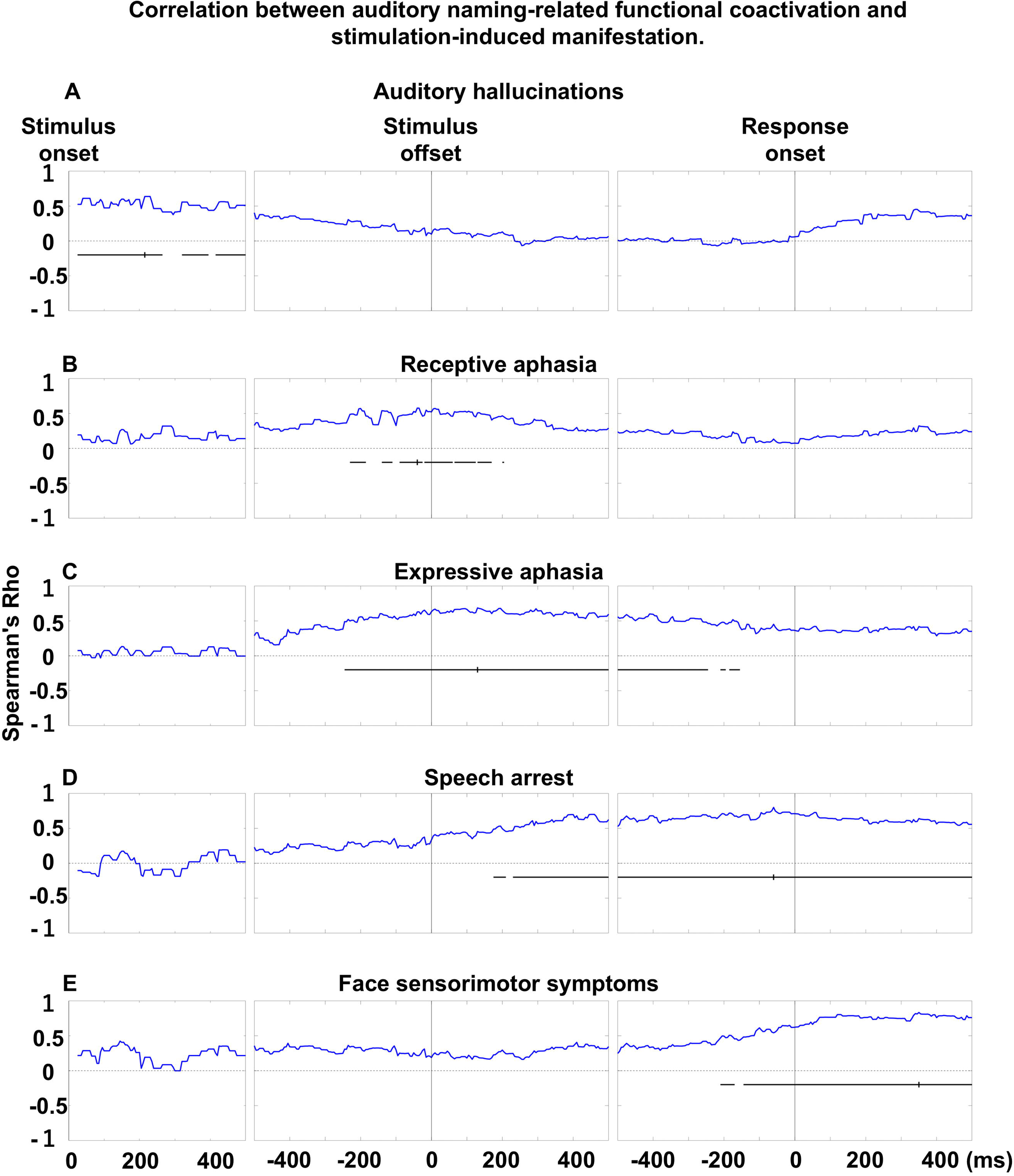
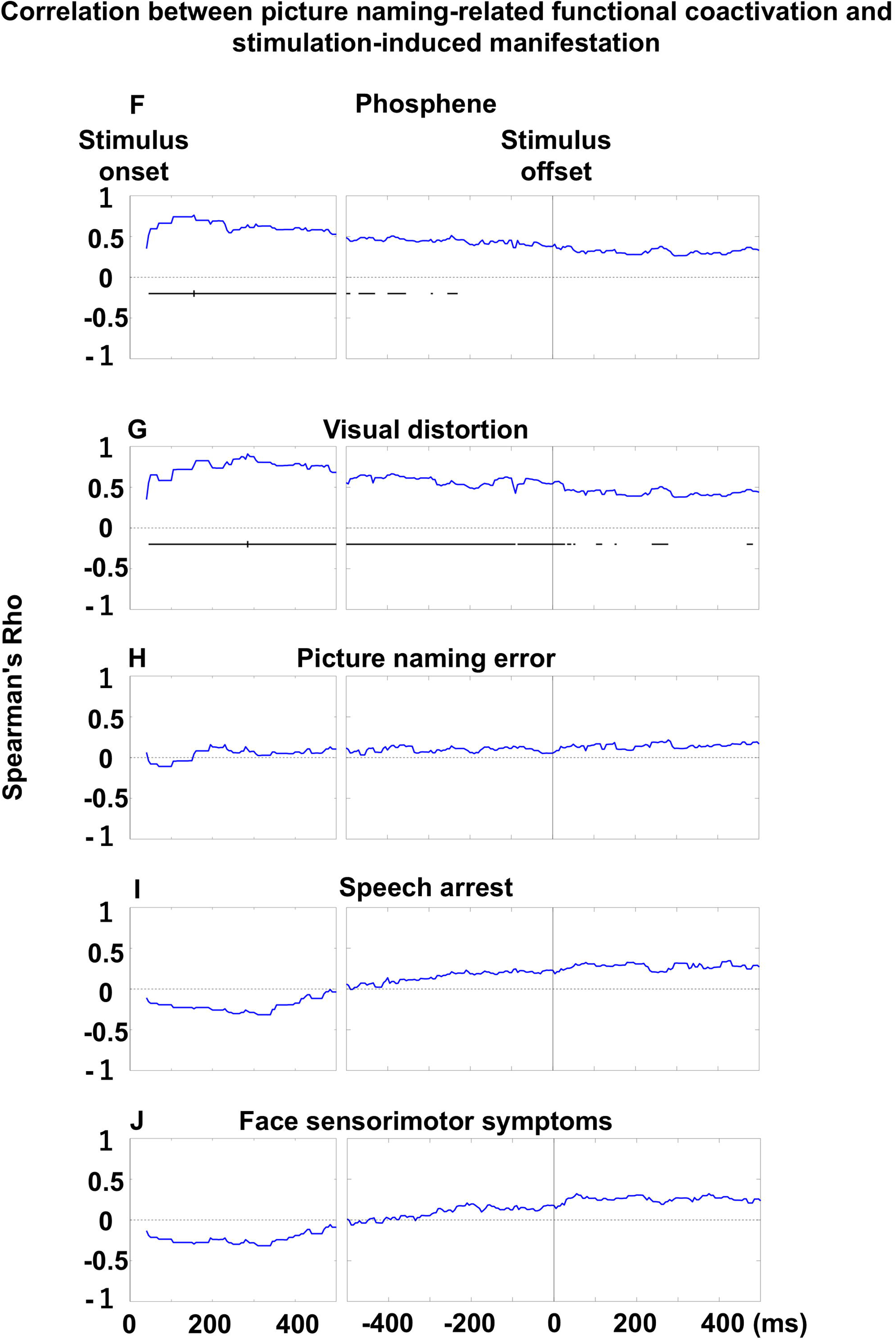
Association between functional coactivation and stimulation-induced manifestations. Each plot shows Spearman’s rho, reflecting the correlation between mean coactivation intensity and the probability of stimulation-induced symptoms across time bins. (A–E) Auditory naming–related correlations. (F–J) Picture naming–related correlations. Horizontal bars indicate significant correlations. Small vertical bars mark the time bin with peak rho.

- Auditory hallucination: 215 ms post-stimulus onset (rho: +0.64; p-value: 1.2×10)

- Receptive aphasia: 40 ms pre-stimulus offset (rho: +0.58; p-value: 4.8×10)

- Expressive aphasia: 130 ms post-stimulus offset (rho: +0.69; p-value: 2.9×10¹)

- Speech arrest: 60 ms pre-response onset (rho: +0.80; p-value: 1.4×10¹)

- Facial sensorimotor symptoms: 350 ms post-response onset (rho: +0.83; p-value: 6.6×10¹).

### Neural correlates of within-individual variability in response times in auditory naming

**Figure 5A** shows the temporal dynamics of high-gamma amplitude and functional coactivation at the following two ROIs, sorted by response times. Faster responses were associated with increased high-gamma and coactivation in the left IFG (pars triangularis) around stimulus offset. Significant negative correlations were found at 19 time bins for local high-gamma (minimum rho: –1.0; p-value: 0.003) and 60 bins for coactivation (minimum rho: –1.0; p-value: 0.003). In contrast, greater high-gamma attenuation in the right rostral MFG (38 bins) was linked to faster responses (maximum rho: +0.99; p-value: 0.006), while co-inactivation in this region was not significant. Among the 66 ROIs, the right rostral MFG was the only region where stronger suppression correlated with faster responses.

**Figure 5.**
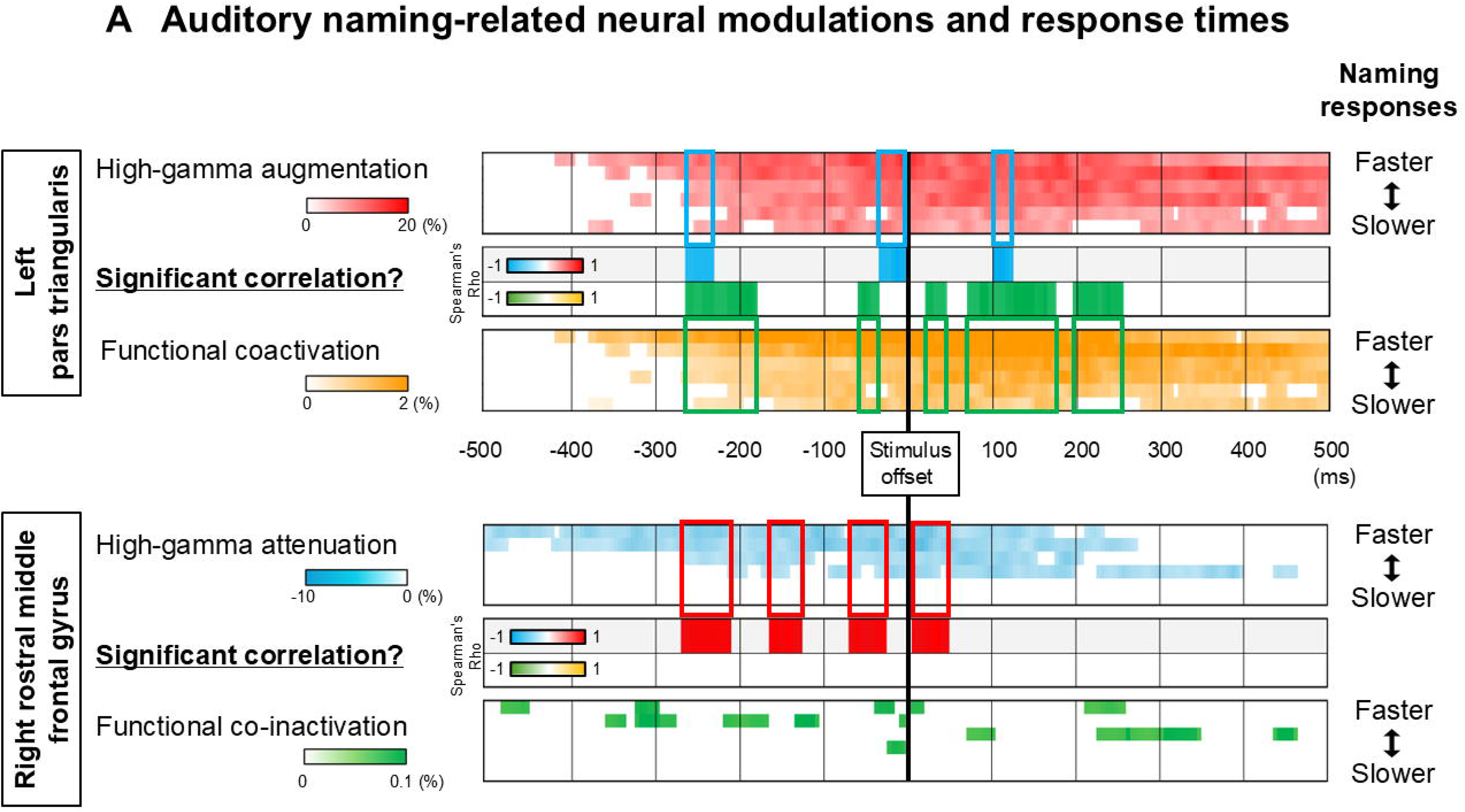
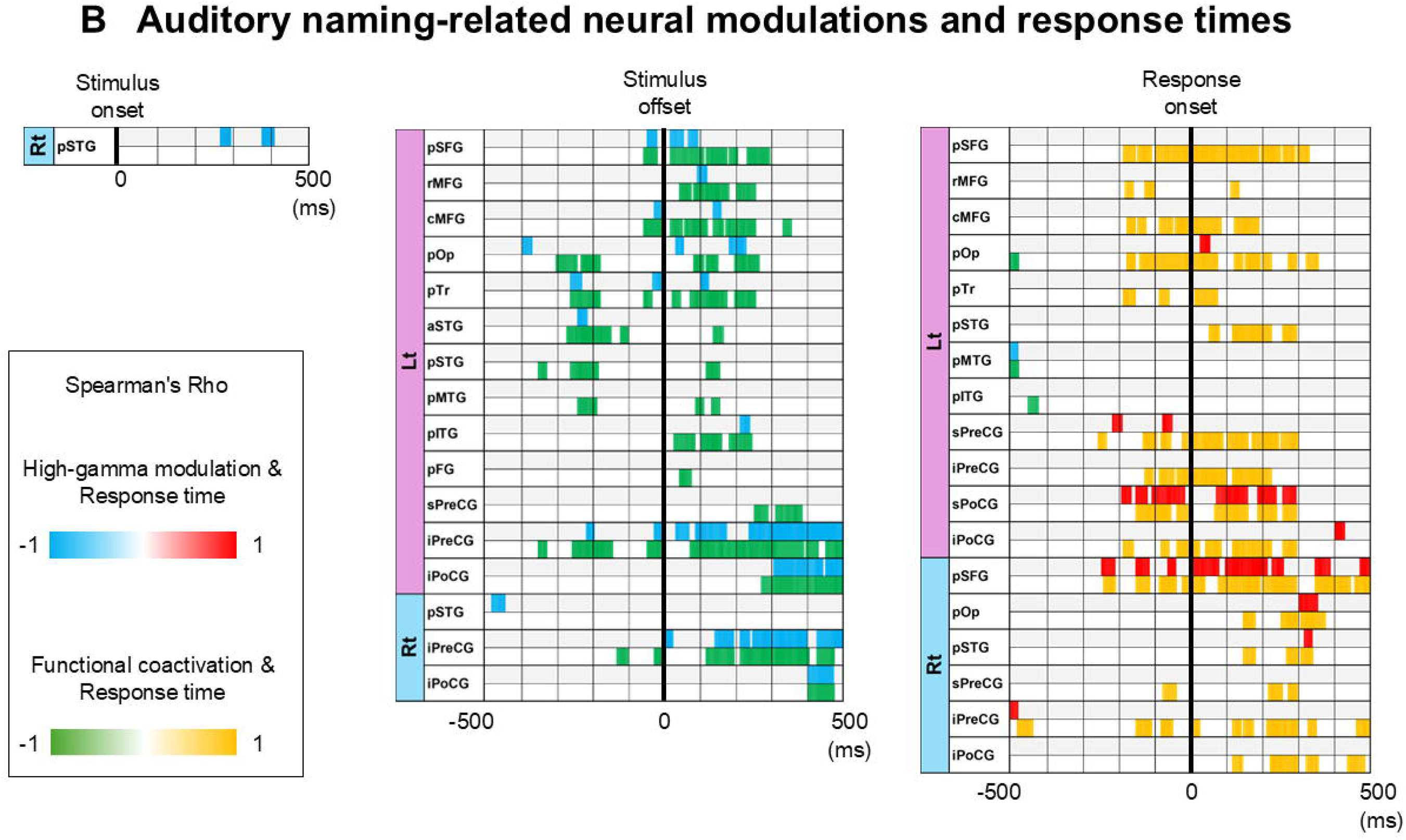
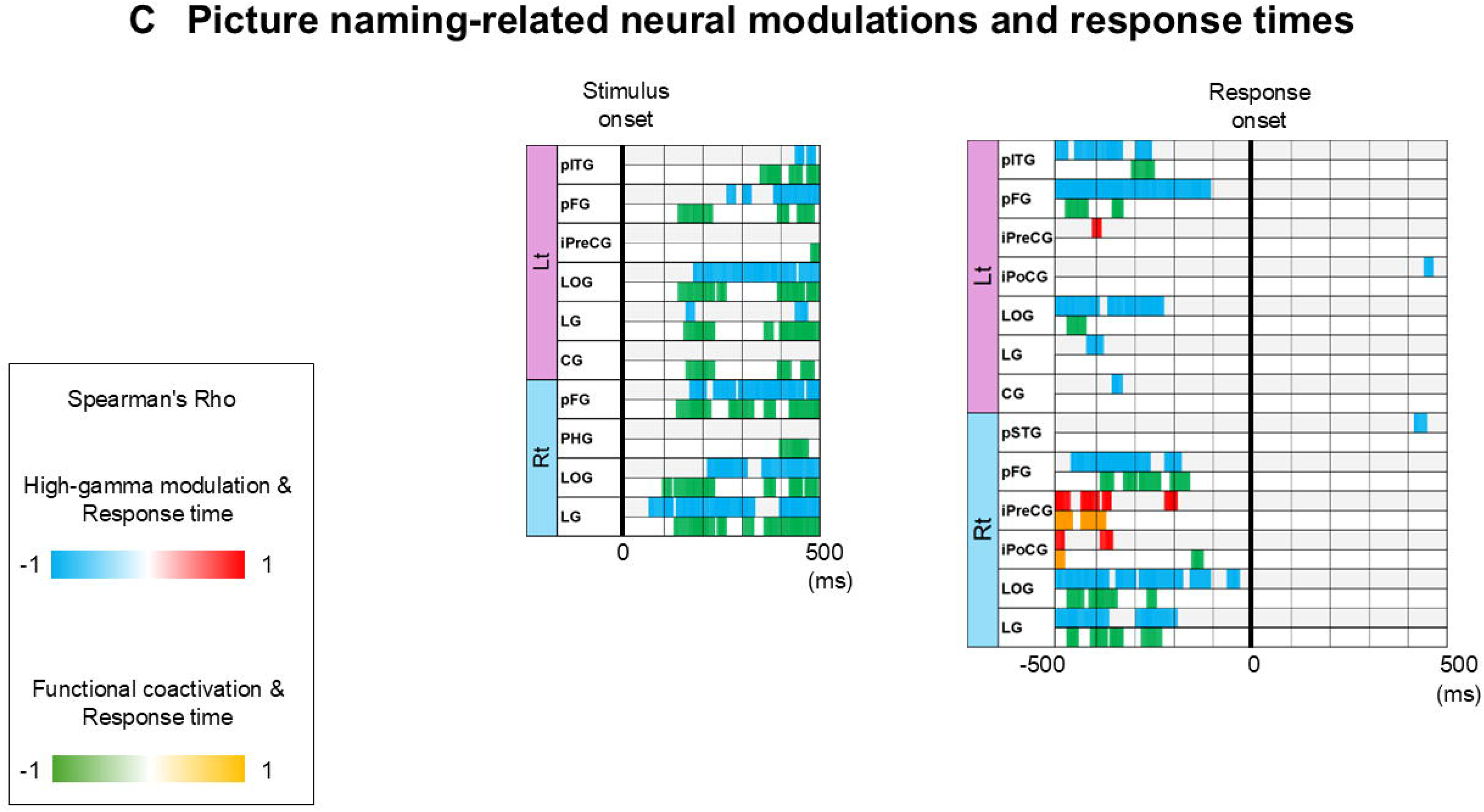
Neural activity and response times. (A) Temporal changes in high-gamma amplitude and coactivation intensity in left pars triangularis and right MFG, grouped by response speed. (B) Spearman’s rho over time showing correlations between neural measures and auditory naming response times. Blue/green: increased neural activity associated with faster responses; red/yellow: slower. (C) Same analysis for picture naming, showing only ROIs with significant correlations. Significance was defined as p <; 0.05 across ≥five consecutive 5-ms time bins. See **eFigure 10** for high-gamma amplitude at each of the 66 ROIs sorted by response time.

**Figure 5B** lists all ROIs with significant correlations between response time and high-gamma or coactivation. In the left hemisphere, functional coactivation around stimulus offset had a greater influence on response time than local high-gamma (median: 37.5 vs. 9.5 time bins; p-value: 0.009, Wilcoxon test). This finding suggests that coactivation intensity was more consistently associated with response time than local high-gamma amplitude. Faster responses were linked to stronger coactivation in left perisylvian regions—left IFG (pars opercularis/triangularis), inferior precentral gyrus, anterior/posterior STG, and posterior MTG—100–300 ms before stimulus offset (minimum rho: –1.0; p-value: 0.003). These perisylvian ROIs are connected via the arcuate fasciculus.

**Figure 5B** also highlights all ROIs where delayed responses were associated with increased high-gamma or coactivation, particularly in frontal areas—posterior SFG, inferior precentral gyrus (bilateral), and left MFG/IFG—200 ms before response onset (maximum rho: +1.0; p-value: 0.003). These frontal ROIs are linked via callosal fibers, the frontal aslant fasciculus, and U-fibers.

### Cortical high-gamma modulations and functional coactivation during picture naming

**eVideo 2** shows the dynamics of high-gamma activity and functional coactivation/co-inactivation during picture naming, with snapshots in **Figure 2**. After stimulus onset, high-gamma increased bilaterally in the medial and lateral occipital gyri, with interhemispheric coactivation via the corpus callosum. Within 300 ms, coactivation extended to posterior temporal regions via the inferior longitudinal fasciculi, while co-inactivation emerged between the left IFG and parietal lobe, and between bilateral IFG regions.

By 400 ms, intra-hemispheric coactivation spread from occipital to frontal areas via the inferior frontal-occipital, superior longitudinal, and arcuate fasciculi, mainly in the left hemisphere. As response onset neared, high-gamma increased in bilateral inferior Rolandic cortices, SFG, and the left IFG (pars opercularis), while occipital activation persisted. Coactivation persisted along the left inferior frontal-occipital, superior longitudinal, and arcuate fasciculi, but declined in the bilateral inferior longitudinal fasciculi. During the response, coactivation was observed between bilateral inferior Rolandic cortices, and between Rolandic, STG, and occipital regions within each hemisphere.

**eFigures 6 and 7** illustrate the proportion of specific white matter pathways that exhibited functional coactivation and co-inactivation. **eFigure 10** illustrates response time–sorted neural dynamics. We also display the temporal dynamics of high-gamma modulations with confidence interval bars (**eFigure 11**) and using a leave-one-out approach (**eFigure 12**).

### Picture naming-related functional coactivation and stimulation-induced manifestations

Correlations between functional coactivation and stimulation-induced manifestations peaked at distinct times:

- Phosphenes at 155 ms (rho: +0.76; p-value: 1.7×10¹³)

- Visual distortions at 285 ms post-stimulus (rho: +0.91; p-value: 1.6×10²)

- No significant correlations were found for picture naming errors, speech arrest, or facial sensorimotor symptoms (p-values>0.05).

### Neural correlates of within-individual variability in response times in picture naming

**Figure 5C** shows all ROIs where picture naming response times correlated with high-gamma or coactivation intensity. Faster responses were linked to greater coactivation in bilateral occipito-temporal regions—including lateral occipital, lingual, and posterior fusiform gyri—100–300 ms after stimulus onset (minimum rho: –1.0; p-value: 0.003). These regions are connected via the inferior longitudinal fasciculi and callosal fibers. Additional coactivation-response time correlations were found in left posterior ITG during 300–500 ms post-stimulus (minimum rho: –1.0; p-value: 0.003).

Conversely, slower responses were associated with increased high-gamma in bilateral inferior Rolandic cortices 200–500 ms before response onset (maximum rho: +0.94 to +1.0; p-values: 0.017–0.003).

## Discussion

### Summary

This study visualized the dynamic engagement of white matter networks during stages of auditory and picture naming. Our findings reconcile competing hypotheses about neural mechanisms by showing that rapid naming involves suppressed and enhanced neural interactions, depending on task modality. Faster auditory naming was linked to marked suppression of the right rostral MFG and increased coactivation along the left arcuate fasciculus around question offset. This suggests that reduced top-down inhibition from the right dorsolateral prefrontal cortex facilitates engagement of left-lateralized perisylvian pathways, expediting lexical retrieval. In contrast, delayed auditory naming showed a late surge in bilateral frontal coactivation, reflecting overload in bifrontal networks despite eventual success. Faster picture naming was associated with early, robust coactivation in bilateral occipitotemporal regions, implicating the inferior longitudinal fasciculi in rapid visual object recognition. White matter networks involving Broca’s area were suppressed within the first 300 ms following picture stimulus onset. These relationships between coactivation and naming speed enhance our understanding of delayed naming in neurological disorders. The strong correlation between coactivation intensity and stimulation-induced symptoms supports the clinical value of our dynamic causal tractography atlases for presurgical language mapping.

### Network dynamics underlying rapid auditory-naming responses

Faster auditory naming was associated with greater high-gamma suppression in the right rostral MFG and increased functional coactivation among left perisylvian regions connected by the arcuate fasciculus. Suppression in the right rostral MFG was most prominent before stimulus offset, while left perisylvian coactivation spanned before and after offset. These findings suggest that reduced domain-general inhibitory monitoring by the right dorsolateral prefrontal cortex may facilitate activation of left-hemispheric language networks linking posterior temporal and frontal regions. This aligns with prior iEEG studies showing increased high-gamma in the right rostral MFG after stop signals, errors, and during tasks requiring inhibitory control.^23–25^ Our recent work showed that elevated high-gamma in this region enhanced response accuracy but delayed responses in a non-verbal task.^6^ Repetitive transcranial magnetic stimulation near the right rostral MFG transiently disrupted inhibitory control in healthy adults.^26^

Compared to local high-gamma amplitudes, functional coactivation among left perisylvian cortices showed a more consistent association with auditory naming response times (**Figure 5B**). These cortices are connected via the arcuate fasciculus, and coactivation intensity correlated with stimulation-induced receptive and expressive aphasia. This suggests that coactivation in these regions reflects semantic analysis and lexical retrieval, which rely more on distributed network coordination than localized activation.^1^ Lesions to the left arcuate fasciculus cause more severe naming deficits than those limited to Broca’s area.^1^ Prior iEEG work demonstrates that the left posterior IFG and medial/lateral temporal cortices activate prior to motor regions during verbal tasks.^2,3^ A study of 36 epilepsy patients showed that high-gamma activity in the left IFG preceded that in the MTG during silent word reading.^3^ In the current study, functional coactivation involving the left posterior IFG showed more consistent associations with response time—spanning 250 ms in the pars opercularis and 300 ms in the pars triangularis—compared to 190 ms in the left posterior MTG (**Figure 5B**).

### Network dynamics underlying rapid picture-naming responses

Picture-naming response times were shaped by network dynamics in posterior regions. Faster responses were associated with stronger functional coactivation among bilateral occipital and posterior fusiform regions during 100–300 ms after stimulus onset (**Figure 8C**). These regions are connected via callosal fibers and the inferior longitudinal fasciculus. Coactivation during this time window also correlated with phosphenes and visual distortions elicited by stimulation, suggesting that enhanced visual attention and interhemispheric transfer facilitate rapid object recognition. Prior iEEG study demonstrated high-gamma augmentation in these regions during sustained visual attention.^27^ The causal role of posterior callosal fibers is supported by naming impairments following callosotomy, attributed to disrupted interhemispheric transfer.^28^ Such impairments are exacerbated when the posterior callosum is included in the disconnection.^29^

Previous research supports the causal role of the inferior longitudinal fasciculus and related cortices in visual naming. Resection of the left fusiform gyrus is linked to postoperative declines in picture naming,^30^ and lesions involving the left inferior longitudinal fasciculus are associated with naming deficits in brain tumor patients.^31^ The right inferior longitudinal fasciculus also contributes to object recognition.^32^ Patients with left-hemispheric lesions may still recognize objects and perform related actions (e.g., opening a refrigerator), suggesting that right-hemispheric visual networks support semantic control of everyday behaviors.^32^

Our analysis showed no evidence of top-down processing from Broca’s area during early visual object recognition. Within 300 ms after stimulus onset, functional co-inactivation involving the left pIFG was observed within and across hemispheres (**Figures 2A-2B**), with no significant correlation between pIFG activity and response times. Since Broca’s area is mainly involved in lexical retrieval and response selection,^33^ our findings suggest that early processing prioritizes object recognition via posterior networks over lexical engagement.

Enhanced functional coactivation in the left posterior ITG was associated with faster picture naming between 350–500 ms after stimulus onset (**Figure 2C**). During this period, coactivation via the left arcuate fasciculus linked the left inferior precentral gyrus with posterior basal temporal regions—including the posterior ITG and fusiform gyri—sites where stimulation often triggered naming errors (**Figure 3H**). Coactivation involving the left IFG pars opercularis and these regions appeared only around 405 ms before response onset (**eVideo 2**), suggesting a more limited role of Broca’s area in rapid picture naming compared to auditory naming.

### Network dynamics associated with delayed naming responses

Our study revealed a late surge in bilateral frontal coactivation immediately preceding auditory-naming responses, which was associated with delayed response times (**Figure 5B**). This finding should be interpreted with caution, as the surge more likely reflects a consequence rather than a cause of delayed responses. One possibility is that it represents subtle non-verbal cues—such as cautious initiation or hesitation.^34^ Our prior iEEG study using the Stroop task showed increased high-gamma activity and coactivation in bilateral posterior SFG, precentral gyri, left MFG, and posterior IFG^24^ which align with the regions showing late surges in the present study. Increased mouth or throat tension—often accompanying hesitation—is also expected to elevate high-gamma amplitude in the inferior Rolandic regions.^17^

Similarly, a late surge in bilateral inferior Rolandic activity was observed with delayed picture-naming responses (**Figure 5C**), though with more limited spatial extent—likely reflecting lower cognitive demand in picture naming compared to auditory naming.

### Functional coactivation dynamics in auditory versus picture naming

This study showed that distinct white matter fasciculi support specific naming stages—such as perception, semantic analysis, lexical retrieval, and articulation—without any single tract remaining active throughout. Instead, fasciculi were selectively engaged depending on the stage. Both auditory and picture naming shared early, symmetric coactivation in intra- and inter-hemispheric pathways involving low-order sensory cortices, supporting perceptual processing. Auditory naming uniquely activated the arcuate fasciculus and callosal fibers from the STG, with coactivation at 215 ms post-stimulus onset best correlating with stimulation-induced auditory hallucinations. This may partly reflect integration of speech timing and rhythm (left STG) with tone and pitch (right STG).^35,36^

In contrast, picture naming activated the inferior longitudinal fasciculus and callosal fibers from the occipital and fusiform gyri, with coactivation at 155 ms best correlating with phosphenes. This early interhemispheric coactivation likely supports transfer of perceptual representations between hemispheres.

Our analysis identified networks underlying cognitive functions. During auditory naming, functional coactivation increased around stimulus offset within major left-lateralized fasciculi—including the arcuate, inferior/middle/superior longitudinal, and inferior fronto-occipital fasciculi. Coactivation around stimulus offset correlated with stimulation-induced receptive and expressive aphasia, suggesting involvement in semantic analysis and lexical retrieval.

In contrast, picture naming showed more symmetric coactivation. At 285 ms post-stimulus, coactivation—especially via the left inferior longitudinal fasciculus—correlated with stimulation-induced visual distortion, reflecting higher-order visual processing. Faster picture naming was linked to increased coactivation between the left posterior fusiform/ITG and the inferior precentral gyrus via the arcuate fasciculus during 350–500 ms post-stimulus. Around 400 ms before response onset, coactivation extended to the left IFG (**eVideo 2**). Given the frequency of picture naming errors during stimulation of these areas (**Figure 3H**)^30^ and supporting lesion studies,^1^ coactivation involving these regions likely reflects lexical retrieval processes.

Just before response, functional coactivation decreased in left-hemispheric fasciculi involved in semantic analysis and visual recognition. Coactivation in the left arcuate fasciculus (auditory naming; **eFigure 6A**) and the left inferior longitudinal fasciculus (picture naming; **eFigure 6F**) declined prior to response. In contrast, coactivation increased in the left frontal aslant, superior longitudinal, and extreme capsule fasciculi—linking the precentral gyrus, SFG, and pars opercularis—in both tasks. Coactivation at 60 ms before auditory-naming response onset correlated most strongly with stimulation-induced speech arrest, suggesting these late dynamics reflect speech initiation networks.

### Utility of the atlas

This study emphasizes the utility of the dynamic causal tractography atlas in predicting cognitive impairments resulting from brain lesions. Damage to white matter pathways may result in more disabling deficits than cortical lesions alone.^1^ Understanding the trajectories connecting eloquent regions could improve outcome prediction and provide an educational resource for students and lay audiences learning how brain networks support cognition.

Our study showed that the left arcuate fasciculus, originating from the posterior STG and MTG, was engaged in lexical retrieval during auditory naming, whereas fibers originating from the posterior fusiform gyrus and ITG were involved in picture naming (**Figure 5; eVideos 1-2**). A study of 48 right-handed patients with left temporal gliomas found that infiltration of the inferior longitudinal fasciculus—originating in part from the fusiform gyrus—was strongly associated with picture-naming deficits, while verbal fluency and auditory naming remained intact.^37^ Similarly, a case of traumatic brain injury involving the left ILF, fusiform, and ITG showed visual naming impairment with preserved auditory naming.^38^ In contrast, stroke-induced damage to the left STG and its underlying white matter selectively impaired auditory comprehension but spared picture naming and reading.^39^ These findings suggest that lesions involving white matter networks from the left posterior STG and MTG predict auditory naming impairments, whereas those affecting the posterior fusiform gyrus and ITG predict visual naming deficits.

### Patient demographics and epilepsy characteristics

This study examined how age, handedness, and epilepsy-related factors influence behavior and neural dynamics. An increase in √age from 3 (9 years) to 4 (16 years) was associated with faster response times by 191 ms and 197 ms during auditory and picture naming, respectively. While we expected √age to correlate with increased high-gamma amplitude in the inferior precentral gyri after stimulus offset, mixed-model analysis revealed no significant √age-related neural effects.

Left-handedness was associated with greater auditory naming–related high-gamma amplitude in the right entorhinal cortex (300–400 ms post-offset), suggesting reduced left-hemispheric dominance during lexical retrieval. This finding aligns with prior fMRI studies reporting decreased left-lateralized language-related activation in left-handed individuals.^40^

Left-hemispheric epileptogenicity was associated with increased picture naming-related high-gamma amplitude in the right posterior STG immediately before response onset, suggesting a possible compensatory role of the right hemisphere in verbal response preparation. This aligns with functional MRI findings of increased right-hemispheric activation during language tasks in left-hemispheric epilepsy.^41^

### Limitations

Combining iEEG data from multiple patients into a unified virtual brain framework is a valid approach, analogous to a cross-sectional design when longitudinal studies are impractical. We previously demonstrated that this approach identifies functional coactivation and co-inactivation networks representative of a broader population, rather than chance findings or outliers.^7^ Since electrode placement is clinically guided, comprehensive bilateral sampling of homotopic nonepileptic regions is neither ethical nor feasible. Thus, individual-level iEEG cannot fully characterize interhemispheric interactions. Prior studies by other investigators addressed this limitation by integrating high-gamma activity across patients into a standardized atlas to infer region-to-region communication.^20,21^ A large sample size reduces individual variability, improved signal-to-noise ratios, and enhanced generalizability.

Moment-to-moment functional coactivation during auditory naming strongly correlated with the probability of stimulation-induced auditory hallucination, receptive aphasia, expressive aphasia, speech arrest, and face sensorimotor symptoms (**Figure 4**). In contrast, during picture naming, coactivation correlated only with phosphenes and visual distortion, not with naming errors, speech arrest, or sensorimotor symptoms. This discrepancy likely reflects task design differences. In auditory naming, patients heard speech only during stimulus presentation, with no auditory input during later stages. In picture naming, visual stimuli remained present until response onset, producing sustained high-gamma activity in bilateral occipital regions. However, electrical stimulation mapping did not identify these occipital areas as critical for naming or motor-related functions. Persistent coactivation in visual cortices likely accounts for the lack of correlation with picture naming error.

## Supporting information

eVideo_1

eVideo_2

## Abbreviations

CI: confidence interval
iEEG: intracranial
EEG IFG: inferior frontal gyrus
ITG: inferior temporal gyrus
MFG: middle frontal gyrus
MTG: middle temporal gyrus
ROIs: regions of interest
SFG: superior frontal gyrus
STG: superior temporal gyrus

## Acknowledgments

We are grateful to Sandeep Sood, MD, Alanna Carlson, MS, LLP, Jamie MacDougall, RN, BSN, CPN at Children’s Hospital of Michigan for the collaboration and assistance in performing the studies described above.

## Data availability

The data are available at https://openneuro.org/datasets/ds006234/versions/1.0.0 and https://openneuro.org/datasets/ds006233/versions/1.0.0.

## Code availability

The codes are available at https://github.com/rkwsu/Project_Auditory_Picture/tree/v1.0.

