## Supplementary_Document for "Dynamic Causal White Matter Atlas of Auditory and Visual Speech Networks at Millisecond Resolution: Intracranial Evidence from 125 Patients"

**in**

This document includes

**eTables 1-2** and **eFigures 1-12.**

| Patient | Age at surgery (years) | Sex | Handedness | MRI | Sampled hemisphere | Resected site | SOZ | Number of antiseizure medication | Median response time (seconds) | |
|---|---|---|---|---|---|---|---|---|---|---|
| | | | | | | | | | Auditory naming | Picture naming |
| 1 | 16 | M | Rt | Tumor | Lt | Lt | Lt F | 3 (LEV, OXC, CBZ) | 0.751 | NA |
| 2 | 17 | F | Rt | Nonlesion | Rt | Rt | Rt O | 2 (OXC, LEV) | 1.001 | NA |
| 3 | 15 | F | Rt | Tumor | Rt | Rt | NA | 1 (OXC) | 0.827 | NA |
| 4 | 8 | M | Rt | Nonlesion | Lt | Lt | Lt FP | 1 (OXC) | 1.353 | NA |
| 5 | 14 | F | Rt | Tumor | Lt | Lt | Lt T | 1 (LEV) | 1.742 | NA |
| 6 | 14 | M | Rt | Tumor | Lt | Lt | NA | 1 (OXC) | 0.668 | NA |
| 7 | 16 | F | Rt | Nonlesion | Lt and Rt | Rt | Rt FTPO | 3 (CZP, TPM, PHT) | 1.450 | 1.425 |
| 8 | 17 | F | Rt | Nonlesion | Lt and Rt | Lt | Lt F | 1 (LTG) | 0.883 | 1.067 |
| 9 | 10 | M | Rt | Dysplasia | Rt | Rt | Rt O | 3 (TPM, LEV, OXC) | 0.949 | NA |
| 10 | 8 | F | Rt | Nonlesion | Rt | Rt | NA | 2 (LTG, OXC) | 1.306 | NA |
| 11 | 8 | M | Rt | Tumor | Lt and Rt | Lt | NA | 1 (OXC) | 2.158 | 2.771 |
| 12 | 11 | F | Rt | Nonlesion | Rt | Rt | Rt T | 2 (OXC, LEV) | 1.339 | 1.798 |
| 13 | 16 | M | Rt | Nonlesion | Rt | Rt | Rt T | 1 (OXC) | 1.799 | NA |
| 14 | 18 | F | Rt | Dysplasia | Rt | Rt | Rt PO | 3 (ZNS, OXC, LTG) | 0.664 | NA |
| 15 | 17 | M | Rt | Tumor | Lt | Lt | Lt T | 1 (OXC) | 0.620 | 1.308 |
| 16 | 8 | M | Lt | Tumor | Lt | Lt | NA | 0 | 1.979 | NA |
| 17 | 17 | M | Rt | Nonlesion | Lt | Lt | NA | 2 (OXC, LEV) | 0.684 | 1.583 |
| 18 | 14 | M | Rt | Nonlesion | Lt | Lt | Lt T | 1 (VPA) | 2.285 | 2.694 |
| 19 | 10 | F | Rt | Tumor | Lt | Lt | Lt TP | 2 (OXC, TPM) | 0.719 | 2.120 |
| 20 | 10 | M | Lt | Nonlesion | Lt and Rt | NA | Lt FT, Rt F | 2 (VPA, CBZ) | 1.265 | 1.031 |
| 21 | 15 | M | Rt | Tumor | Lt | Lt | Lt T | 2 (LEV, LTG) | 1.155 | NA |
| 22 | 19 | F | Rt | Nonlesion | Lt and Rt | Rt | Rt P | 3 (OXC, LTG, LCM) | 0.762 | NA |
| 23 | 6 | F | Rt | Nonlesion | Lt and Rt | Rt | NA | 2 (CLB, PHT) | 1.459 | 1.441 |
| 24 | 14 | F | Rt | Nonlesion | Lt | Lt | Lt T | 3 (OXC, LTG, TPM) | 1.285 | 1.283 |
| 25 | 11 | M | Rt | Dysplasia | Rt | Rt* | Rt FP | 2 (OXC, LEV) | 0.808 | 1.237 |
| 26 | 13 | M | Rt | Tumor | Rt | Rt | Rt FP | 1 (OXC) | 0.627 | 0.927 |
| 27 | 23 | M | Rt | Nonlesion | Lt and Rt | Lt | Lt F | 2 (LCM, LEV) | 0.685 | 1.386 |
| 28 | 10 | M | Rt | Tumor | Rt | Rt | Rt T | 1 (OXC) | 1.220 | 1.467 |
| 29 | 5 | M | Rt | Nonlesion | Lt | Lt | Lt F | 3 (OXC, LEV, VPA) | 1.831 | 1.764 |
| 30 | 16 | F | Rt | Nonlesion | Lt and Rt | Lt | Lt PO | 2 (LEV, OXC) | 0.904 | 0.816 |
| 31 | 16 | M | Rt | Nonlesion | Rt | Rt | Rt TPO | 2 (LEV, OXC) | 0.983 | 1.327 |
| 32 | 37 | F | Rt | Tumor | Lt | Lt | NA | 1 (LEV) | 0.832 | 0.925 |

| | | | | | | | | | | |
|---|---|---|---|---|---|---|---|---|---|---|
| 33 | 14 | M | Rt | Nonlesion | Lt and Rt | Rt | Rt FP and Lt FP | 3 (LCM, OXC , VPA) | 1.127 | 1.935 |
| 34 | 5 | F | Rt | Others | Lt | Lt | Lt FT | 5 (ZNS, LEV, OXC , TPM, CLB) | 2.178 | 1.825 |
| 35 | 11 | F | Rt | Nonlesion | Rt | Rt | Rt FP | 2 (LEV, LTG) | 2.290 | 1.955 |
| 36 | 21 | F | Lt | Tumor | Lt | Lt | Lt O | 1 (LEV) | 0.446 | 1.067 |
| 37 | 17 | M | Rt | Dysplasia | Rt | Rt | Rt TPO | 2 (LCM, OXC) | 1.023 | 1.285 |
| 38 | 15 | F | Rt | Tumor | Lt | Lt | Lt T | 1 (LEV) | 1.301 | 1.227 |
| 39 | 44 | M | Ambidextrous | Dysplasia | Rt | Rt | Rt TP | 2 (OXC, LEV) | 1.121 | 1.157 |
| 40 | 37 | F | Rt | Dysplasia | Lt | Lt | Lt T | 2 (LTG, LCM) | 1.499 | 1.255 |
| 41 | 14 | F | Rt | Dysplasia | Lt | Lt | NA | 3 (LEV, OXC, LCM) | 1.099 | 1.042 |
| 42 | 28 | F | Rt | Nonlesion | Lt | Lt | Lt T | 2 (CBZ,LCM) | 2.563 | 3.064 |
| 43 | 20 | M | Rt | Nonlesion | Lt | Lt | Lt T | 2 (CBZ LCM) | 1.015 | 1.187 |
| 44 | 14 | F | Rt | Others | Lt and Rt | Rt | Rt T | 2 (LEV, TPM) | 1.425 | 1.188 |
| 45 | 13 | F | Rt | Nonlesion | Rt | Rt | NA | 3 (LEV, LTG, OXC) | 2.958 | 2.306 |
| 46 | 41 | F | Rt | Nonlesion | Lt | Lt | NA | 2 (LEV, PHT) | 0.736 | 2.087 |
| 47 | 12 | M | Rt | Nonlesion | Lt | Lt | Lt T | 3 (LCM, OXC, VPA) | 2.289 | 1.813 |
| 48 | 8 | M | Rt | Nonlesion | Lt and Rt | Rt | Rt F | 1 (LCM) | 1.329 | 2.272 |
| 49 | 10 | M | Lt | Dysplasia | Rt | Rt | Rt P | 3 (LCM, VPA, TRP) | NA | 3.280 |
| 50 | 10 | M | Rt | Others | Rt | Rt | NA | 1 (OXC) | 1.058 | 1.837 |
| 51 | 12 | M | Rt | Nonlesion | Rt | Rt | Rt T | 2 (VPA, LCM) | 1.870 | 1.728 |
| 52 | 9 | M | Rt | Others | Lt | Lt | Lt P | 2 (OXC, LTG) | 1.451 | 1.330 |
| 53 | 28 | M | Rt | Tumor | Rt | Rt | Rt T | 1 (CBZ) | 0.523 | 0.757 |
| 54 | 27 | F | Rt | Tumor | Lt and Rt | Lt | Lt F | 2 (LEV, LCM) | 1.023 | 1.025 |
| 55 | 17 | M | Rt | Others | Rt | Rt | Rt T | 2 (LEV, LCM) | 0.900 | 1.085 |
| 56 | 15 | F | Rt | Nonlesion | Lt and Rt | Rt | Rt T | 3 (LTG, LEV, ZNS) | 1.629 | 1.350 |
| 57 | 6 | F | Lt | Others | Lt and Rt | Lt | Lt T | 2 (VPA, LTG) | 1.321 | 1.454 |
| 58 | 12 | F | Rt | Others | Lt | Lt | Lt T | 2 (LTG, LCM) | 0.914 | 1.107 |
| 59 | 5 | F | Rt | Others | Lt | Lt | Lt T | 2 (LEV, LCM) | 1.996 | 2.516 |
| 60 | 9 | F | Rt | Tumor | Lt | Lt | Lt T | 1 (CBZ) | 1.508 | 1.313 |
| 61 | 30 | M | Rt | Nonlesion | Lt and Rt | Rt | Rt T | 3 (PHT, CBZ, LCM) | 1.360 | 1.052 |
| 62 | 21 | F | Rt | Nonlesion | Lt | Lt | Lt T | 2 (PHT, LEV) | 1.041 | 1.148 |
| 63 | 13 | M | Rt | Nonlesion | Rt | Rt | NA | 1 (LTG) | 0.718 | 2.710 |
| 64 | 12 | M | Rt | Dysplasia | Lt | Lt | Lt P | 2 (LEV, VPA) | NA | 1.099 |
| 65 | 11 | F | Rt | Tumor | Lt | Lt | Lt T | 2 (OXC, LEV) | 1.337 | 1.030 |
| 66 | 17 | M | Lt | Nonlesion | Rt | Rt | Rt O | 2 (LTG, LCM) | 0.544 | 0.763 |
| 67 | 16 | M | Rt | Nonlesion | Lt | Lt | Lt F | 2 (LTG, OXC) | 1.016 | 1.078 |
| 68 | 17 | F | Rt | Dysplasia | Lt | Lt | Lt T | 1 (LTG) | NA | 1.361 |

| | | | | | | | | | | |
|---|---|---|---|---|---|---|---|---|---|---|
| 69 | 8 | M | Rt | Tumor | Rt | Rt | Rt TO | 1 (OXC) | 2.273 | 2.508 |
| 70 | 13 | F | Rt | Nonlesion | Lt | Lt | Lt T | 1 (LEV) | 1.167 | 1.052 |
| 71 | 14 | F | Rt | Nonlesion | Rt | Rt | Rt P | 2 (LCM, LEV) | 1.164 | NA |
| 72 | 12 | M | Rt | Nonlesion | Lt | Lt | Lt T | 1 (VPA) | 2.217 | 2.050 |
| 73 | 13 | F | Rt | Dysplasia | Rt | Rt | Rt T | 4 (LEV, CLB, PHT , LTG) | 2.732 | 1.837 |
| 74 | 11 | F | Rt | Tumor | Lt | Lt | NA | 2 (OXC, LEV) | 1.384 | 1.124 |
| 75 | 15 | M | Rt | Nonlesion | Rt | Rt | Rt F | 2 (LCM, VPA) | 1.688 | 1.159 |
| 76 | 14 | F | Rt | Dysplasia | Lt | Lt | Lt T | 2 (OXC, VPA) | 3.067 | 1.498 |
| 77 | 11 | F | Rt | Nonlesion | Rt | Rt | Rt P | 1 (LEV) | 1.127 | 1.466 |
| 78 | 12 | F | Rt | Tumor | Rt | Rt | Rt P | 1 (LEV) | 1.152 | 1.170 |
| 79 | 17 | M | Lt | Nonlesion | Lt | Lt | Lt F | 2 (OXC, LTG) | 1.048 | 1.198 |
| 80 | 11 | F | Rt | Nonlesion | Rt | Rt | Rt F | 2 (OXC, LEV) | 1.860 | 1.578 |
| 81 | 17 | M | Rt | Nonlesion | Lt | Lt | Lt T | 3 (OXC, LEV, LCM) | 0.780 | 0.799 |
| 82 | 10 | F | Rt | Dysplasia | Rt | Rt | NA | 3 (LEV, LCM,ZNS) | 1.762 | 0.996 |
| 83 | 13 | M | Rt | Dysplasia | Rt | Rt | Rt TPO | 3 (LEV, LCM, PHT) | 2.359 | NA |
| 84 | 11 | M | Rt | Dysplasia | Rt | Rt | Rt P | 1 (LEV) | 1.964 | 3.123 |
| 85 | 16 | M | Rt | Dysplasia | Lt | Lt | Lt TO | 3 (OXC, LEV, LCM) | 1.833 | 1.383 |
| 86 | 15 | M | Rt | Nonlesion | Lt and Rt | Lt | Lt T | 1 (VPA) | 1.583 | 1.688 |
| 87 | 6 | M | Rt | Dysplasia | Lt | Lt | Lt F | 1 (CLB) | NA | 1.692 |
| 88 | 15 | M | Rt | Nonlesion | Rt | Lt | Lt T | 2 (VPA, LCM) | 1.129 | 2.076 |
| 89 | 10 | M | Rt | Nonlesion | Rt | Rt | Rt P | 1 (LCM) | 0.987 | 1.286 |
| 90 | 10 | M | Rt | Others | Rt | Rt | Rt T | 2 (OXC, LTG) | 1.068 | 0.761 |
| 91 | 16 | M | Rt | Nonlesion | Lt and Rt | NA | NA | 1 (OXC) | 0.724 | 1.181 |
| 92 | 12 | F | Rt | Dysplasia | Rt | Rt | Rt T | 2 (OXC, ZNS) | 1.487 | 1.426 |
| 93 | 8 | M | Rt | Tumor | Lt | Lt | Lt T | 2 (VPA, LEV) | 1.282 | 1.189 |
| 94 | 14 | F | Rt | Dysplasia | Rt | Rt | Rt T | 2 (OXC, LCM) | 2.191 | 1.487 |
| 95 | 12 | F | Rt | Tumor | Lt and Rt | NA | Lt F | 2 (LCM, LTG) | 1.292 | NA |
| 96 | 19 | M | Rt | Dysplasia | Rt | Rt | Rt O | 2 (LEV, OXC) | 1.635 | 1.594 |
| 97 | 6 | F | Rt | Nonlesion | Lt | Lt | Lt PO | 2 (OXC, LCM) | 1.332 | NA |
| 98 | 16 | F | Rt | Dysplasia | Lt | Lt | Lt TP | 1 (OXC) | 1.120 | 1.649 |
| 99 | 8 | M | Rt | Nonlesion | Rt | Rt | NA | 2 (LTG, LCM) | 1.807 | 1.633 |
| 100 | 5 | F | Rt | Nonlesion | Lt | Lt | Lt TPO | 3 (OXC, LEV, CLB) | NA | 4.073 |
| 101 | 16 | M | Rt | Nonlesion | Rt | Rt | Rt F | 1 (OXC) | 2.267 | 2.307 |
| 102 | 19 | F | Rt | Nonlesion | Lt and Rt | NA | Lt F | 1 (OXC) | 1.919 | 1.476 |
| 103 | 15 | F | Rt | Nonlesion | Rt | Rt | Rt P | 2 (ZNS, LTG) | 1.497 | 1.260 |
| 104 | 14 | F | Rt | Tumor | Rt | Rt | Rt I | 2 (LCM, LTG) | 1.729 | 1.210 |
| 105 | 5 | F | Lt | Dysplasia | Lt and Rt | Rt | Rt F | 1 (LEV) | 2.338 | 2.016 |

| | | | | | | | | | | |
|---|---|---|---|---|---|---|---|---|---|---|
| 106 | 16 | F | Rt | Nonlesion | Rt | NA | NA | 2 (TPM, CLB) | 1.363 | 1.207 |
| 107 | 13 | M | Rt | Dysplasia | Lt | Lt | Lt O | 3 (LEV, LCM, OXC) | 1.563 | 0.761 |
| 108 | 16 | F | Rt | Nonlesion | Lt and Rt | Rt | Rt F | 2 (OXC, LTG) | 0.915 | 1.815 |
| 109 | 9 | M | Rt | Tumor | Lt | Lt | Lt T | 1 (OXC) | 2.304 | 1.272 |
| 110 | 13 | M | Rt | Nonlesion | Lt and Rt | Rt | Rt F | 2 (OXC, LEV) | 1.078 | 0.861 |
| 111 | 11 | M | Rt | Dysplasia | Lt | Lt | Lt TO | 5 (OXC, CLB, LCM , VPA, PER) | 2.339 | 1.343 |
| 112 | 13 | F | Rt | Nonlesion | Rt | Rt | Rt TO | 3 (CBZ, OXC, ESL) | 2.279 | 1.282 |
| 113 | 20 | M | Rt | Nonlesion | Rt | Rt | Rt F | 2 (LCM, ESL) | 1.332 | 1.140 |
| 114 | 49 | F | Rt | Tumor | Lt and Rt | Lt | Lt TP | 2 (LEV, CLB) | 1.435 | 1.385 |
| 115 | 12 | F | Rt | Dysplasia | Lt and Rt | Lt | Lt F | 2 (LTG, LEV) | 1.283 | 1.017 |
| 116 | 15 | F | Rt | Nonlesion | Lt and Rt | Rt | Rt TPO and Lt TO | 2 (OXC, TPM) | 1.577 | 1.951 |
| 117 | 8 | F | Rt | Dysplasia | Lt and Rt | Rt | Rt P | 3 (OXC, CLB, LCM) | 2.196 | 0.909 |
| 118 | 16 | M | Rt | Tumor | Lt | Lt | NA | 1 (LEV) | 2.867 | 2.420 |
| 119 | 17 | M | Lt | Tumor | Lt | Lt | NA | 3 (LEV, OXC, CLB) | 1.384 | 1.036 |
| 120 | 13 | M | Rt | Nonlesion | Lt and Rt | Rt | Rt F | 3 (DZP, VPA, OXC) | 1.994 | 1.282 |
| 121 | 7 | F | Rt | Tumor | Rt | Rt | NA | 2 (LEV, CLB) | 2.195 | 1.514 |
| 122 | 6 | M | Rt | Dysplasia | Lt and Rt | Lt | Lt F | 2 (LEV, CLB) | NA | 2.626 |
| 123 | 17 | M | Rt | Nonlesion | Lt | Lt** | NA | 2 (LEV, OXC) | 1.162 | 0.859 |
| 124 | 19 | M | Rt | Others | Lt | Lt | Lt T | 3 (CLB, OXC, ZNS) | 2.667 | 1.425 |
| 125 | 15 | F | Rt | Others | Lt and Rt | Rt | Rt TPO | 2 (LEV, CLB) | 1.767 | 1.703 |

**eTable 1. Patient profiles.** Sixty-one females (F), and 64 males (M) were included in the study. Lt: left. Rt: right. SOZ: seizure onset zone. NA: not available. MRI lesions other than dysplasia and tumor included focal cortical atrophy (patient 34, 52, 125), hippocampal sclerosis (patients 44, 57, 58, 59, 90, 124), arachnoid cyst (patient 50), and arteriovenous malformations (patient 55). *: multiple subpial transections on the right Rolandic areas. **: responsive neurostimulation employed to the left temporal region. F: Frontal. T: Temporal. O: Occipital. P: Parietal. I: insula. CBZ: Carbamazepine. CLB: Clobazam. CZP: Clonazepam. DZP: Diazepam. ESL: Eslicarbazepine. LCM: Lacosamide. LEV: Levetiracetam. LTG: Lamotrigine. OXC: Oxcarbazepine. PER: Perampanel. PHT: Phenytoin. TPM: Topiramate. VPA: Valproic acid. ZNS: Zonisamide.

| Auditory naming | | | Picture naming | | |
|---|---|---|---|---|---|
| **Regions of interest** | **Left** | **Right** | **Regions of interest** | **Left** | **Right** |
| aSFG: anterior superior frontal gyrus | 82 (19) | 143 (35) | aSFG: anterior superior frontal gyrus | 91 (21) | 136 (32) |
| pSFG: posterior superior frontal gyrus | 79 (23) | 99 (35) | pSFG: posterior superior frontal gyrus | 85 (25) | 88 (30) |
| rMFG: rostral middle frontal gyrus | 254 (47) | 339 (51) | rMFG: rostral middle frontal gyrus | 259 (43) | 296 (46) |
| cMFG: caudal middle frontal gyrus | 229 (51) | 259 (54) | cMFG: caudal middle frontal gyrus | 225 (47) | 229 (48) |
| pOp: pars opercularis | 158 (47) | 105 (44) | pOp: pars opercularis | 152 (43) | 94 (36) |
| pTr: pars triangularis | 110 (38) | 177 (43) | pTr: pars triangularis | 108 (41) | 154 (37) |
| pOrb: pars orbitalis | 83 (32) | 83 (26) | pOrb: pars orbitalis | 82 (31) | 72 (24) |
| LOrbF: lateral orbitofrontal gyrus | 120 (38) | 100 (33) | LOrbF: lateral orbitofrontal gyrus | 116 (36) | 97 (30) |
| MOrbF: medial orbitofrontal gyrus | 27 (19) | 40 (24) | MOrbF: medial orbitofrontal gyrus | 28 (20) | 39 (24) |
| aCG: anterior cingulate gyrus | 13 (8) | 18 (9) | aCG: anterior cingulate gyrus | 13 (8) | 17 (9) |
| IG: insular gyrus | 4 (2) | 1 (1) | IG: insular gyrus | 6 (3) | 3 (2) |
| aSTG: anterior superior temporal gyrus | 162 (47) | 224 (50) | aSTG: anterior superior temporal gyrus | 172 (46) | 185 (42) |
| pSTG: posterior superior temporal gyrus | 264 (53) | 207 (49) | pSTG: posterior superior temporal gyrus | 247 (49) | 158 (40) |
| aMTG: anterior middle temporal gyrus | 87 (31) | 89 (32) | aMTG: anterior middle temporal gyrus | 97 (31) | 85 (32) |
| pMTG: posterior middle temporal gyrus | 258 (50) | 131 (39) | pMTG: posterior middle temporal gyrus | 231 (46) | 88 (31) |
| aITG: anterior inferior temporal gyrus | 65 (26) | 99 (35) | aITG: anterior inferior temporal gyrus | 65 (25) | 81 (30) |
| pITG: posterior inferior temporal gyrus | 168 (52) | 103 (40) | pITG: posterior inferior temporal gyrus | 174 (49) | 89 (36) |
| aFG: anterior fusiform gyrus | 44 (24) | 76 (36) | aFG: anterior fusiform gyrus | 47 (24) | 63 (29) |
| pFG: posterior fusiform gyrus | 198 (47) | 159 (46) | pFG: posterior fusiform gyrus | 201 (47) | 150 (41) |
| EG: entorhinal gyrus | 46 (22) | 69 (28) | EG: entorhinal gyrus | 47 (23) | 60 (23) |
| PHG: parahippocampal gyrus | 35 (18) | 27 (20) | PHG: parahippocampal gyrus | 34 (17) | 24 (17) |
| sPreCG: superior precentral gyrus | 110 (39) | 127 (42) | sPreCG: superior precentral gyrus | 112 (41) | 105 (38) |
| iPreCG: inferior precentral gyrus | 358 (58) | 404 (57) | iPreCG: inferior precentral gyrus | 363 (56) | 359 (49) |
| sPoCG: superior postcentral gyrus | 111 (40) | 89 (40) | sPoCG: superior postcentral gyrus | 105 (37) | 86 (34) |
| iPoCG: inferior postcentral gyrus | 323 (58) | 320 (54) | iPoCG: inferior postcentral gyrus | 330 (55) | 281 (48) |
| PCL: paracentral lobule | 28 (14) | 65 (24) | PCL: paracentral lobule | 38 (17) | 54 (21) |
| SPG: superior parietal gyrus | 69 (19) | 57 (19) | SPG: superior parietal gyrus | 65 (18) | 33 (14) |
| IPG: inferior parietal gyrus | 94 (25) | 160 (37) | IPG: inferior parietal gyrus | 89 (25) | 127 (33) |
| SMG: supramarginal gyrus | 352 (55) | 339 (51) | SMG: supramarginal gyrus | 342 (52) | 290 (46) |
| LOG: lateral occipital gyrus | 229 (41) | 217 (42) | LOG: lateral occipital gyrus | 244 (41) | 205 (38) |
| LG: lingual gyrus | 122 (34) | 168 (45) | LG: lingual gyrus | 132 (36) | 157 (41) |
| CG: cuneus gyrus | 23 (12) | 39 (21) | CG: cuneus gyrus | 26 (15) | 35 (18) |
| Pcun: precuneus gyrus | 52 (23) | 72 (28) | Pcun: precuneus gyrus | 49 (24) | 63 (24) |
| pCG: posterior cingulate gyrus | 64 (21) | 83 (27) | pCG: posterior cingulate gyrus | 64 (22) | 73 (25) |
| Total | 4421 (72) | 4688 (71) | Total | 4439 (69) | 4076 (62) |

**eTable 2. Number of artifact-free, nonepileptic electrode sites in regions of interest (ROIs).** Parenthetical values represent the number of patients who contributed data. IG was not included in the ROI-based analysis because the number of contributing patients was fewer than three in a hemisphere.

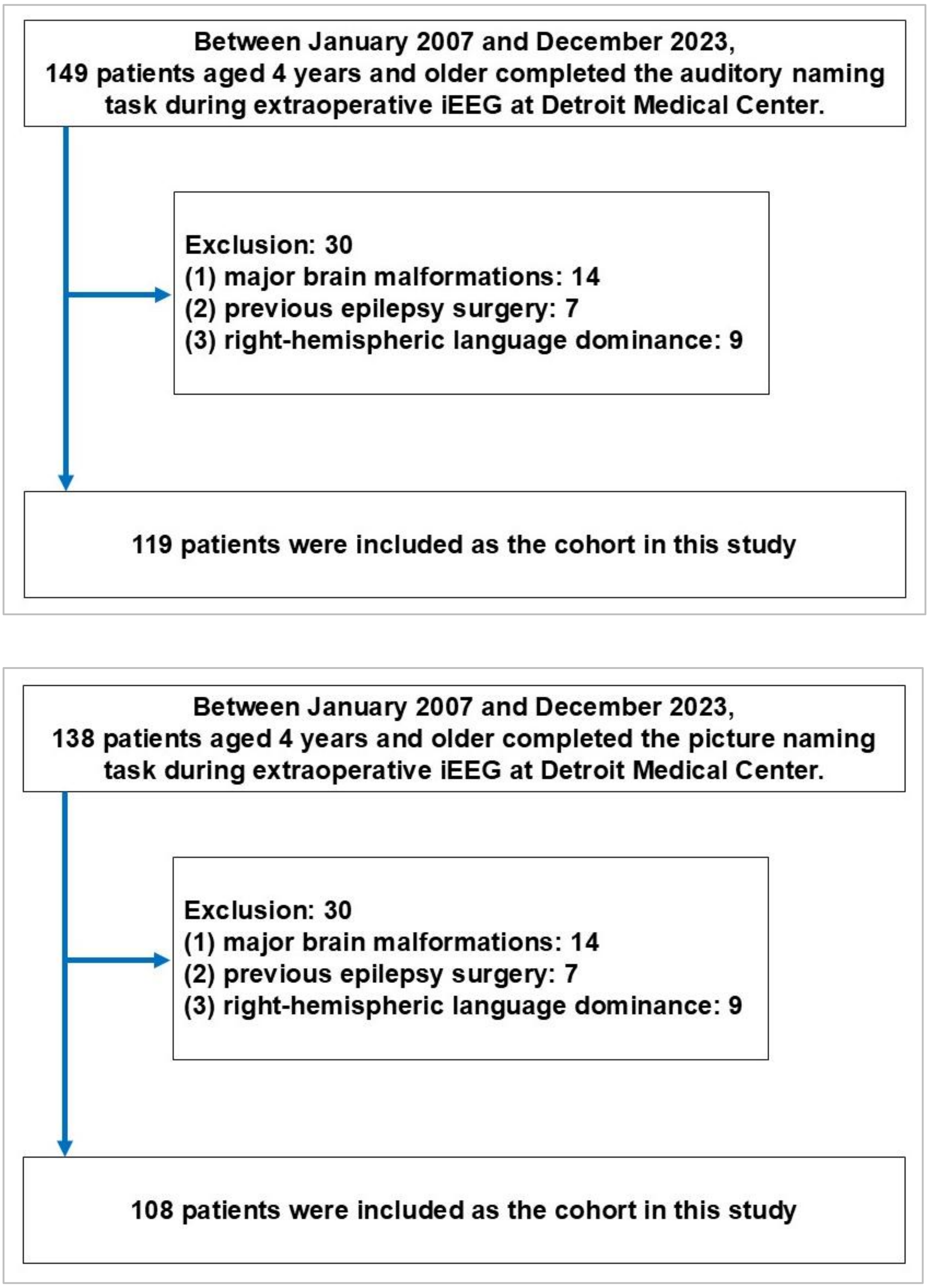


**eFigure 1. Flowcharts illustrating the selection of study patients who met eligibility criteria.**

A total of 149 native English-speaking patients aged ≥4 years completed the auditory naming task during extraoperative intracranial EEG recording. Of these, 14 were excluded due to major brain malformations, 7 due to a history of prior epilepsy surgery, and 9 due to right-hemispheric language dominance. Consequently, 119 patients were included in the present study.

Similarly, 138 patients completed the picture naming task. Among them, 14 were excluded due to major brain malformations, 7 due to a history of prior epilepsy surgery, and 9 due to right-hemispheric language dominance. As a result, 108 patients were included in the present study.

A total of 125 patients were included in the present study; 102 patients completed both auditory and picture naming tasks; 17 completed the auditory naming task alone; 6 completed the picture naming task alone.

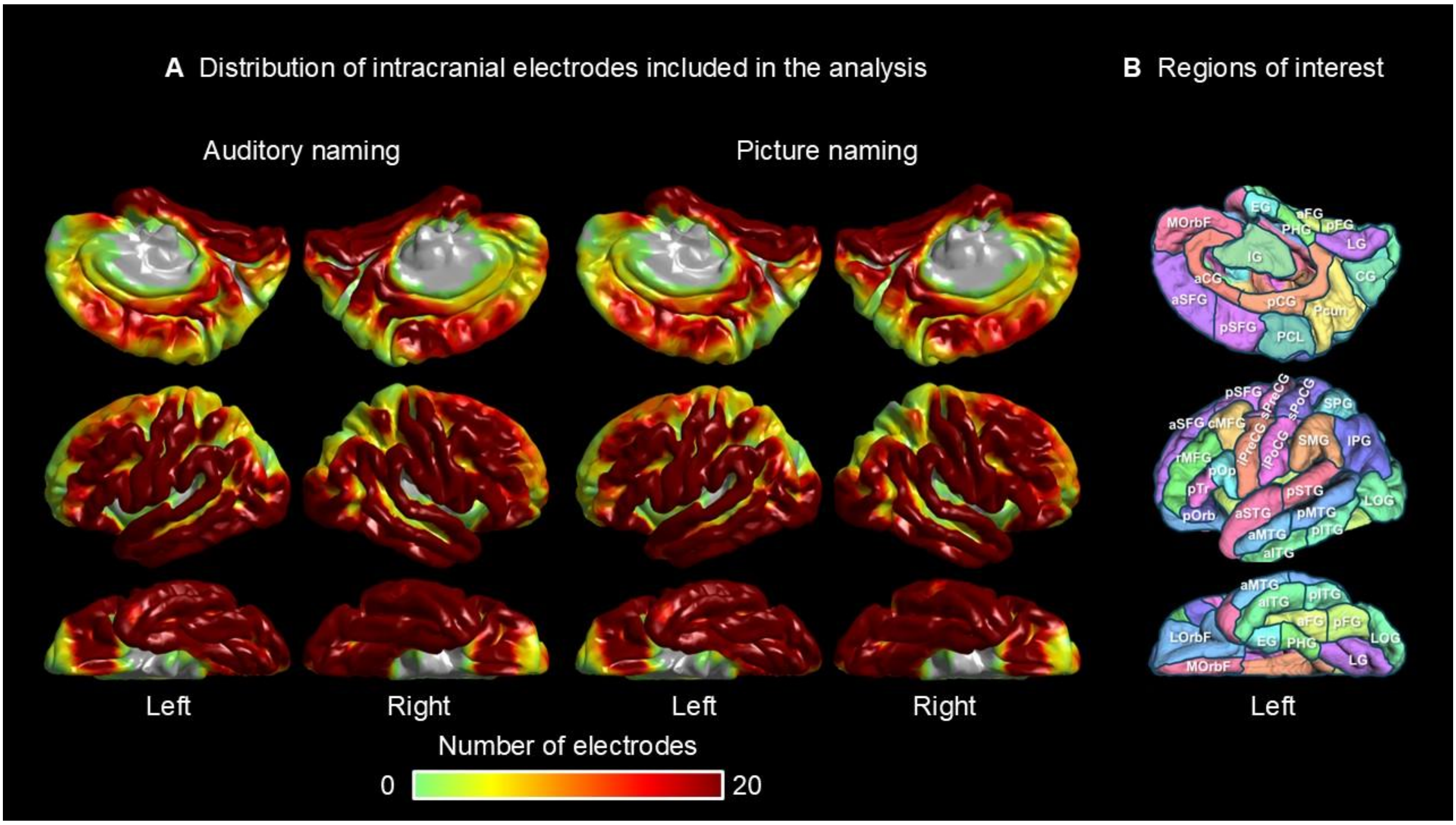


**eFigure 2. Intracranial electrode coverage.** (A) Number of artifact-free, non-epileptic sites analyzed for high-gamma activity during auditory (left) and picture naming (right). (B) Left hemisphere ROIs based on the Desikan-Killiany atlas[S1]; full names listed in **eTable 2**. Compared to our previous iEEG study,[S2] we increased the number of ROIs by subdividing temporal regions into anterior/posterior and Rolandic regions into superior/inferior portions, enabling finer assessment of neural engagement and coactivation. For example, during auditory naming, posterior STG and MTG showed earlier high-gamma augmentation and coactivation than anterior portions (**eFigures 8A** and **8G**), and the inferior left precentral gyrus activated earlier and more strongly than its superior counterpart (**eFigures 8B** and **8H**). In picture naming, posterior fusiform and ITG were engaged earlier than anterior regions (**eFigures 8M** and **8Q**). However, given variability in ROI sizes, comparisons of coactivation intensity should be limited to different time windows within the same fasciculus, as cross-fasciculus comparisons may be misleading.

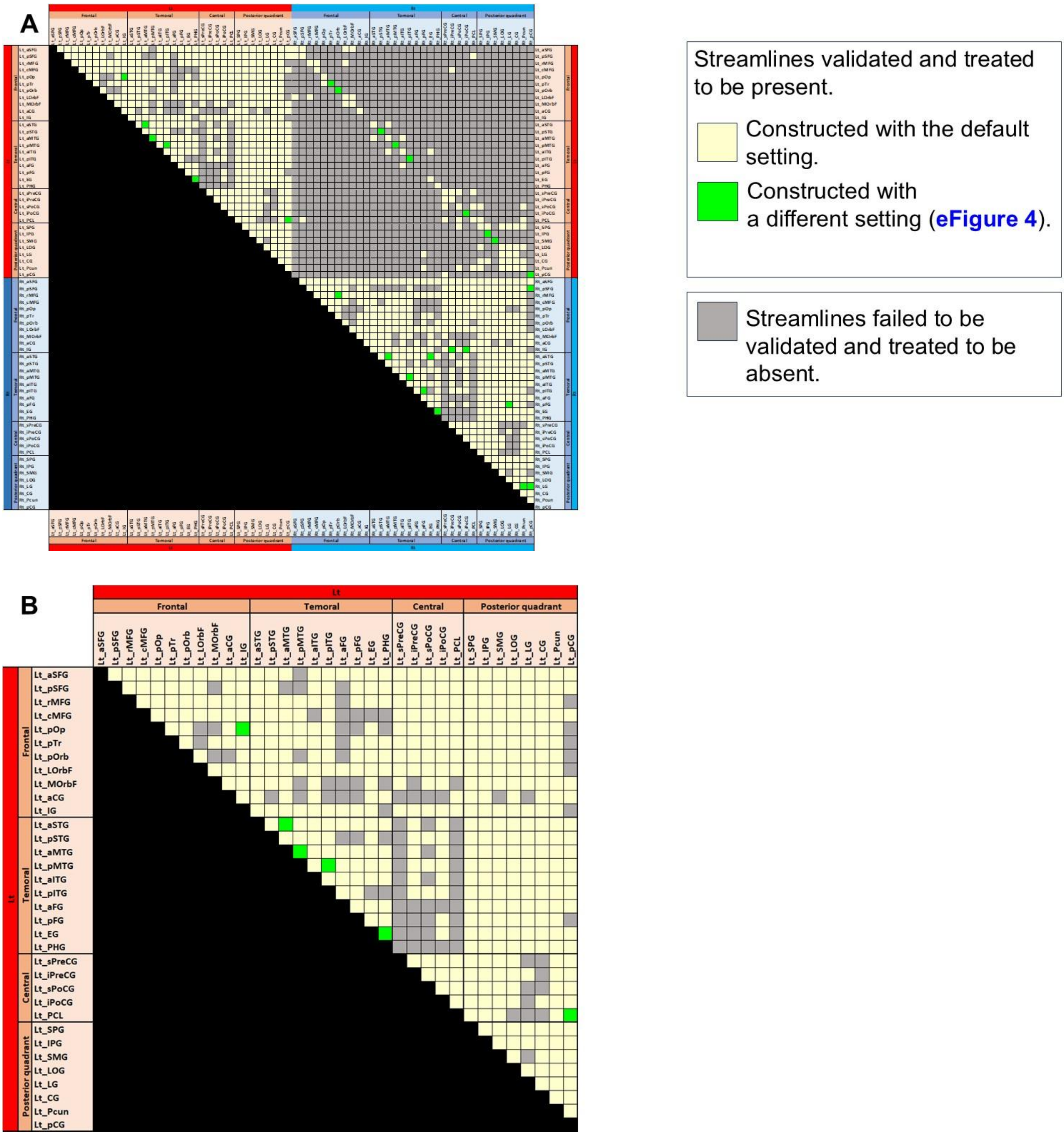
A
Streamlines validated and treated to be present.
Constructed with the default setting.
Constructed with a different setting (eFigure 4).
Streamlines failed to be validated and treated to be absent.
B
Lt
Frontal
Temoral
Central
Posterior quadrant
Lt_aSFG
Lt_pSFG
Lt_rMFG
Lt_cMFG
Lt_pOp
Lt_pTr
Lt_pOrb
Lt_LOrbF
Lt_MOrbF
Lt_aCG
Lt_IG
Lt_aSTG
Lt_pSTG
Lt_aMTG
Lt_pMTG
Lt_aITG
Lt_pITG
Lt_aFG
Lt_pFG
Lt_EG
Lt_PHG
Lt_sPreCG
Lt_iPreCG
Lt_sPoCG
Lt_iPoCG
Lt_PCL
Lt_SPG
Lt_IPG
Lt_SMG
Lt_LOG
Lt_LG
Lt_CG
Lt_Pcun
Lt_pCG

eFigure 3

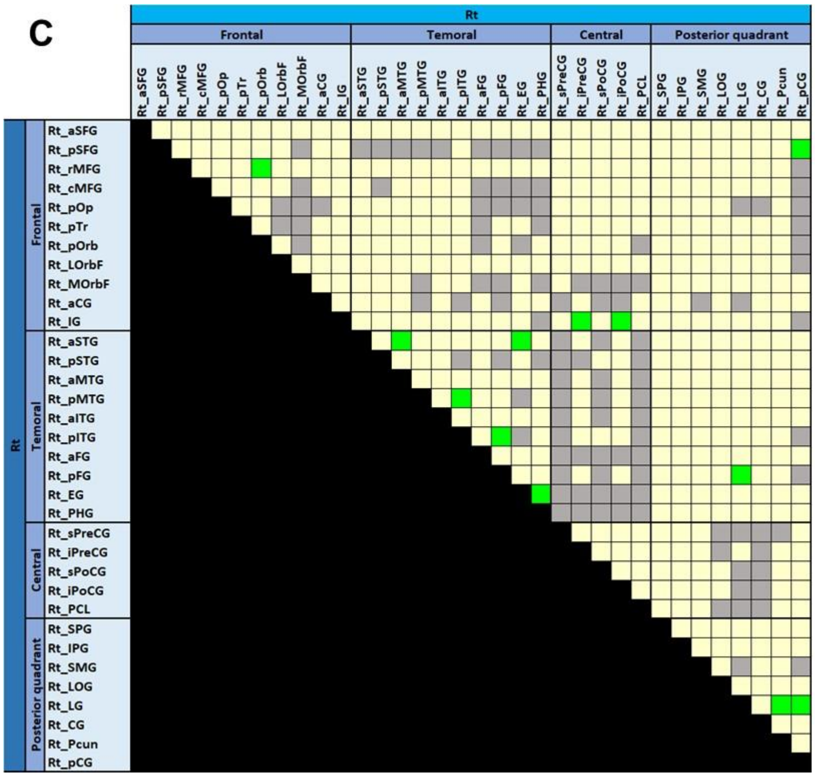

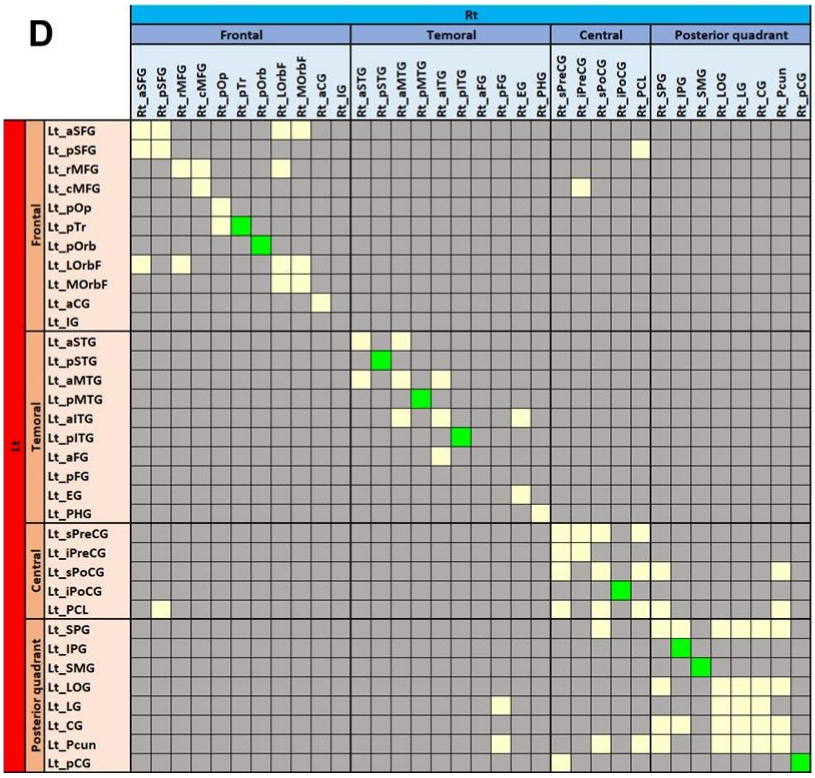

**eFigure 3. Construction and visualization of structural connectivity using white matter tractography.** **A** Structural connectivity matrix used to examine functional connectivity modulations. **B** Magnified view of intra-hemispheric white matter streamlines in the left hemisphere. **C** Magnified view of intra-hemispheric streamlines in the right hemisphere. **D** Magnified view of inter-hemispheric streamlines. Cream- and light green–colored squares indicate pairs of regions of interest (ROIs) for which white matter streamlines were successfully identified. Specifically, the cream-colored entries represent streamlines generated using the default parameter set, while the light green entries denote streamlines reconstructed using different parameters (see **eFigure 4**). Gray squares mark ROI pairs without validated streamlines. A list of the abbreviated cortical ROIs is provided in **eTable 2**. The complete structural

connectivity template is openly available (https://github.com/rkwsu/Project_Auditory_Picture/tree/v1.0). Using DSI Studio (http://dsi-studio.labsolver.org/) within the Montreal Neurological Institute (MNI) standard space, we identified and visualized the shortest among all detected streamlines for each ROI pair, applying default parameters of a 0.05 quantitative anisotropy (QA) threshold, a maximum turning angle of 70°, and a permissible tract length of 20–250 mm. Board-certified neurosurgeons (R.K. and A.K.) then validated each reconstructed streamline, excluding artifactual trajectories and those extending into the brainstem, basal ganglia, or thalamus. Artifacts were defined as excessively looping streamlines, paths terminating in improbable locations, or other irregular patterns.[S3–S8] Whenever local U-fibers between directly adjacent ROIs or callosal fibers connecting precisely homotopic ROIs were not recovered using the default parameters, an alternative parameter set was employed. **eFigure 4** documents the distinct settings used to generate each validated streamline. In line with prior evidence that demonstrates direct connectivity between immediately neighboring ROIs[S9,S10] and between homotopic cortical areas[S11–S26], our goal was to comprehensively visualize all plausible white matter pathways. This approach was also informed by previous findings[S8] indicating that standard parameters may fail to reconstruct some legitimate local U-fibers and inter-hemispheric fibers.

S16. Caminiti R, Carducci F, Piervincenzi C, et al. Diameter, length, speed, and conduction delay of callosal axons in macaque monkeys and humans: comparing data from histology and magnetic resonance imaging diffusion

tractography. J Neurosci. 2013;33:14501-14511.
S17. O'Reilly JX, Croxson PL, Jbabdi S, et al. Causal effect of disconnection lesions on interhemispheric functional connectivity in rhesus monkeys. Proc Natl Acad Sci U S A. 2013;110:13982-13987.
S18. Fenlon LR, Richards LJ. Contralateral targeting of the corpus callosum in normal and pathological brain function. Trends Neurosci. 2015;38:264-272.
S19. Horowitz A, Barazany D, Tavor I, Bernstein M, Yovel G, Assaf Y. In vivo correlation between axon diameter and conduction velocity in the human brain. Brain Struct Funct. 2015;220:1777-1788.
S20. Lacuey N, Zonjy B, Kahriman ES, Kaffashi F, Miller J, Lüders HO. Functional connectivity between right and left mesial temporal structures. Brain Struct Funct. 2015;220:2617-2623.
S21. Lacuey N, Zonjy B, Kahriman ES, et al. Homotopic reciprocal functional connectivity between anterior human insulae. Brain Struct Funct. 2016;221:2695-2701.
S22. Ruddy KL, Leemans A, Carson RG. Transcallosal connectivity of the human cortical motor network. Brain Struct Funct. 2017;222:1243-1252.
S23. Lehner KR, Yeagle EM, Argyelan M, et al. Validation of corpus callosotomy after laser interstitial thermal therapy: a multimodal approach. J Neurosurg. 2018;131:1095-1105.
S24. Mancuso L, Uddin LQ, Nani A, Costa T, Cauda F. Brain functional connectivity in individuals with callosotomy and agenesis of the corpus callosum: A systematic review. Neurosci Biobehav Rev. 2019;105:231-248.
S25. Mitsuhashi T, Sonoda M, Jeong JW, et al. Four-dimensional tractography animates propagations of neural activation via distinct interhemispheric pathways. Clin Neurophysiol. 2021;132:520-529.
S26. Innocenti GM, Schmidt K, Milleret C, et al. The functional characterization of callosal connections. Prog Neurobiol. 2022;208:102186.

| Parameter setting | ROI | ROI | Tracking threshold | Angular threshold (°) | Step size | Smoothing | Min length (mm) | Max length (mm) |
|---|---|---|---|---|---|---|---|---|
| Default | All others | | 0.05 | 0 | 0 | 0 | 20 | 250 |
| Local u-fibers between immediately adjacent ROIs | Lt_aMTG | Lt_aSTG | 0.02 | 0 | 0.7 | 0 | 20 | 250 |
| | Lt_aMTG | Lt_pMTG | 0.05 | 0 | 0 | 0 | 10 | 250 |
| | Lt_EG | Lt_PHG | 0.02 | 60 | 0.3 | 0 | 10 | 250 |
| | Lt_IG | Lt_pOp | 0.05 | 90 | 0 | 0 | 10 | 100 |
| | Lt_PCL | Lt_pCG | 0.05 | 0 | 0 | 0 | 10 | 250 |
| | Lt_pITG | Lt_pMTG | 0.07 | 90 | 0.3 | 0 | 10 | 250 |
| | Rt_aMTG | Rt_aSTG | 0.04 | 0 | 0 | 0 | 20 | 250 |
| | Rt_aSTG | Rt_EG | 0.05 | 90 | 0 | 0 | 10 | 100 |
| | Rt_EG | Rt_PHG | 0.02 | 60 | 0.3 | 0 | 10 | 250 |
| | Rt_iPoCG | Rt_IG | 0.04 | 0 | 0 | 0 | 20 | 250 |
| | Rt_iPreCG | Rt_IG | 0.04 | 0 | 0 | 0 | 20 | 250 |
| | Rt_LG | Rt_pCG | 0.05775 | 0 | 0 | 0 | 10 | 250 |
| | Rt_LG | Rt_pFG | 0.04 | 0 | 0 | 0 | 20 | 250 |
| | Rt_LG | Rt_Pcun | 0.04 | 0 | 0 | 0 | 20 | 250 |
| | Rt_pOrb | Rt_rMFG | 0.04 | 0 | 0 | 0 | 20 | 250 |
| | Rt_pCG | Rt_pSFG | 0.04 | 0 | 0 | 0 | 20 | 250 |
| | Rt_pFG | Rt_pITG | 0.05 | 0 | 0 | 0 | 10 | 250 |
| | Rt_pITG | Rt_pMTG | 0.07 | 90 | 0.3 | 0 | 10 | 250 |
| Inter-hemispheric fibers between precisely homotopic cortical ROIs | Lt_iPoCG | Rt_iPoCG | 0.02 | 0 | 0 | 0 | 20 | 250 |
| | Lt_IPG | Rt_IPG | 0.02 | 0 | 1 | 0 | 20 | 300 |
| | Lt_pOrb | Rt_pOrb | 0.02 | 70 | 1 | 0 | 50 | 300 |
| | Lt_pTr | Rt_pTr | 0.02 | 0 | 1 | 0 | 20 | 300 |
| | Lt_pCG | Rt_pCG | 0.07 | 70 | 0.3 | 0 | 20 | 80 |
| | Lt_pITG | Rt_pITG | 0.02 | 90 | 1 | 0 | 50 | 300 |
| | Lt_pMTG | Rt_pMTG | 0.02 | 90 | 1 | 0 | 50 | 300 |
| | Lt_pSTG | Rt_pSTG | 0.01 | 90 | 1 | 0 | 50 | 300 |
| | Lt_SMG | Rt_SMG | 0.02 | 0 | 1 | 0 | 50 | 300 |

**eFigure 4. Fiber tracking parameters for structural connectivity.** A total of 974 white matter streamlines were reconstructed and validated using the default parameter configuration. An additional 27 streamlines—18 representing local U-fibers between adjacent cortical regions of interest (ROIs) and 9 connecting precisely homotopic ROIs across hemispheres—were generated with a modified parameter setting. These alternate streamlines underwent the same visual validation procedure.

For visual clarity, we delineated only one white matter streamline per ROI pair. While this approach may result in false negatives by omitting genuine streamlines, it was necessary to effectively visualize major fasciculi in bird's-eye, posterior, and lateral views. Including too many lateral fasciculi would obscure medial structures, and excessive streamlines overall would block the visibility of posterior tracts.

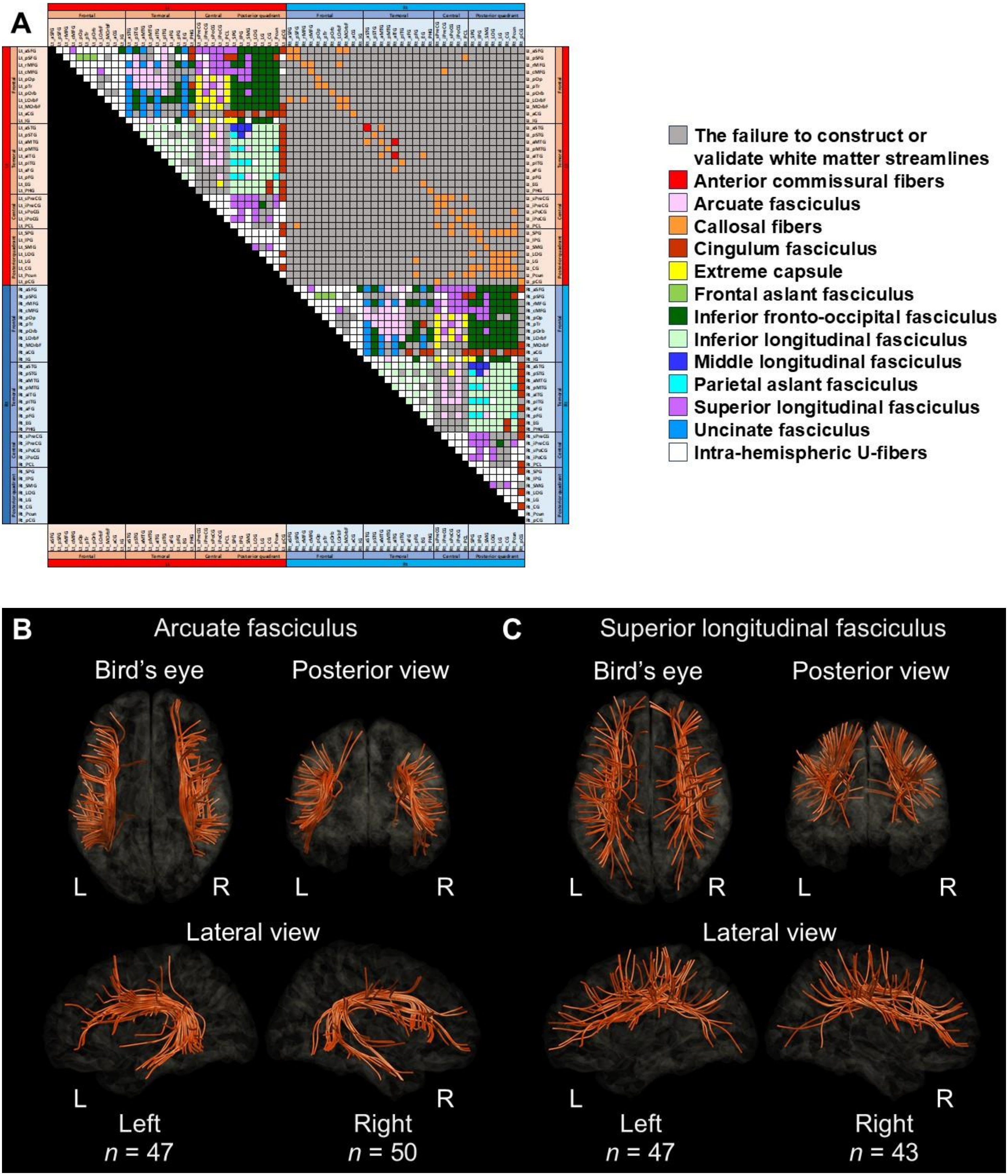
A
The failure to construct or validate white matter streamlines
Anterior commissural fibers
Arcuate fasciculus
Callosal fibers
Cingulum fasciculus
Extreme capsule
Frontal aslant fasciculus
Inferior fronto-occipital fasciculus
Inferior longitudinal fasciculus
Middle longitudinal fasciculus
Parietal aslant fasciculus
Superior longitudinal fasciculus
Uncinate fasciculus
Intra-hemispheric U-fibers
B
Arcuate fasciculus
Bird's eye
Posterior view
L
R
L
R
Lateral view
L
R
Left
n = 47
Right
n = 50
C
Superior longitudinal fasciculus
Bird's eye
Posterior view
L
R
L
R
Lateral view
L
R
Left
n = 47
Right
n = 43

**eFigure 5**

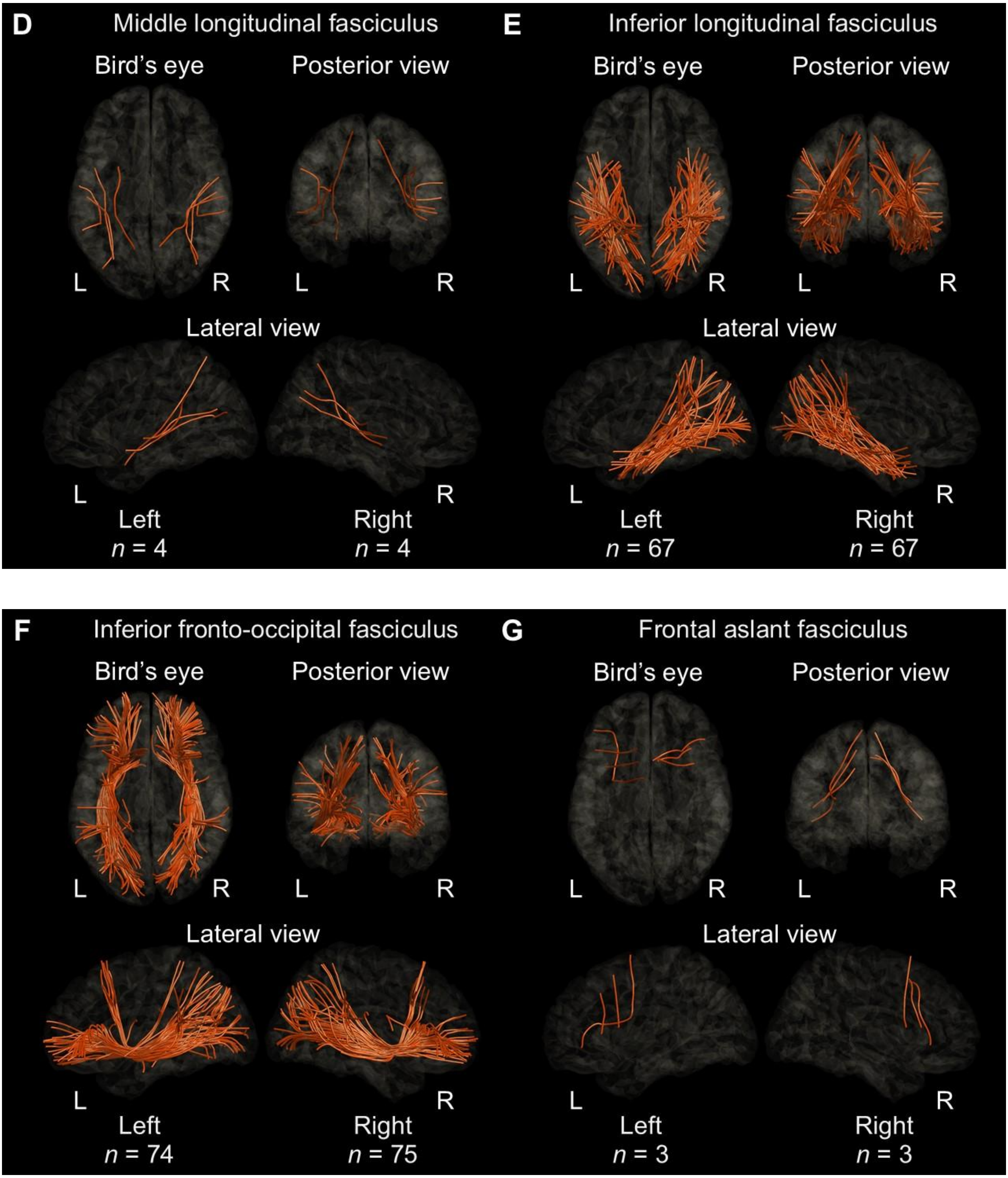
D
Middle longitudinal fasciculus
Bird's eye
Posterior view
L
R
L
R
Lateral view
L
R
Left
n = 4
Right
n = 4
E
Inferior longitudinal fasciculus
Bird's eye
Posterior view
L
R
L
R
Lateral view
L
R
Left
n = 67
Right
n = 67
F
Inferior fronto-occipital fasciculus
Bird's eye
Posterior view
L
R
L
R
Lateral view
L
R
Left
n = 74
Right
n = 75
G
Frontal aslant fasciculus
Bird's eye
Posterior view
L
R
L
R
Lateral view
L
R
Left
n = 3
Right
n = 3

**eFigure 5**

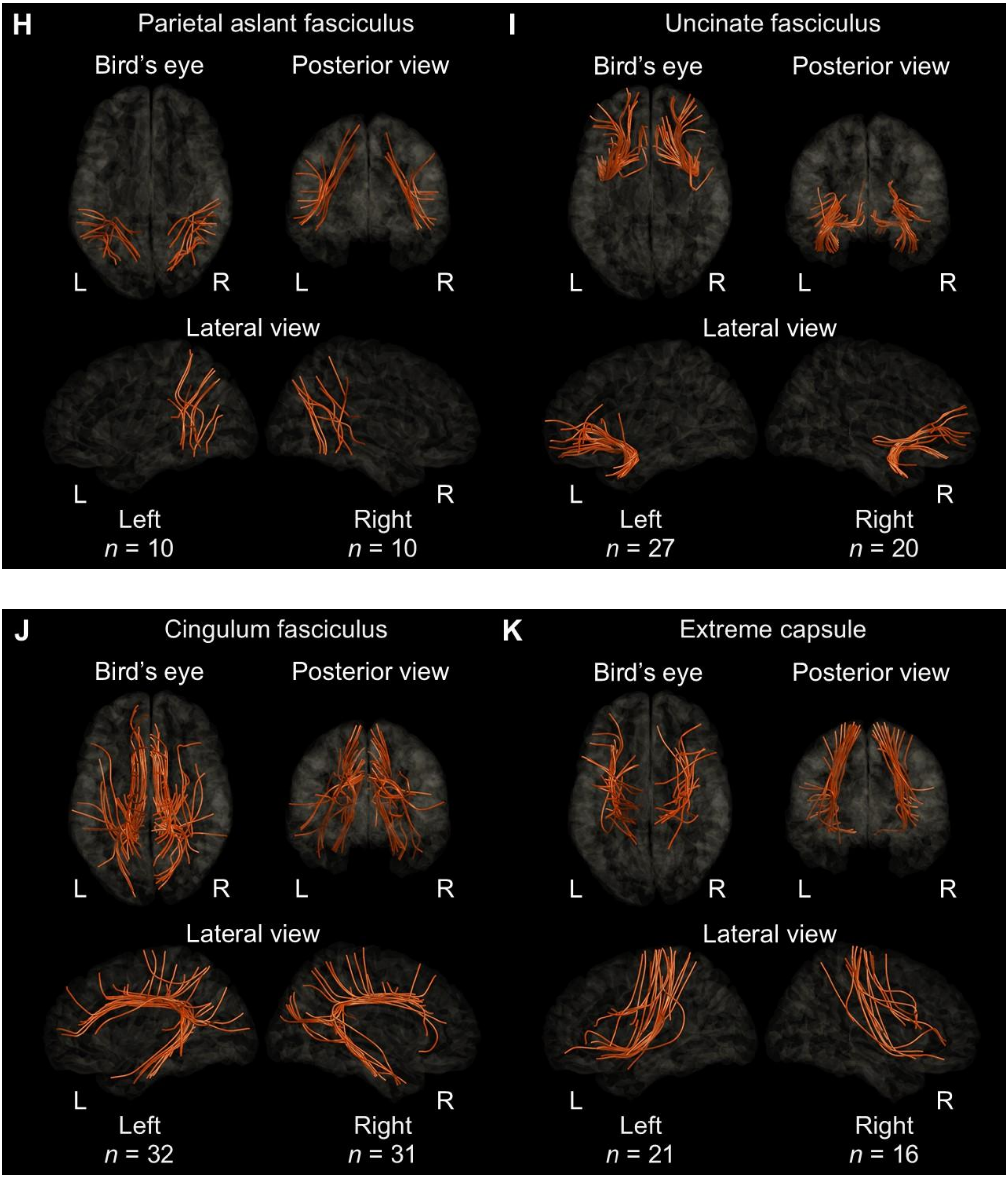
H
Parietal aslant fasciculus
Bird's eye
Posterior view
L
R
L
R
Lateral view
L
R
Left
n = 10
Right
n = 10
I
Uncinate fasciculus
Bird's eye
Posterior view
L
R
L
R
Lateral view
L
R
Left
n = 27
Right
n = 20
J
Cingulum fasciculus
Bird's eye
Posterior view
L
R
L
R
Lateral view
L
R
Left
n = 32
Right
n = 31
K
Extreme capsule
Bird's eye
Posterior view
L
R
L
R
Lateral view
L
R
Left
n = 21
Right
n = 16

**eFigure 5**

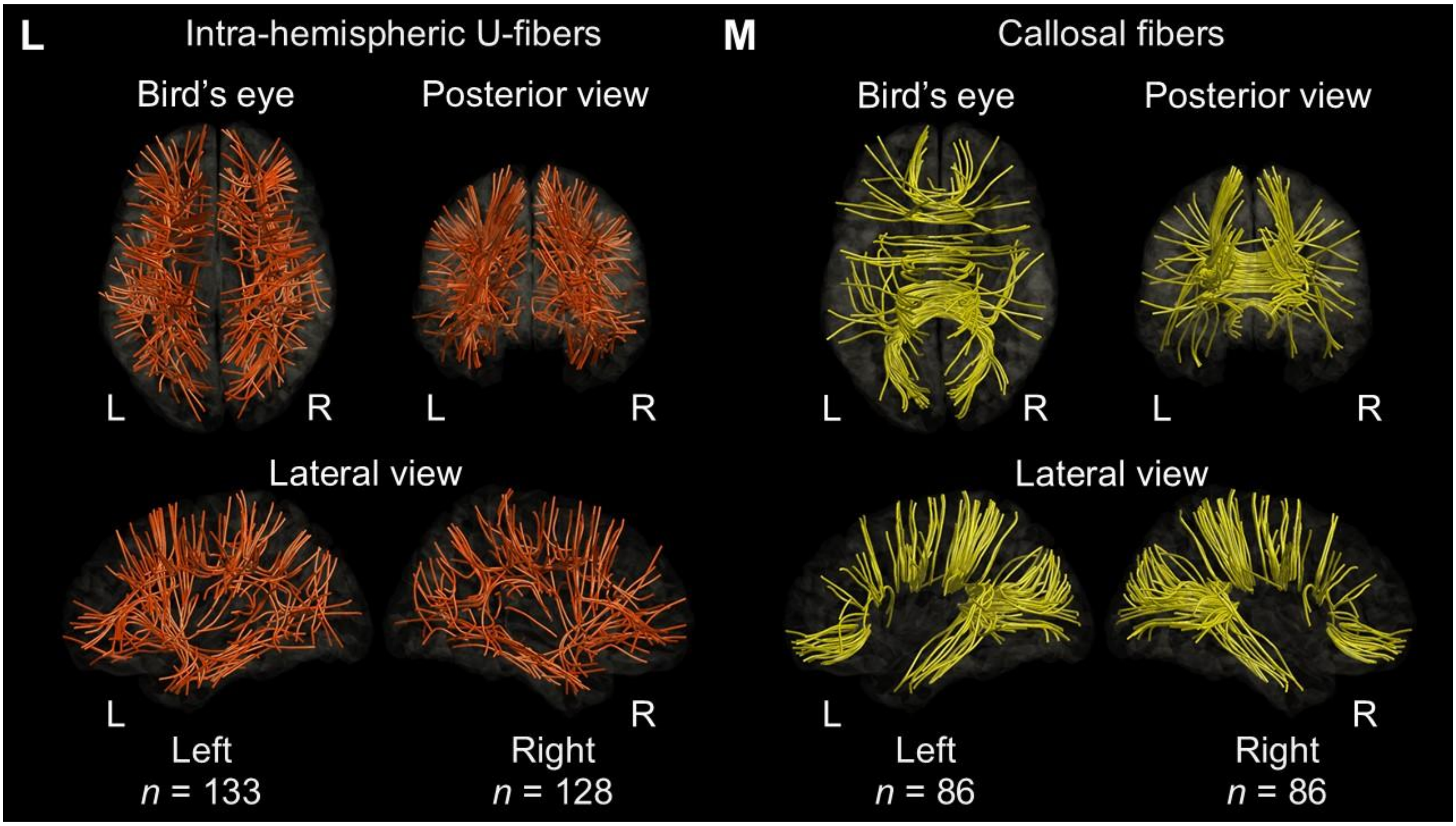


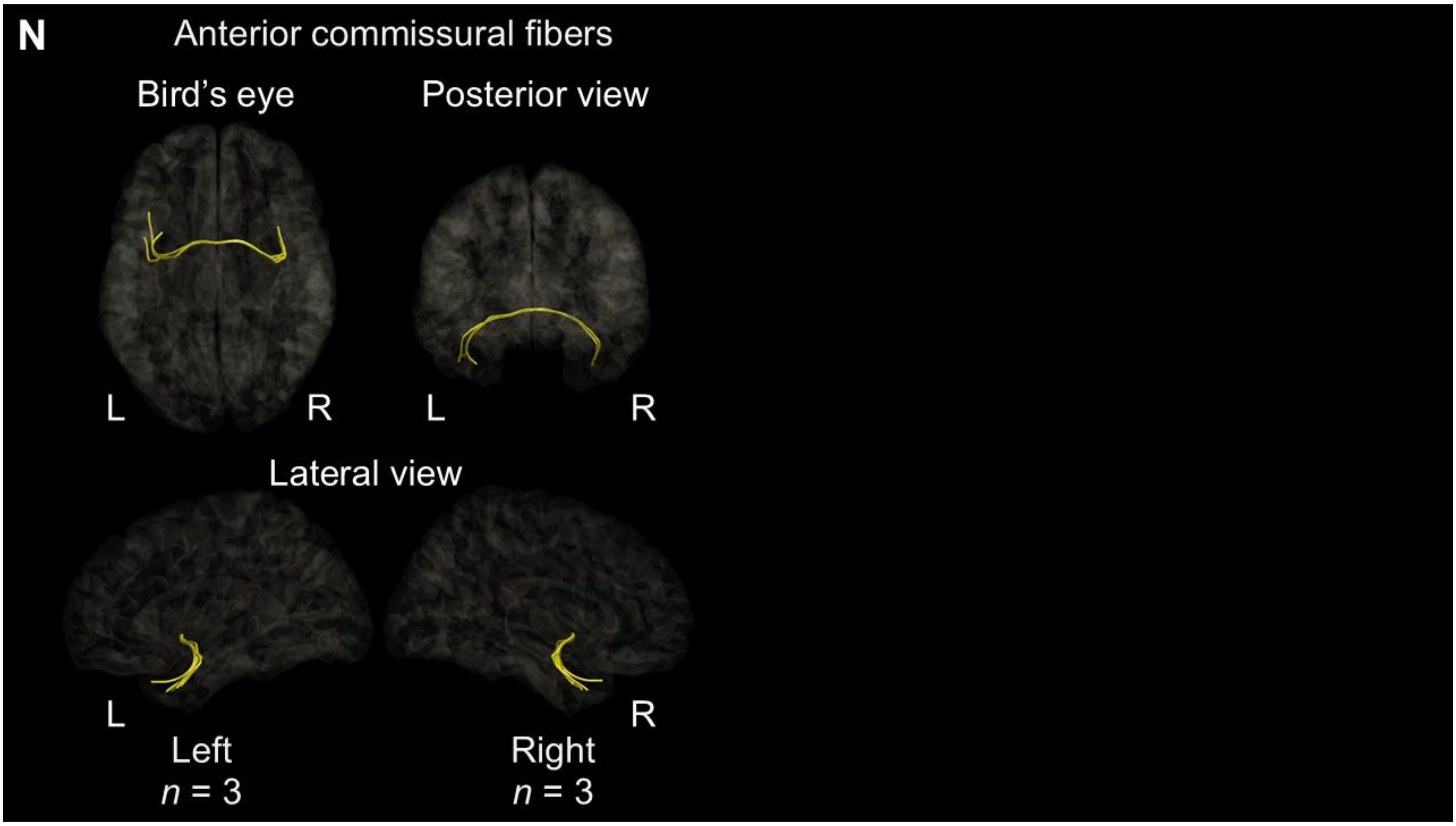


**eFigure 5. Structural connectivity classification**. (A) Each pair of cortical regions of interest (ROIs) in this structural connectivity matrix is assigned a label corresponding to a specific fasciculus. (B) Arcuate fasciculus. (C) Superior longitudinal fasciculus. (D) Middle longitudinal fasciculus. (E) Inferior longitudinal fasciculus. (F) Inferior fronto-occipital fasciculus. (G) Frontal aslant fasciculus. (H) Parietal aslant fasciculus. (I) Uncinate fasciculi. (J) Cingulum fasciculus. (K) Extreme capsule. (L) Intrahemispheric U-fibers. (M) Callosal fibers. (N) Anterior commissural fibers.

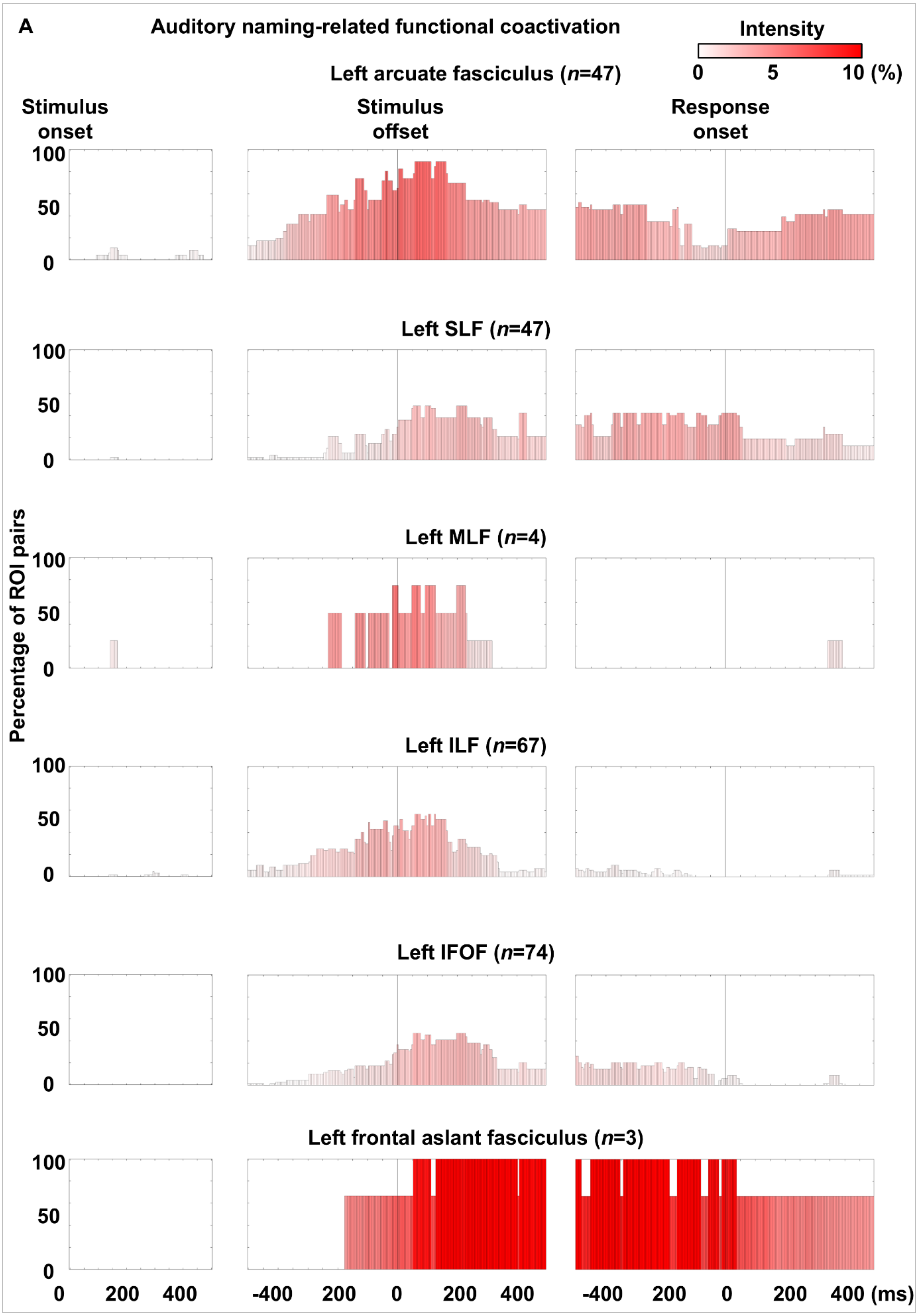
A
Auditory naming-related functional coactivation
Intensity
0
5
10 (%)
Left arcuate fasciculus (n=47)
Stimulus onset
Stimulus offset
Response onset
Left SLF (n=47)
Left MLF (n=4)
Left ILF (n=67)
Left IFOF (n=74)
Left frontal aslant fasciculus (n=3)
Percentage of ROI pairs
100
50
0
200
400
-400
(ms)

eFigure 6

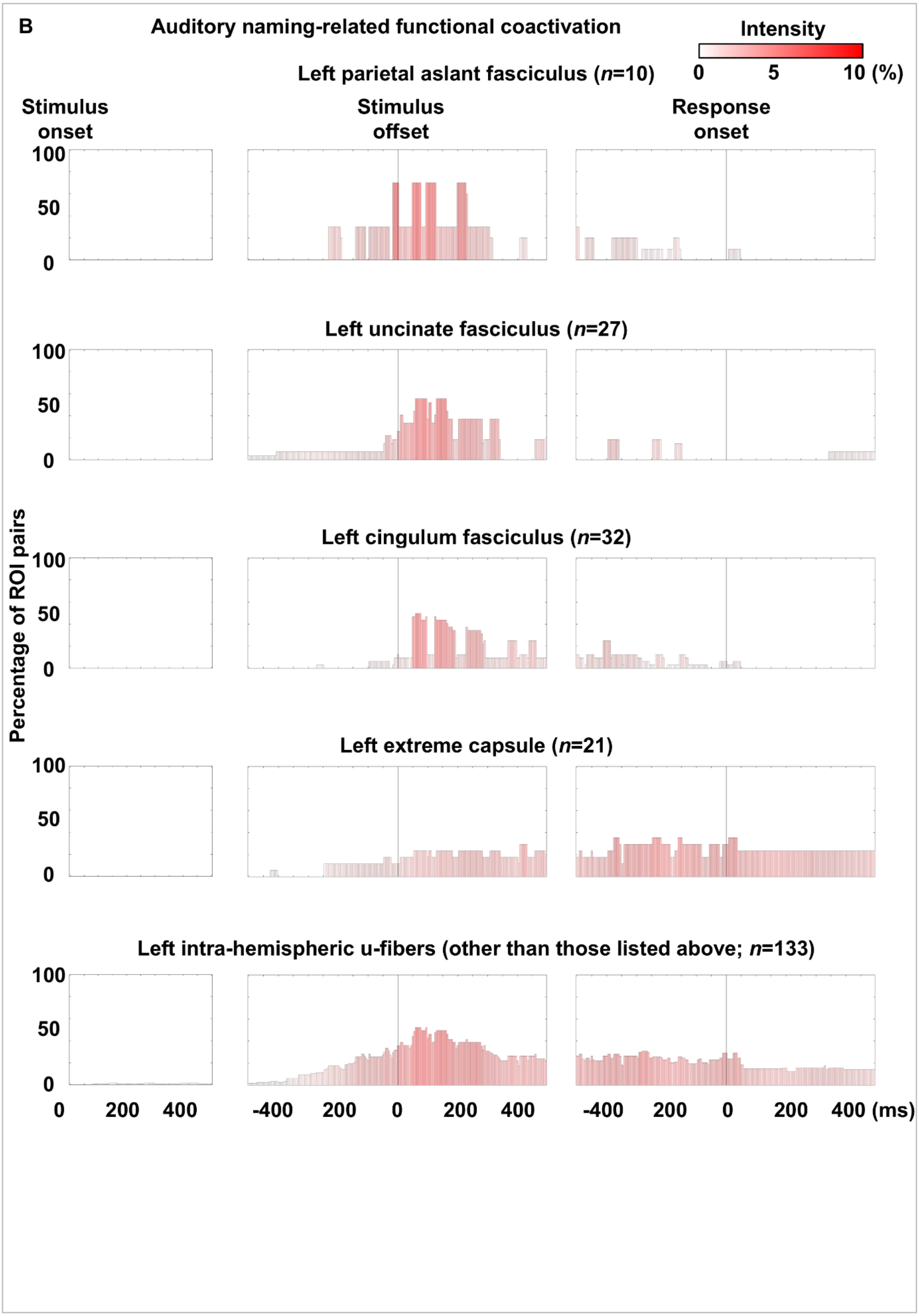
B
Auditory naming-related functional coactivation
Intensity
0
5
10 (%)
Left parietal aslant fasciculus (n=10)
Stimulus onset
Stimulus offset
Response onset
100
50
0
Left uncinate fasciculus (n=27)
Left cingulum fasciculus (n=32)
Percentage of ROI pairs
Left extreme capsule (n=21)
Left intra-hemispheric u-fibers (other than those listed above; n=133)
0
200
400
-400
200
0
200
400 (ms)

eFigure 6

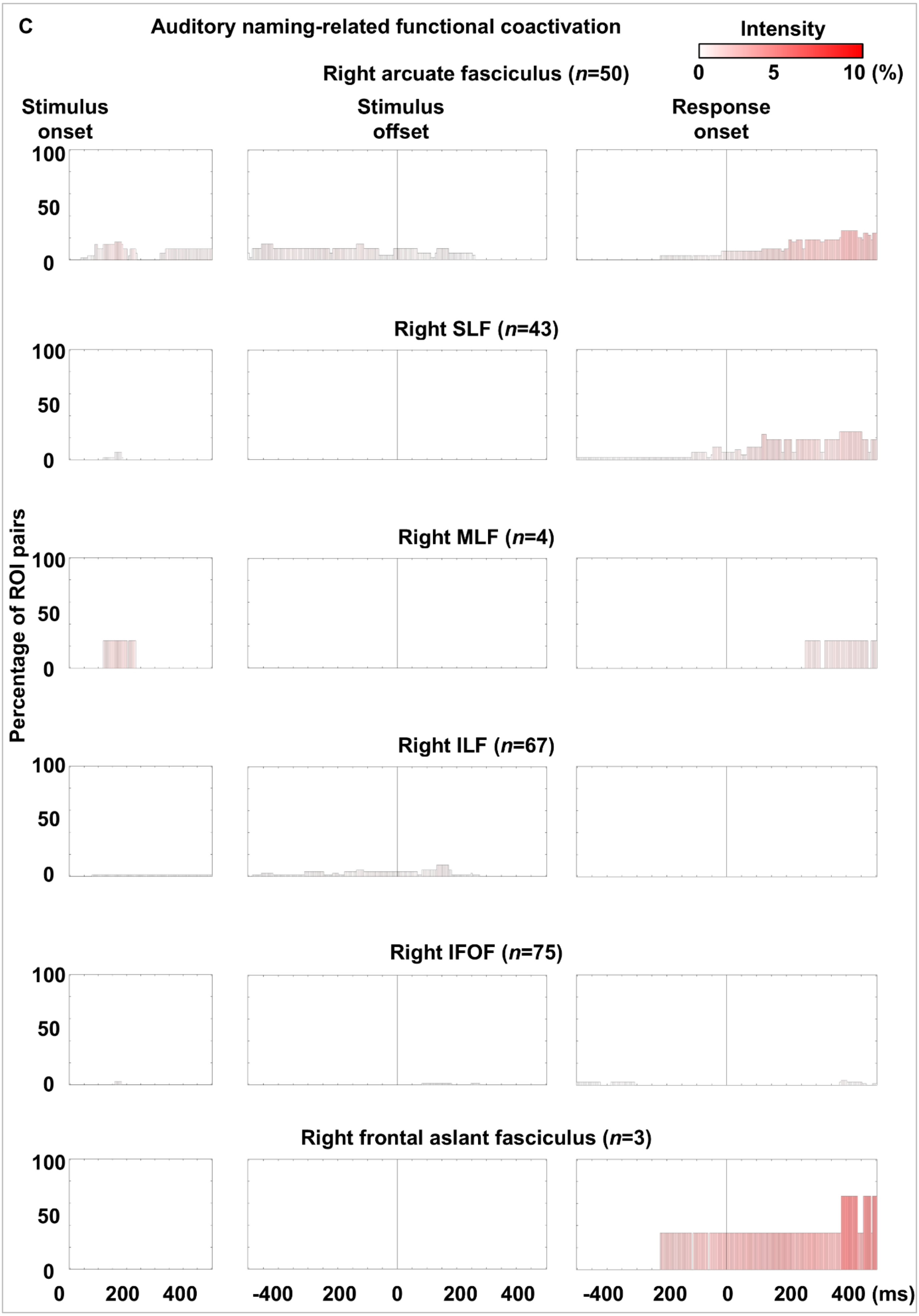
C
Auditory naming-related functional coactivation
Intensity
0
5
10 (%)
Right arcuate fasciculus (n=50)
Stimulus onset
Stimulus offset
Response onset
Right SLF (n=43)
Right MLF (n=4)
Right ILF (n=67)
Right IFOF (n=75)
Right frontal aslant fasciculus (n=3)
Percentage of ROI pairs
100
50
0
0
200
400
-400
200
0
200
400
-400
200
0
200
400 (ms)

eFigure 6

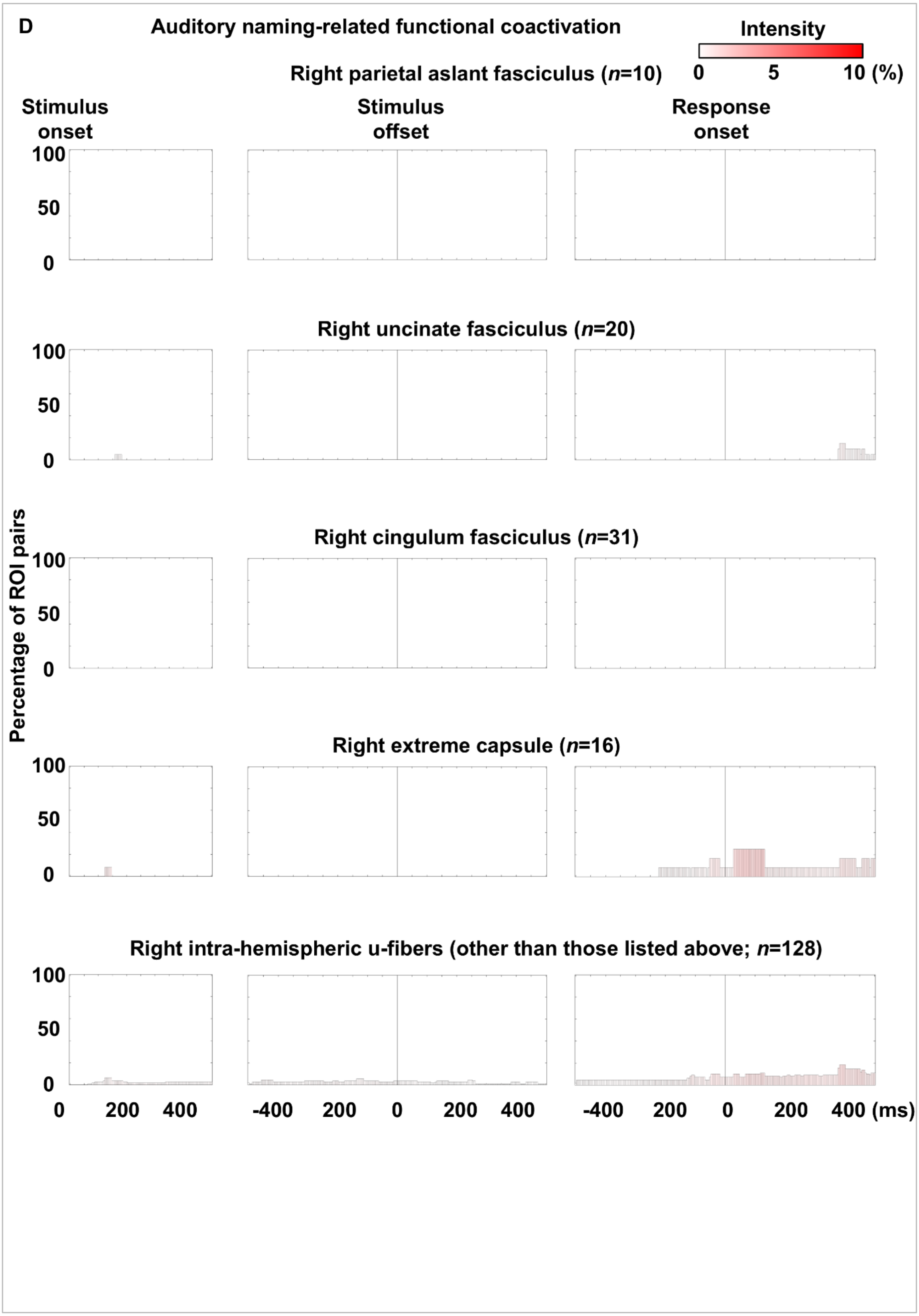

D
Auditory naming-related functional coactivation
Intensity
0
5
10 (%)
Right parietal aslant fasciculus (n=10)
Stimulus onset
Stimulus offset
Response onset
100
50
0
Right uncinate fasciculus (n=20)
Right cingulum fasciculus (n=31)
Right extreme capsule (n=16)
Right intra-hemispheric u-fibers (other than those listed above; n=128)
Percentage of ROI pairs
0
200
400
-400
200
0
200
400 (ms)


eFigure 6

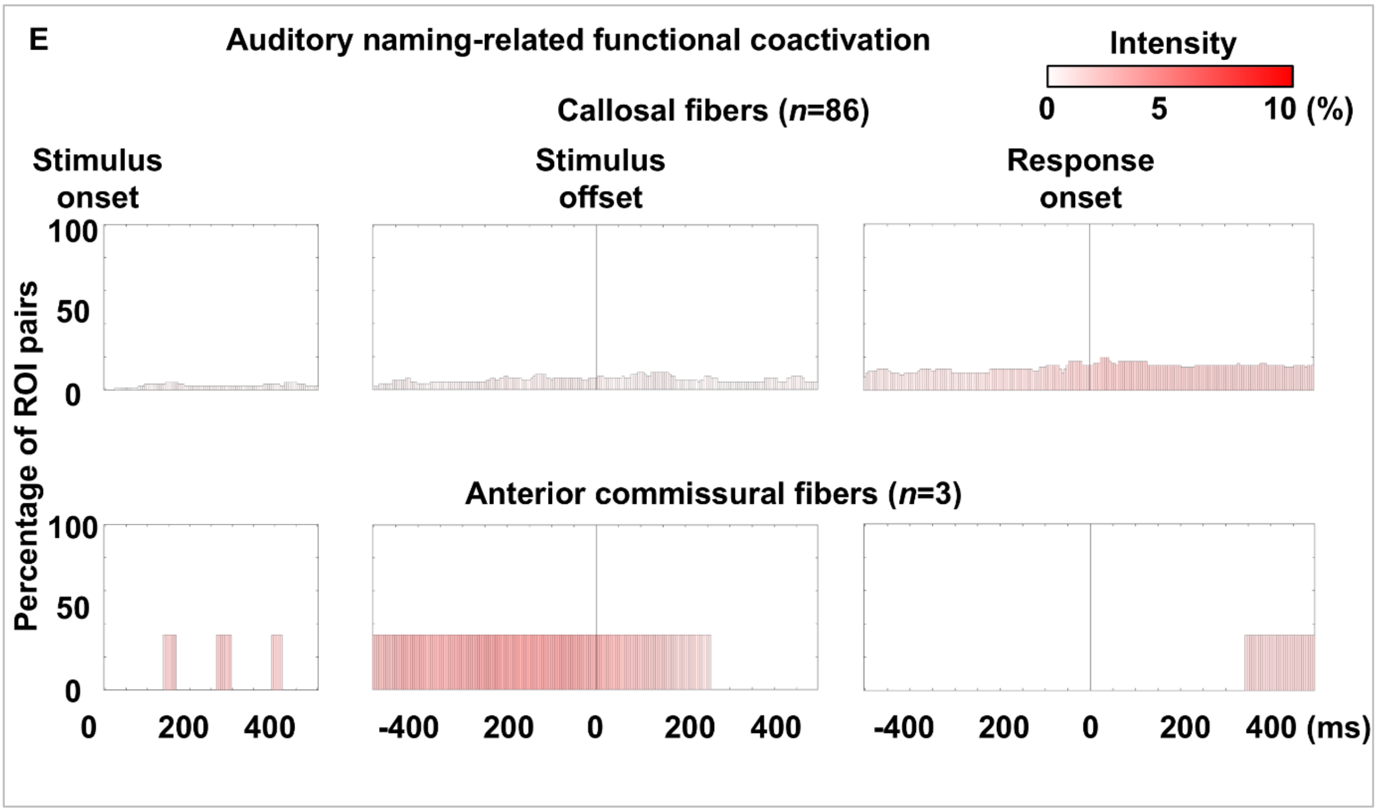
E
Auditory naming-related functional coactivation
Intensity
0 5 10 (%)
Callosal fibers (n=86)
Stimulus onset
Stimulus offset
Response onset
Percentage of ROI pairs
100
50
0
Anterior commissural fibers (n=3)
0 200 400
-400 200 0 200 400
-400 200 0 200 400 (ms)

eFigure 6

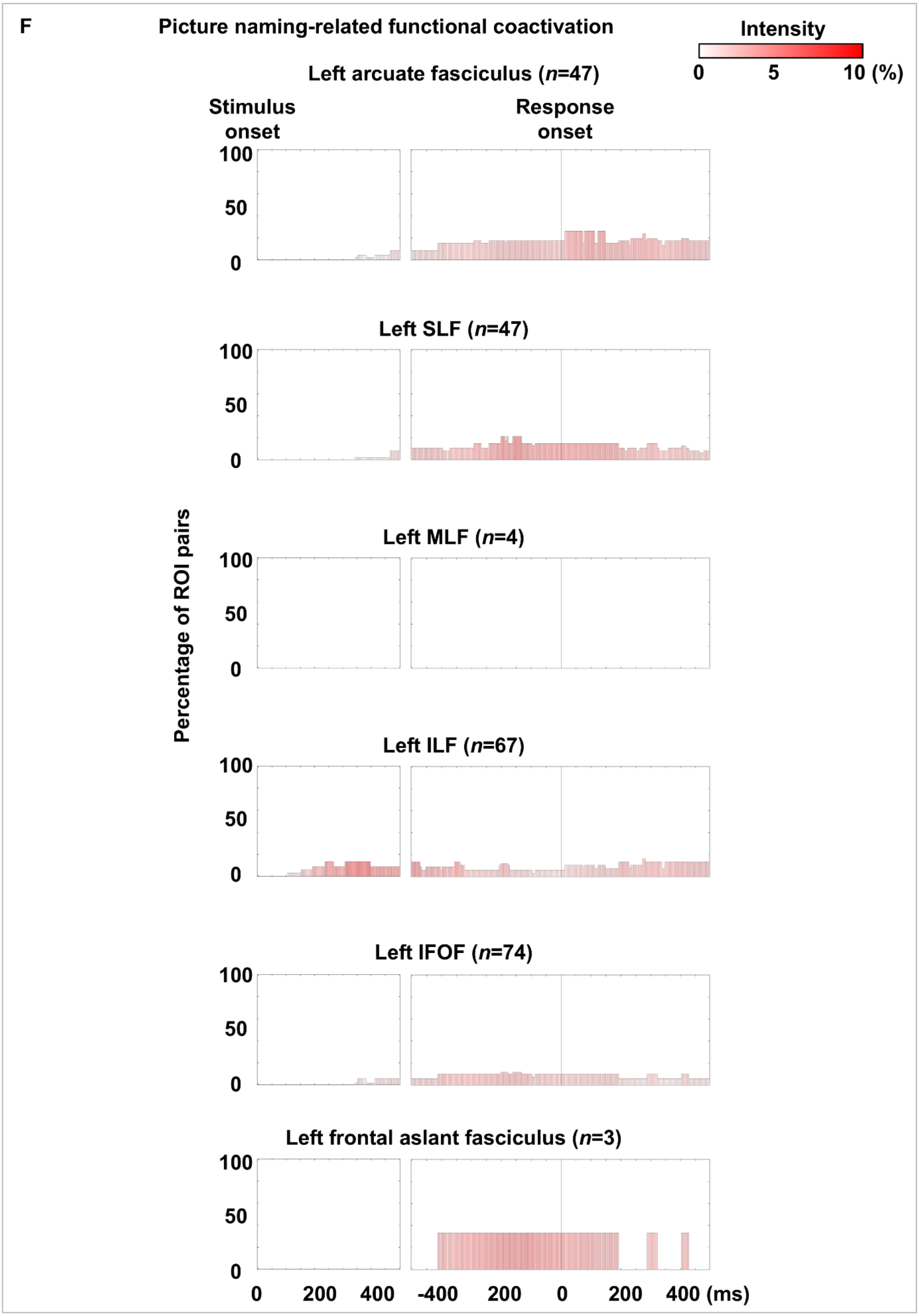
F
Picture naming-related functional coactivation
Intensity
0 5 10 (%)
Left arcuate fasciculus (n=47)
Stimulus onset
Response onset
Left SLF (n=47)
Left MLF (n=4)
Left ILF (n=67)
Left IFOF (n=74)
Left frontal aslant fasciculus (n=3)
Percentage of ROI pairs
100
50
0
0 200 400 -400 200 0 200 400 (ms)

eFigure 6

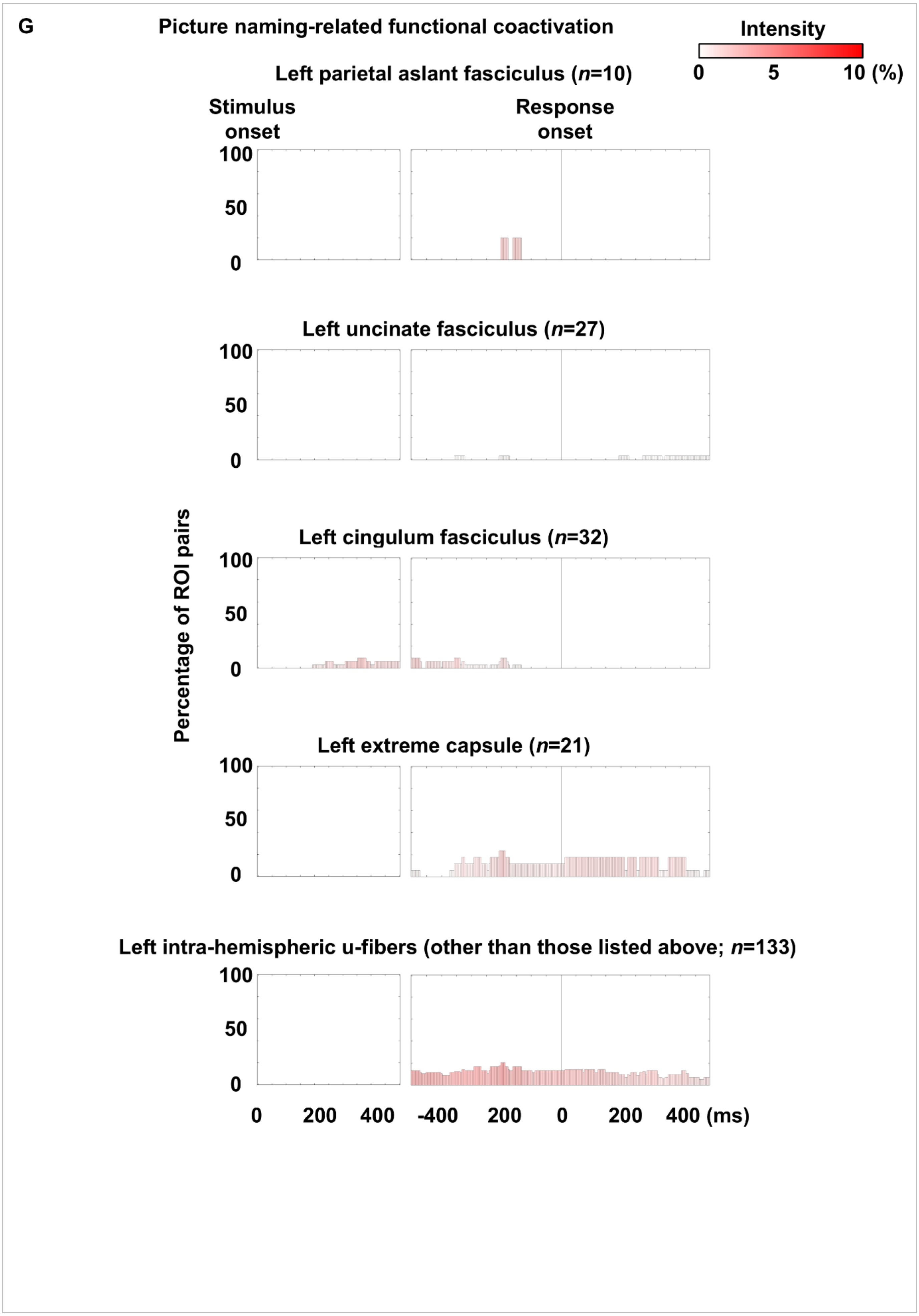
G
Picture naming-related functional coactivation
Intensity
0
5
10 (%)
Left parietal aslant fasciculus (n=10)
Stimulus onset
Response onset
100
50
0
Left uncinate fasciculus (n=27)
Left cingulum fasciculus (n=32)
Left extreme capsule (n=21)
Left intra-hemispheric u-fibers (other than those listed above; n=133)
Percentage of ROI pairs
0
200
400
-400
200
0
200
400 (ms)

eFigure 6

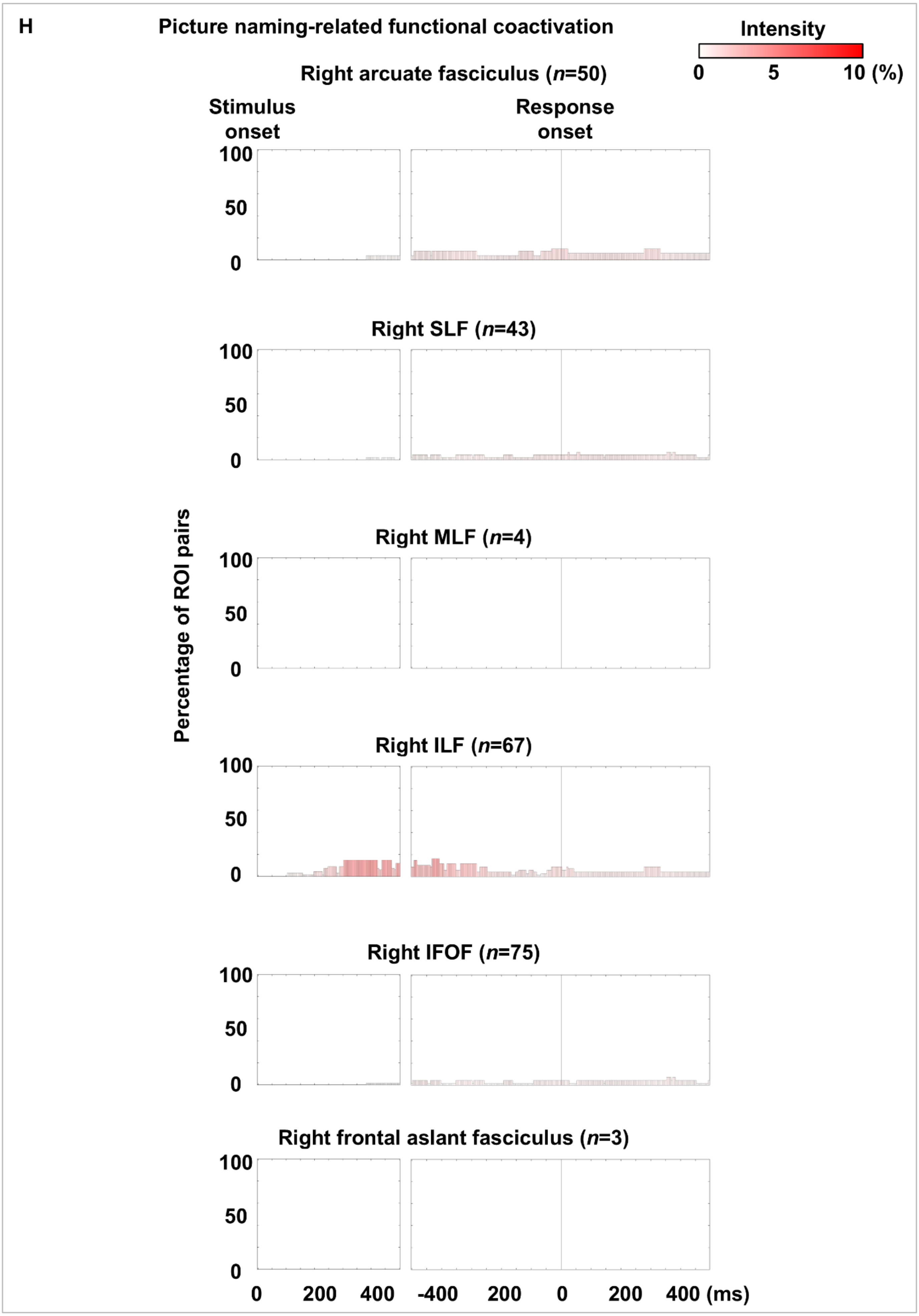
H
Picture naming-related functional coactivation
Intensity
0
5
10 (%)
Right arcuate fasciculus (n=50)
Stimulus onset
Response onset
Right SLF (n=43)
Right MLF (n=4)
Right ILF (n=67)
Right IFOF (n=75)
Right frontal aslant fasciculus (n=3)
Percentage of ROI pairs
100
50
0
0
200
400
-400
200
0
200
400 (ms)

eFigure 6

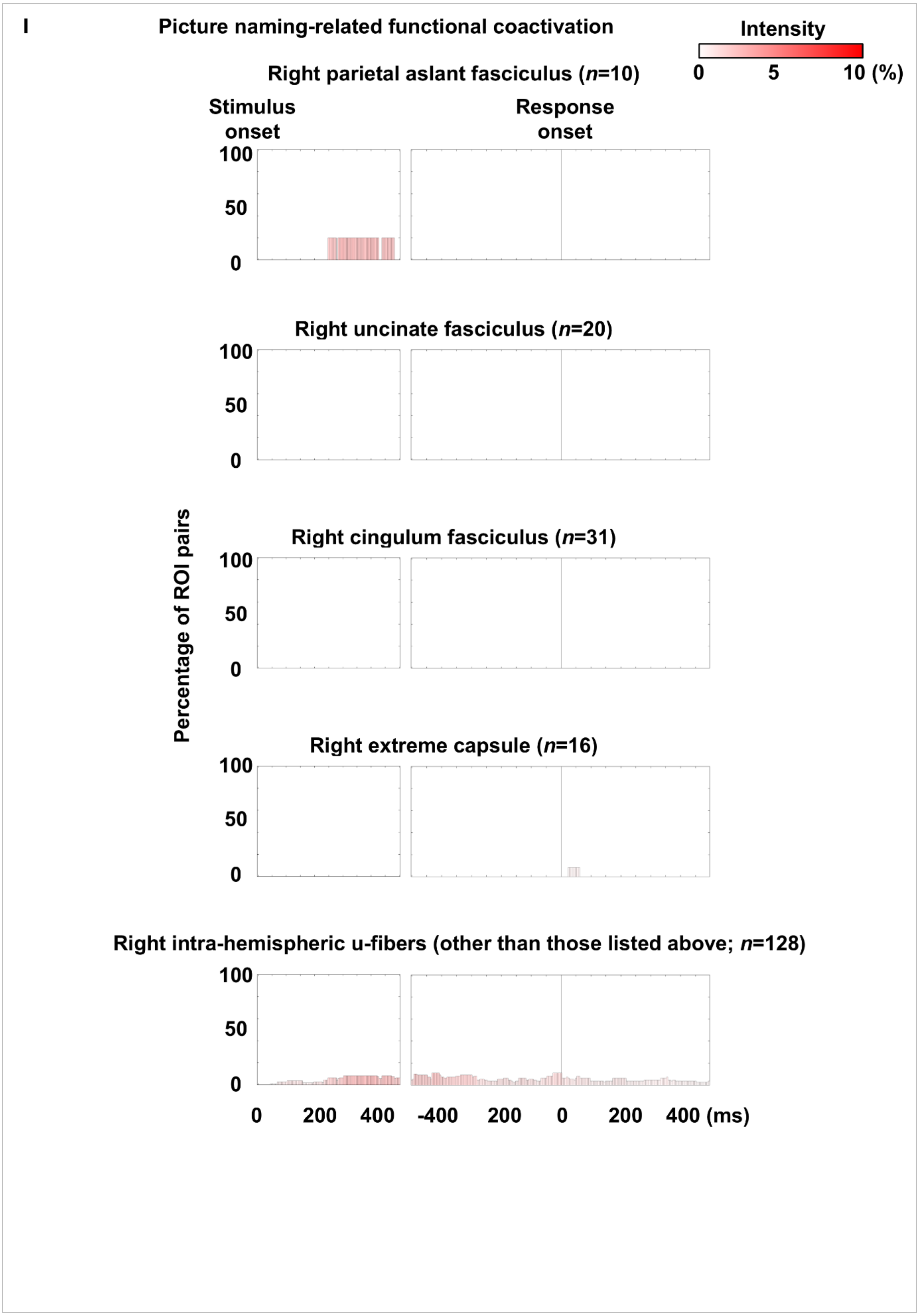
I
Picture naming-related functional coactivation
Intensity
0
5
10 (%)
Right parietal aslant fasciculus (n=10)
Stimulus onset
Response onset
100
50
0
Right uncinate fasciculus (n=20)
100
50
0
Right cingulum fasciculus (n=31)
100
50
0
Percentage of ROI pairs
Right extreme capsule (n=16)
100
50
0
Right intra-hemispheric u-fibers (other than those listed above; n=128)
100
50
0
0
200
400
-400
200
0
200
400 (ms)

eFigure 6

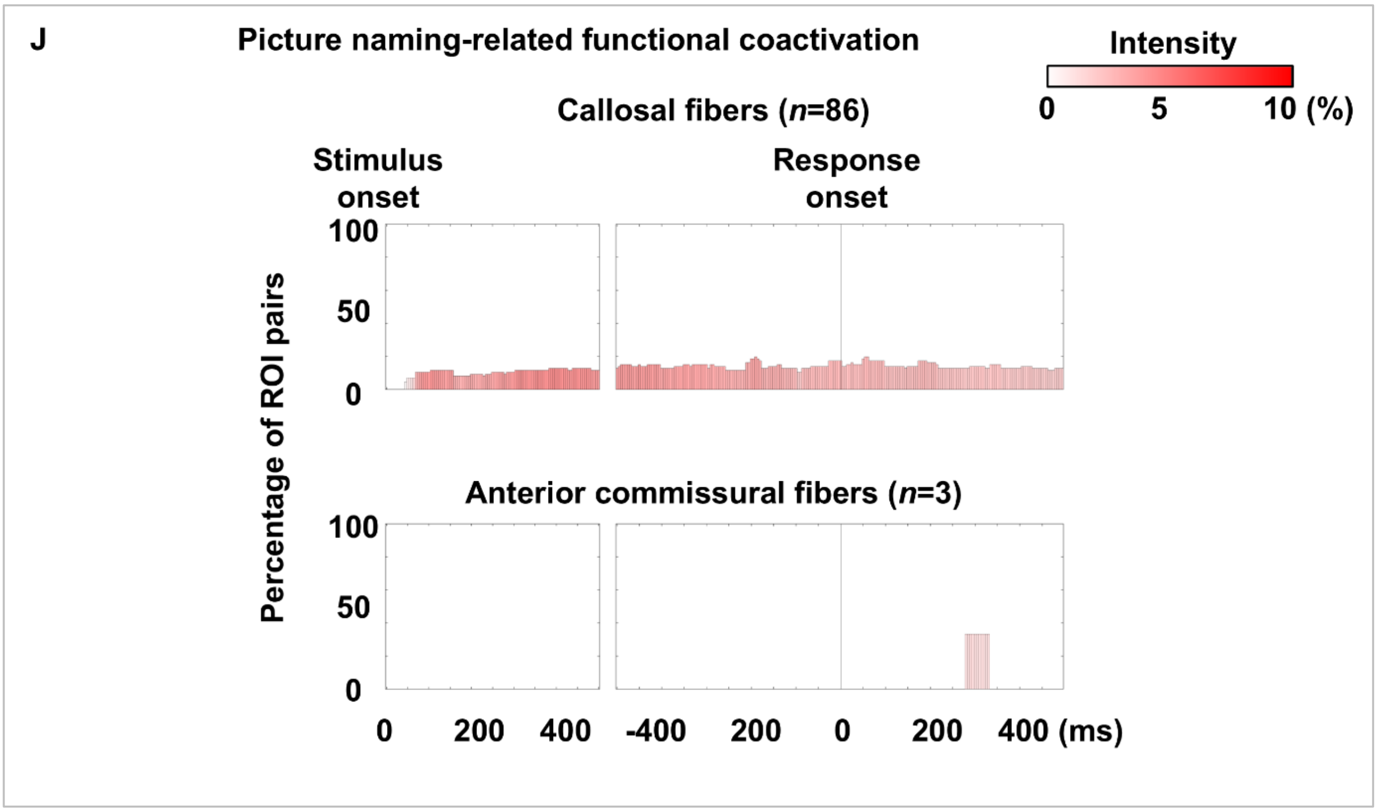


**eFigure 6. Auditory and picture naming-related functional coactivation through each fasciculus.** Bar height indicates the proportion of pathways showing functional coactivation within each fasciculus per time bin; bar color reflects the average coactivation intensity. (A-E) Auditory naming. (F-J) Picture naming. (A, B, F, G) Left intra-hemispheric pathways. (C, D, H, I) Right intra-hemispheric pathways. (E, J) Inter-hemispheric pathways.

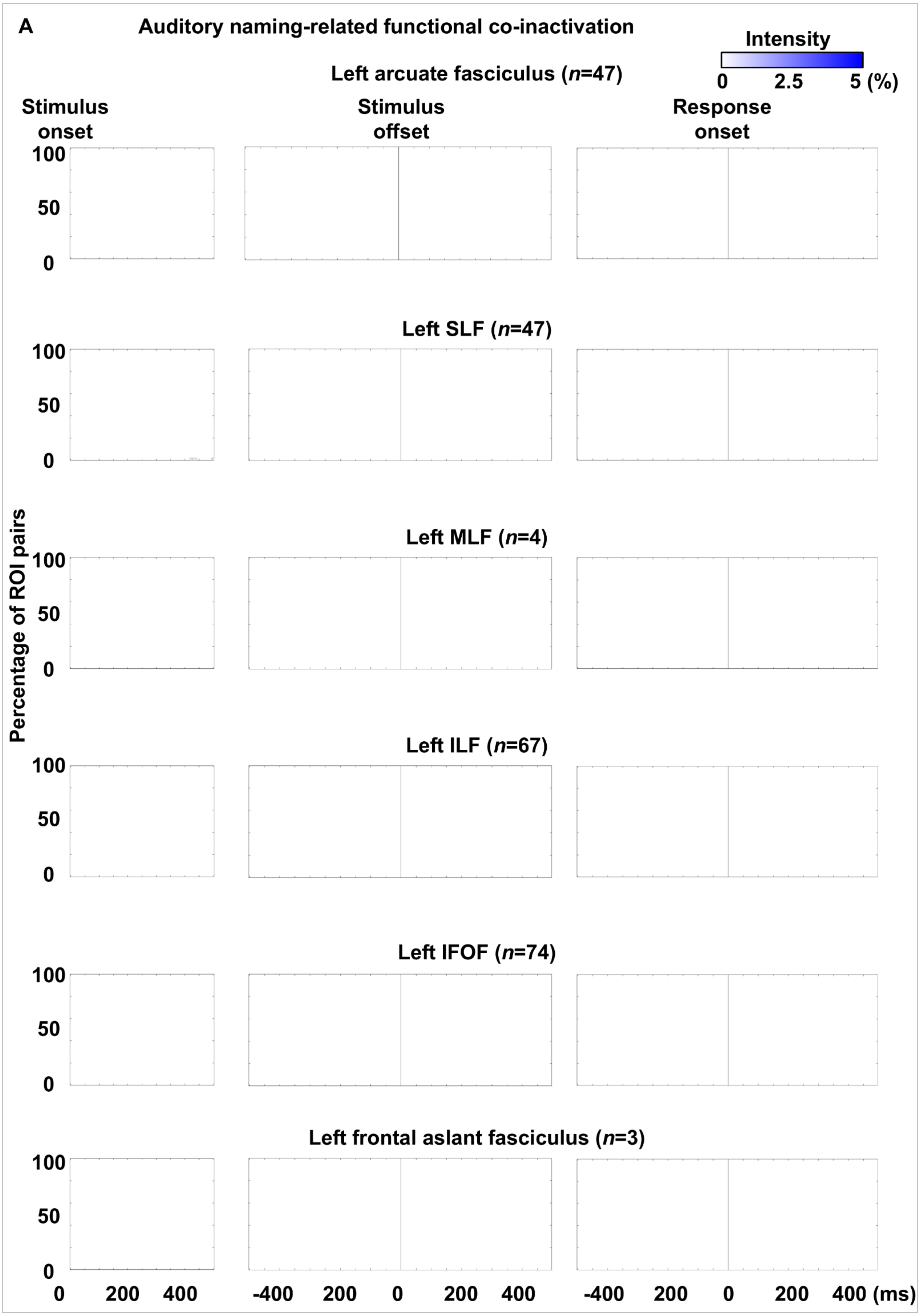
A
Auditory naming-related functional co-inactivation
Intensity
0 2.5 5 (%)
Left arcuate fasciculus (n=47)
Stimulus onset
Stimulus offset
Response onset
Left SLF (n=47)
Left MLF (n=4)
Left ILF (n=67)
Left IFOF (n=74)
Left frontal aslant fasciculus (n=3)
Percentage of ROI pairs
100
50
0
0 200 400
-400 200 0 200 400
-400 200 0 200 400 (ms)

eFigure 7

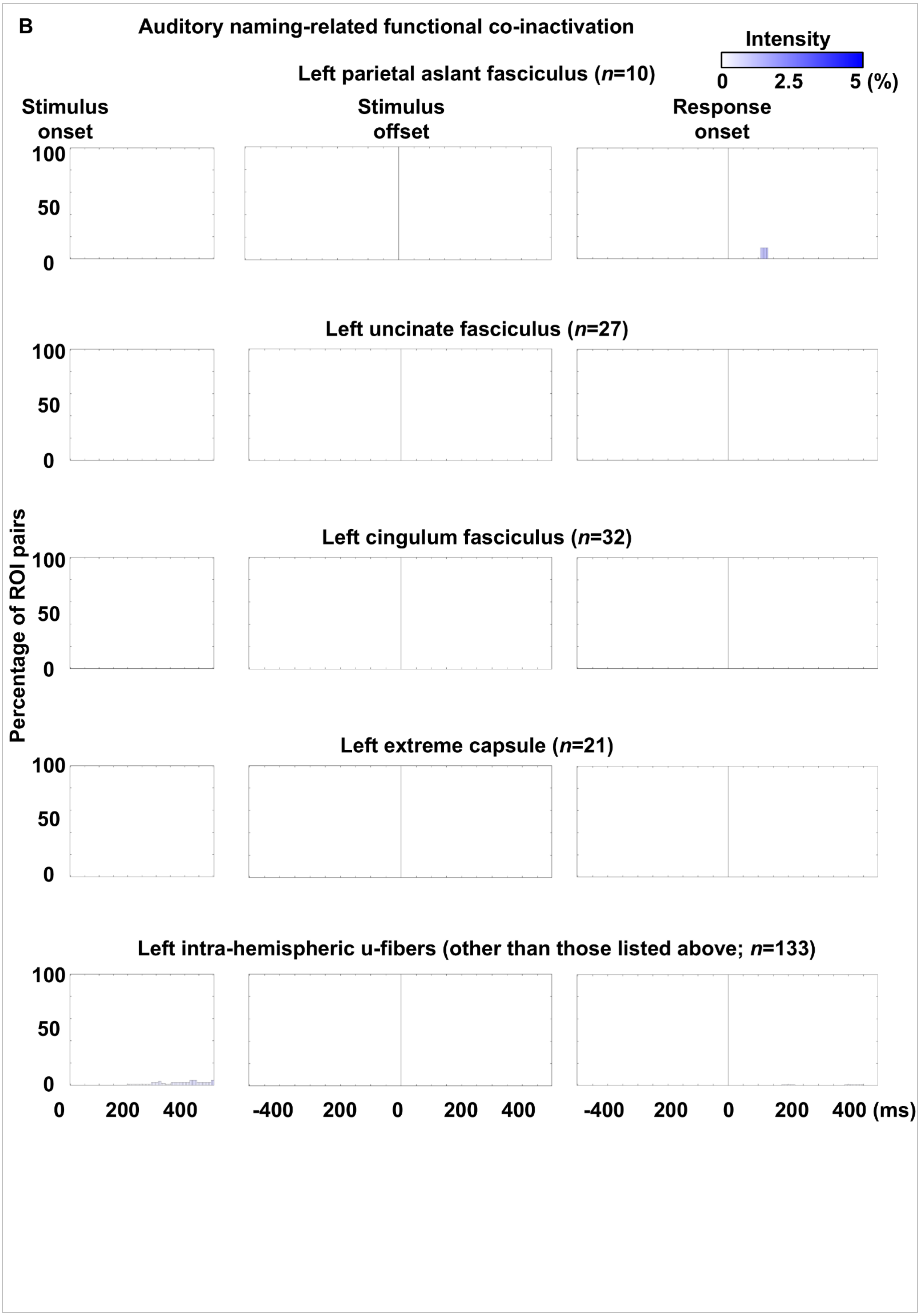

B
Auditory naming-related functional co-inactivation
Intensity
0
2.5
5 (%)
Left parietal aslant fasciculus (n=10)
Stimulus onset
Stimulus offset
Response onset
Left uncinate fasciculus (n=27)
Left cingulum fasciculus (n=32)
Left extreme capsule (n=21)
Left intra-hemispheric u-fibers (other than those listed above; n=133)
Percentage of ROI pairs
100
50
0
0
200
400
-400
200
0
200
400 (ms)


eFigure 7

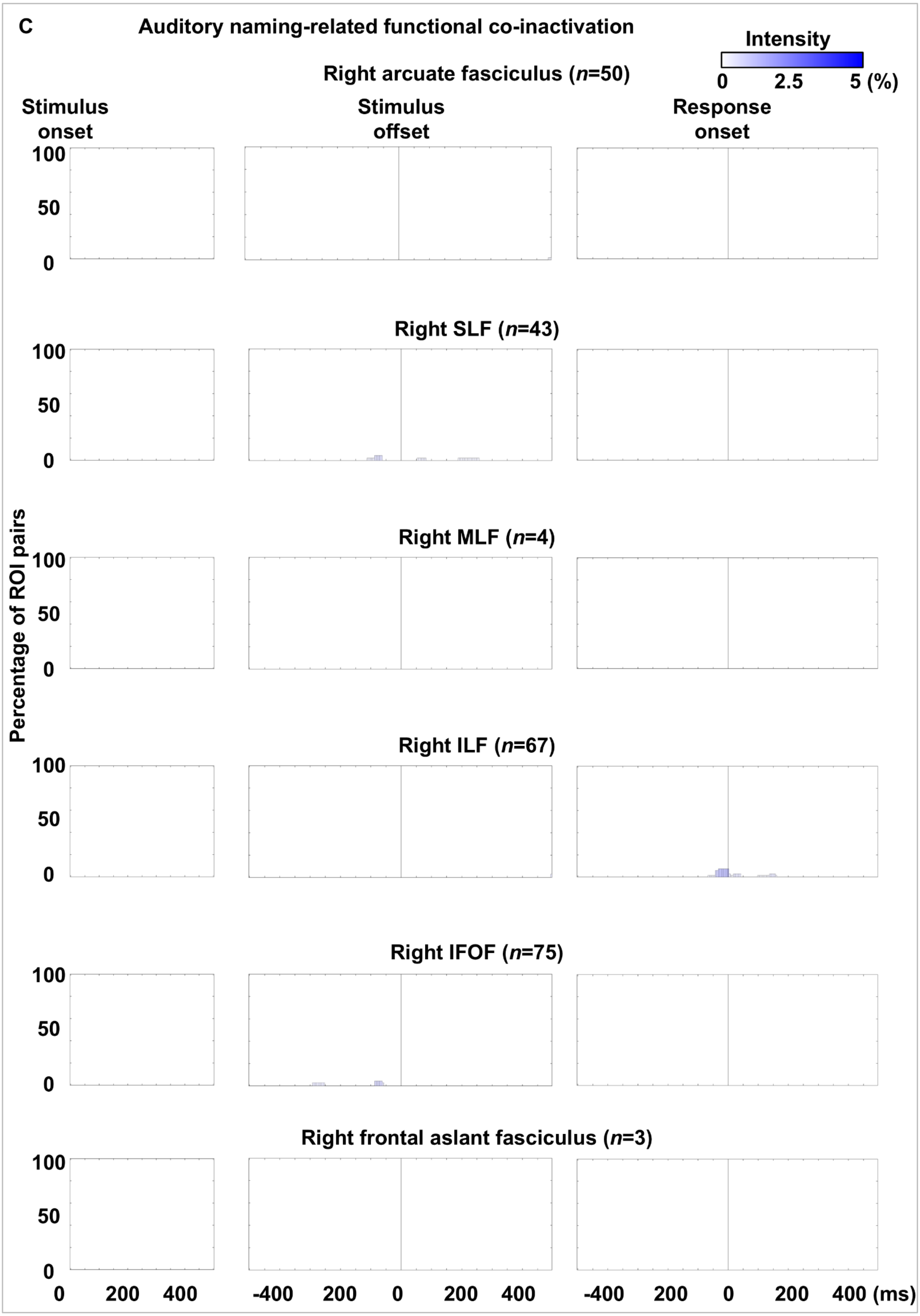
C
Auditory naming-related functional co-inactivation
Intensity
0
2.5
5 (%)
Right arcuate fasciculus (n=50)
Stimulus onset
Stimulus offset
Response onset
Right SLF (n=43)
Right MLF (n=4)
Right ILF (n=67)
Right IFOF (n=75)
Right frontal aslant fasciculus (n=3)
Percentage of ROI pairs
100
50
0
200
400
-400
(ms)

eFigure 7

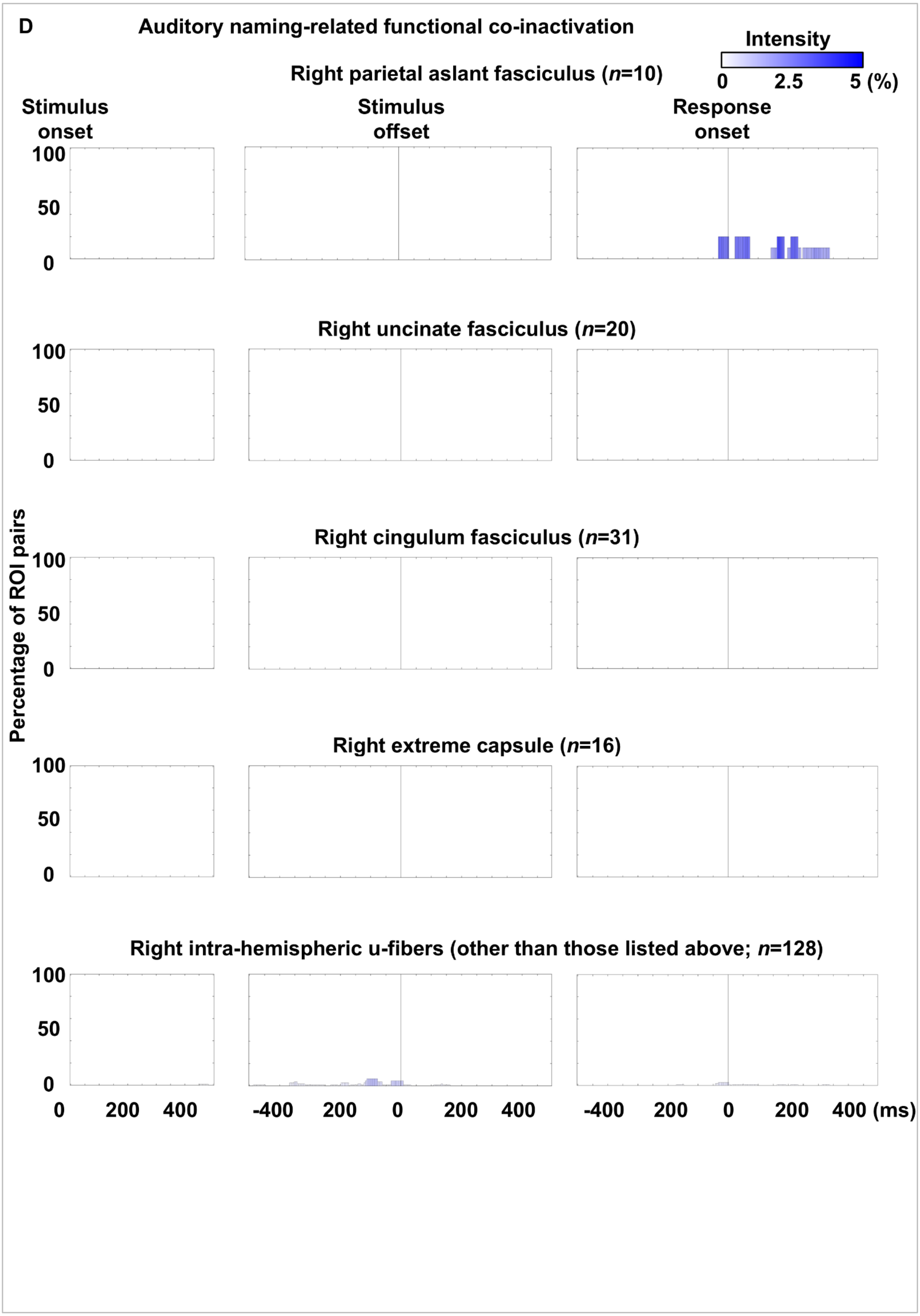
D
Auditory naming-related functional co-inactivation
Intensity
0 2.5 5 (%)
Right parietal aslant fasciculus (n=10)
Stimulus onset
Stimulus offset
Response onset
Right uncinate fasciculus (n=20)
Right cingulum fasciculus (n=31)
Right extreme capsule (n=16)
Right intra-hemispheric u-fibers (other than those listed above; n=128)
Percentage of ROI pairs
100
50
0
0 200 400
-400 200 0 200 400
-400 200 0 200 400 (ms)

eFigure 7

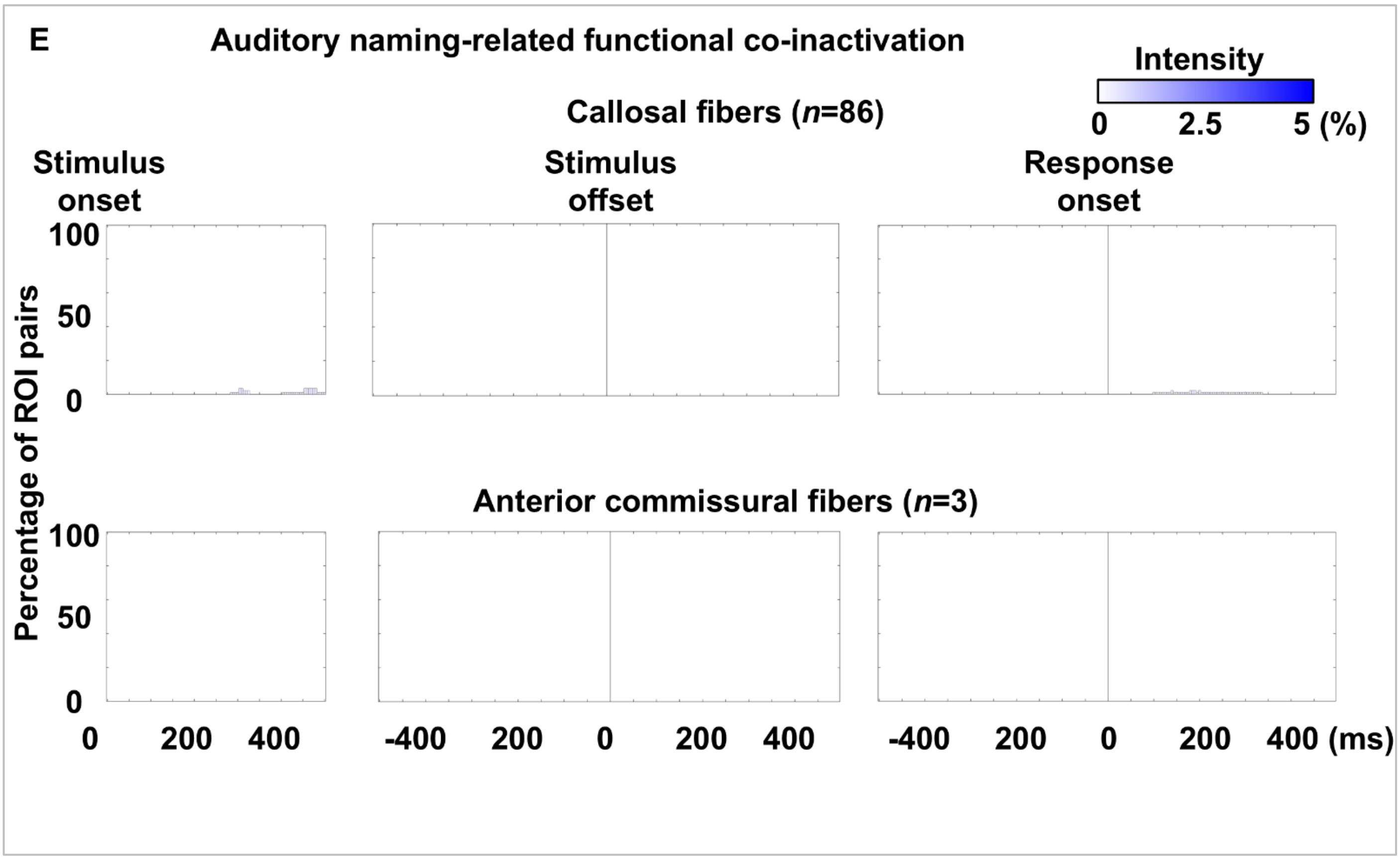
E
Auditory naming-related functional co-inactivation
Intensity
0 2.5 5 (%)
Callosal fibers (n=86)
Stimulus onset
Stimulus offset
Response onset
Percentage of ROI pairs
100
50
0
Anterior commissural fibers (n=3)
100
50
0
0 200 400
-400 200 0 200 400
-400 200 0 200 400 (ms)

eFigure 7

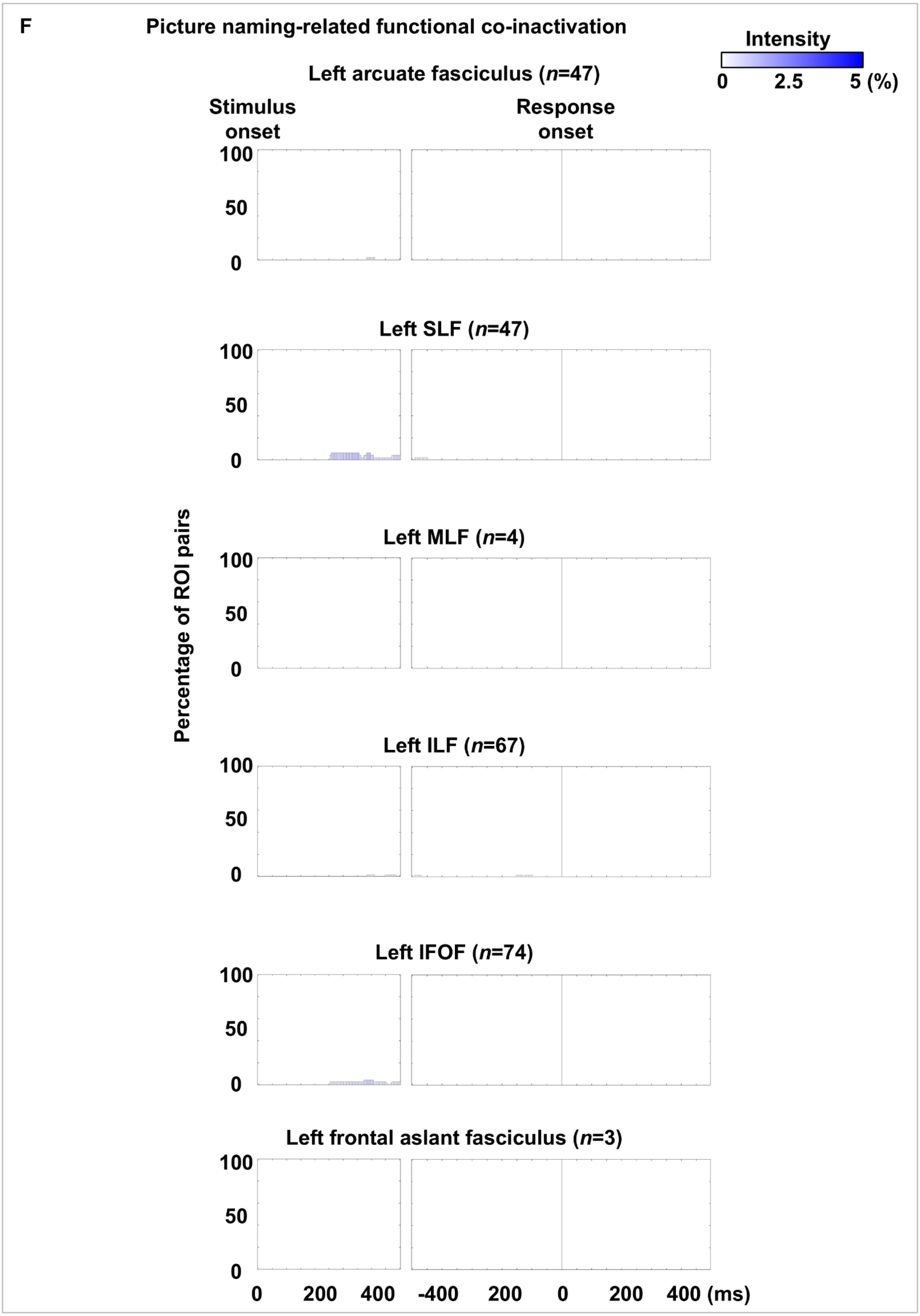
F
Picture naming-related functional co-inactivation
Intensity
0
2.5
5 (%)
Left arcuate fasciculus (n=47)
Stimulus onset
Response onset
Left SLF (n=47)
Left MLF (n=4)
Left ILF (n=67)
Left IFOF (n=74)
Left frontal aslant fasciculus (n=3)
Percentage of ROI pairs
100
50
0
0
200
400
-400
200
0
200
400 (ms)

eFigure 7

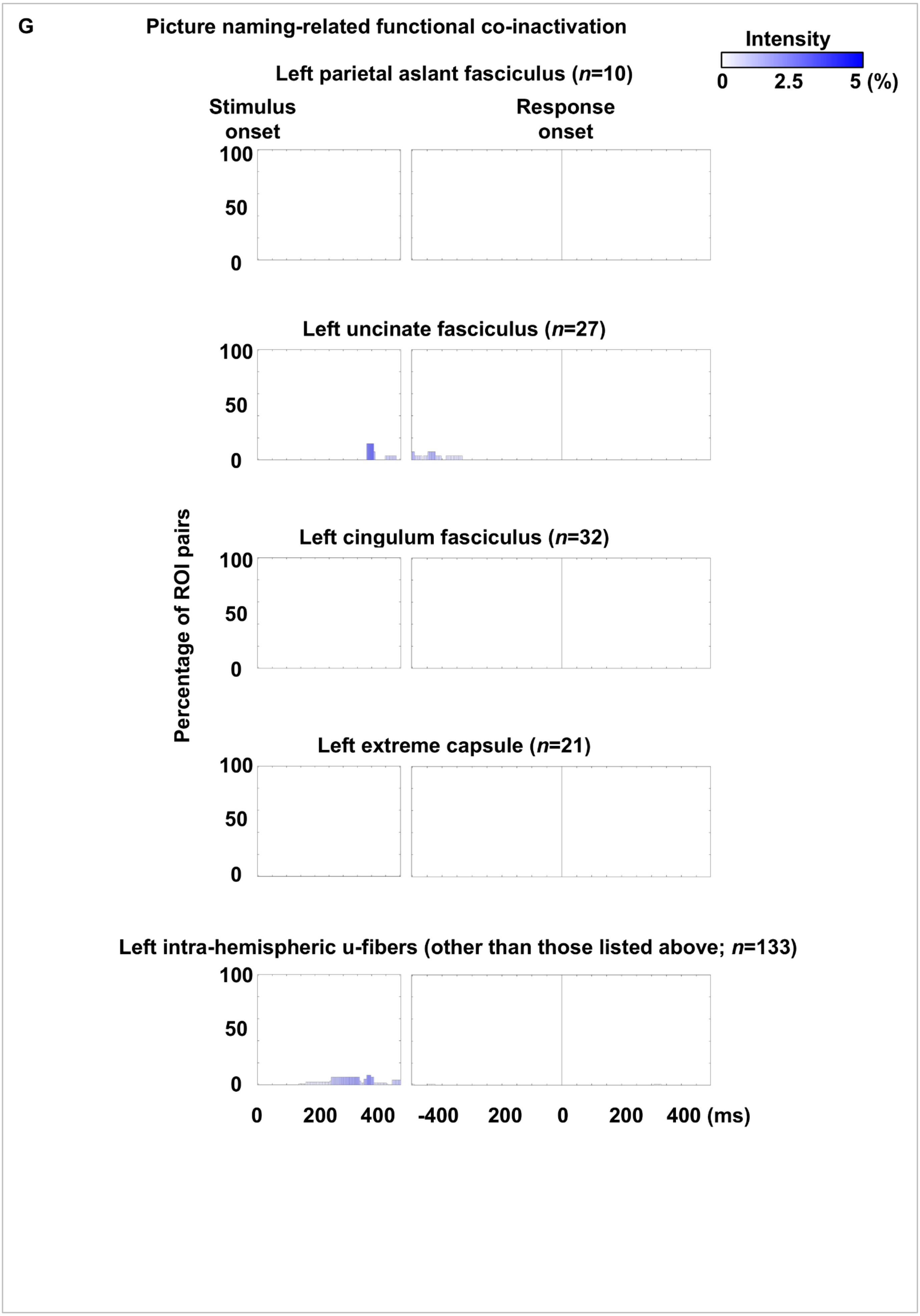
G
Picture naming-related functional co-inactivation
Intensity
0
2.5
5 (%)
Left parietal aslant fasciculus (n=10)
Stimulus onset
Response onset
Left uncinate fasciculus (n=27)
Left cingulum fasciculus (n=32)
Left extreme capsule (n=21)
Left intra-hemispheric u-fibers (other than those listed above; n=133)
Percentage of ROI pairs
100
50
0
0
200
400
-400
200
0
200
400 (ms)

eFigure 7

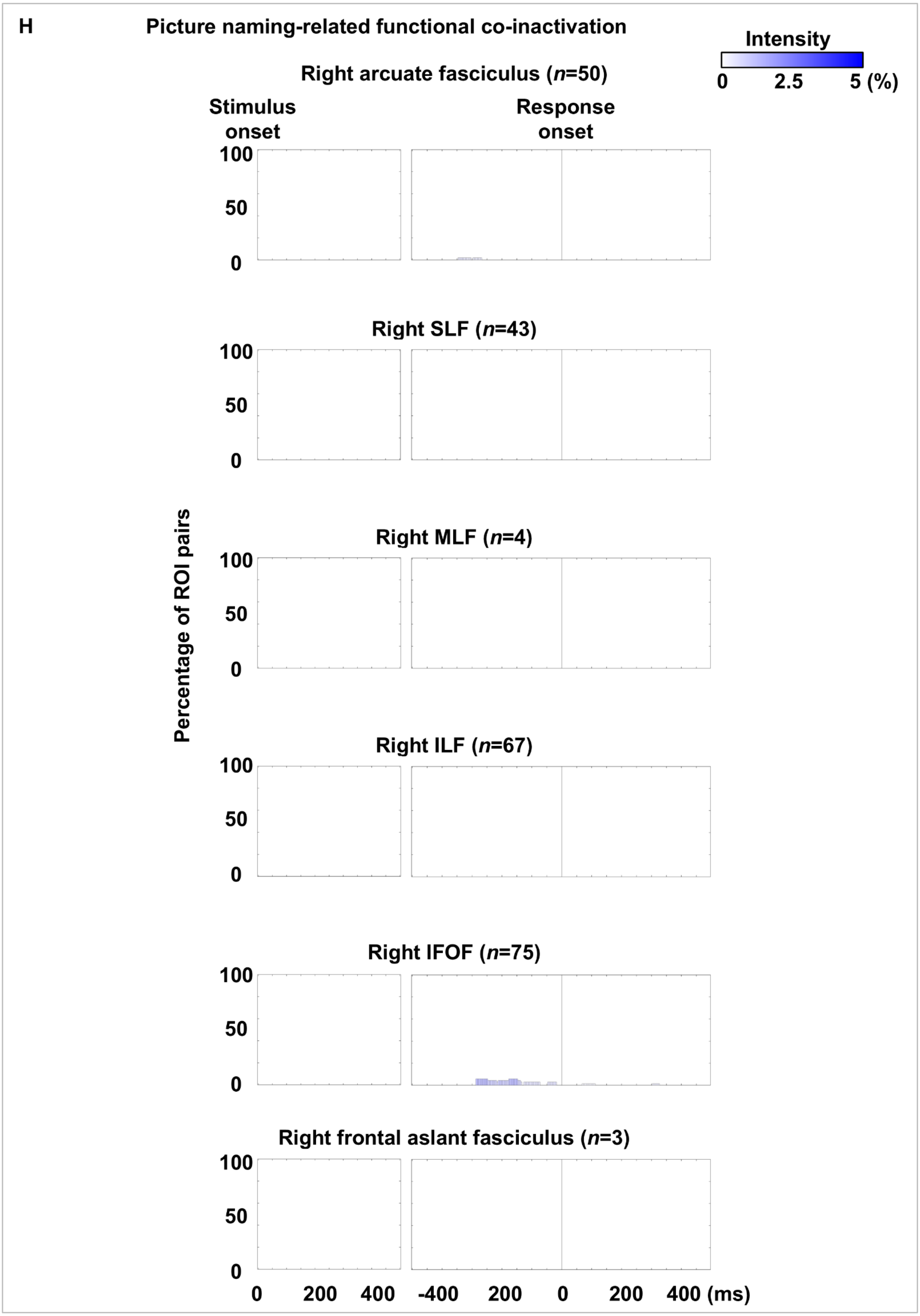
H
Picture naming-related functional co-inactivation
Intensity
0
2.5
5 (%)
Right arcuate fasciculus (n=50)
Stimulus onset
Response onset
100
50
0
Right SLF (n=43)
Right MLF (n=4)
Right ILF (n=67)
Right IFOF (n=75)
Right frontal aslant fasciculus (n=3)
Percentage of ROI pairs
0
200
400
-400
200
0
200
400 (ms)

eFigure 7

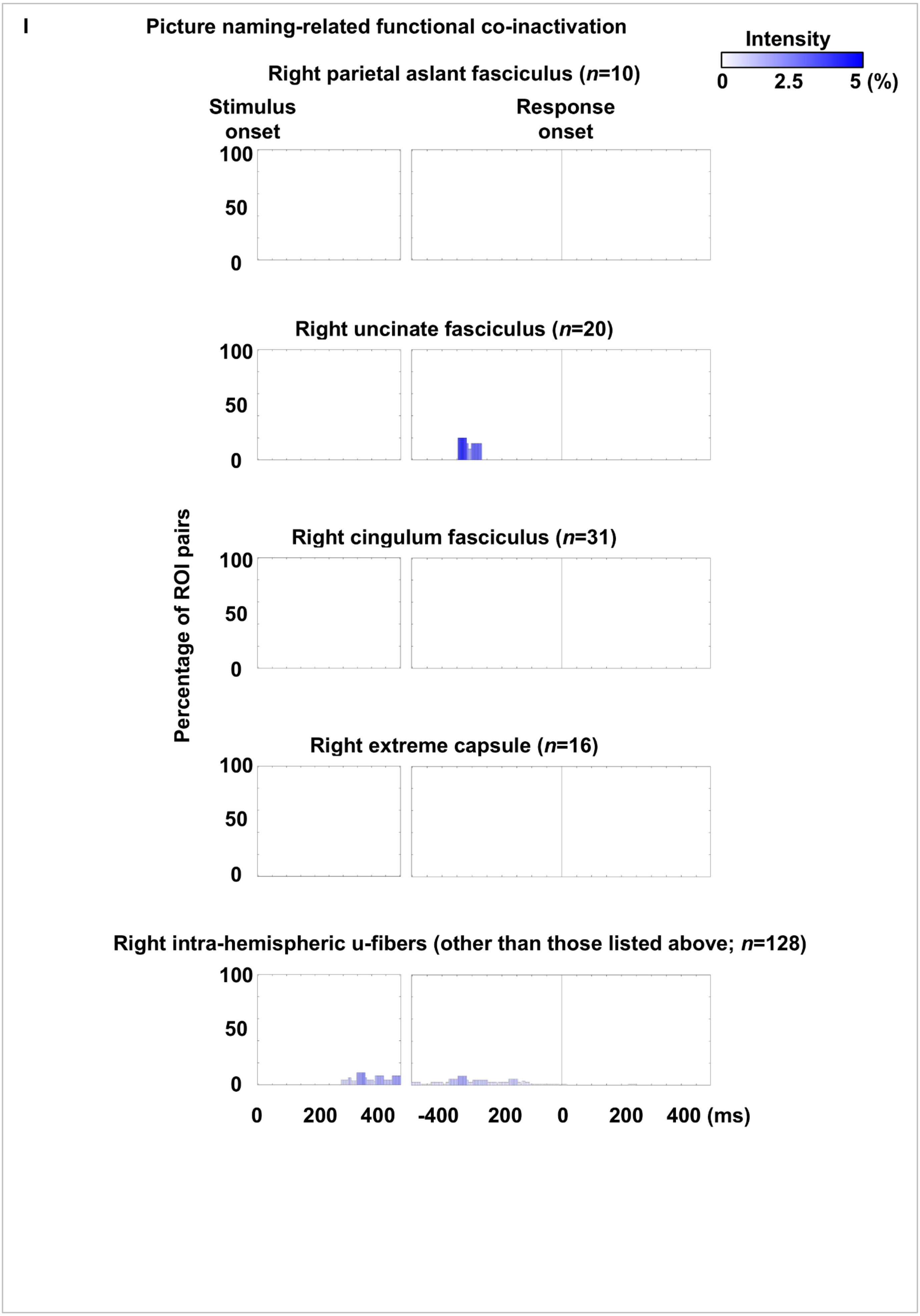
I
Picture naming-related functional co-inactivation
Intensity
0
2.5
5 (%)
Right parietal aslant fasciculus (n=10)
Stimulus onset
Response onset
100
50
0
Right uncinate fasciculus (n=20)
Percentage of ROI pairs
Right cingulum fasciculus (n=31)
Right extreme capsule (n=16)
Right intra-hemispheric u-fibers (other than those listed above; n=128)
0
200
400
-400
200
0
200
400 (ms)

eFigure 7

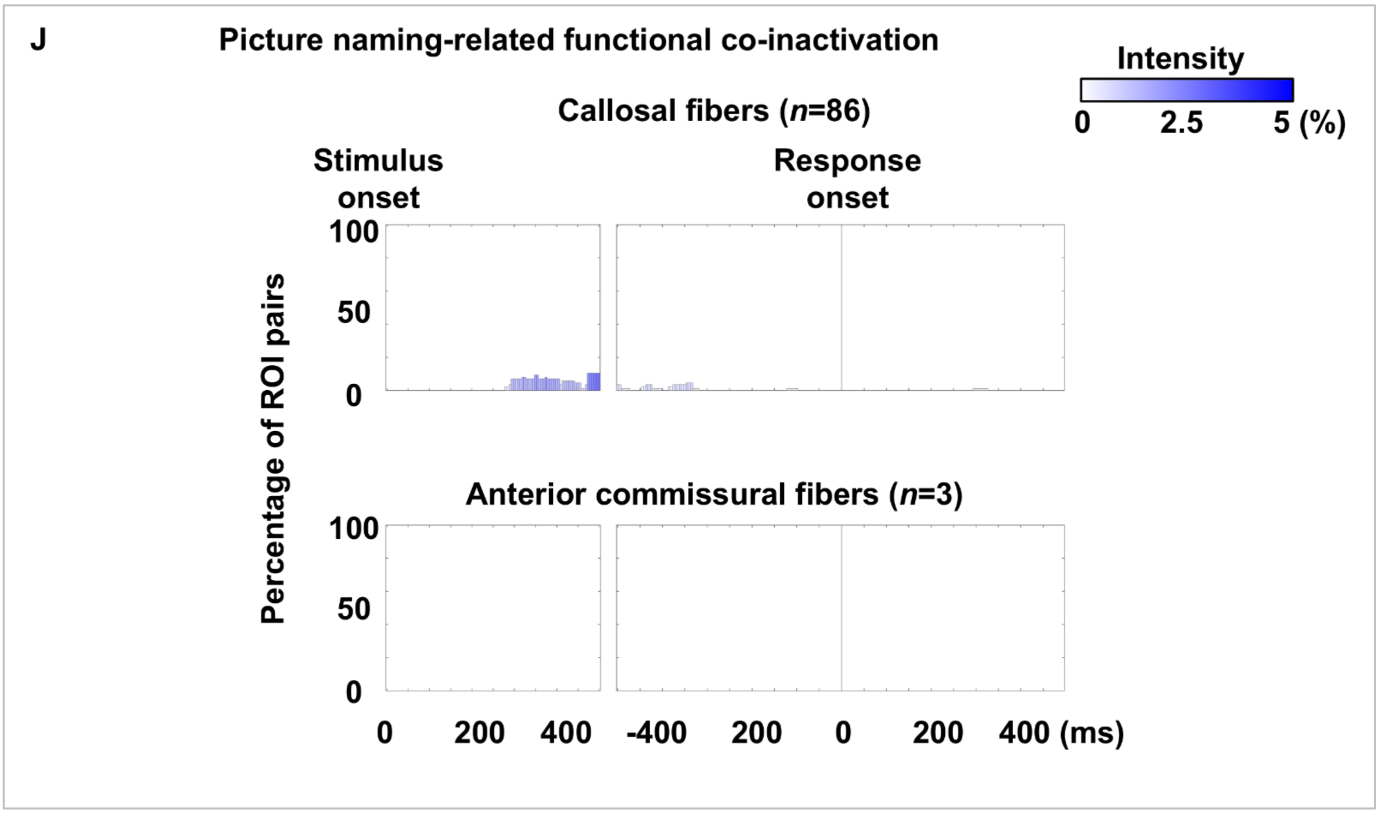


**eFigure 7. Auditory and picture naming-related functional co-inactivation through each fasciculus.** The bar height indicates the proportion of functional co-inactivation within each fasciculus during a given time bin, while the bar color represents the average intensity of functional co-inactivation. **A-E** Auditory naming. **F-J** Picture naming. **A, B, F, G** Left intra-hemispheric pathways. **C, D, H, I** Right intra-hemispheric pathways. **E, J** Inter-hemispheric pathways.

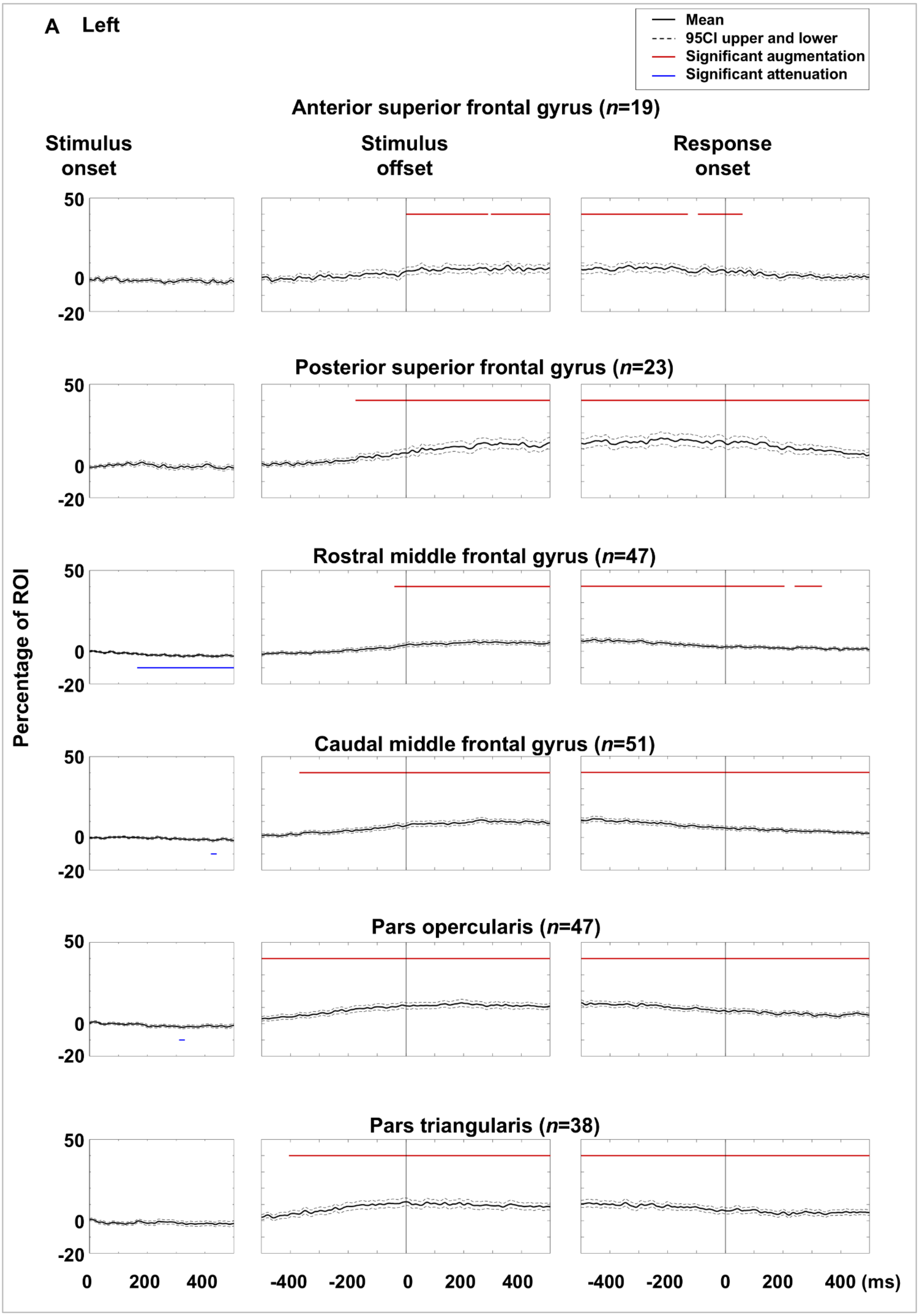
A Left
Mean
95CI upper and lower
Significant augmentation
Significant attenuation
Anterior superior frontal gyrus (n=19)
Stimulus onset
Stimulus offset
Response onset
Posterior superior frontal gyrus (n=23)
Rostral middle frontal gyrus (n=47)
Caudal middle frontal gyrus (n=51)
Pars opercularis (n=47)
Pars triangularis (n=38)
Percentage of ROI
50
0
-20
0
200
400
-400
-200
400 (ms)

eFigure 8

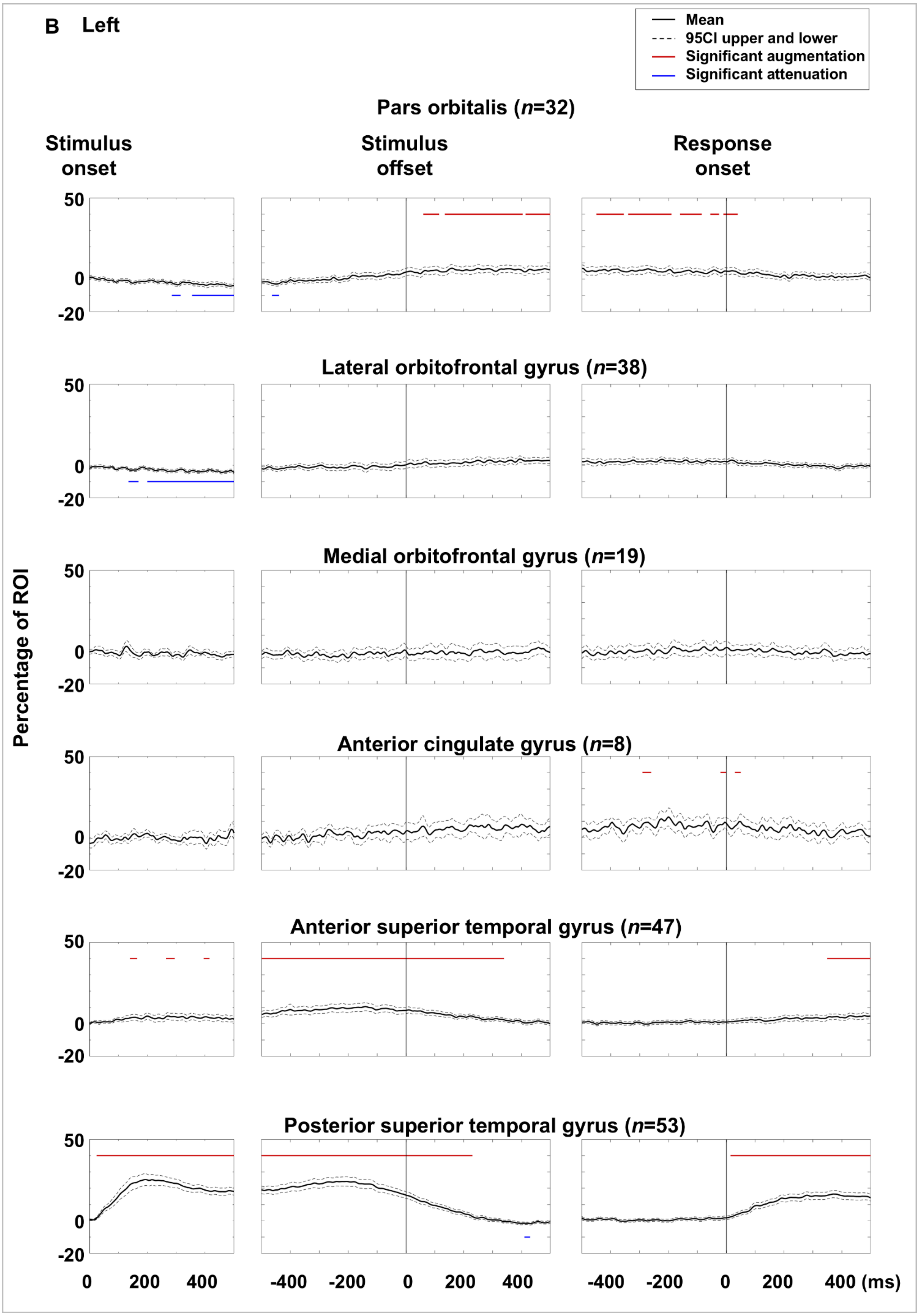
B Left
Mean
95CI upper and lower
Significant augmentation
Significant attenuation
Pars orbitalis (n=32)
Stimulus onset
Stimulus offset
Response onset
Lateral orbitofrontal gyrus (n=38)
Medial orbitofrontal gyrus (n=19)
Anterior cingulate gyrus (n=8)
Anterior superior temporal gyrus (n=47)
Posterior superior temporal gyrus (n=53)
Percentage of ROI
50
0
-20
0
200
400
-400
-200
0
200
400
-400
-200
0
200
400 (ms)

eFigure 8

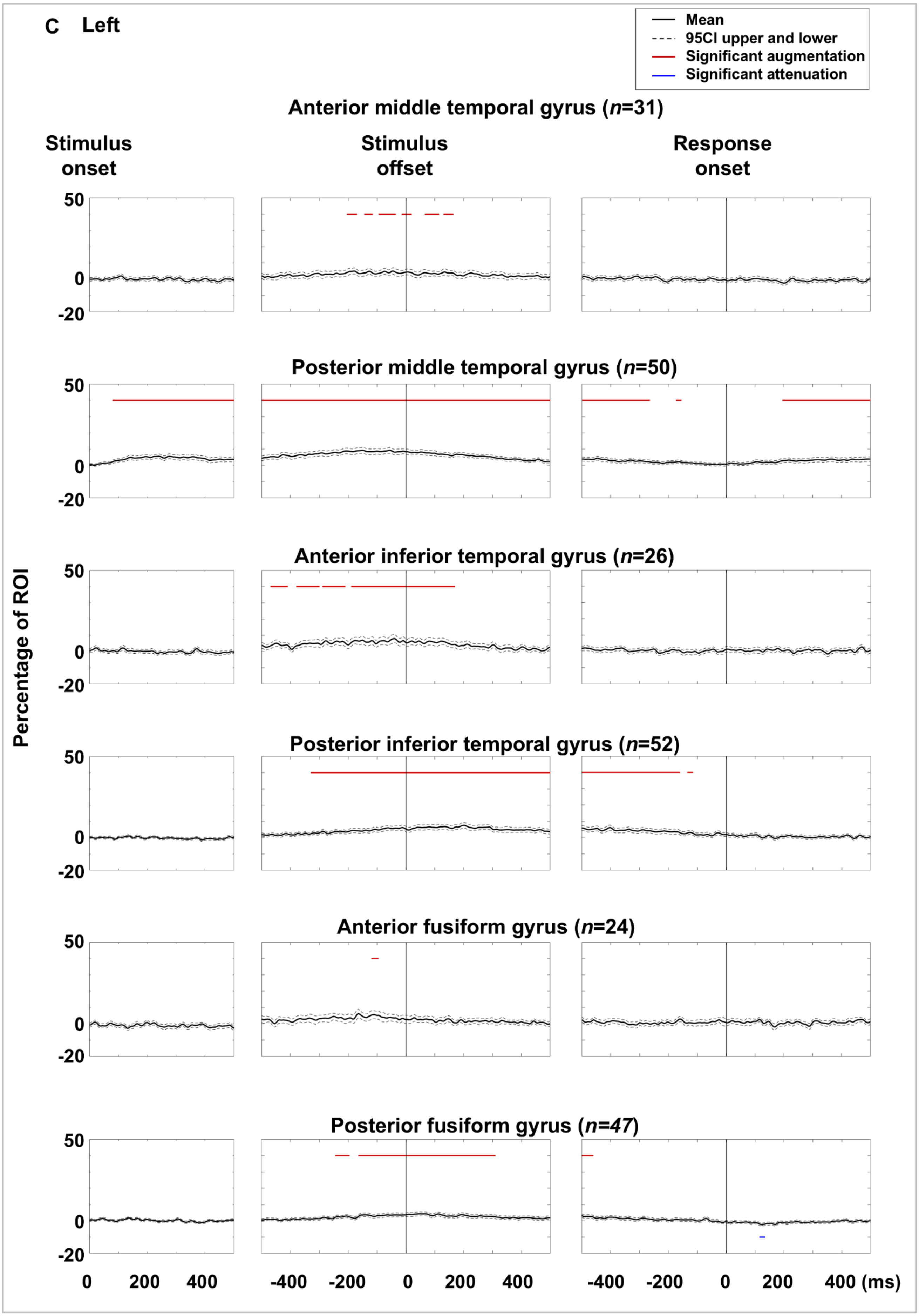
C Left
Mean
95CI upper and lower
Significant augmentation
Significant attenuation
Anterior middle temporal gyrus (n=31)
Stimulus onset
Stimulus offset
Response onset
Posterior middle temporal gyrus (n=50)
Anterior inferior temporal gyrus (n=26)
Posterior inferior temporal gyrus (n=52)
Anterior fusiform gyrus (n=24)
Posterior fusiform gyrus (n=47)
Percentage of ROI
50
0
-20
0 200 400
-400 -200 0 200 400
-400 -200 0 200 400 (ms)

eFigure 8

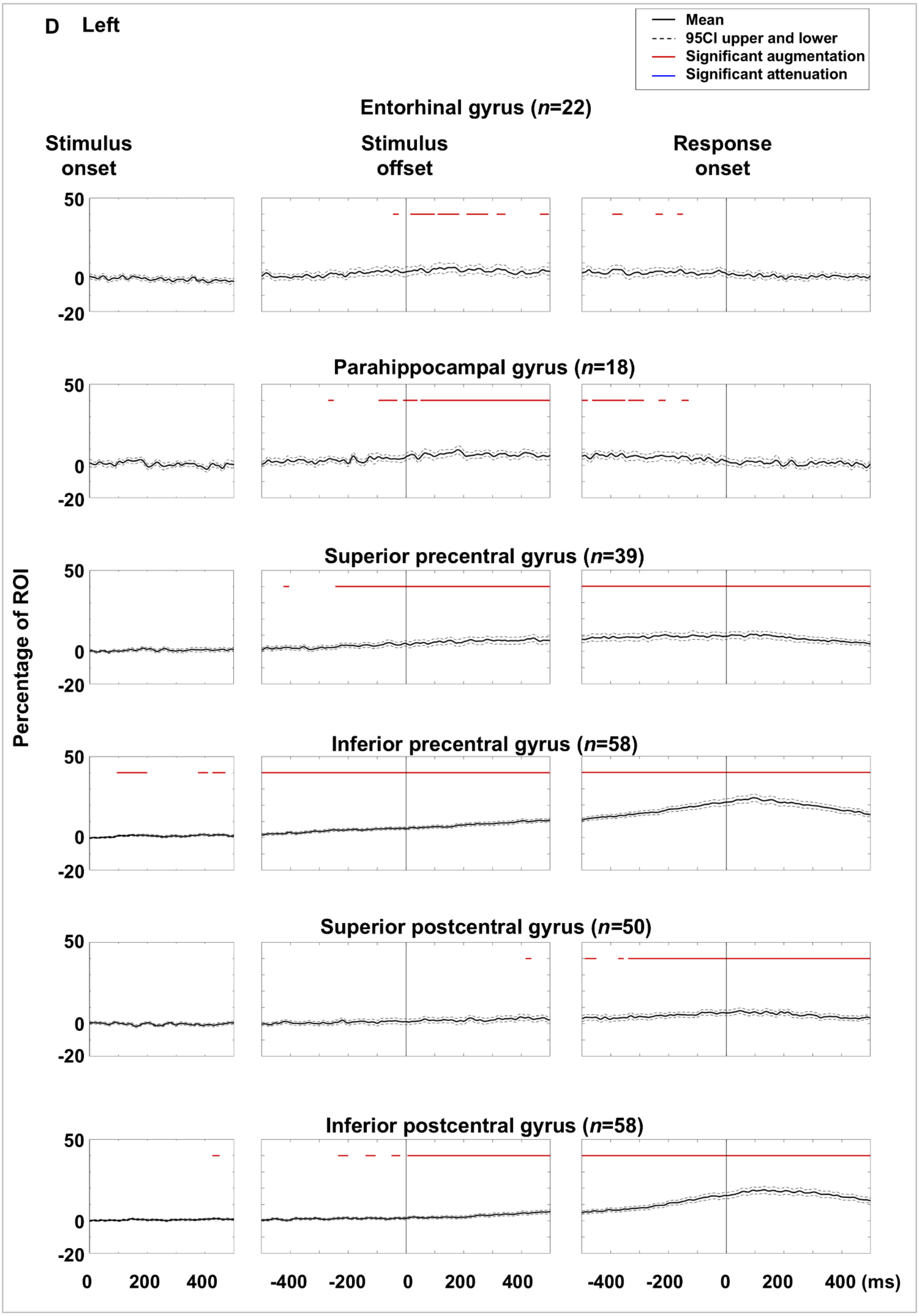
D Left
Mean
95CI upper and lower
Significant augmentation
Significant attenuation
Entorhinal gyrus (n=22)
Stimulus onset
Stimulus offset
Response onset
Parahippocampal gyrus (n=18)
Superior precentral gyrus (n=39)
Inferior precentral gyrus (n=58)
Superior postcentral gyrus (n=50)
Inferior postcentral gyrus (n=58)
Percentage of ROI
50
0
-20
0
200
400
-400
-200
400 (ms)

eFigure 8

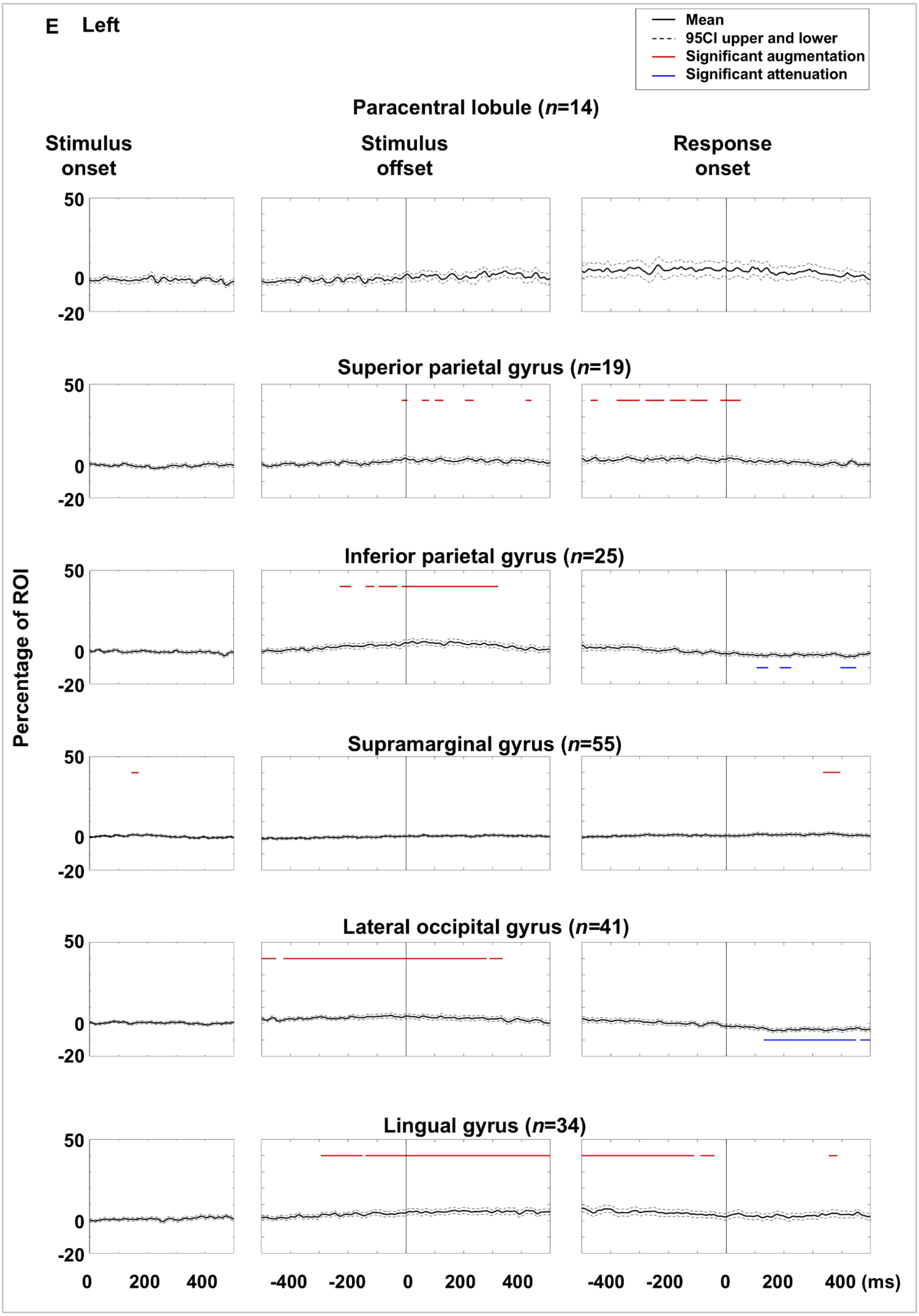
E Left
Mean
95CI upper and lower
Significant augmentation
Significant attenuation
Paracentral lobule (n=14)
Stimulus onset
Stimulus offset
Response onset
Superior parietal gyrus (n=19)
Inferior parietal gyrus (n=25)
Supramarginal gyrus (n=55)
Lateral occipital gyrus (n=41)
Lingual gyrus (n=34)
Percentage of ROI
50
0
-20
0
200
400
-400
-200
400 (ms)

eFigure 8

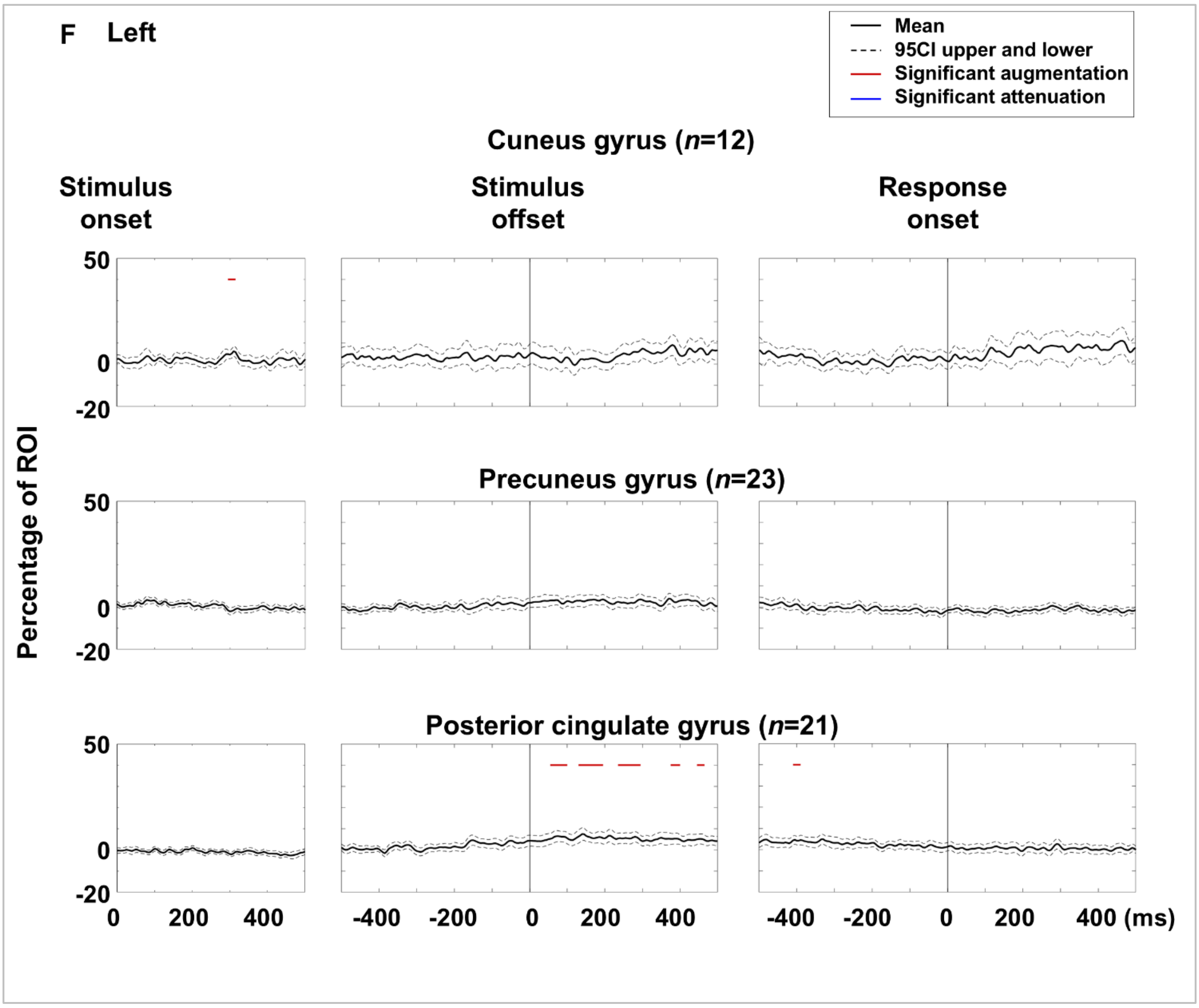
F Left
Mean
95CI upper and lower
Significant augmentation
Significant attenuation
Cuneus gyrus (n=12)
Stimulus onset
Stimulus offset
Response onset
Precuneus gyrus (n=23)
Posterior cingulate gyrus (n=21)
Percentage of ROI
50
0
-20
0
200
400
-400
-200
400 (ms)

**eFigure 8**

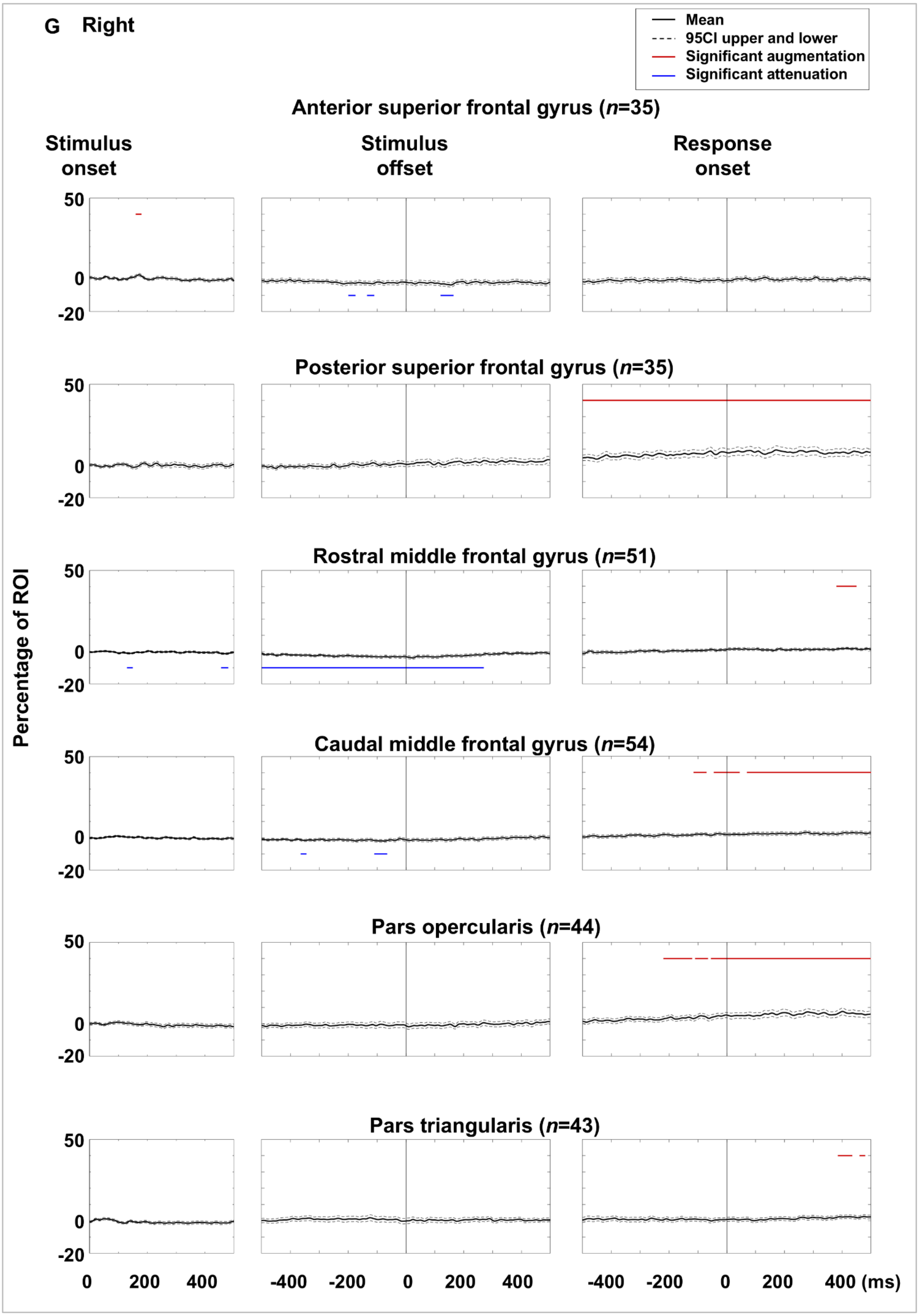
G Right
Mean
95CI upper and lower
Significant augmentation
Significant attenuation
Anterior superior frontal gyrus (n=35)
Stimulus onset
Stimulus offset
Response onset
Posterior superior frontal gyrus (n=35)
Rostral middle frontal gyrus (n=51)
Caudal middle frontal gyrus (n=54)
Pars opercularis (n=44)
Pars triangularis (n=43)
Percentage of ROI
50
0
-20
0 200 400
-400 -200 0 200 400
-400 -200 0 200 400 (ms)

eFigure 8

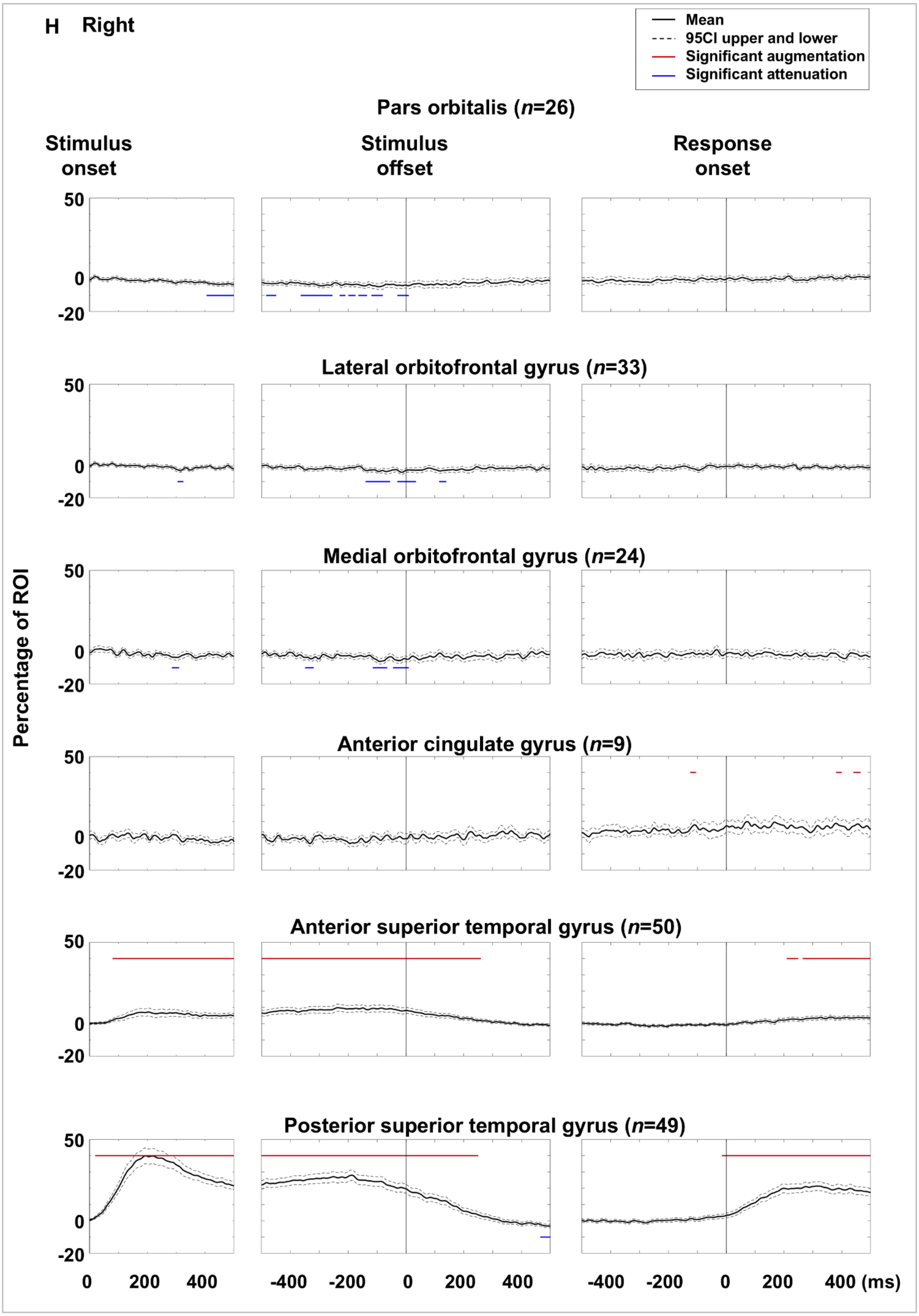
H Right
Mean
95CI upper and lower
Significant augmentation
Significant attenuation
Pars orbitalis (n=26)
Stimulus onset
Stimulus offset
Response onset
Lateral orbitofrontal gyrus (n=33)
Medial orbitofrontal gyrus (n=24)
Anterior cingulate gyrus (n=9)
Anterior superior temporal gyrus (n=50)
Posterior superior temporal gyrus (n=49)
Percentage of ROI
50
0
-20
0 200 400
-400 -200 0 200 400
-400 -200 0 200 400 (ms)

eFigure 8

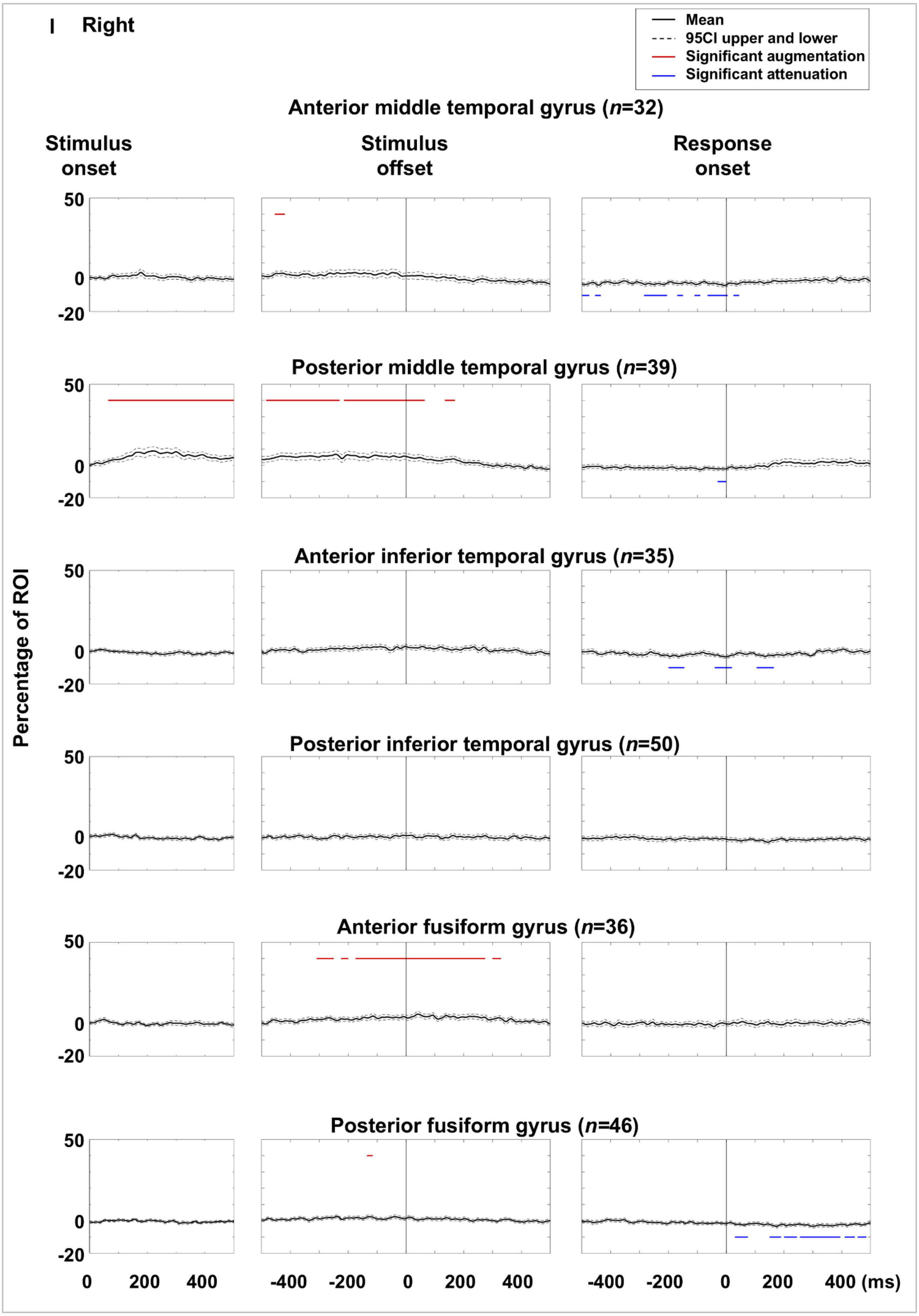
I Right
Mean
95CI upper and lower
Significant augmentation
Significant attenuation
Anterior middle temporal gyrus (n=32)
Stimulus onset
Stimulus offset
Response onset
Posterior middle temporal gyrus (n=39)
Anterior inferior temporal gyrus (n=35)
Posterior inferior temporal gyrus (n=50)
Anterior fusiform gyrus (n=36)
Posterior fusiform gyrus (n=46)
Percentage of ROI
50
0
-20
0 200 400
-400 -200 0 200 400
-400 -200 0 200 400 (ms)

eFigure 8

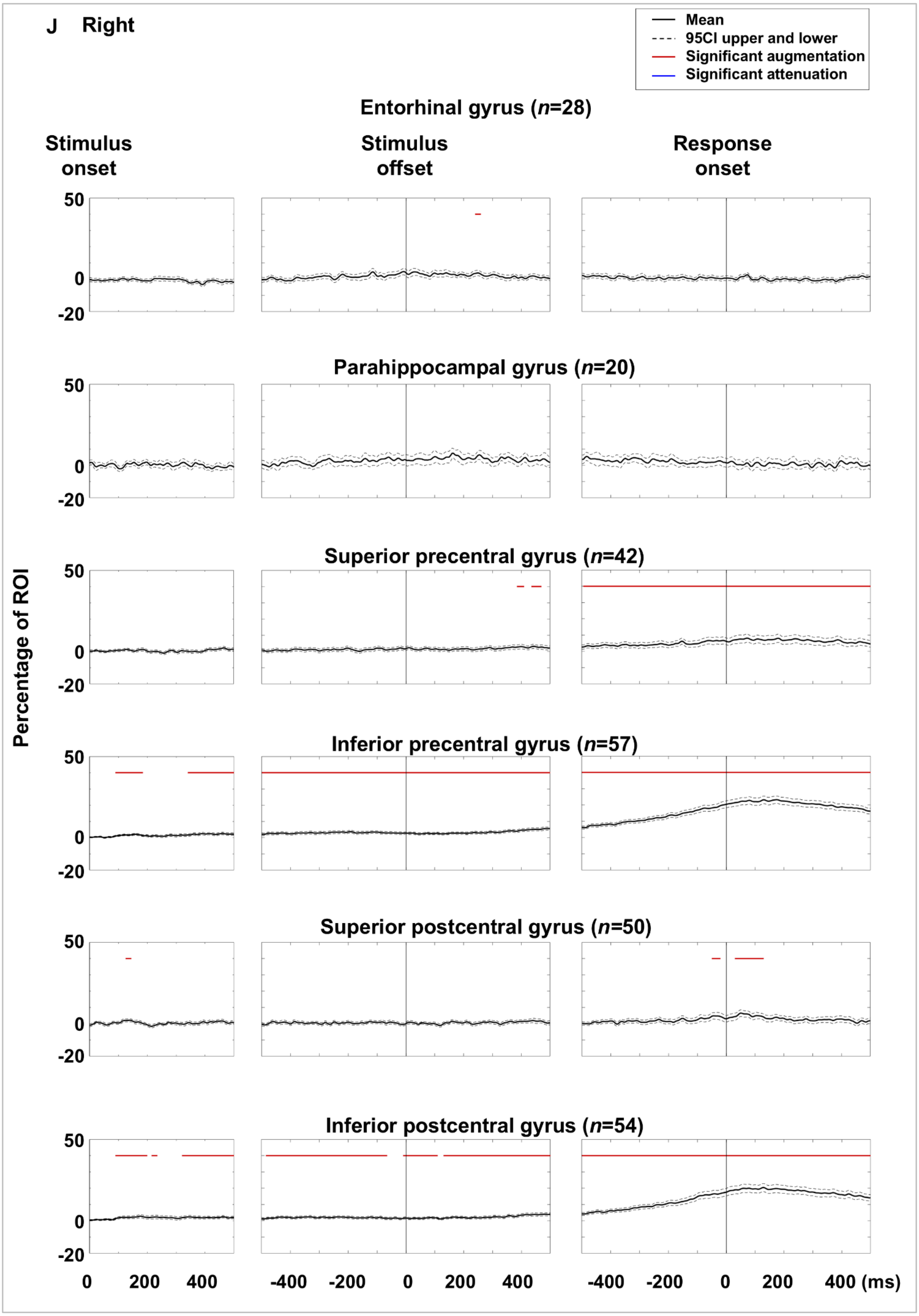
J Right
Mean
95CI upper and lower
Significant augmentation
Significant attenuation
Entorhinal gyrus (n=28)
Stimulus onset
Stimulus offset
Response onset
Parahippocampal gyrus (n=20)
Superior precentral gyrus (n=42)
Inferior precentral gyrus (n=57)
Superior postcentral gyrus (n=50)
Inferior postcentral gyrus (n=54)
Percentage of ROI
50
0
-20
0 200 400
-400 -200 0 200 400
-400 -200 0 200 400 (ms)

**eFigure 8**

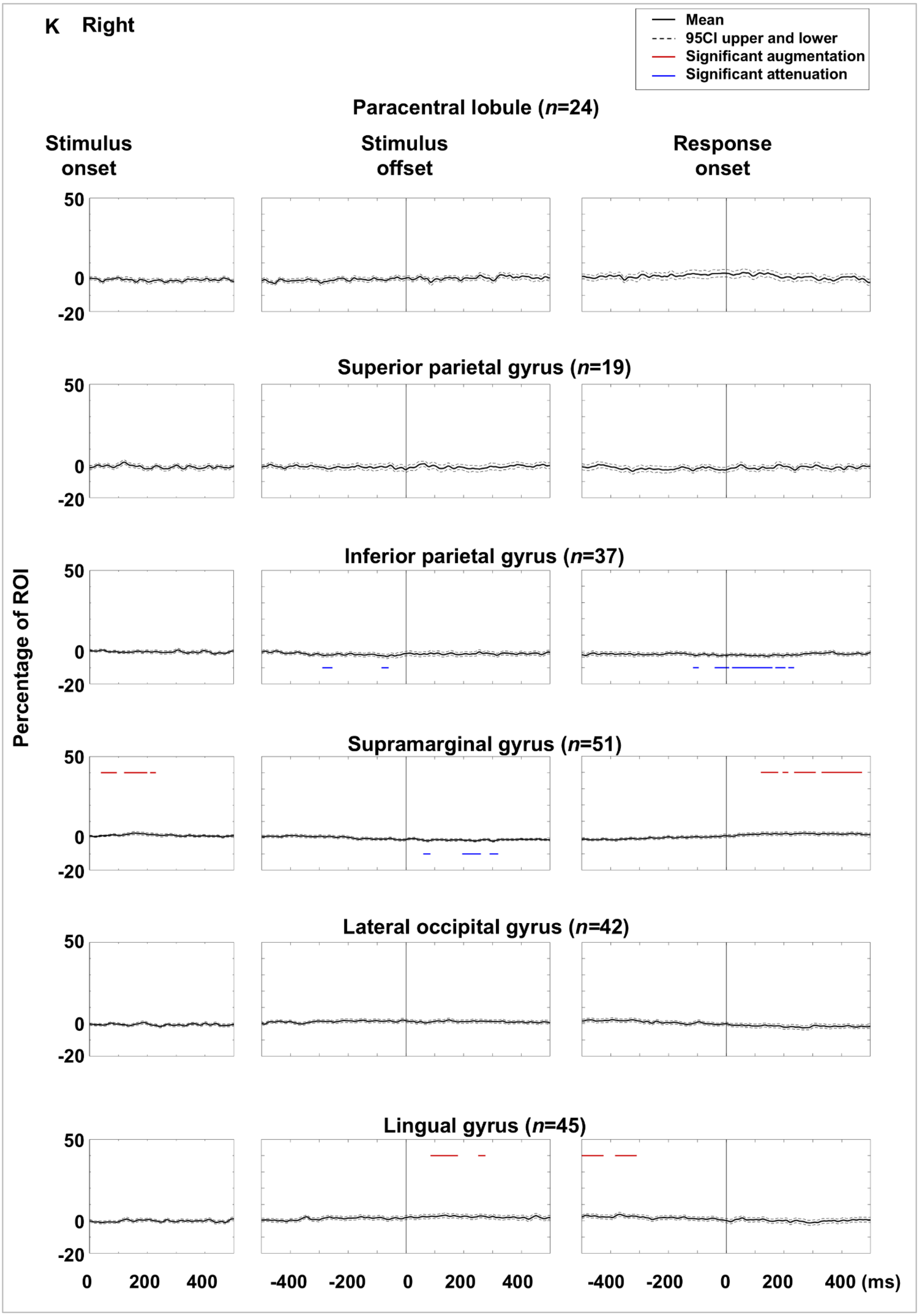

K Right
Mean
95CI upper and lower
Significant augmentation
Significant attenuation
Paracentral lobule (*n*=24)
Stimulus onset
Stimulus offset
Response onset
Superior parietal gyrus (*n*=19)
Inferior parietal gyrus (*n*=37)
Supramarginal gyrus (*n*=51)
Lateral occipital gyrus (*n*=42)
Lingual gyrus (*n*=45)
Percentage of ROI
50
0
-20
0
200
400
-400
-200
400 (ms)


eFigure 8

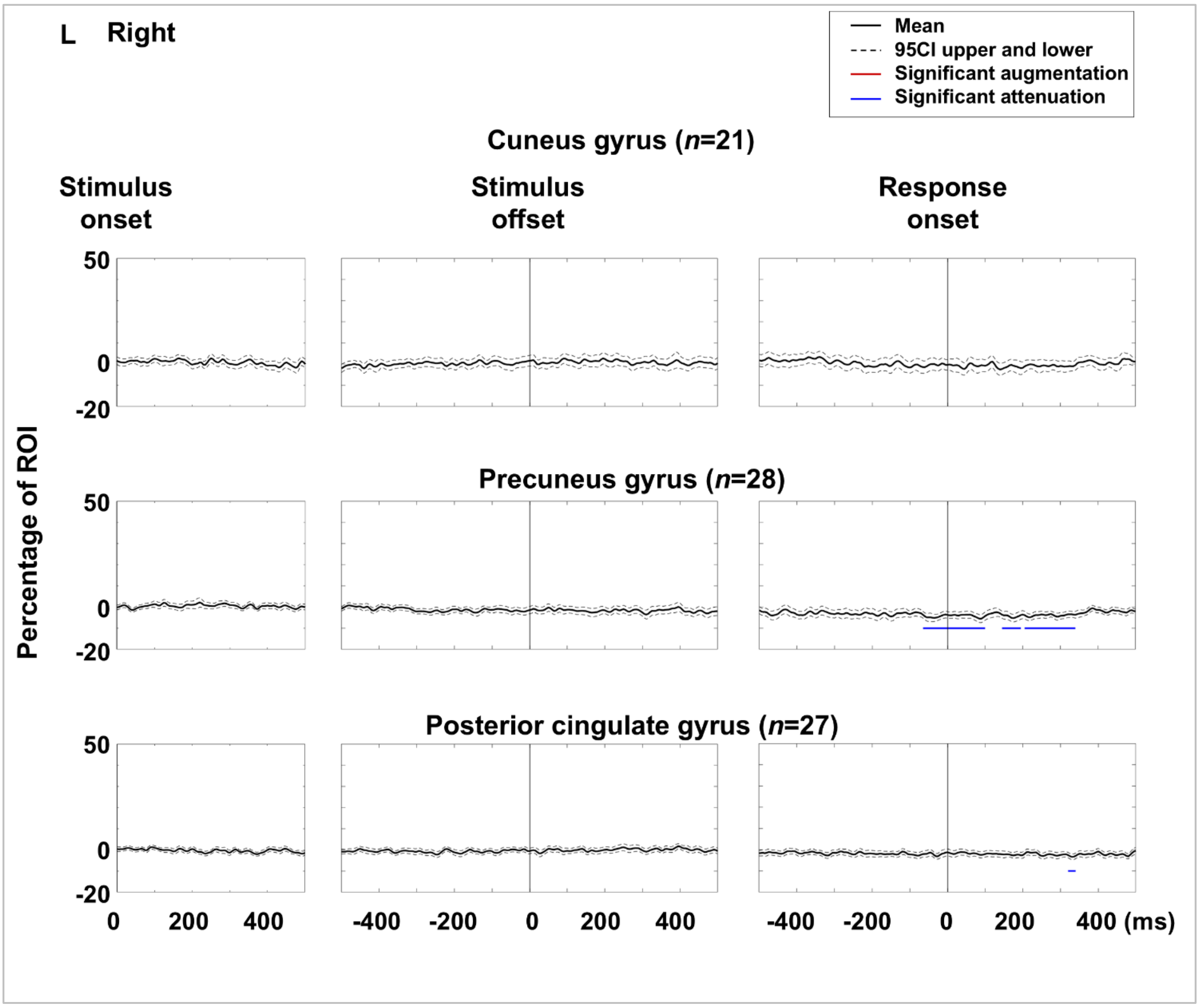


**eFigure 8. Auditory naming-related high-gamma amplitude dynamics at regions of interest (ROIs).** Mean high-gamma amplitude (% change relative to baseline) at each ROI is shown with 95% confidence intervals. **A–F** Left hemispheric ROIs. **G–L** Right hemispheric ROIs.

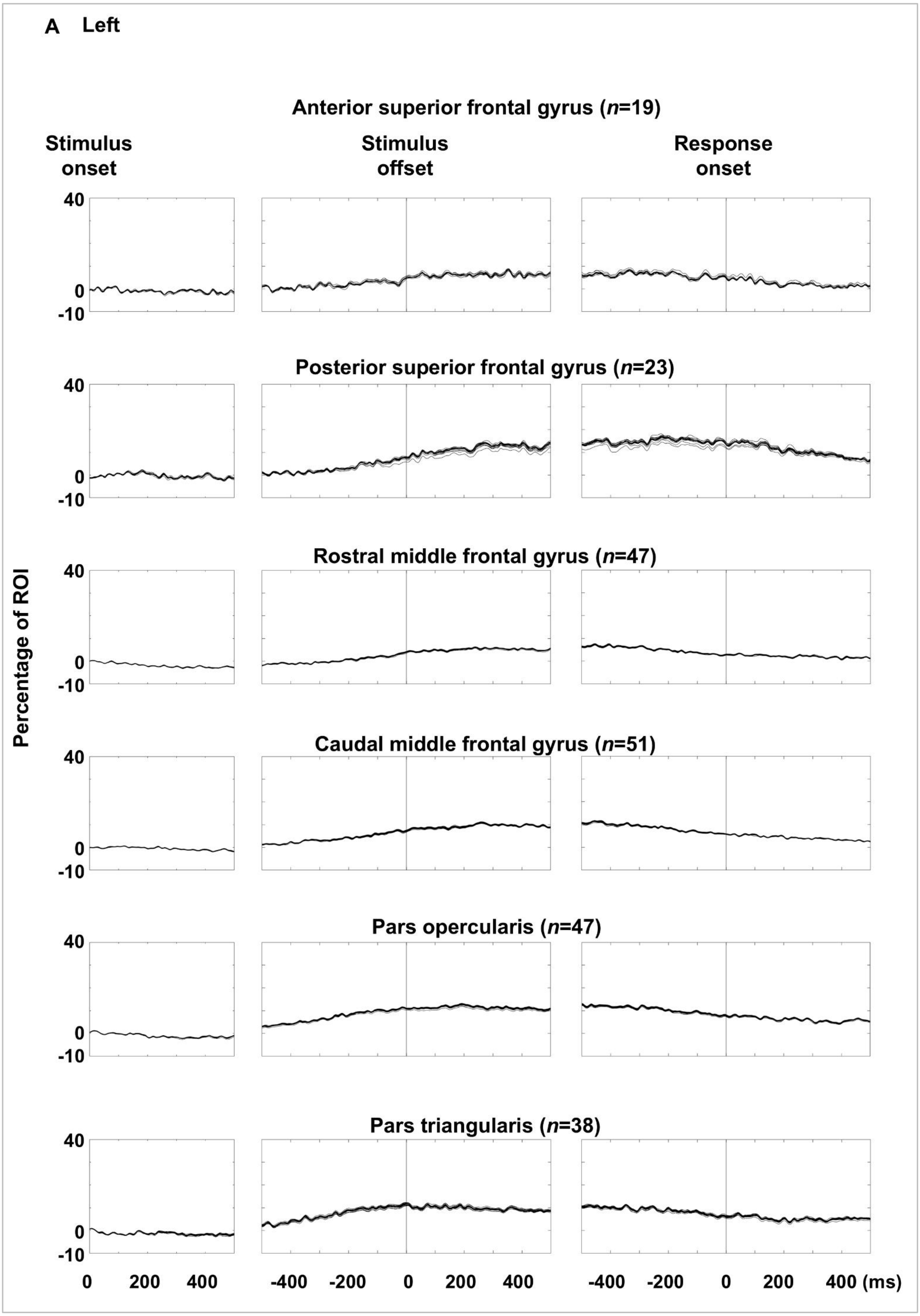

A Left
Anterior superior frontal gyrus (n=19)
Stimulus onset
Stimulus offset
Response onset
Posterior superior frontal gyrus (n=23)
Rostral middle frontal gyrus (n=47)
Caudal middle frontal gyrus (n=51)
Pars opercularis (n=47)
Pars triangularis (n=38)
Percentage of ROI
40
0
-10
0 200 400
-400 -200 0 200 400
-400 -200 0 200 400 (ms)


eFigure 9

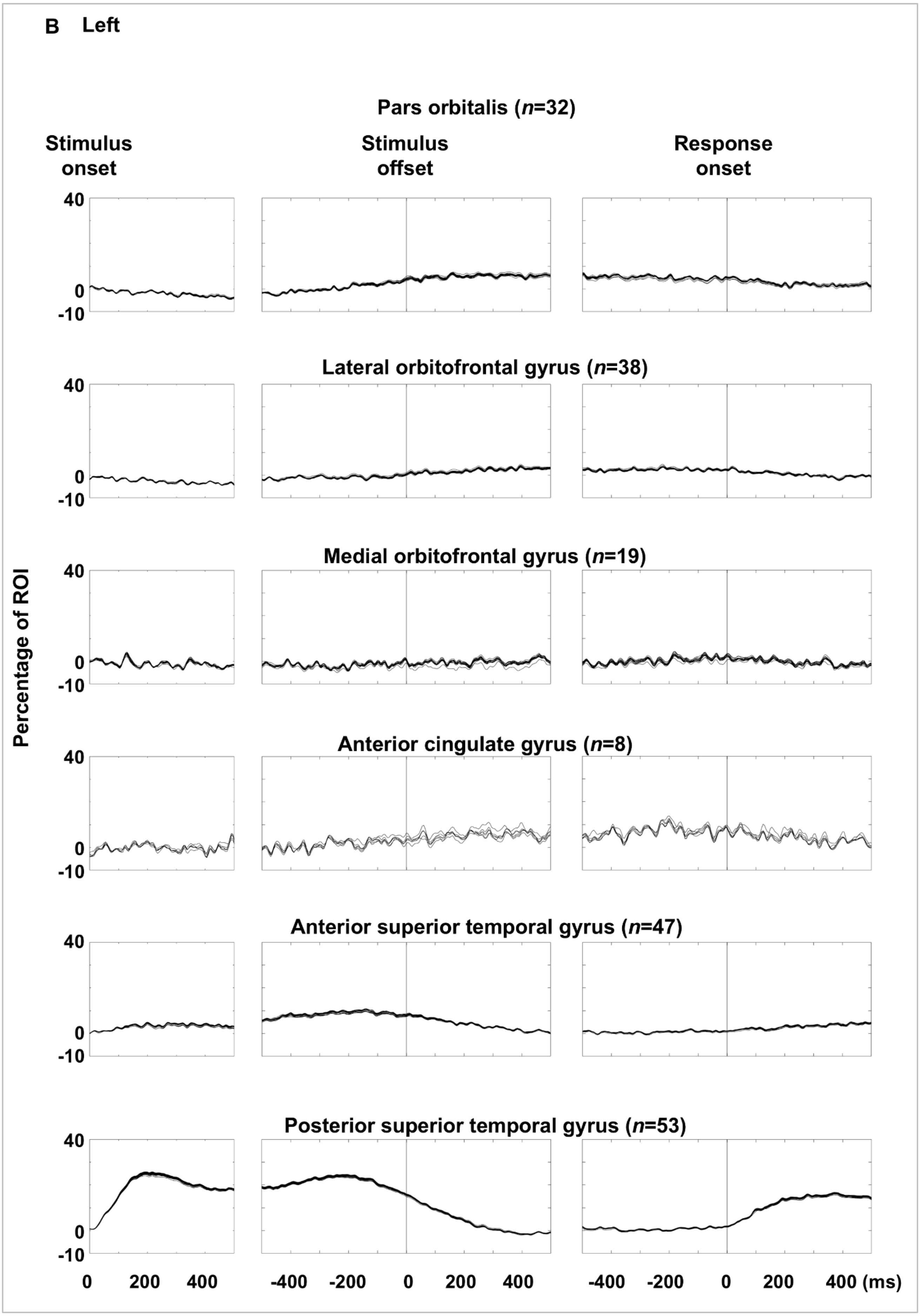
B Left
Pars orbitalis (n=32)
Stimulus onset
Stimulus offset
Response onset
Lateral orbitofrontal gyrus (n=38)
Medial orbitofrontal gyrus (n=19)
Anterior cingulate gyrus (n=8)
Anterior superior temporal gyrus (n=47)
Posterior superior temporal gyrus (n=53)
Percentage of ROI
40
0
-10
0 200 400
-400 -200 0 200 400
-400 -200 0 200 400 (ms)

eFigure 9

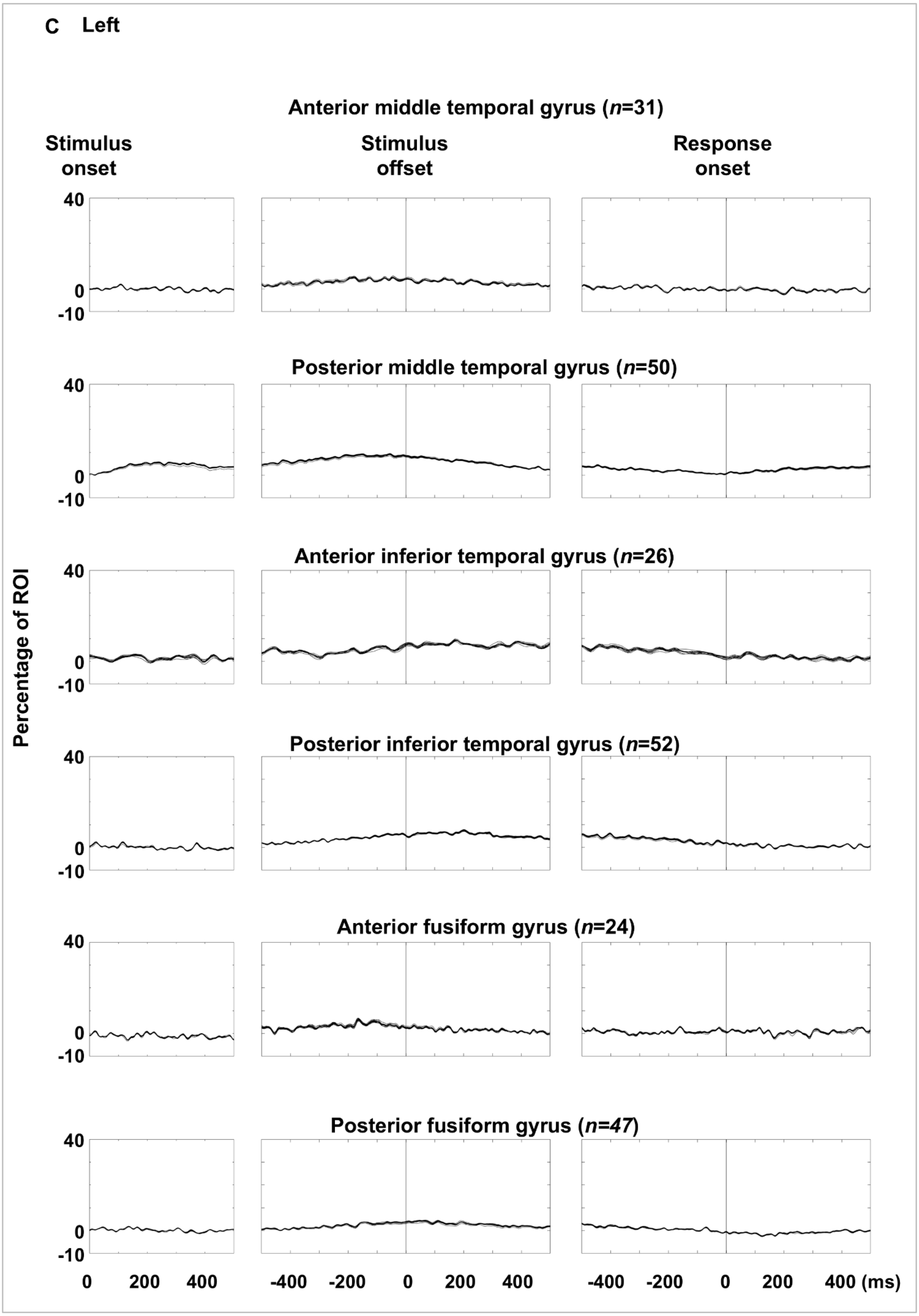
C Left
Anterior middle temporal gyrus (n=31)
Stimulus onset
Stimulus offset
Response onset
Posterior middle temporal gyrus (n=50)
Anterior inferior temporal gyrus (n=26)
Posterior inferior temporal gyrus (n=52)
Anterior fusiform gyrus (n=24)
Posterior fusiform gyrus (n=47)
Percentage of ROI
40
0
-10
0 200 400
-400 -200 0 200 400
-400 -200 0 200 400 (ms)

eFigure 9

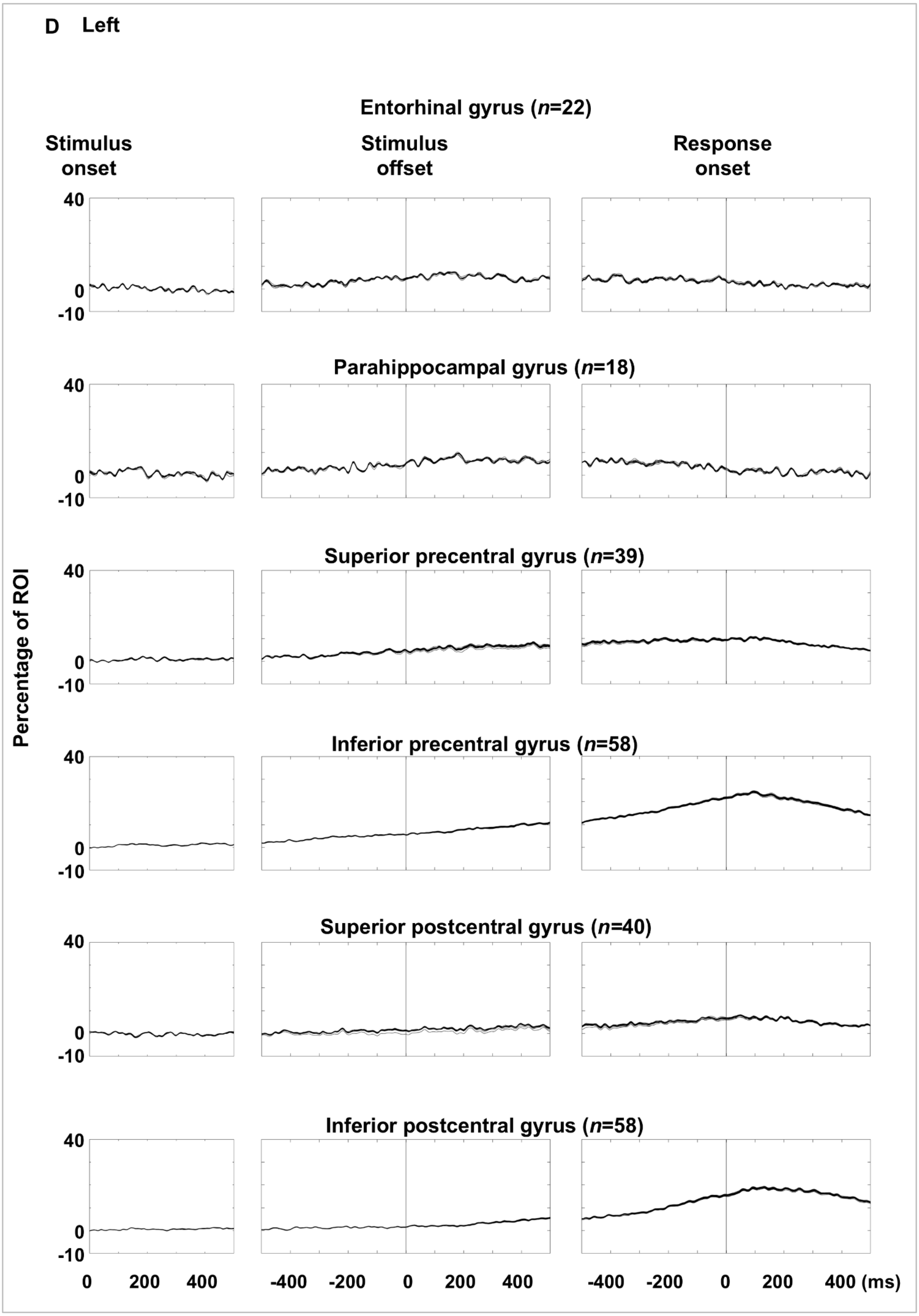
D Left
Entorhinal gyrus (n=22)
Stimulus onset
Stimulus offset
Response onset
Parahippocampal gyrus (n=18)
Superior precentral gyrus (n=39)
Inferior precentral gyrus (n=58)
Superior postcentral gyrus (n=40)
Inferior postcentral gyrus (n=58)
Percentage of ROI
40
0
-10
0
200
400
-400
-200
400 (ms)

eFigure 9

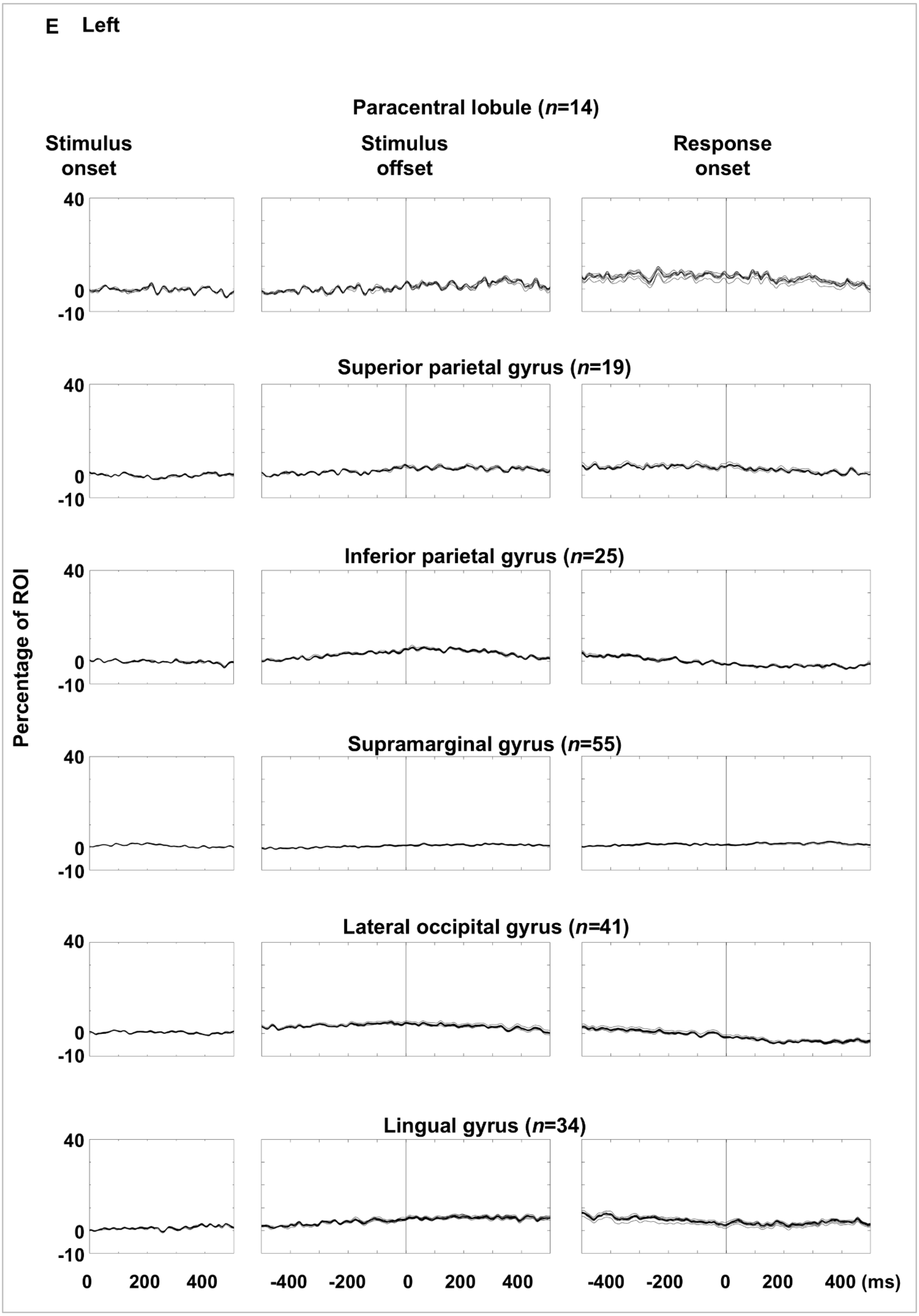
E Left
Paracentral lobule (n=14)
Stimulus onset
Stimulus offset
Response onset
Superior parietal gyrus (n=19)
Inferior parietal gyrus (n=25)
Supramarginal gyrus (n=55)
Lateral occipital gyrus (n=41)
Lingual gyrus (n=34)
Percentage of ROI
40
0
-10
0
200
400
-400
-200
400 (ms)

eFigure 9

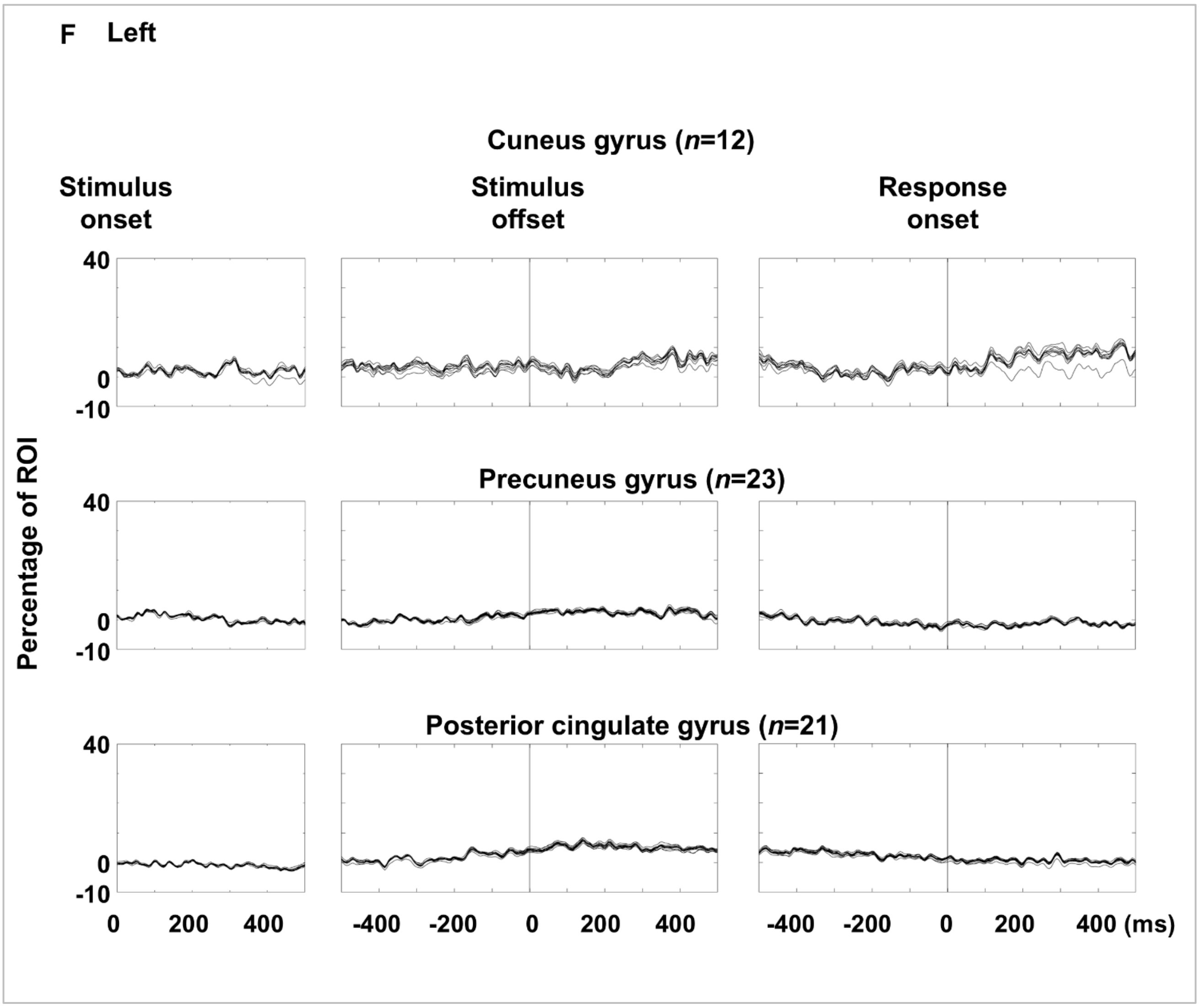
F Left
Cuneus gyrus (n=12)
Stimulus onset
Stimulus offset
Response onset
40
0
-10
Percentage of ROI
Precuneus gyrus (n=23)
Posterior cingulate gyrus (n=21)
0
200
400
-400
-200
400 (ms)

**eFigure 9**

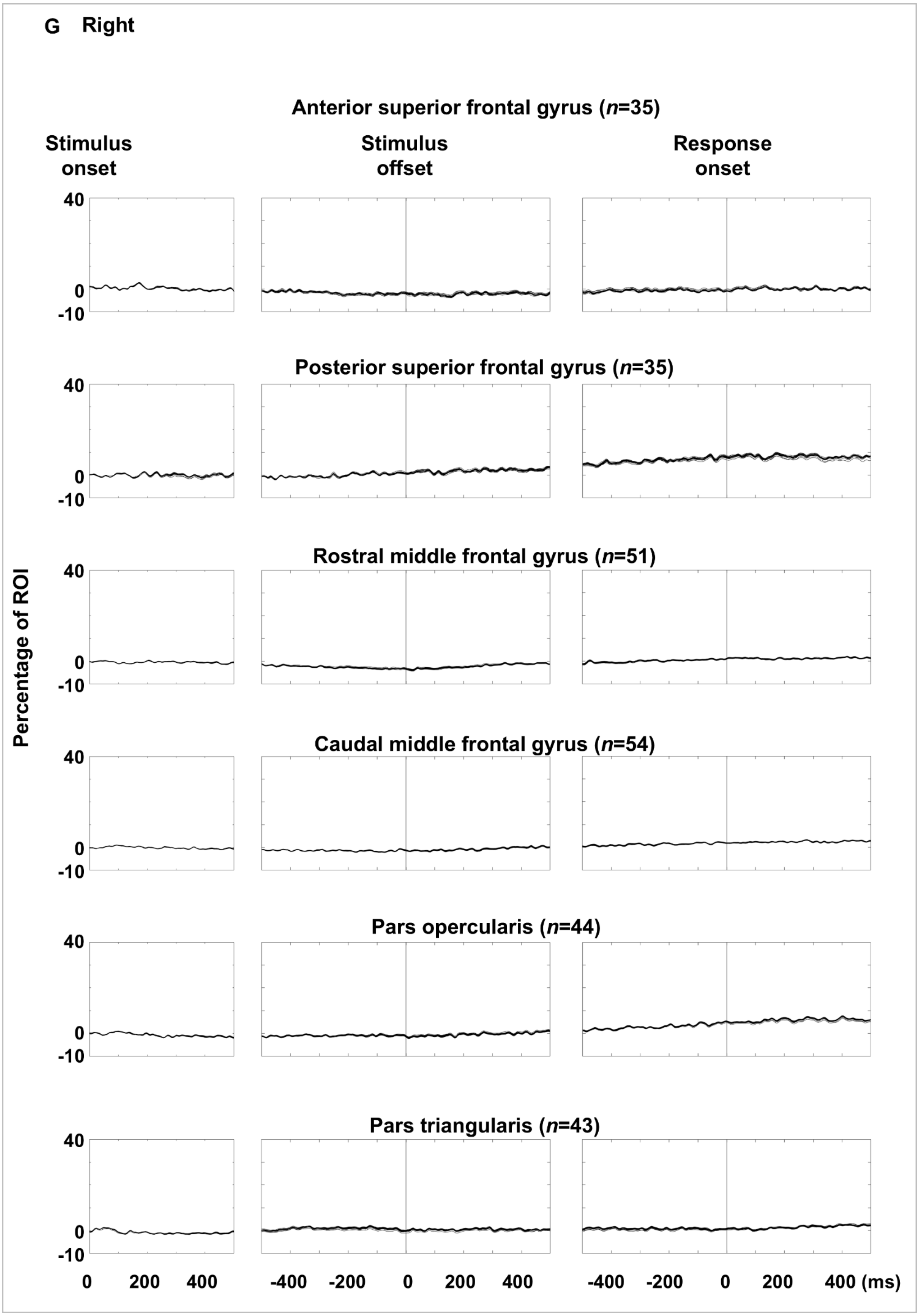
G Right
Anterior superior frontal gyrus (n=35)
Stimulus onset
Stimulus offset
Response onset
Posterior superior frontal gyrus (n=35)
Rostral middle frontal gyrus (n=51)
Caudal middle frontal gyrus (n=54)
Pars opercularis (n=44)
Pars triangularis (n=43)
Percentage of ROI
40
0
-10
0
200
400
-400
-200
400 (ms)

eFigure 9

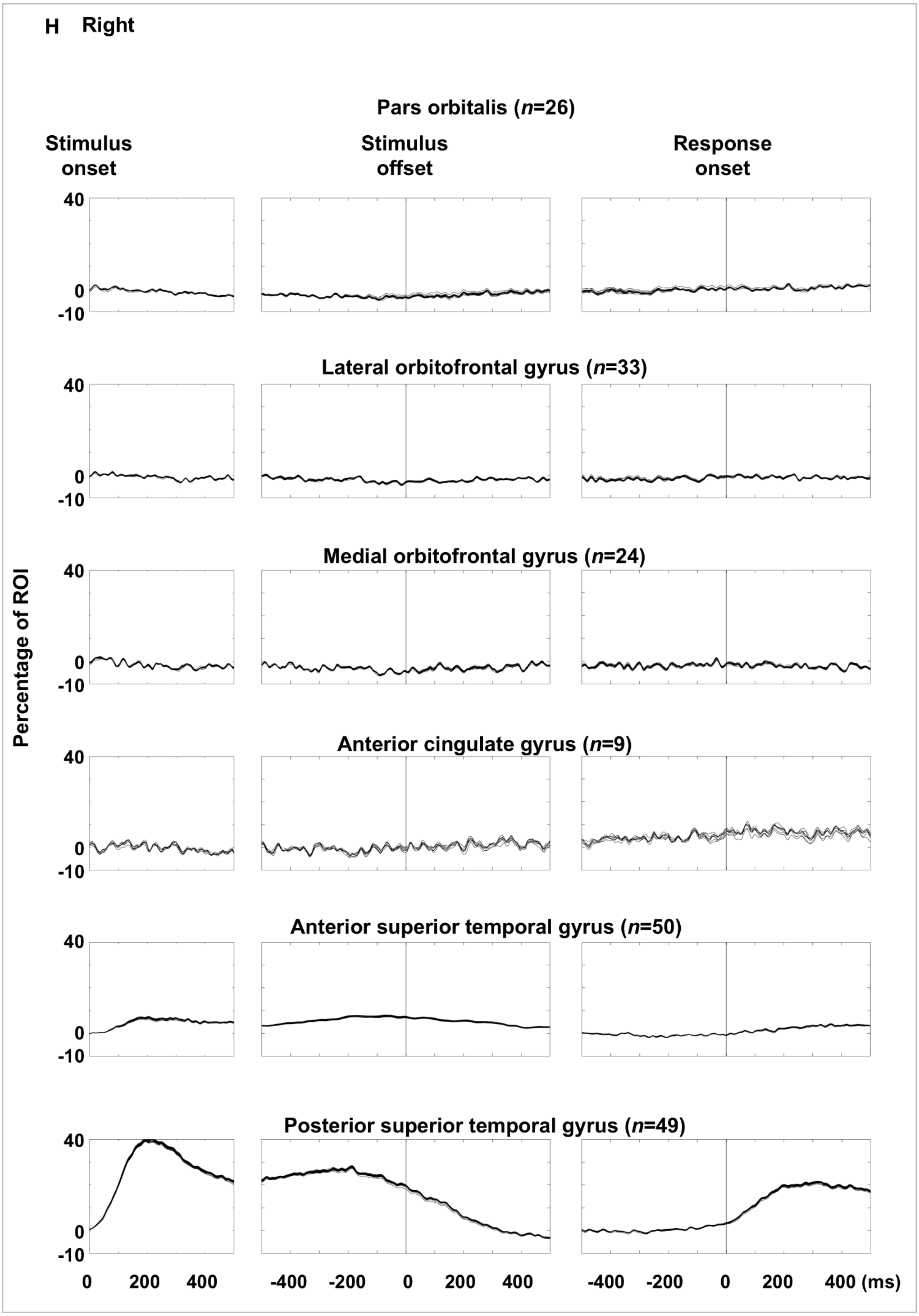
H Right
Pars orbitalis (n=26)
Stimulus onset
Stimulus offset
Response onset
Lateral orbitofrontal gyrus (n=33)
Medial orbitofrontal gyrus (n=24)
Anterior cingulate gyrus (n=9)
Anterior superior temporal gyrus (n=50)
Posterior superior temporal gyrus (n=49)
Percentage of ROI
40
0
-10
0
200
400
-400
-200
400 (ms)

eFigure 9

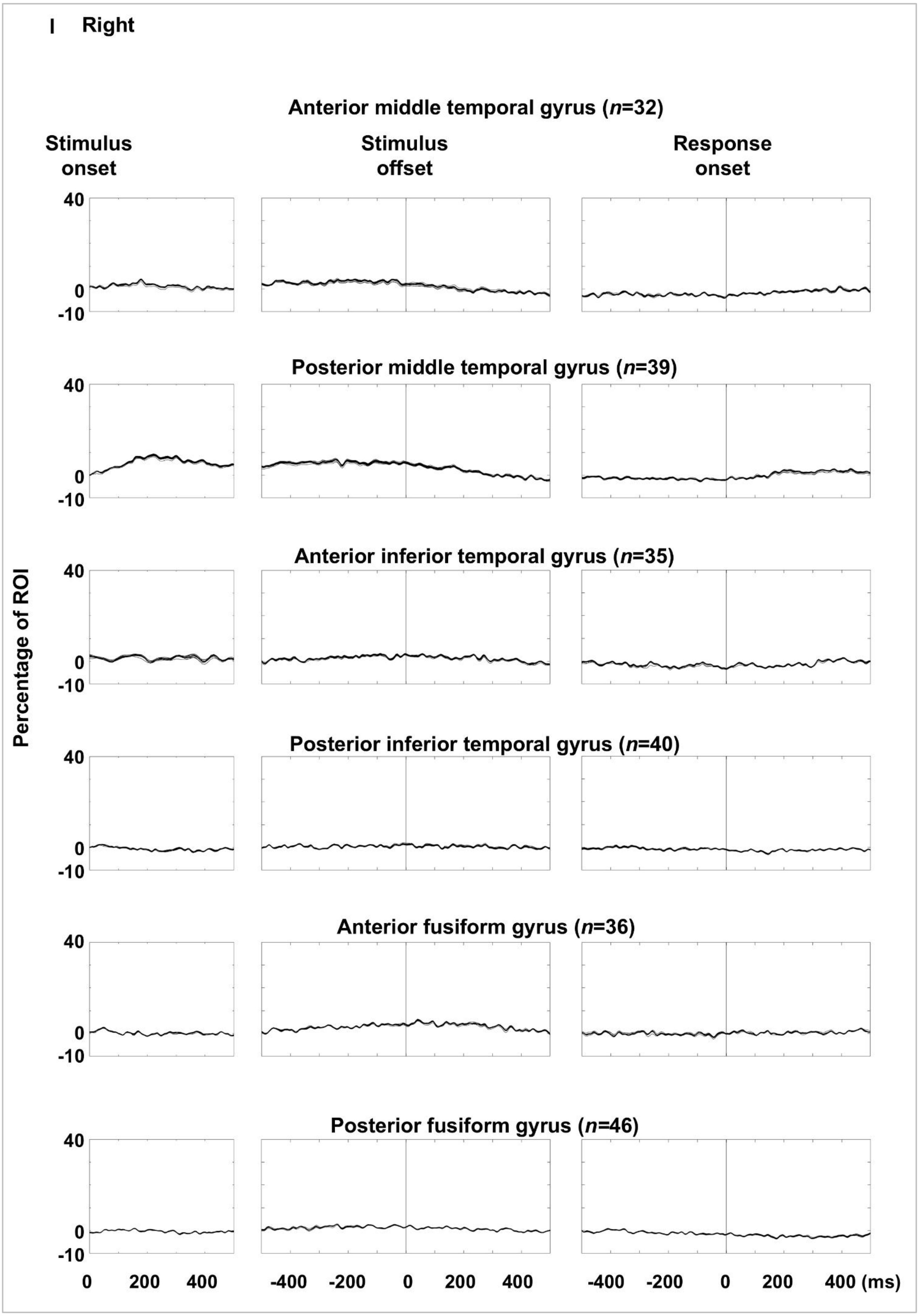
I Right
Anterior middle temporal gyrus (n=32)
Stimulus onset
Stimulus offset
Response onset
Posterior middle temporal gyrus (n=39)
Anterior inferior temporal gyrus (n=35)
Posterior inferior temporal gyrus (n=40)
Anterior fusiform gyrus (n=36)
Posterior fusiform gyrus (n=46)
Percentage of ROI
40
0
-10
0 200 400
-400 -200 0 200 400
-400 -200 0 200 400 (ms)

eFigure 9

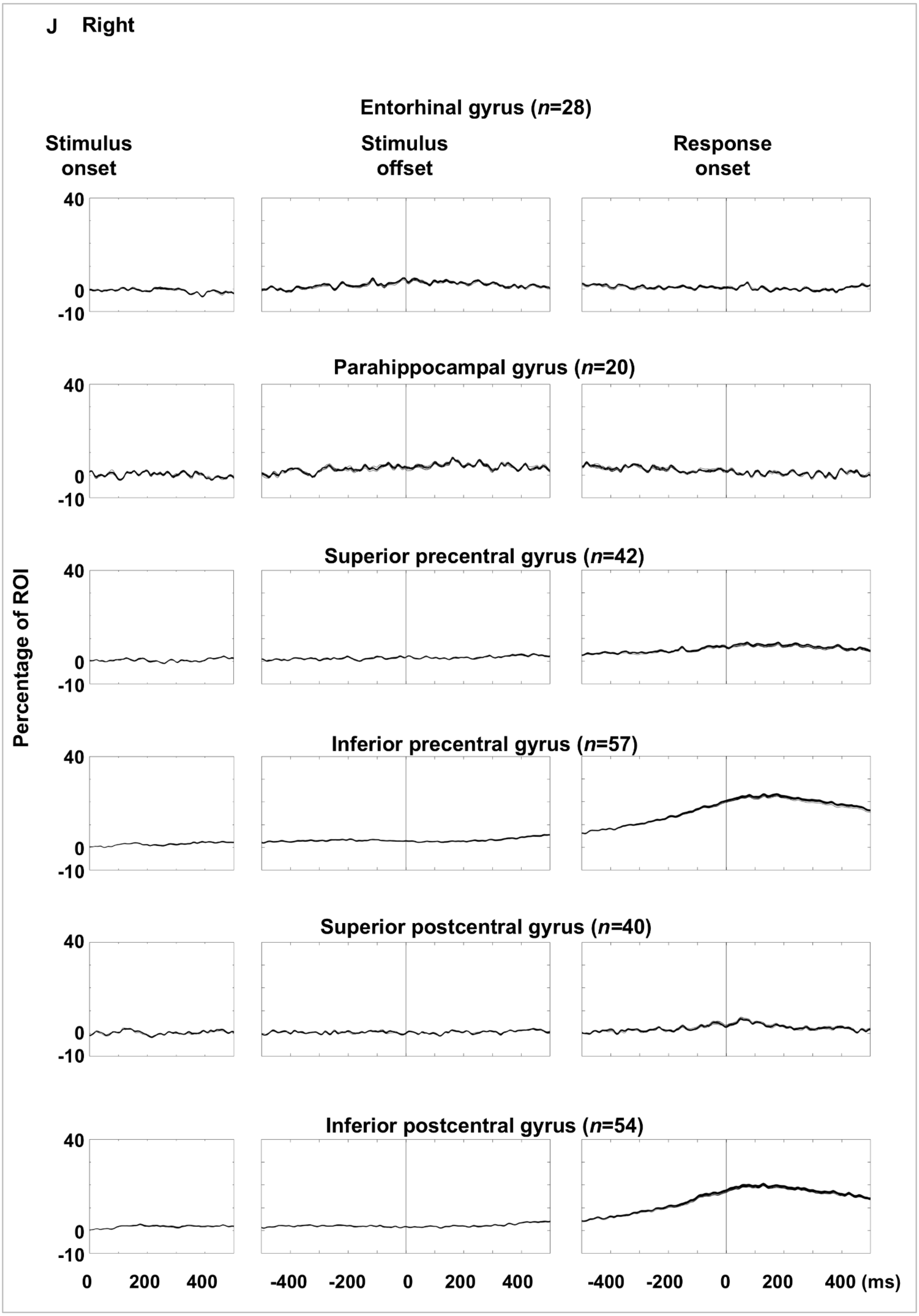
J Right
Entorhinal gyrus (n=28)
Stimulus onset
Stimulus offset
Response onset
Parahippocampal gyrus (n=20)
Superior precentral gyrus (n=42)
Inferior precentral gyrus (n=57)
Superior postcentral gyrus (n=40)
Inferior postcentral gyrus (n=54)
Percentage of ROI
40
0
-10
0 200 400
-400 -200 0 200 400
-400 -200 0 200 400 (ms)

eFigure 9

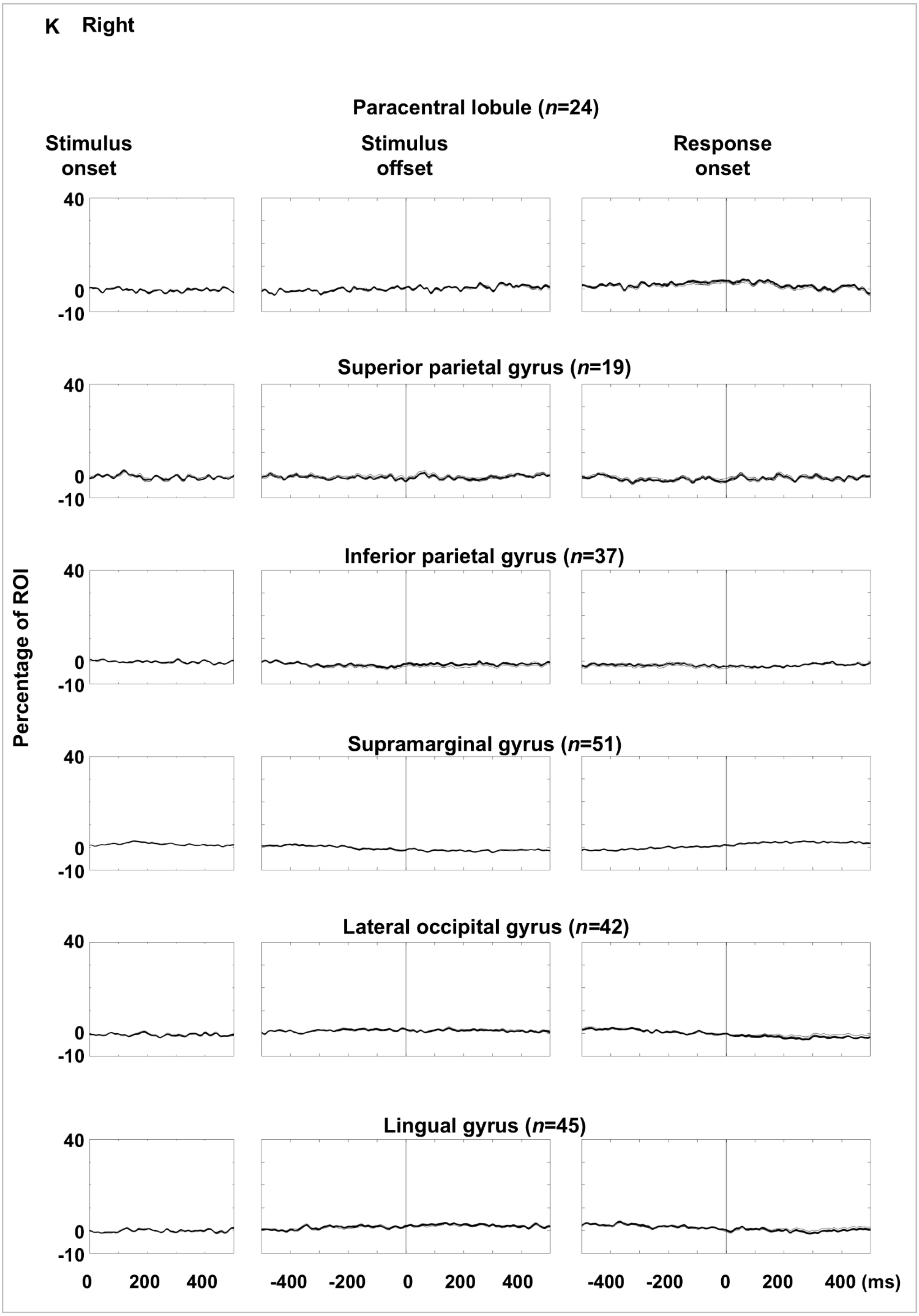
K Right
Paracentral lobule (n=24)
Stimulus onset
Stimulus offset
Response onset
Superior parietal gyrus (n=19)
Inferior parietal gyrus (n=37)
Supramarginal gyrus (n=51)
Lateral occipital gyrus (n=42)
Lingual gyrus (n=45)
Percentage of ROI
40
0
-10
0
200
400
-400
-200
400 (ms)

eFigure 9

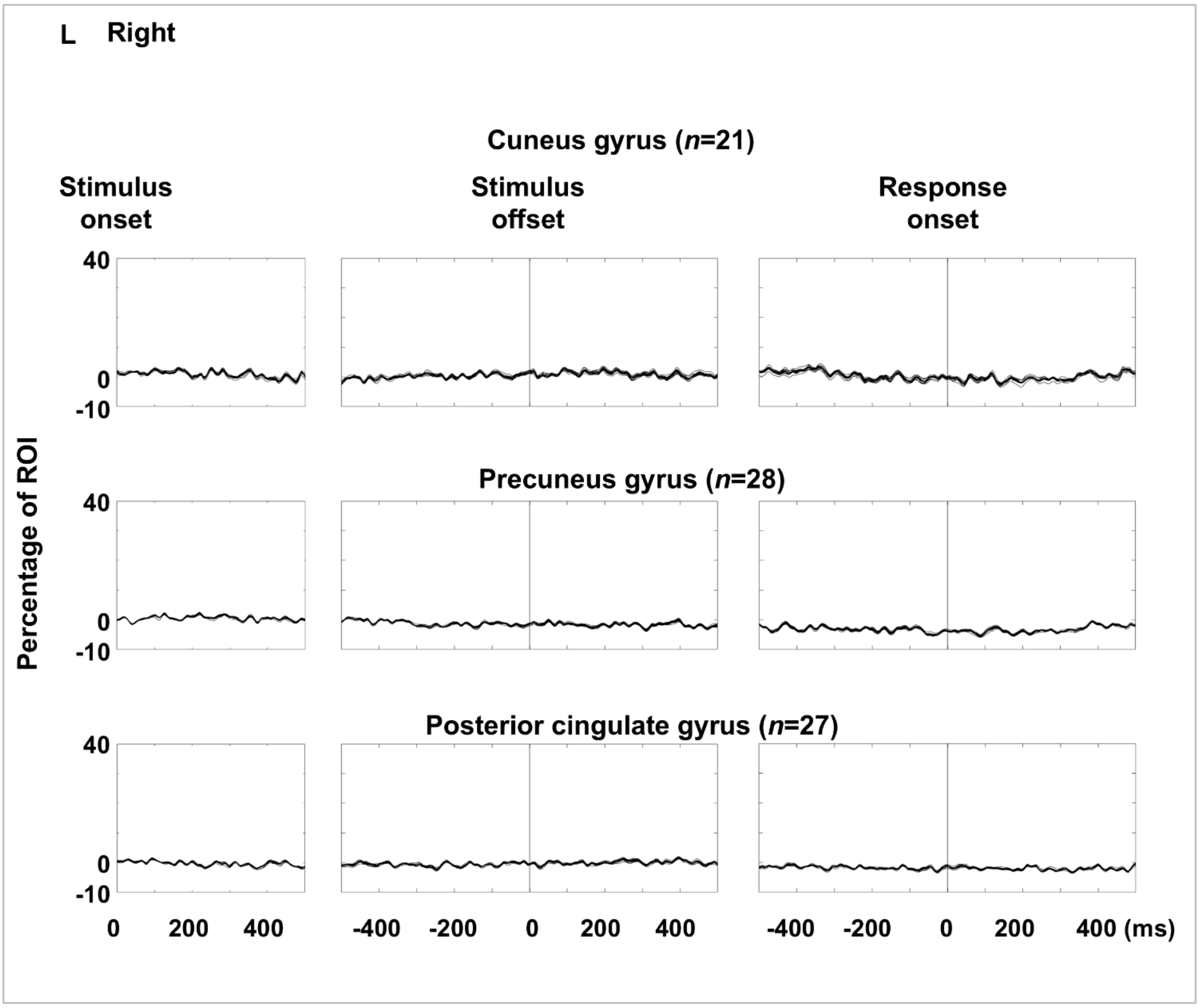


**eFigure 9. Auditory naming-related high-gamma amplitude dynamics at regions of interest (ROIs): leave-one-patient-out analysis.** Each plot shows the dynamics of high-gamma amplitude at each ROI (% change relative to baseline) averaged across all patients except one excluded patient. **A–F** Left hemispheric ROIs. **G-L** Right hemispheric ROIs. To demonstrate that high-gamma dynamics are robust and not driven disproportionately by individual patients, we computed Spearman's correlation coefficient (rho value) repeatedly, each time excluding a different patient's data (leave-one-patient-out approach). The mean Spearman's rho value at each ROI ranged from 0.946 to 0.999. One-sample *t*-tests confirmed mean rho values significantly greater than zero at all 66 ROIs (p-value range: $5.12 \times 10^{-254}$ to $3.02 \times 10^{-13}$).

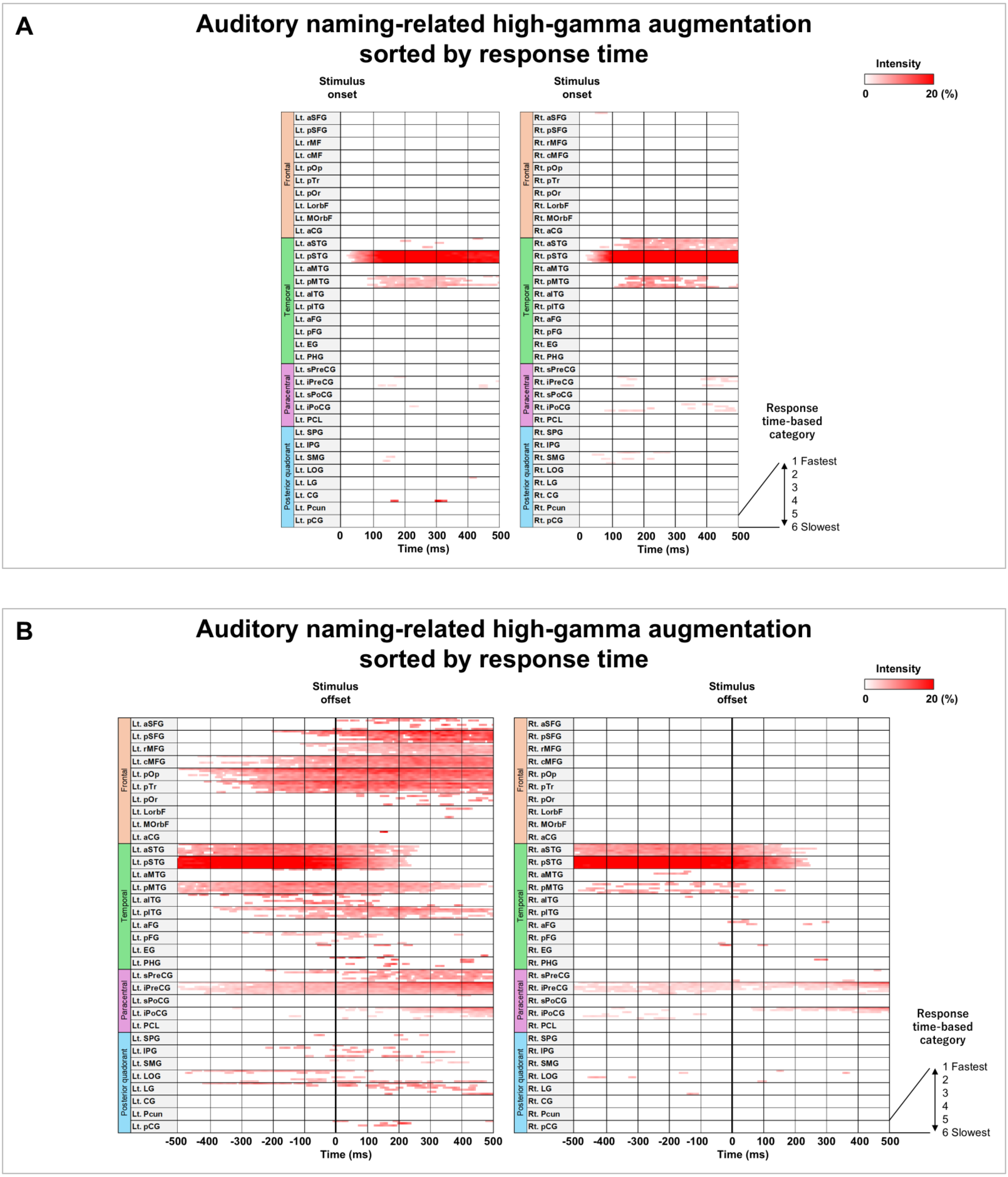
A
Auditory naming-related high-gamma augmentation sorted by response time
Stimulus onset
Stimulus onset
Intensity
0
20 (%)
Frontal
Temporal
Paracentral
Posterior quadorant
Lt. aSFG
Lt. pSFG
Lt. rMF
Lt. cMF
Lt. pOp
Lt. pTr
Lt. pOr
Lt. LorbF
Lt. MOrbF
Lt. aCG
Lt. aSTG
Lt. pSTG
Lt. aMTG
Lt. pMTG
Lt. aITG
Lt. pITG
Lt. aFG
Lt. pFG
Lt. EG
Lt. PHG
Lt. sPreCG
Lt. iPreCG
Lt. sPoCG
Lt. iPoCG
Lt. PCL
Lt. SPG
Lt. IPG
Lt. SMG
Lt. LOG
Lt. LG
Lt. CG
Lt. Pcun
Lt. pCG
Rt. aSFG
Rt. pSFG
Rt. rMFG
Rt. cMFG
Rt. pOp
Rt. pTr
Rt. pOr
Rt. LorbF
Rt. MOrbF
Rt. aCG
Rt. aSTG
Rt. pSTG
Rt. aMTG
Rt. pMTG
Rt. aITG
Rt. pITG
Rt. aFG
Rt. pFG
Rt. EG
Rt. PHG
Rt. sPreCG
Rt. iPreCG
Rt. sPoCG
Rt. iPoCG
Rt. PCL
Rt. SPG
Rt. IPG
Rt. SMG
Rt. LOG
Rt. LG
Rt. CG
Rt. Pcun
Rt. pCG
0 100 200 300 400 500
Time (ms)
Response time-based category
1 Fastest
2
3
4
5
6 Slowest
B
Auditory naming-related high-gamma augmentation sorted by response time
Stimulus offset
Stimulus offset
Intensity
0
20 (%)
Lt. aSFG
Lt. pSFG
Lt. rMFG
Lt. cMFG
Lt. pOp
Lt. pTr
Lt. pOr
Lt. LorbF
Lt. MOrbF
Lt. aCG
Rt. aSFG
Rt. pSFG
Rt. rMFG
Rt. cMFG
Rt. pOp
Rt. pTr
Rt. pOr
Rt. LorbF
Rt. MOrbF
Rt. aCG
-500 -400 -300 -200 -100 0 100 200 300 400 500
Time (ms)
Response time-based category
1 Fastest
2
3
4
5
6 Slowest

**eFigure 10**

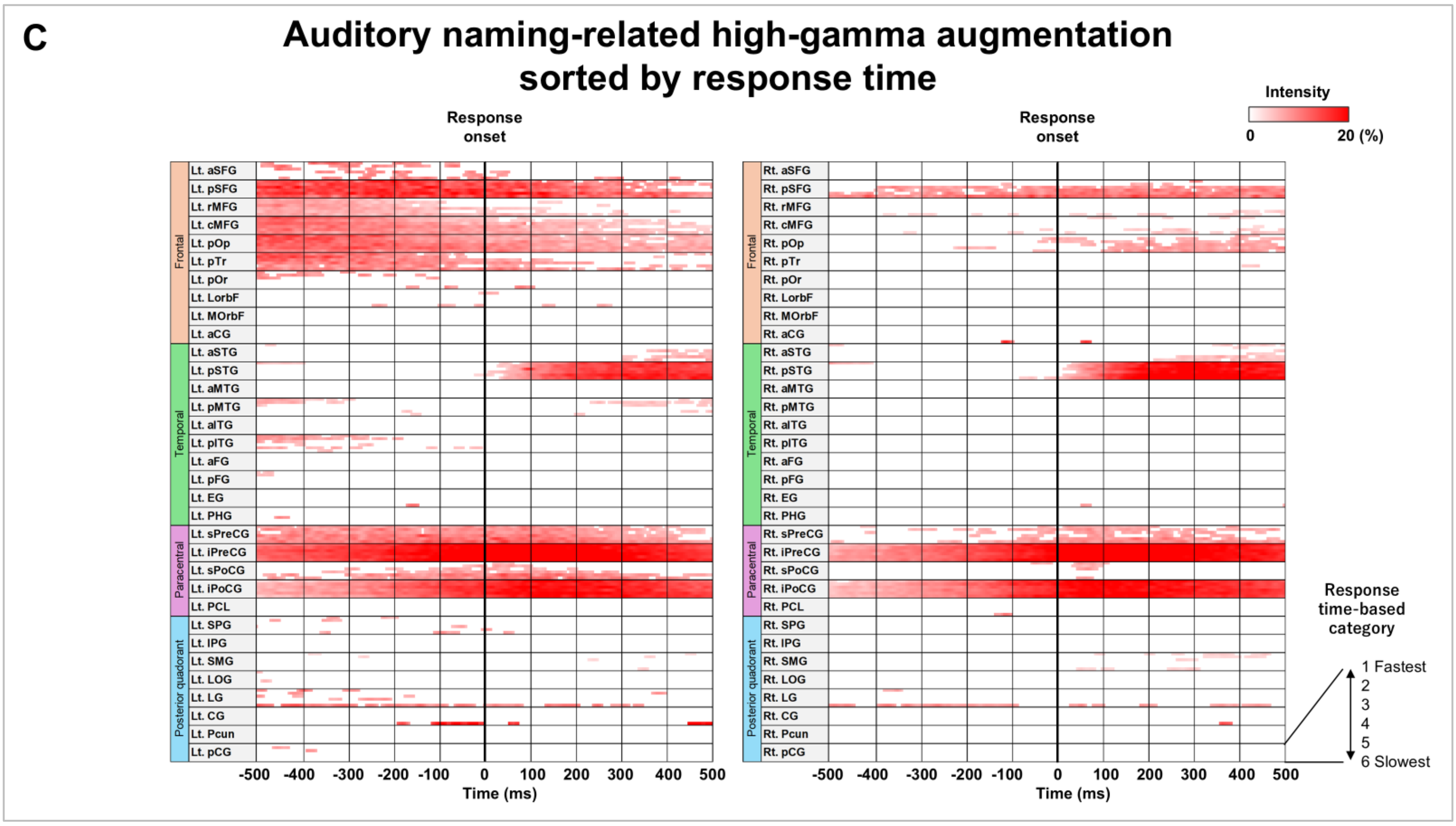
C
Auditory naming-related high-gamma augmentation
sorted by response time
Response onset
Response onset
Intensity
0
20 (%)
Lt. aSFG
Lt. pSFG
Lt. rMFG
Lt. cMFG
Lt. pOp
Lt. pTr
Lt. pOr
Lt. LorbF
Lt. MOrbF
Lt. aCG
Lt. aSTG
Lt. pSTG
Lt. aMTG
Lt. pMTG
Lt. aITG
Lt. pITG
Lt. aFG
Lt. pFG
Lt. EG
Lt. PHG
Lt. sPreCG
Lt. iPreCG
Lt. sPoCG
Lt. iPoCG
Lt. PCL
Lt. SPG
Lt. IPG
Lt. SMG
Lt. LOG
Lt. LG
Lt. CG
Lt. Pcun
Lt. pCG
Rt. aSFG
Rt. pSFG
Rt. rMFG
Rt. cMFG
Rt. pOp
Rt. pTr
Rt. pOr
Rt. LorbF
Rt. MOrbF
Rt. aCG
Rt. aSTG
Rt. pSTG
Rt. aMTG
Rt. pMTG
Rt. aITG
Rt. pITG
Rt. aFG
Rt. pFG
Rt. EG
Rt. PHG
Rt. sPreCG
Rt. iPreCG
Rt. sPoCG
Rt. iPoCG
Rt. PCL
Rt. SPG
Rt. IPG
Rt. SMG
Rt. LOG
Rt. LG
Rt. CG
Rt. Pcun
Rt. pCG
Frontal
Temporal
Paracentral
Posterior quadorant
-500 -400 -300 -200 -100 0 100 200 300 400 500
Time (ms)
Response time-based category
1 Fastest
2
3
4
5
6 Slowest

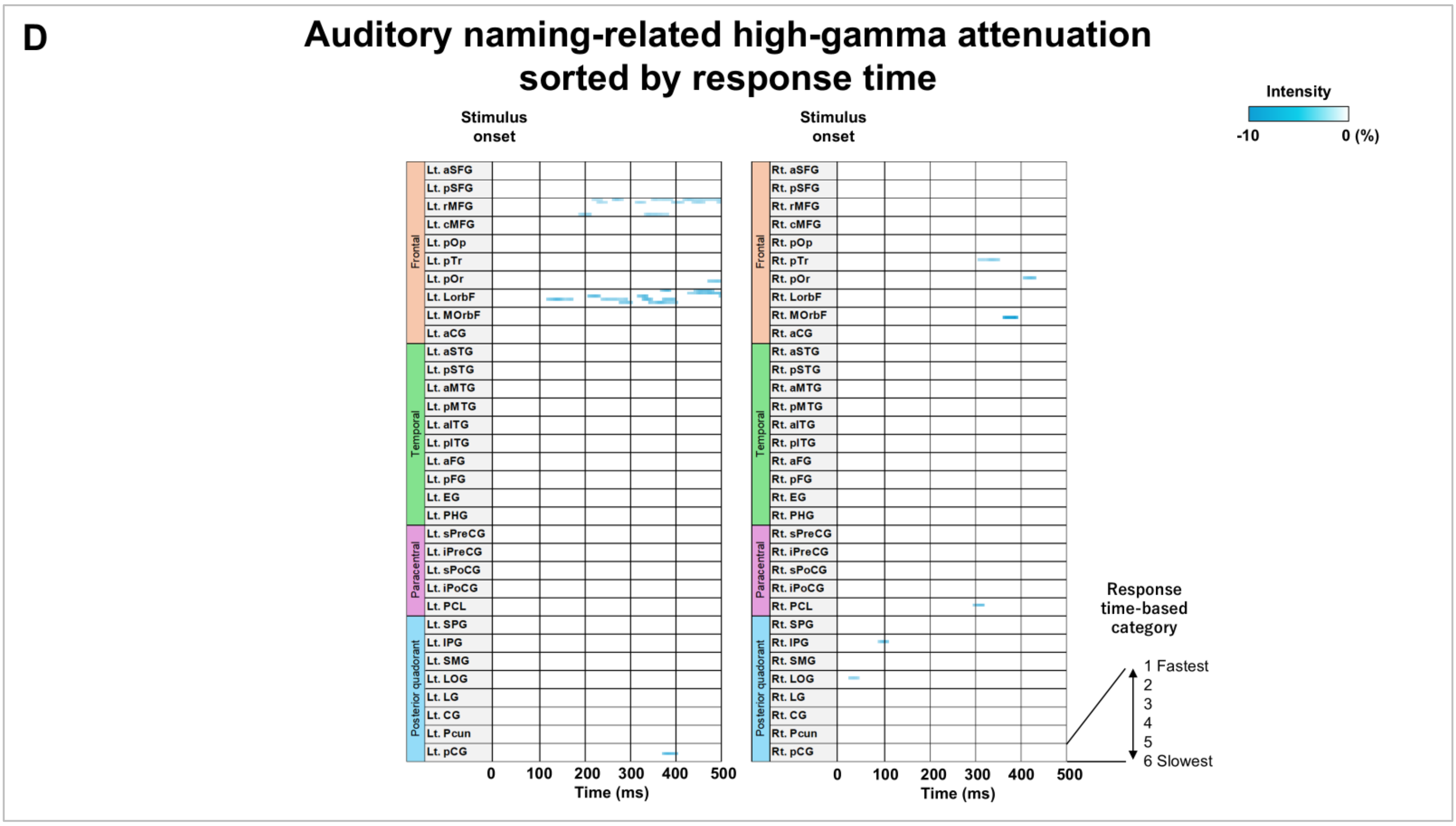
D
Auditory naming-related high-gamma attenuation
sorted by response time
Stimulus onset
Stimulus onset
Intensity
-10
0 (%)
Frontal
Temporal
Paracentral
Posterior quadorant
0 100 200 300 400 500
Time (ms)
Response time-based category
1 Fastest
2
3
4
5
6 Slowest

**eFigure 10**

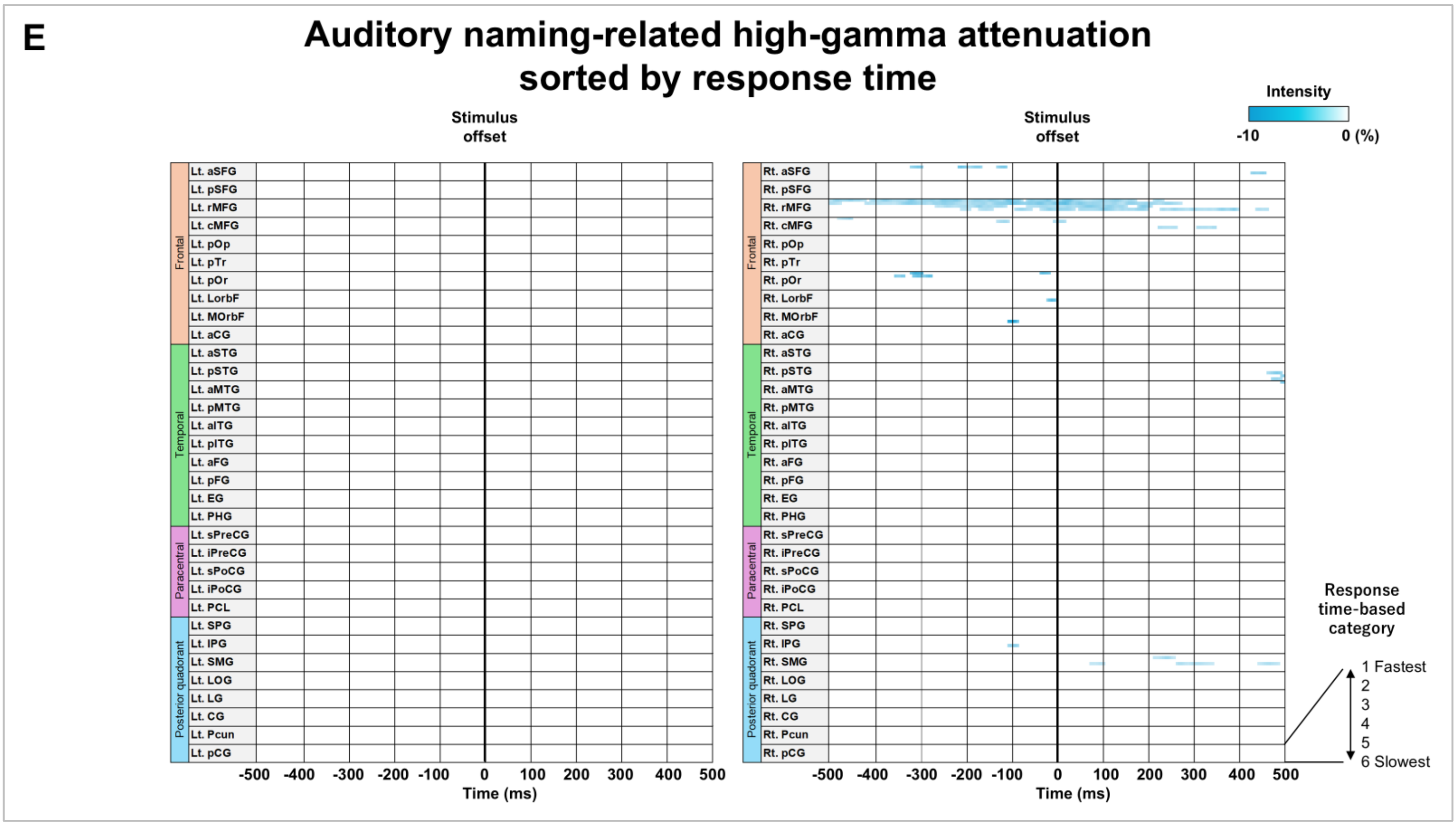
E
Auditory naming-related high-gamma attenuation sorted by response time
Stimulus offset
Stimulus offset
Intensity
-10
0 (%)
Frontal
Temporal
Paracentral
Posterior quadorant
Lt. aSFG
Lt. pSFG
Lt. rMFG
Lt. cMFG
Lt. pOp
Lt. pTr
Lt. pOr
Lt. LorbF
Lt. MOrbF
Lt. aCG
Lt. aSTG
Lt. pSTG
Lt. aMTG
Lt. pMTG
Lt. aITG
Lt. pITG
Lt. aFG
Lt. pFG
Lt. EG
Lt. PHG
Lt. sPreCG
Lt. iPreCG
Lt. sPoCG
Lt. iPoCG
Lt. PCL
Lt. SPG
Lt. IPG
Lt. SMG
Lt. LOG
Lt. LG
Lt. CG
Lt. Pcun
Lt. pCG
Rt. aSFG
Rt. pSFG
Rt. rMFG
Rt. cMFG
Rt. pOp
Rt. pTr
Rt. pOr
Rt. LorbF
Rt. MOrbF
Rt. aCG
Rt. aSTG
Rt. pSTG
Rt. aMTG
Rt. pMTG
Rt. aITG
Rt. pITG
Rt. aFG
Rt. pFG
Rt. EG
Rt. PHG
Rt. sPreCG
Rt. iPreCG
Rt. sPoCG
Rt. iPoCG
Rt. PCL
Rt. SPG
Rt. IPG
Rt. SMG
Rt. LOG
Rt. LG
Rt. CG
Rt. Pcun
Rt. pCG
-500 -400 -300 -200 -100 0 100 200 300 400 500
Time (ms)
Response time-based category
1 Fastest
2
3
4
5
6 Slowest

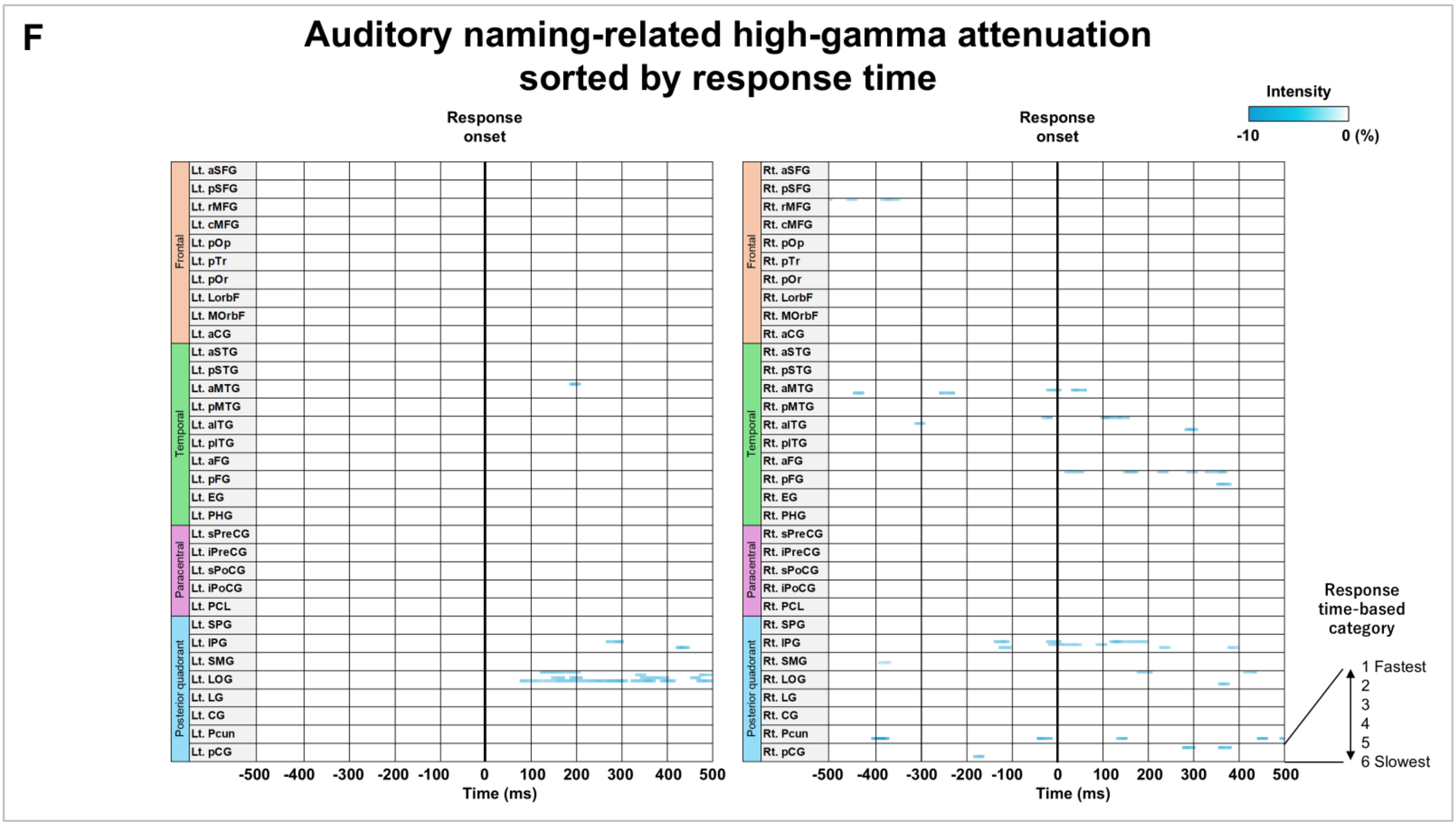
F
Auditory naming-related high-gamma attenuation sorted by response time
Response onset
Response onset
Intensity
-10
0 (%)
Frontal
Temporal
Paracentral
Posterior quadorant
-500 -400 -300 -200 -100 0 100 200 300 400 500
Time (ms)
Response time-based category
1 Fastest
2
3
4
5
6 Slowest

eFigure 10

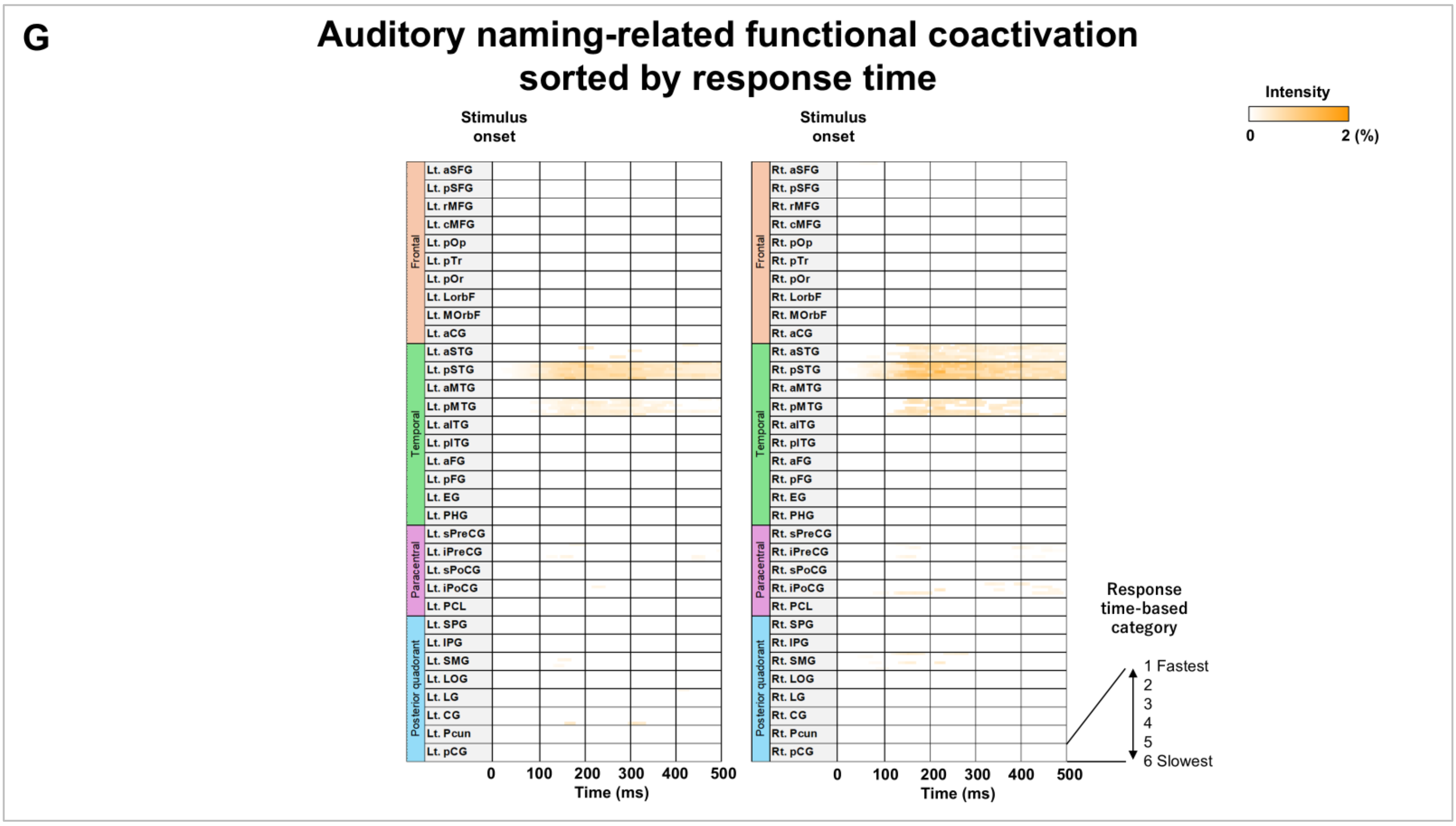
G
Auditory naming-related functional coactivation sorted by response time
Stimulus onset
Stimulus onset
Intensity
0
2 (%)
Frontal
Temporal
Paracentral
Posterior quadorant
Lt. aSFG
Lt. pSFG
Lt. rMFG
Lt. cMFG
Lt. pOp
Lt. pTr
Lt. pOr
Lt. LorbF
Lt. MOrbF
Lt. aCG
Lt. aSTG
Lt. pSTG
Lt. aMTG
Lt. pMTG
Lt. aITG
Lt. pITG
Lt. aFG
Lt. pFG
Lt. EG
Lt. PHG
Lt. sPreCG
Lt. iPreCG
Lt. sPoCG
Lt. iPoCG
Lt. PCL
Lt. SPG
Lt. IPG
Lt. SMG
Lt. LOG
Lt. LG
Lt. CG
Lt. Pcun
Lt. pCG
Rt. aSFG
Rt. pSFG
Rt. rMFG
Rt. cMFG
Rt. pOp
Rt. pTr
Rt. pOr
Rt. LorbF
Rt. MOrbF
Rt. aCG
Rt. aSTG
Rt. pSTG
Rt. aMTG
Rt. pMTG
Rt. aITG
Rt. pITG
Rt. aFG
Rt. pFG
Rt. EG
Rt. PHG
Rt. sPreCG
Rt. iPreCG
Rt. sPoCG
Rt. iPoCG
Rt. PCL
Rt. SPG
Rt. IPG
Rt. SMG
Rt. LOG
Rt. LG
Rt. CG
Rt. Pcun
Rt. pCG
0 100 200 300 400 500
Time (ms)
Response time-based category
1 Fastest
2
3
4
5
6 Slowest

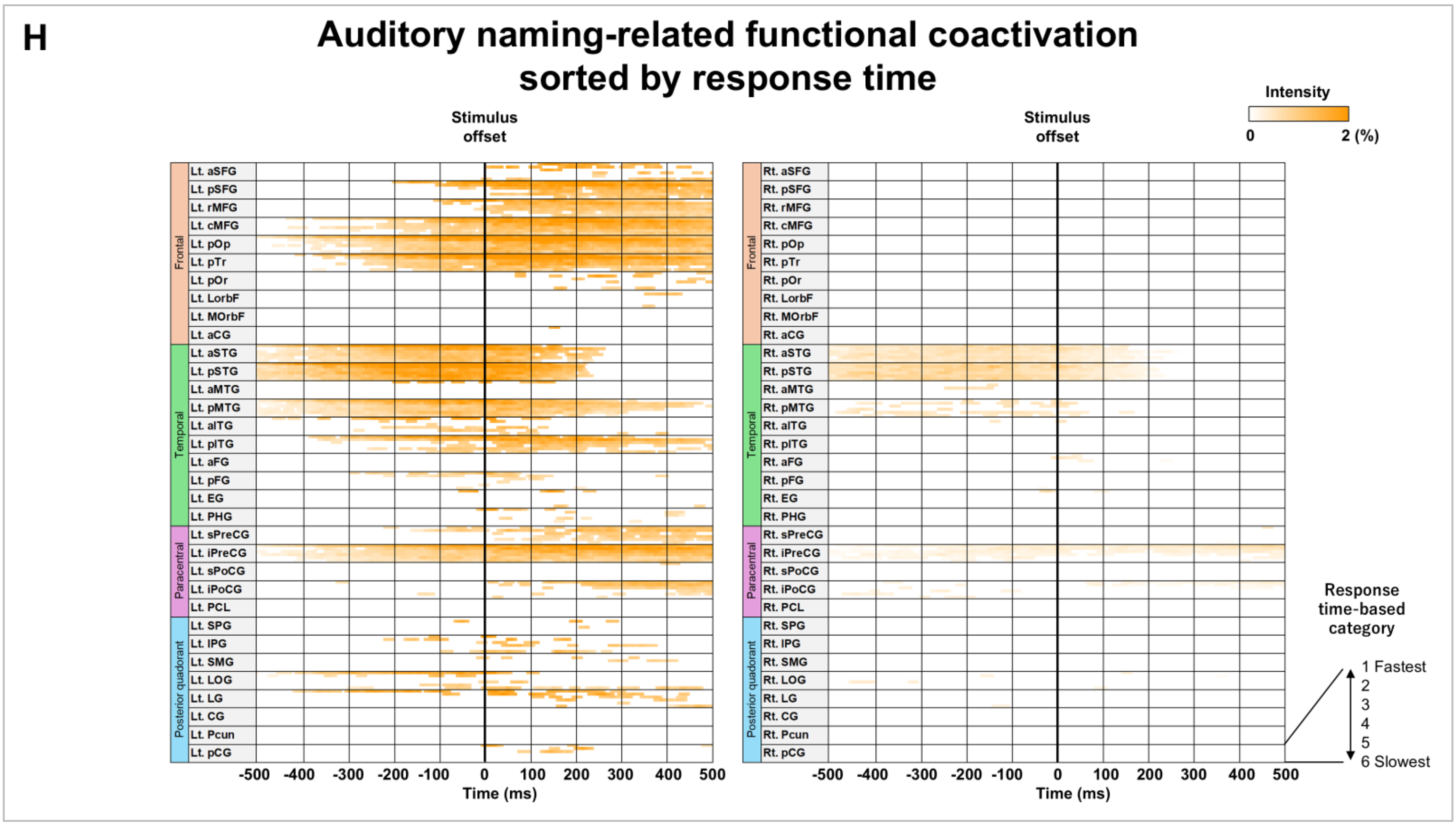
H
Auditory naming-related functional coactivation sorted by response time
Stimulus offset
Stimulus offset
Intensity
0
2 (%)
Frontal
Temporal
Paracentral
Posterior quadorant
-500 -400 -300 -200 -100 0 100 200 300 400 500
Time (ms)
Response time-based category
1 Fastest
2
3
4
5
6 Slowest

**eFigure 10**

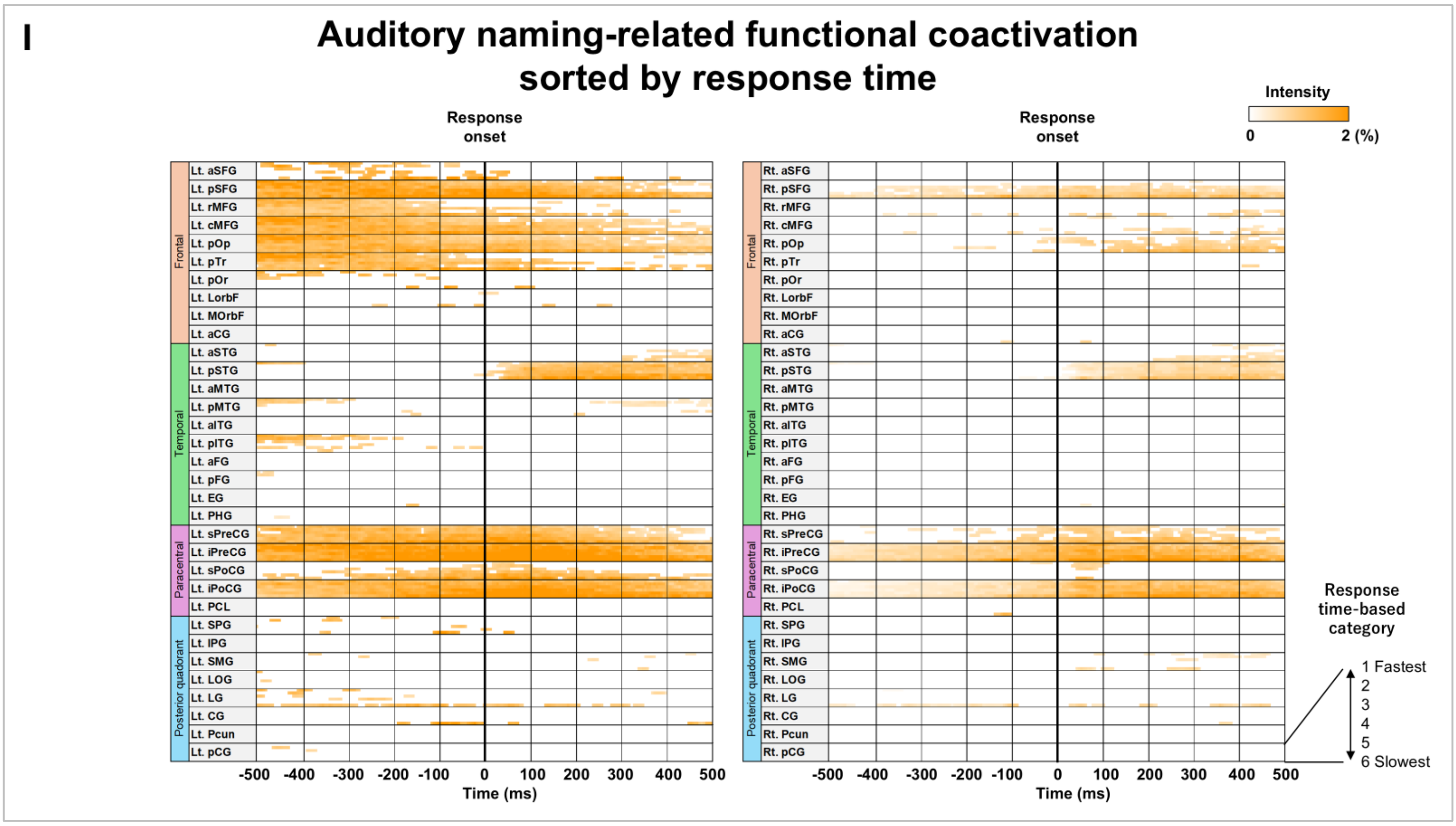
I
Auditory naming-related functional coactivation sorted by response time
Response onset
Response onset
Intensity
0
2 (%)
Lt. aSFG
Lt. pSFG
Lt. rMFG
Lt. cMFG
Lt. pOp
Lt. pTr
Lt. pOr
Lt. LorbF
Lt. MOrbF
Lt. aCG
Lt. aSTG
Lt. pSTG
Lt. aMTG
Lt. pMTG
Lt. aITG
Lt. pITG
Lt. aFG
Lt. pFG
Lt. EG
Lt. PHG
Lt. sPreCG
Lt. iPreCG
Lt. sPoCG
Lt. iPoCG
Lt. PCL
Lt. SPG
Lt. IPG
Lt. SMG
Lt. LOG
Lt. LG
Lt. CG
Lt. Pcun
Lt. pCG
Frontal
Temporal
Paracentral
Posterior quadorant
Rt. aSFG
Rt. pSFG
Rt. rMFG
Rt. cMFG
Rt. pOp
Rt. pTr
Rt. pOr
Rt. LorbF
Rt. MOrbF
Rt. aCG
Rt. aSTG
Rt. pSTG
Rt. aMTG
Rt. pMTG
Rt. aITG
Rt. pITG
Rt. aFG
Rt. pFG
Rt. EG
Rt. PHG
Rt. sPreCG
Rt. iPreCG
Rt. sPoCG
Rt. iPoCG
Rt. PCL
Rt. SPG
Rt. IPG
Rt. SMG
Rt. LOG
Rt. LG
Rt. CG
Rt. Pcun
Rt. pCG
-500 -400 -300 -200 -100 0 100 200 300 400 500
Time (ms)
Response time-based category
1 Fastest
2
3
4
5
6 Slowest

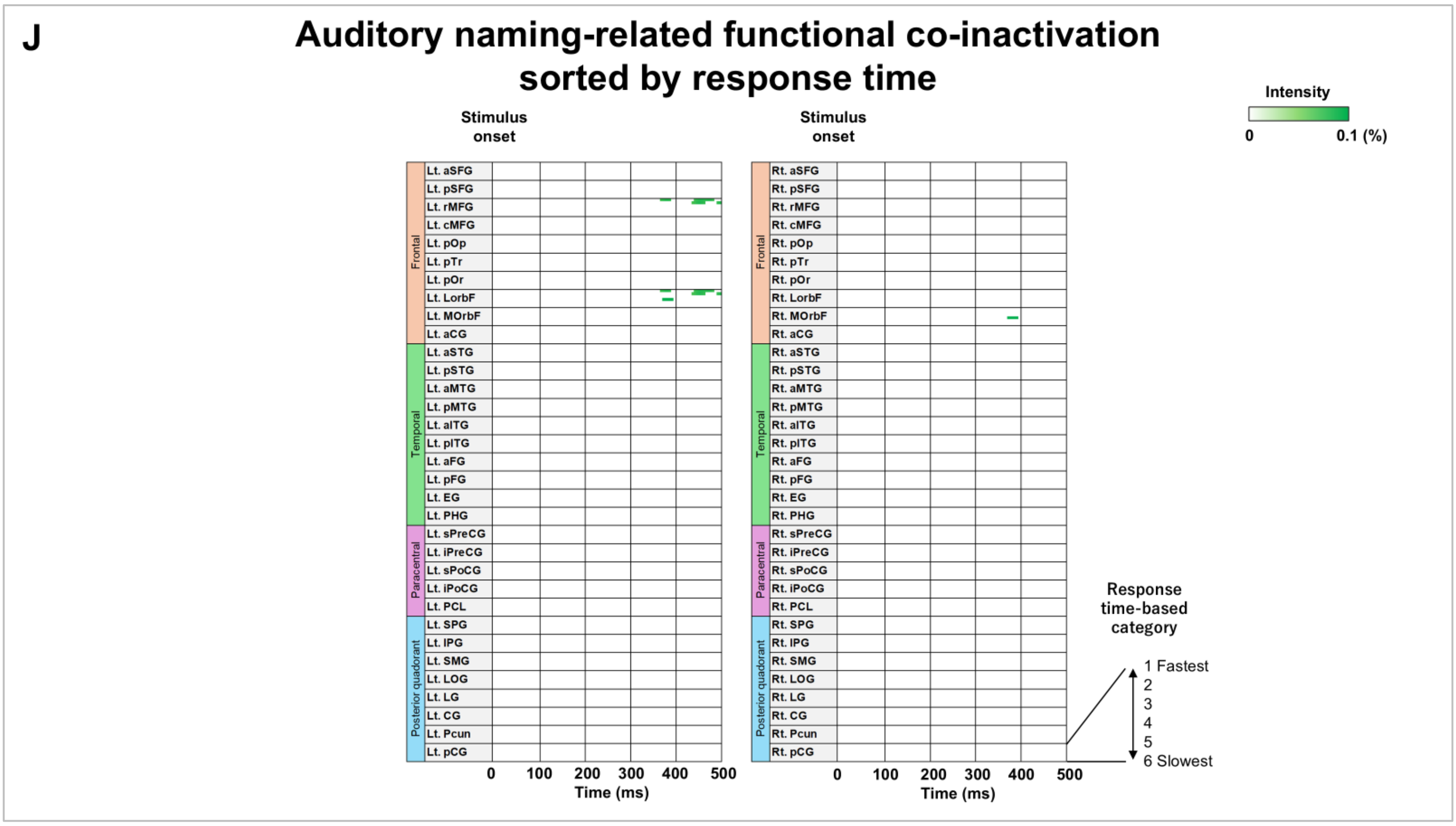
J
Auditory naming-related functional co-inactivation sorted by response time
Stimulus onset
Stimulus onset
Intensity
0
0.1 (%)
Lt. aSFG
Lt. pSFG
Lt. rMFG
Lt. cMFG
Lt. pOp
Lt. pTr
Lt. pOr
Lt. LorbF
Lt. MOrbF
Lt. aCG
Lt. aSTG
Lt. pSTG
Lt. aMTG
Lt. pMTG
Lt. aITG
Lt. pITG
Lt. aFG
Lt. pFG
Lt. EG
Lt. PHG
Lt. sPreCG
Lt. iPreCG
Lt. sPoCG
Lt. iPoCG
Lt. PCL
Lt. SPG
Lt. IPG
Lt. SMG
Lt. LOG
Lt. LG
Lt. CG
Lt. Pcun
Lt. pCG
Frontal
Temporal
Paracentral
Posterior quadorant
Rt. aSFG
Rt. pSFG
Rt. rMFG
Rt. cMFG
Rt. pOp
Rt. pTr
Rt. pOr
Rt. LorbF
Rt. MOrbF
Rt. aCG
Rt. aSTG
Rt. pSTG
Rt. aMTG
Rt. pMTG
Rt. aITG
Rt. pITG
Rt. aFG
Rt. pFG
Rt. EG
Rt. PHG
Rt. sPreCG
Rt. iPreCG
Rt. sPoCG
Rt. iPoCG
Rt. PCL
Rt. SPG
Rt. IPG
Rt. SMG
Rt. LOG
Rt. LG
Rt. CG
Rt. Pcun
Rt. pCG
0 100 200 300 400 500
Time (ms)
Response time-based category
1 Fastest
2
3
4
5
6 Slowest

**eFigure 10**

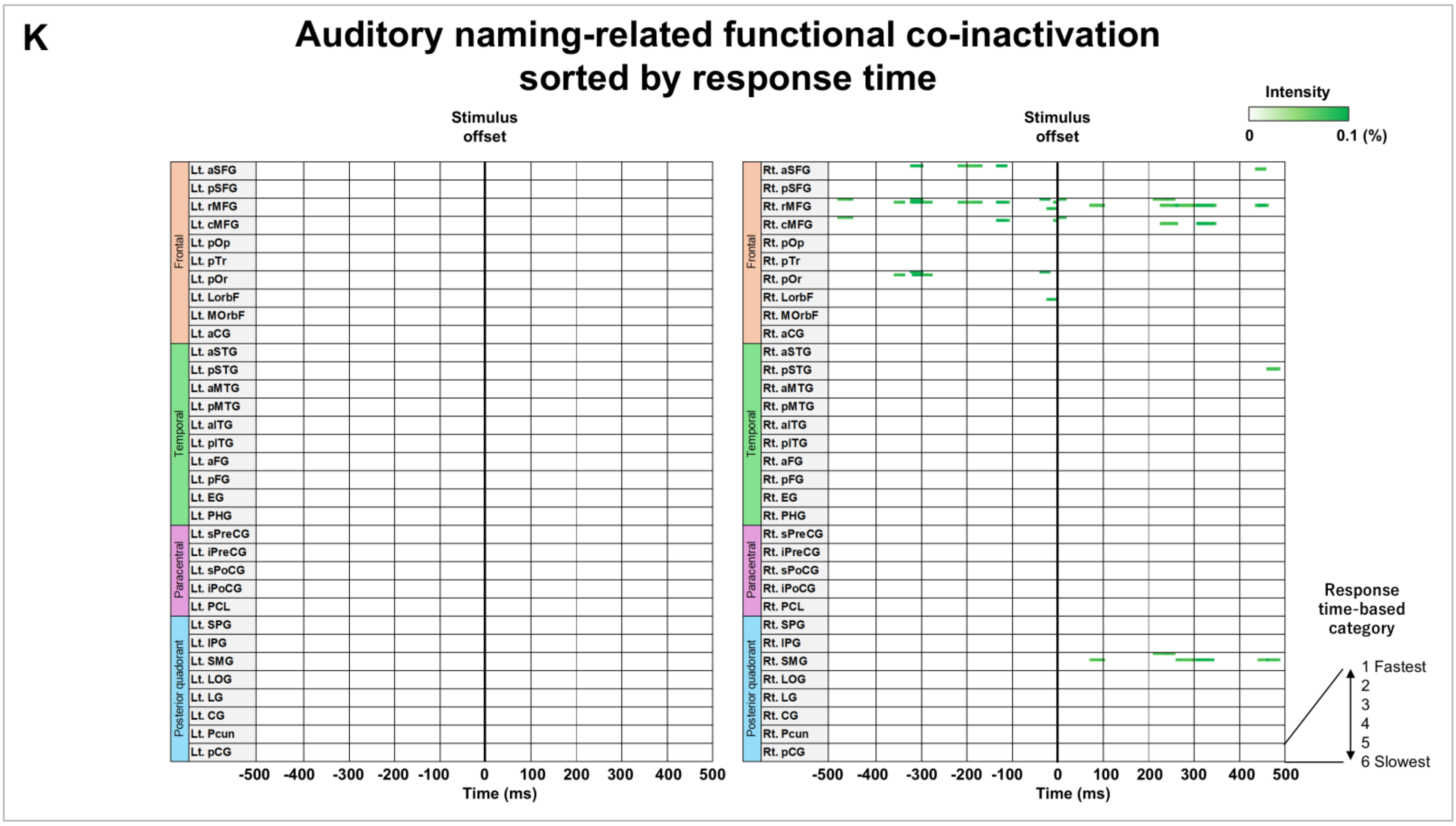
K
Auditory naming-related functional co-inactivation sorted by response time
Stimulus offset
Intensity
0
0.1 (%)
Frontal
Temporal
Paracentral
Posterior quadorant
Lt. aSFG
Lt. pSFG
Lt. rMFG
Lt. cMFG
Lt. pOp
Lt. pTr
Lt. pOr
Lt. LorbF
Lt. MOrbF
Lt. aCG
Lt. aSTG
Lt. pSTG
Lt. aMTG
Lt. pMTG
Lt. aITG
Lt. pITG
Lt. aFG
Lt. pFG
Lt. EG
Lt. PHG
Lt. sPreCG
Lt. iPreCG
Lt. sPoCG
Lt. iPoCG
Lt. PCL
Lt. SPG
Lt. IPG
Lt. SMG
Lt. LOG
Lt. LG
Lt. CG
Lt. Pcun
Lt. pCG
Rt. aSFG
Rt. pSFG
Rt. rMFG
Rt. cMFG
Rt. pOp
Rt. pTr
Rt. pOr
Rt. LorbF
Rt. MOrbF
Rt. aCG
Rt. aSTG
Rt. pSTG
Rt. aMTG
Rt. pMTG
Rt. aITG
Rt. pITG
Rt. aFG
Rt. pFG
Rt. EG
Rt. PHG
Rt. sPreCG
Rt. iPreCG
Rt. sPoCG
Rt. iPoCG
Rt. PCL
Rt. SPG
Rt. IPG
Rt. SMG
Rt. LOG
Rt. LG
Rt. CG
Rt. Pcun
Rt. pCG
-500 -400 -300 -200 -100 0 100 200 300 400 500
Time (ms)
Response time-based category
1 Fastest
2
3
4
5
6 Slowest

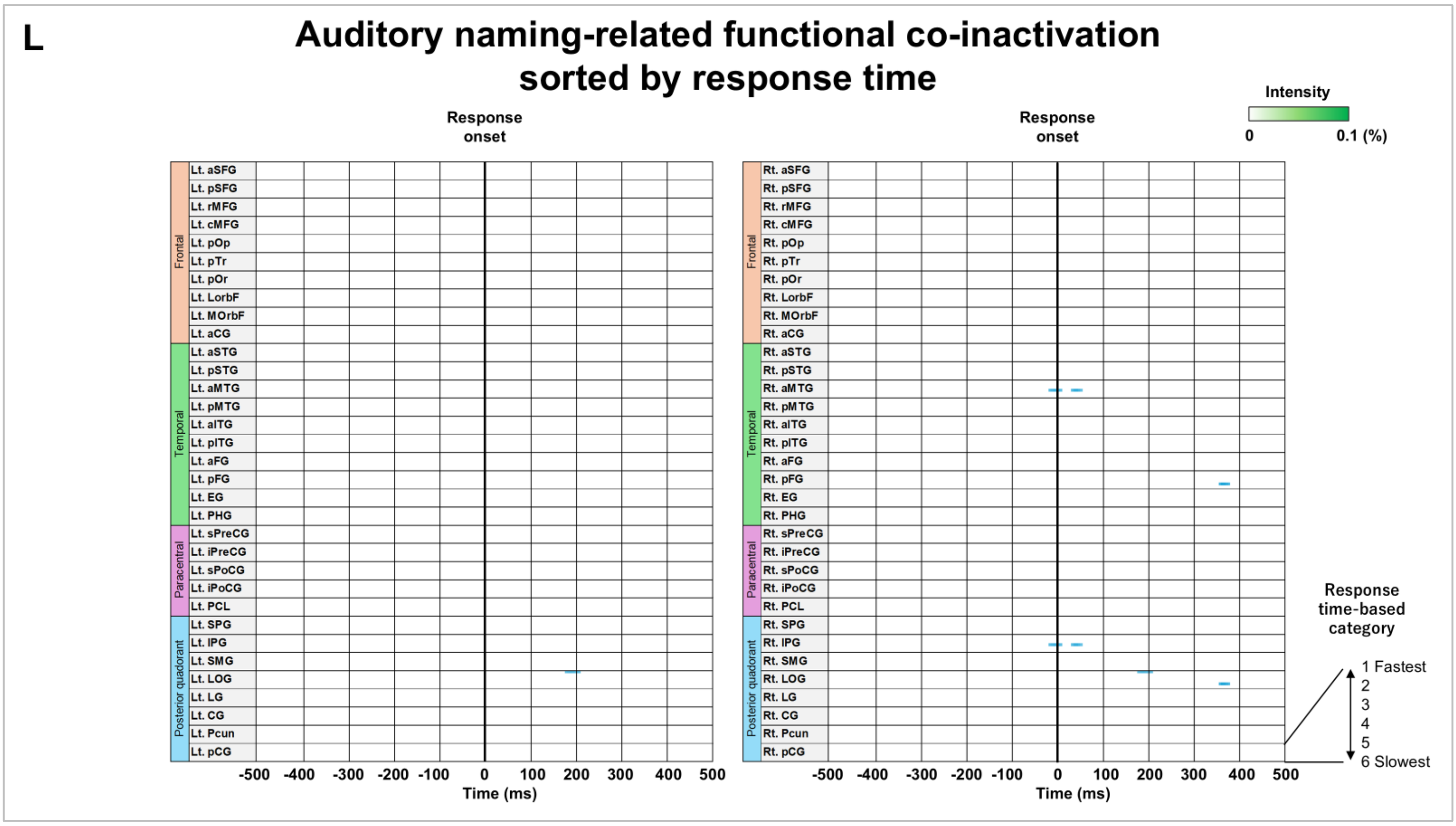
L
Auditory naming-related functional co-inactivation sorted by response time
Response onset
Intensity
0
0.1 (%)
Frontal
Temporal
Paracentral
Posterior quadorant
Lt. aSFG
Lt. pSFG
Lt. rMFG
Lt. cMFG
Lt. pOp
Lt. pTr
Lt. pOr
Lt. LorbF
Lt. MOrbF
Lt. aCG
Lt. aSTG
Lt. pSTG
Lt. aMTG
Lt. pMTG
Lt. aITG
Lt. pITG
Lt. aFG
Lt. pFG
Lt. EG
Lt. PHG
Lt. sPreCG
Lt. iPreCG
Lt. sPoCG
Lt. iPoCG
Lt. PCL
Lt. SPG
Lt. IPG
Lt. SMG
Lt. LOG
Lt. LG
Lt. CG
Lt. Pcun
Lt. pCG
Rt. aSFG
Rt. pSFG
Rt. rMFG
Rt. cMFG
Rt. pOp
Rt. pTr
Rt. pOr
Rt. LorbF
Rt. MOrbF
Rt. aCG
Rt. aSTG
Rt. pSTG
Rt. aMTG
Rt. pMTG
Rt. aITG
Rt. pITG
Rt. aFG
Rt. pFG
Rt. EG
Rt. PHG
Rt. sPreCG
Rt. iPreCG
Rt. sPoCG
Rt. iPoCG
Rt. PCL
Rt. SPG
Rt. IPG
Rt. SMG
Rt. LOG
Rt. LG
Rt. CG
Rt. Pcun
Rt. pCG
-500 -400 -300 -200 -100 0 100 200 300 400 500
Time (ms)
Response time-based category
1 Fastest
2
3
4
5
6 Slowest

eFigure 10

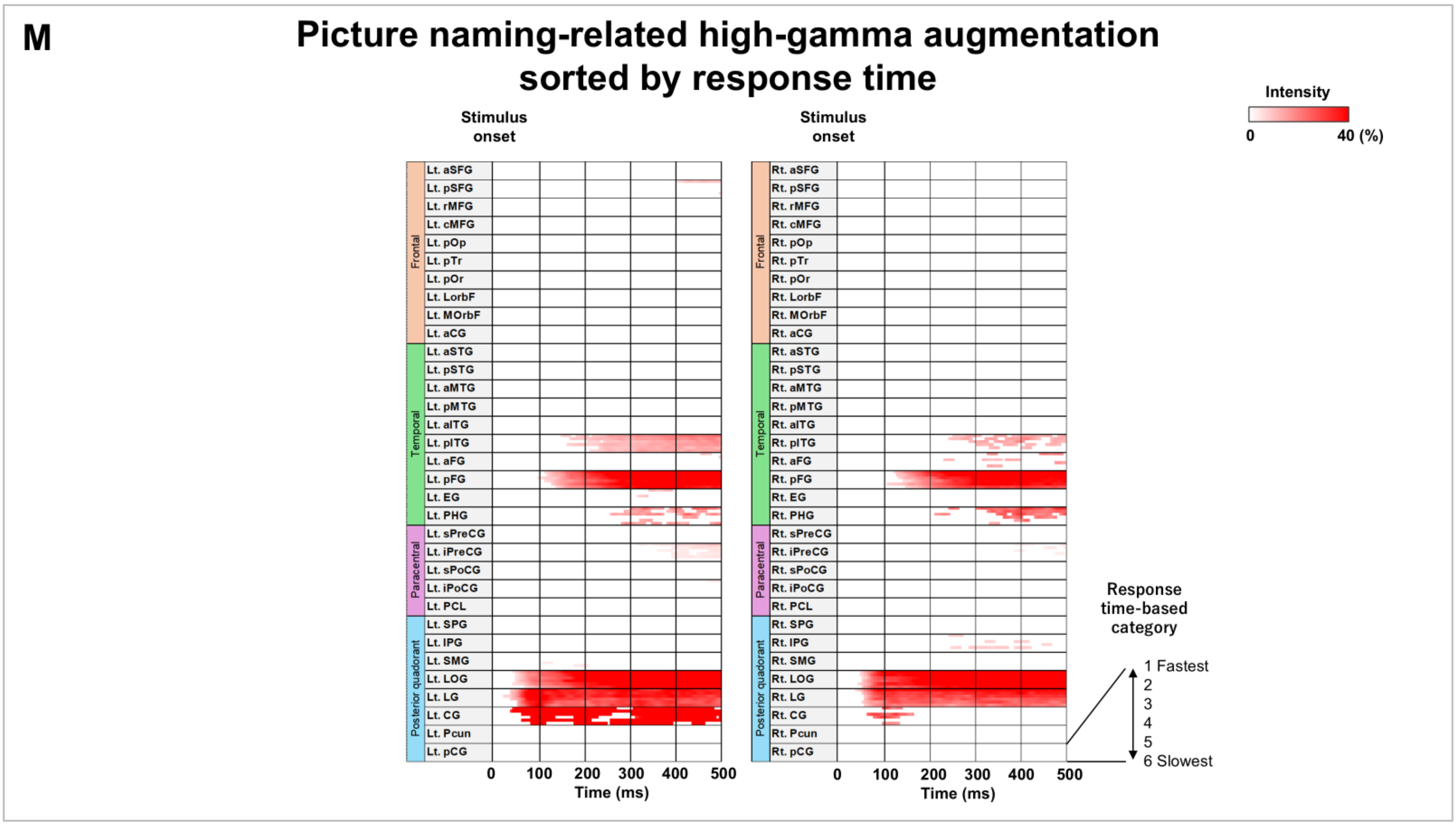
M
Picture naming-related high-gamma augmentation sorted by response time
Stimulus onset
Stimulus onset
Intensity
0
40 (%)
Frontal
Temporal
Paracentral
Posterior quadorant
Lt. aSFG
Lt. pSFG
Lt. rMFG
Lt. cMFG
Lt. pOp
Lt. pTr
Lt. pOr
Lt. LorbF
Lt. MOrbF
Lt. aCG
Lt. aSTG
Lt. pSTG
Lt. aMTG
Lt. pMTG
Lt. aITG
Lt. pITG
Lt. aFG
Lt. pFG
Lt. EG
Lt. PHG
Lt. sPreCG
Lt. iPreCG
Lt. sPoCG
Lt. iPoCG
Lt. PCL
Lt. SPG
Lt. IPG
Lt. SMG
Lt. LOG
Lt. LG
Lt. CG
Lt. Pcun
Lt. pCG
Rt. aSFG
Rt. pSFG
Rt. rMFG
Rt. cMFG
Rt. pOp
Rt. pTr
Rt. pOr
Rt. LorbF
Rt. MOrbF
Rt. aCG
Rt. aSTG
Rt. pSTG
Rt. aMTG
Rt. pMTG
Rt. aITG
Rt. pITG
Rt. aFG
Rt. pFG
Rt. EG
Rt. PHG
Rt. sPreCG
Rt. iPreCG
Rt. sPoCG
Rt. iPoCG
Rt. PCL
Rt. SPG
Rt. IPG
Rt. SMG
Rt. LOG
Rt. LG
Rt. CG
Rt. Pcun
Rt. pCG
0 100 200 300 400 500
Time (ms)
0 100 200 300 400 500
Time (ms)
Response time-based category
1 Fastest
2
3
4
5
6 Slowest

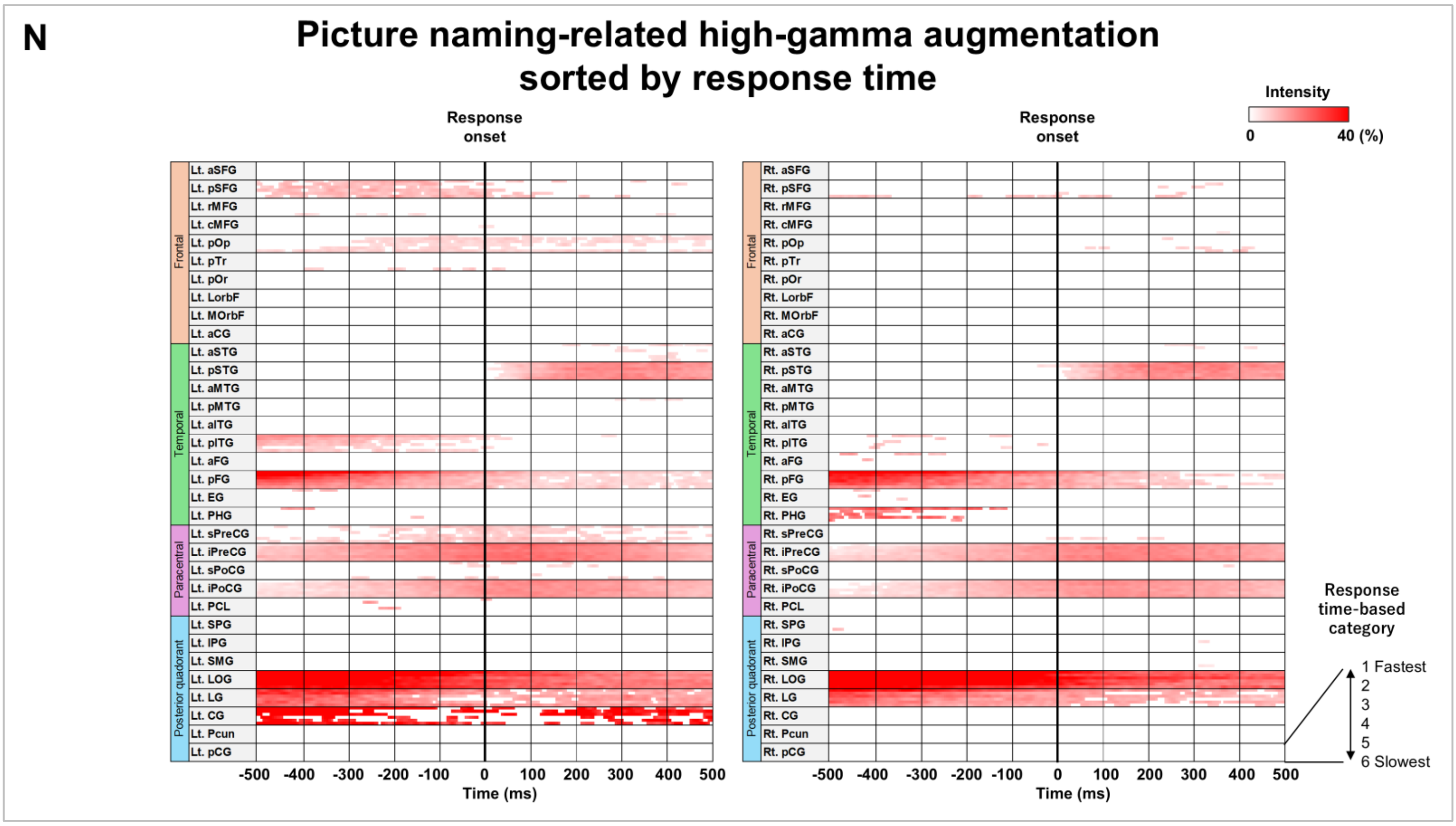
N
Picture naming-related high-gamma augmentation sorted by response time
Response onset
Response onset
Intensity
0
40 (%)
Frontal
Temporal
Paracentral
Posterior quadorant
Lt. aSFG
Lt. pSFG
Lt. rMFG
Lt. cMFG
Lt. pOp
Lt. pTr
Lt. pOr
Lt. LorbF
Lt. MOrbF
Lt. aCG
Lt. aSTG
Lt. pSTG
Lt. aMTG
Lt. pMTG
Lt. aITG
Lt. pITG
Lt. aFG
Lt. pFG
Lt. EG
Lt. PHG
Lt. sPreCG
Lt. iPreCG
Lt. sPoCG
Lt. iPoCG
Lt. PCL
Lt. SPG
Lt. IPG
Lt. SMG
Lt. LOG
Lt. LG
Lt. CG
Lt. Pcun
Lt. pCG
Rt. aSFG
Rt. pSFG
Rt. rMFG
Rt. cMFG
Rt. pOp
Rt. pTr
Rt. pOr
Rt. LorbF
Rt. MOrbF
Rt. aCG
Rt. aSTG
Rt. pSTG
Rt. aMTG
Rt. pMTG
Rt. aITG
Rt. pITG
Rt. aFG
Rt. pFG
Rt. EG
Rt. PHG
Rt. sPreCG
Rt. iPreCG
Rt. sPoCG
Rt. iPoCG
Rt. PCL
Rt. SPG
Rt. IPG
Rt. SMG
Rt. LOG
Rt. LG
Rt. CG
Rt. Pcun
Rt. pCG
-500 -400 -300 -200 -100 0 100 200 300 400 500
Time (ms)
-500 -400 -300 -200 -100 0 100 200 300 400 500
Time (ms)
Response time-based category
1 Fastest
2
3
4
5
6 Slowest

**eFigure 10**

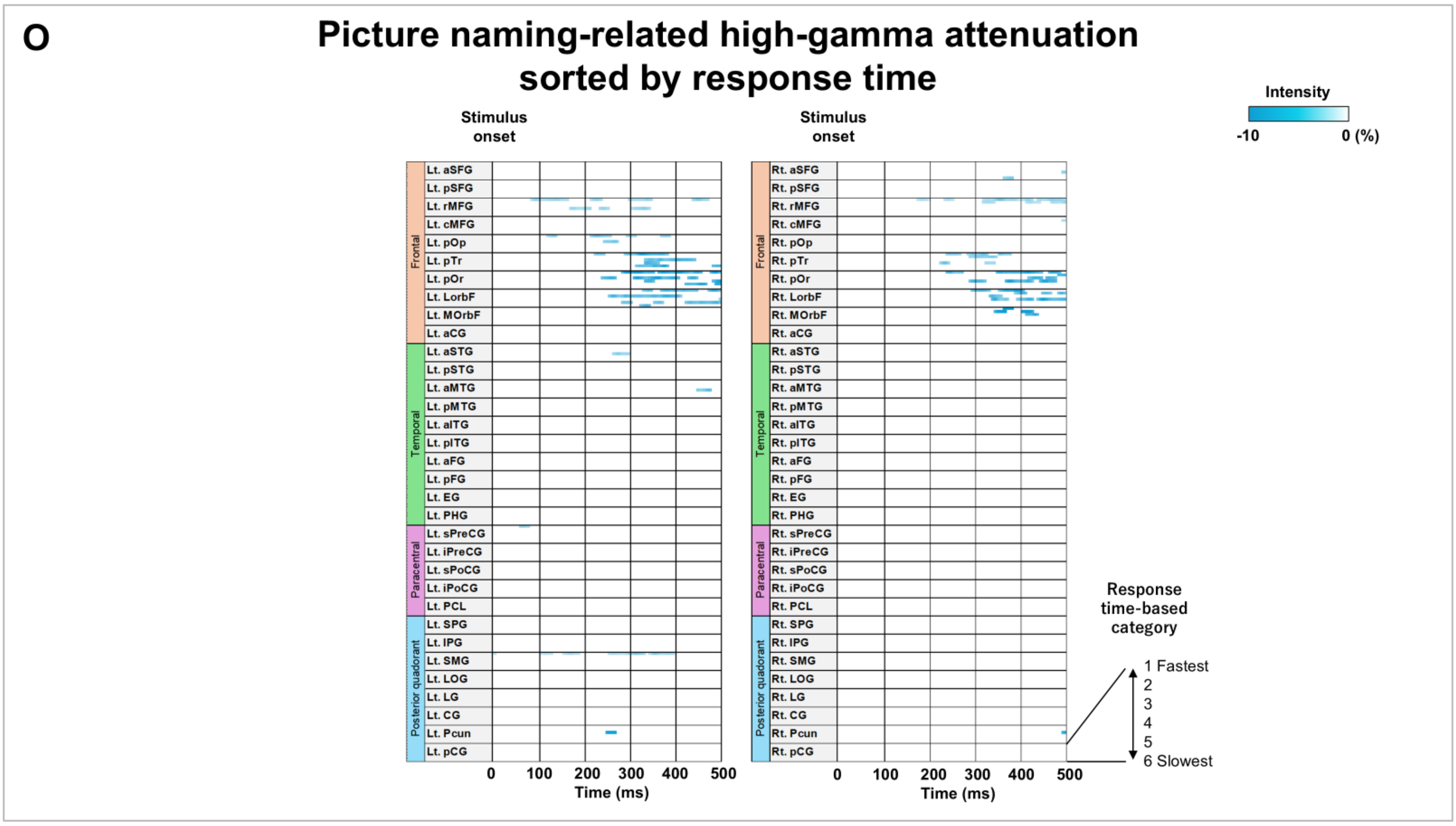

O
Picture naming-related high-gamma attenuation
sorted by response time
Stimulus onset
Stimulus onset
Intensity
-10
0 (%)
Frontal
Temporal
Paracentral
Posterior quadorant
Lt. aSFG
Lt. pSFG
Lt. rMFG
Lt. cMFG
Lt. pOp
Lt. pTr
Lt. pOr
Lt. LorbF
Lt. MOrbF
Lt. aCG
Lt. aSTG
Lt. pSTG
Lt. aMTG
Lt. pMTG
Lt. aITG
Lt. pITG
Lt. aFG
Lt. pFG
Lt. EG
Lt. PHG
Lt. sPreCG
Lt. iPreCG
Lt. sPoCG
Lt. iPoCG
Lt. PCL
Lt. SPG
Lt. IPG
Lt. SMG
Lt. LOG
Lt. LG
Lt. CG
Lt. Pcun
Lt. pCG
Rt. aSFG
Rt. pSFG
Rt. rMFG
Rt. cMFG
Rt. pOp
Rt. pTr
Rt. pOr
Rt. LorbF
Rt. MOrbF
Rt. aCG
Rt. aSTG
Rt. pSTG
Rt. aMTG
Rt. pMTG
Rt. aITG
Rt. pITG
Rt. aFG
Rt. pFG
Rt. EG
Rt. PHG
Rt. sPreCG
Rt. iPreCG
Rt. sPoCG
Rt. iPoCG
Rt. PCL
Rt. SPG
Rt. IPG
Rt. SMG
Rt. LOG
Rt. LG
Rt. CG
Rt. Pcun
Rt. pCG
0 100 200 300 400 500
Time (ms)
0 100 200 300 400 500
Time (ms)
Response time-based category
1 Fastest
2
3
4
5
6 Slowest


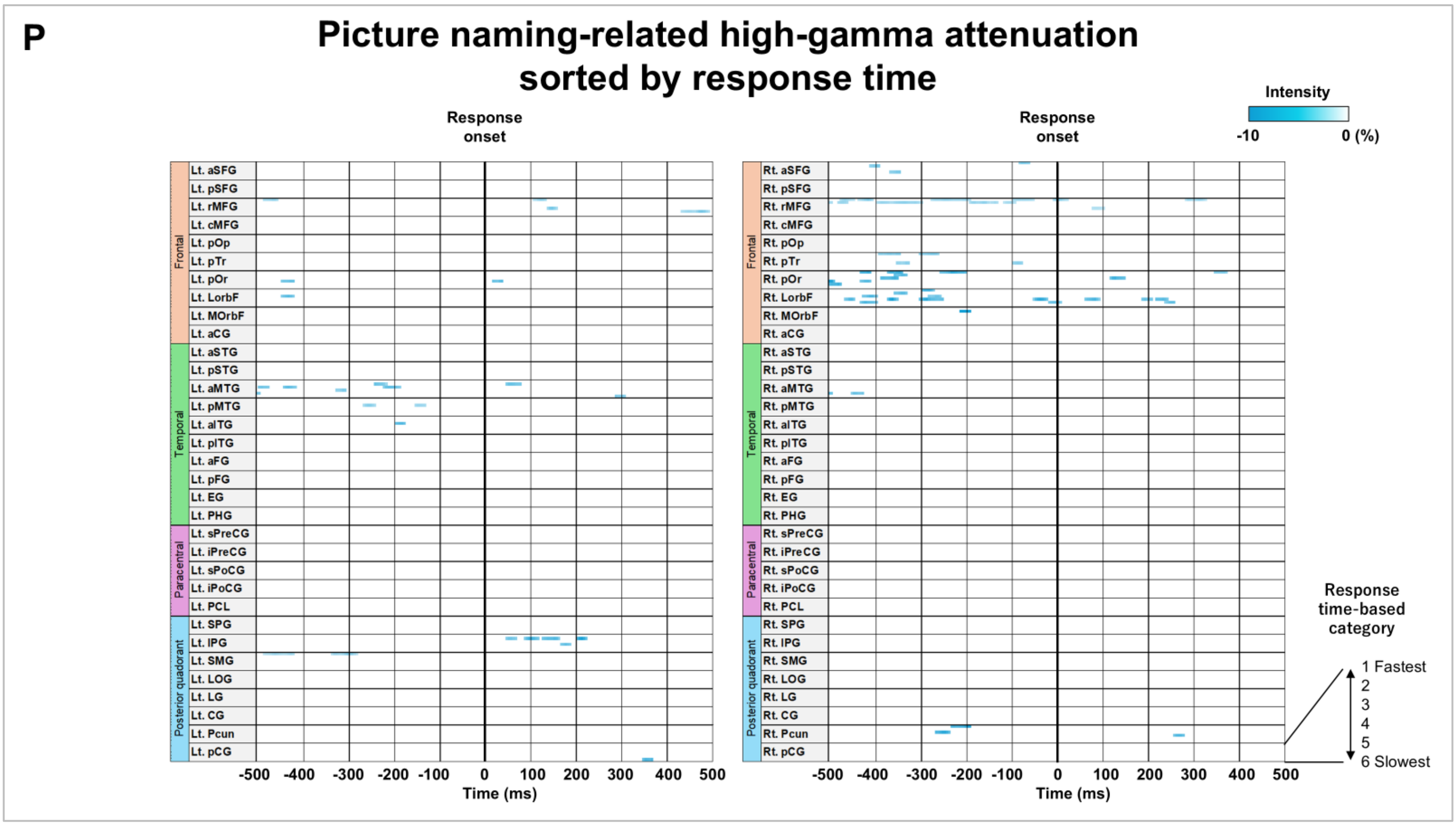

P
Picture naming-related high-gamma attenuation
sorted by response time
Response onset
Response onset
Intensity
-10
0 (%)
Frontal
Temporal
Paracentral
Posterior quadorant
Lt. aSFG
Lt. pSFG
Lt. rMFG
Lt. cMFG
Lt. pOp
Lt. pTr
Lt. pOr
Lt. LorbF
Lt. MOrbF
Lt. aCG
Lt. aSTG
Lt. pSTG
Lt. aMTG
Lt. pMTG
Lt. aITG
Lt. pITG
Lt. aFG
Lt. pFG
Lt. EG
Lt. PHG
Lt. sPreCG
Lt. iPreCG
Lt. sPoCG
Lt. iPoCG
Lt. PCL
Lt. SPG
Lt. IPG
Lt. SMG
Lt. LOG
Lt. LG
Lt. CG
Lt. Pcun
Lt. pCG
Rt. aSFG
Rt. pSFG
Rt. rMFG
Rt. cMFG
Rt. pOp
Rt. pTr
Rt. pOr
Rt. LorbF
Rt. MOrbF
Rt. aCG
Rt. aSTG
Rt. pSTG
Rt. aMTG
Rt. pMTG
Rt. aITG
Rt. pITG
Rt. aFG
Rt. pFG
Rt. EG
Rt. PHG
Rt. sPreCG
Rt. iPreCG
Rt. sPoCG
Rt. iPoCG
Rt. PCL
Rt. SPG
Rt. IPG
Rt. SMG
Rt. LOG
Rt. LG
Rt. CG
Rt. Pcun
Rt. pCG
-500 -400 -300 -200 -100 0 100 200 300 400 500
Time (ms)
-500 -400 -300 -200 -100 0 100 200 300 400 500
Time (ms)
Response time-based category
1 Fastest
2
3
4
5
6 Slowest


eFigure 10

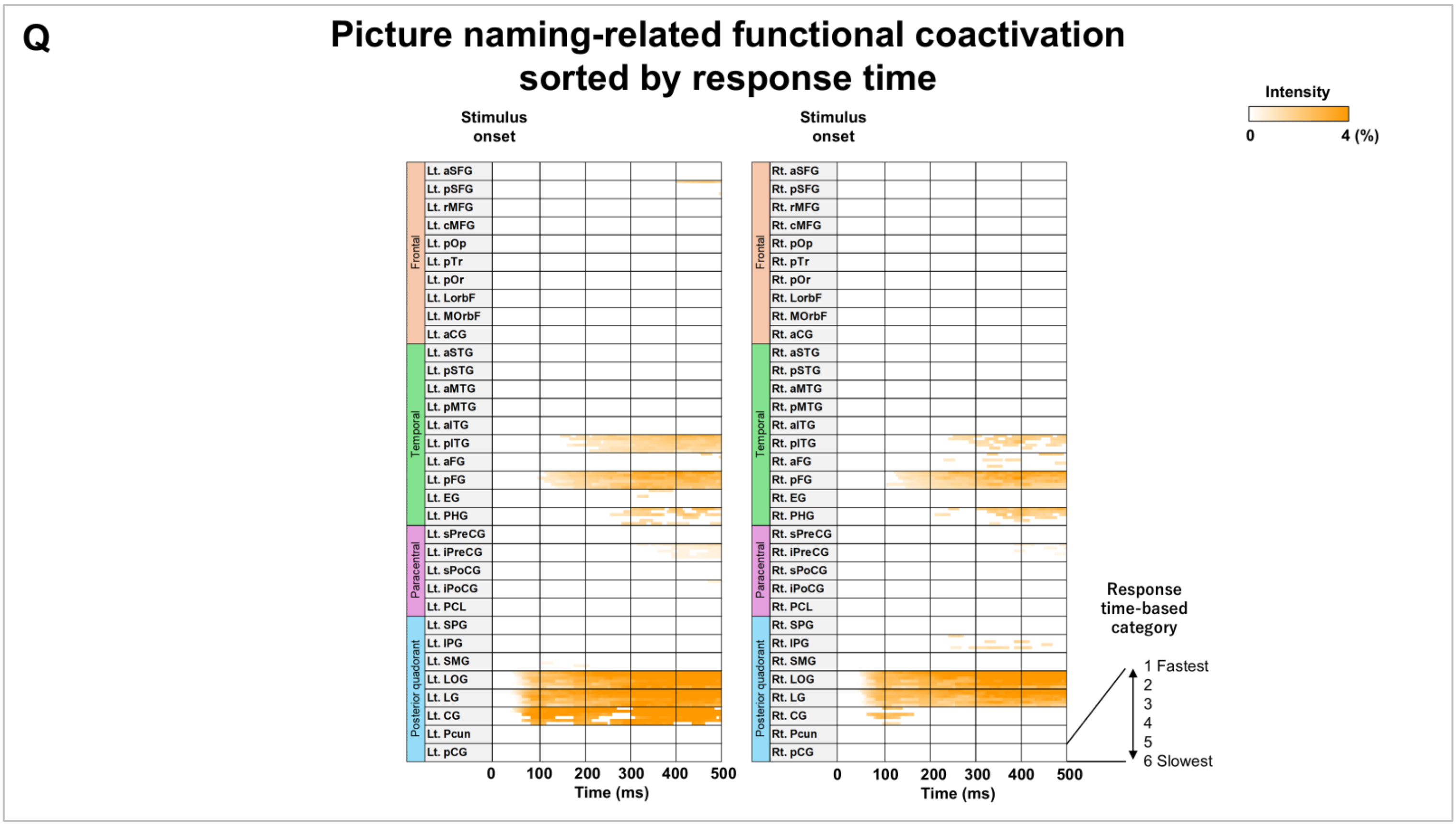
Q
Picture naming-related functional coactivation sorted by response time
Stimulus onset
Stimulus onset
Intensity
0
4 (%)
Frontal
Temporal
Paracentral
Posterior quadorant
Lt. aSFG
Lt. pSFG
Lt. rMFG
Lt. cMFG
Lt. pOp
Lt. pTr
Lt. pOr
Lt. LorbF
Lt. MOrbF
Lt. aCG
Lt. aSTG
Lt. pSTG
Lt. aMTG
Lt. pMTG
Lt. aITG
Lt. pITG
Lt. aFG
Lt. pFG
Lt. EG
Lt. PHG
Lt. sPreCG
Lt. iPreCG
Lt. sPoCG
Lt. iPoCG
Lt. PCL
Lt. SPG
Lt. IPG
Lt. SMG
Lt. LOG
Lt. LG
Lt. CG
Lt. Pcun
Lt. pCG
Rt. aSFG
Rt. pSFG
Rt. rMFG
Rt. cMFG
Rt. pOp
Rt. pTr
Rt. pOr
Rt. LorbF
Rt. MOrbF
Rt. aCG
Rt. aSTG
Rt. pSTG
Rt. aMTG
Rt. pMTG
Rt. aITG
Rt. pITG
Rt. aFG
Rt. pFG
Rt. EG
Rt. PHG
Rt. sPreCG
Rt. iPreCG
Rt. sPoCG
Rt. iPoCG
Rt. PCL
Rt. SPG
Rt. IPG
Rt. SMG
Rt. LOG
Rt. LG
Rt. CG
Rt. Pcun
Rt. pCG
0 100 200 300 400 500
Time (ms)
0 100 200 300 400 500
Time (ms)
Response time-based category
1 Fastest
2
3
4
5
6 Slowest

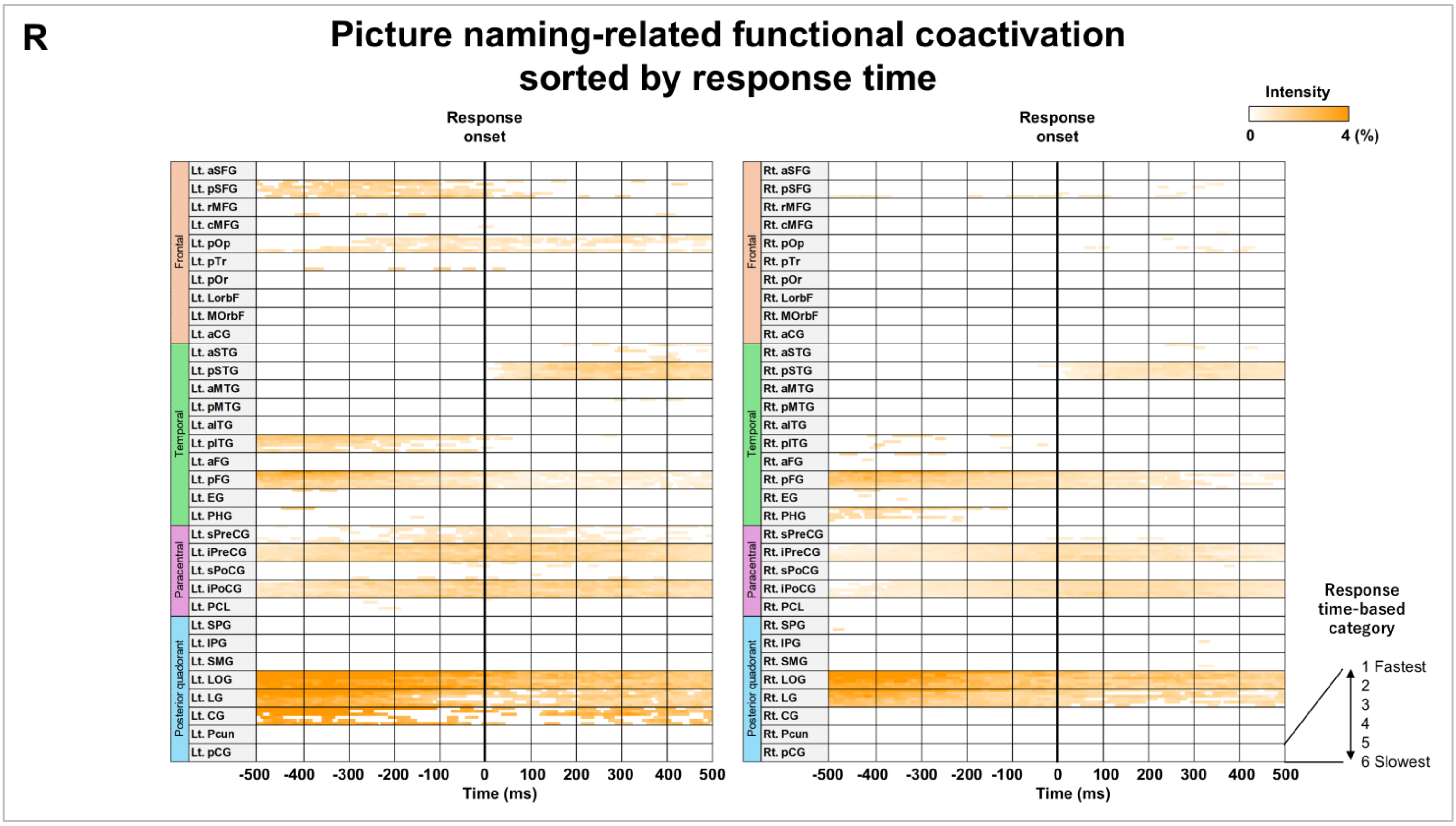
R
Picture naming-related functional coactivation sorted by response time
Response onset
Response onset
Intensity
0
4 (%)
Frontal
Temporal
Paracentral
Posterior quadorant
Lt. aSFG
Lt. pSFG
Lt. rMFG
Lt. cMFG
Lt. pOp
Lt. pTr
Lt. pOr
Lt. LorbF
Lt. MOrbF
Lt. aCG
Lt. aSTG
Lt. pSTG
Lt. aMTG
Lt. pMTG
Lt. aITG
Lt. pITG
Lt. aFG
Lt. pFG
Lt. EG
Lt. PHG
Lt. sPreCG
Lt. iPreCG
Lt. sPoCG
Lt. iPoCG
Lt. PCL
Lt. SPG
Lt. IPG
Lt. SMG
Lt. LOG
Lt. LG
Lt. CG
Lt. Pcun
Lt. pCG
Rt. aSFG
Rt. pSFG
Rt. rMFG
Rt. cMFG
Rt. pOp
Rt. pTr
Rt. pOr
Rt. LorbF
Rt. MOrbF
Rt. aCG
Rt. aSTG
Rt. pSTG
Rt. aMTG
Rt. pMTG
Rt. aITG
Rt. pITG
Rt. aFG
Rt. pFG
Rt. EG
Rt. PHG
Rt. sPreCG
Rt. iPreCG
Rt. sPoCG
Rt. iPoCG
Rt. PCL
Rt. SPG
Rt. IPG
Rt. SMG
Rt. LOG
Rt. LG
Rt. CG
Rt. Pcun
Rt. pCG
-500 -400 -300 -200 -100 0 100 200 300 400 500
Time (ms)
-500 -400 -300 -200 -100 0 100 200 300 400 500
Time (ms)
Response time-based category
1 Fastest
2
3
4
5
6 Slowest

eFigure 10

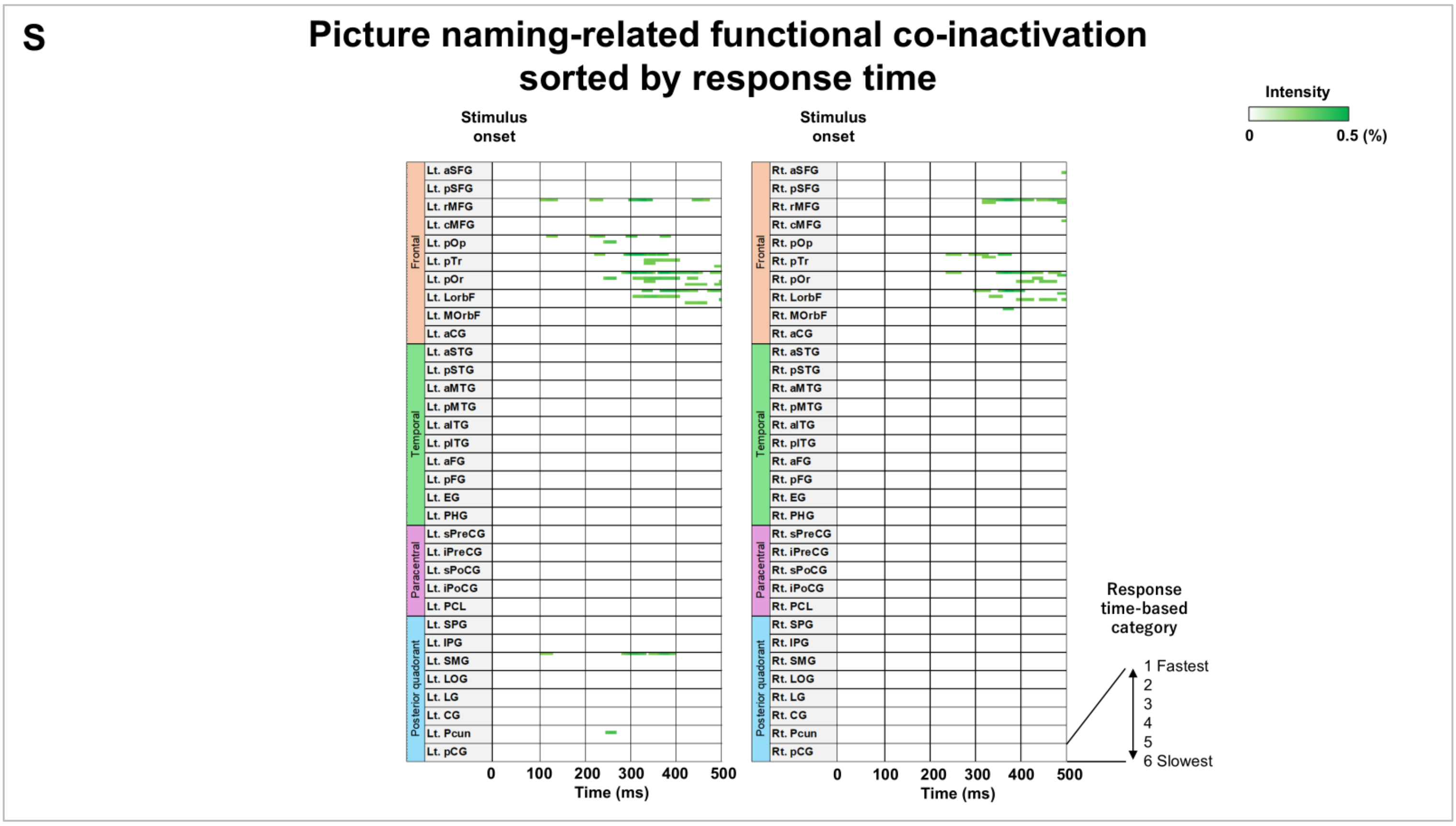


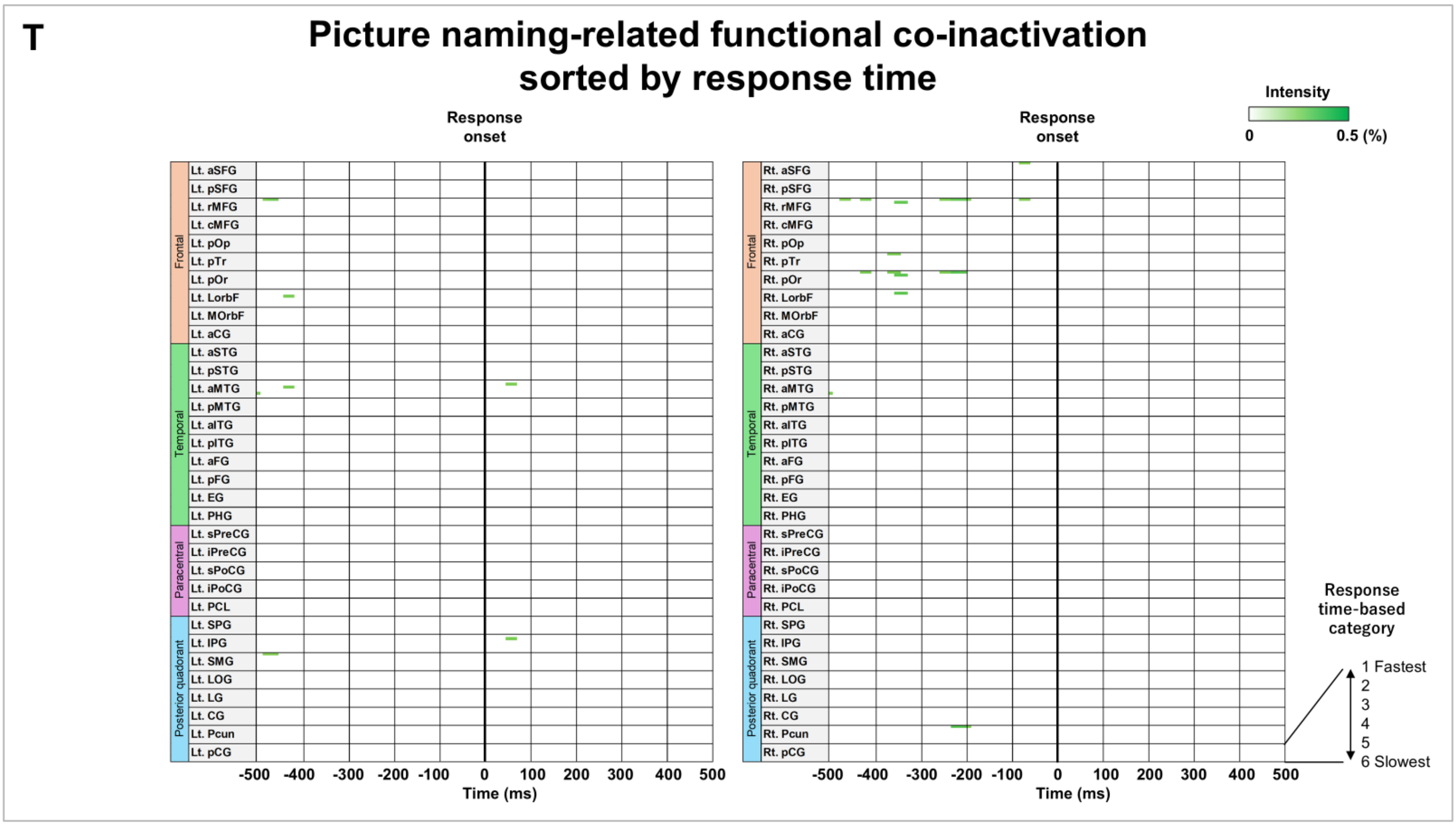


**eFigure 10. Temporal dynamics of significant neural modulations sorted by response times.** **A-C** Auditory naming-related high-gamma augmentation. **D-F** Auditory naming-related high-gamma attenuation. **G-I** Auditory naming-related functional coactivation. **J-L** Auditory naming-related functional co-inactivation. **M-N** Picture naming-related high-gamma augmentation. **O-P** Picture naming-related high-gamma attenuation. **Q-R** Picture naming-related functional coactivation. **S-T** Picture naming-related functional co-inactivation.

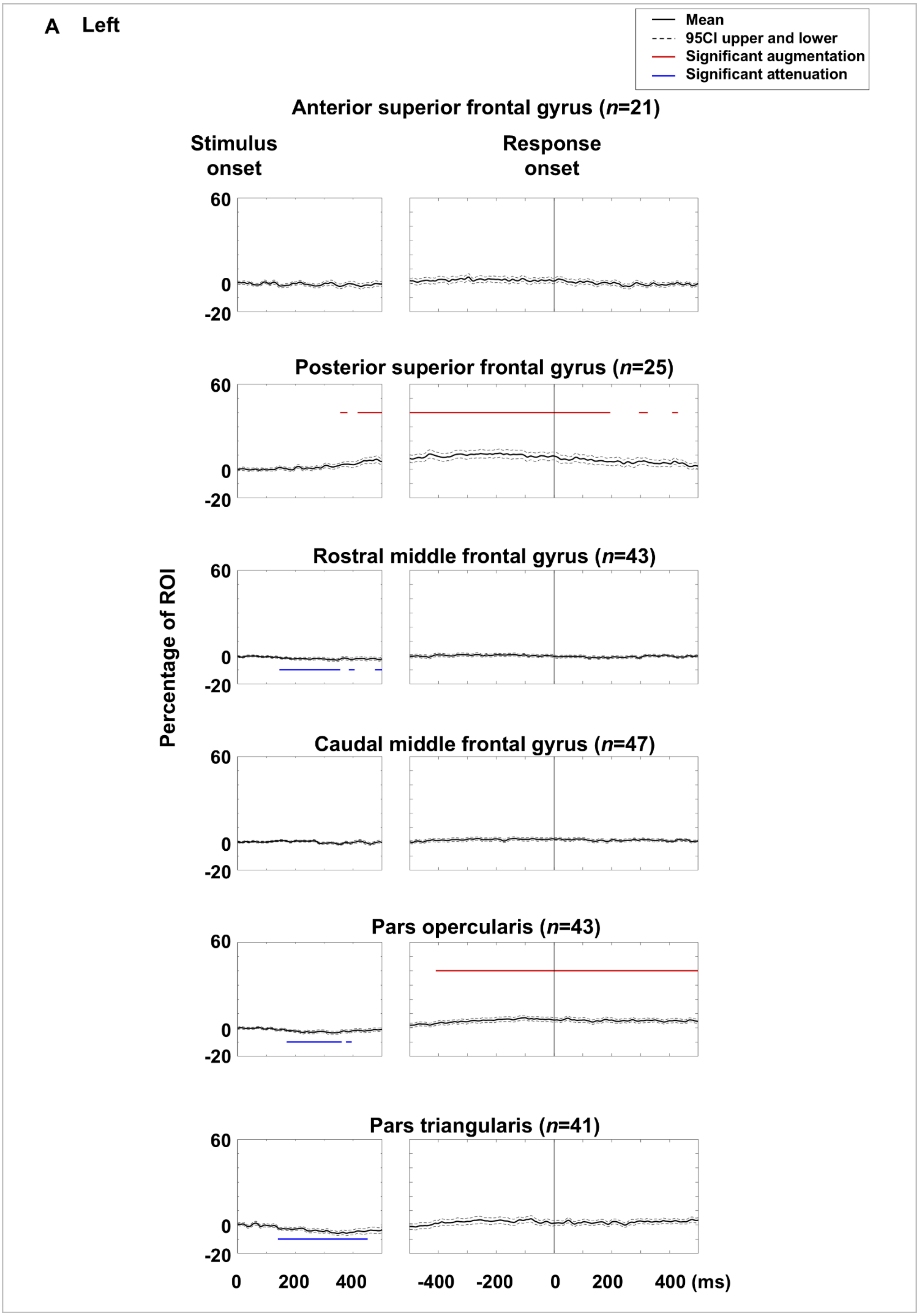
A Left
Mean
95CI upper and lower
Significant augmentation
Significant attenuation
Anterior superior frontal gyrus (n=21)
Stimulus onset
Response onset
Posterior superior frontal gyrus (n=25)
Rostral middle frontal gyrus (n=43)
Caudal middle frontal gyrus (n=47)
Pars opercularis (n=43)
Pars triangularis (n=41)
Percentage of ROI
60
0
-20
0 200 400
-400 -200 0 200 400 (ms)

eFigure 11

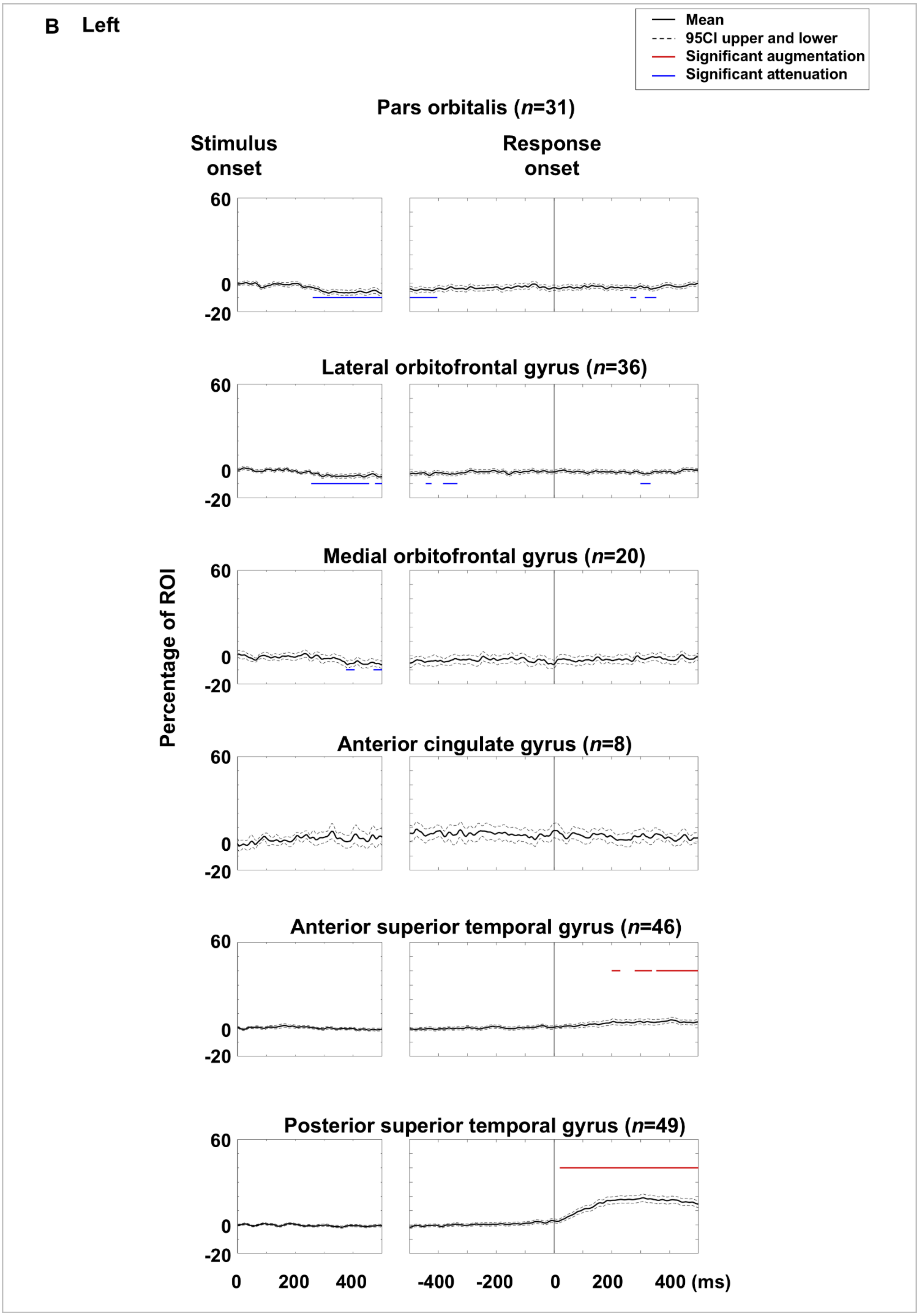
B Left
Mean
95CI upper and lower
Significant augmentation
Significant attenuation
Pars orbitalis (n=31)
Stimulus onset
Response onset
Lateral orbitofrontal gyrus (n=36)
Medial orbitofrontal gyrus (n=20)
Anterior cingulate gyrus (n=8)
Anterior superior temporal gyrus (n=46)
Posterior superior temporal gyrus (n=49)
Percentage of ROI
60
0
-20
0
200
400
-400
-200
0
200
400 (ms)

eFigure 11

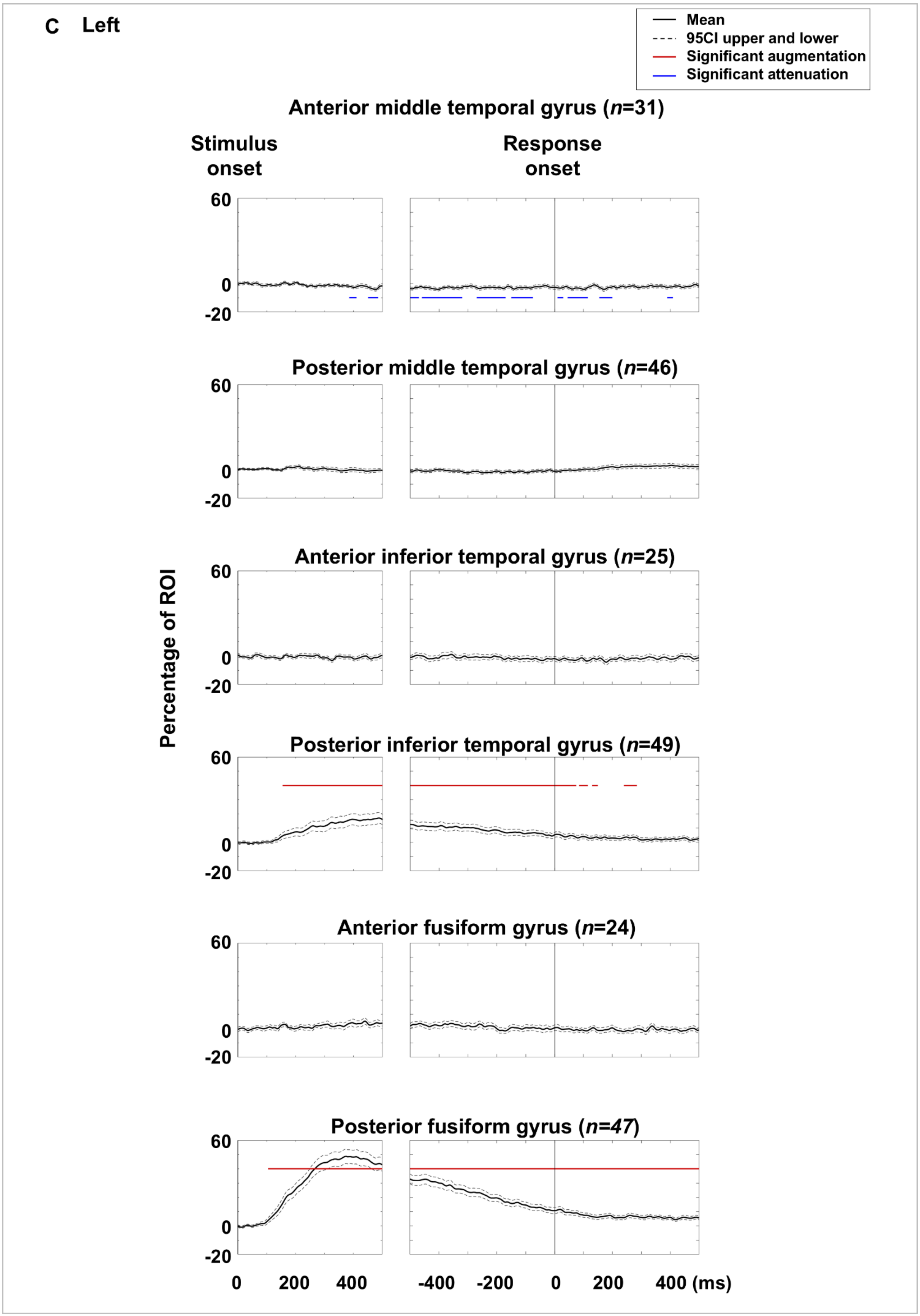
C Left
Mean
95CI upper and lower
Significant augmentation
Significant attenuation
Anterior middle temporal gyrus (n=31)
Stimulus onset
Response onset
Posterior middle temporal gyrus (n=46)
Anterior inferior temporal gyrus (n=25)
Posterior inferior temporal gyrus (n=49)
Anterior fusiform gyrus (n=24)
Posterior fusiform gyrus (n=47)
Percentage of ROI
60
0
-20
0
200
400
-400
-200
0
200
400 (ms)

eFigure 11

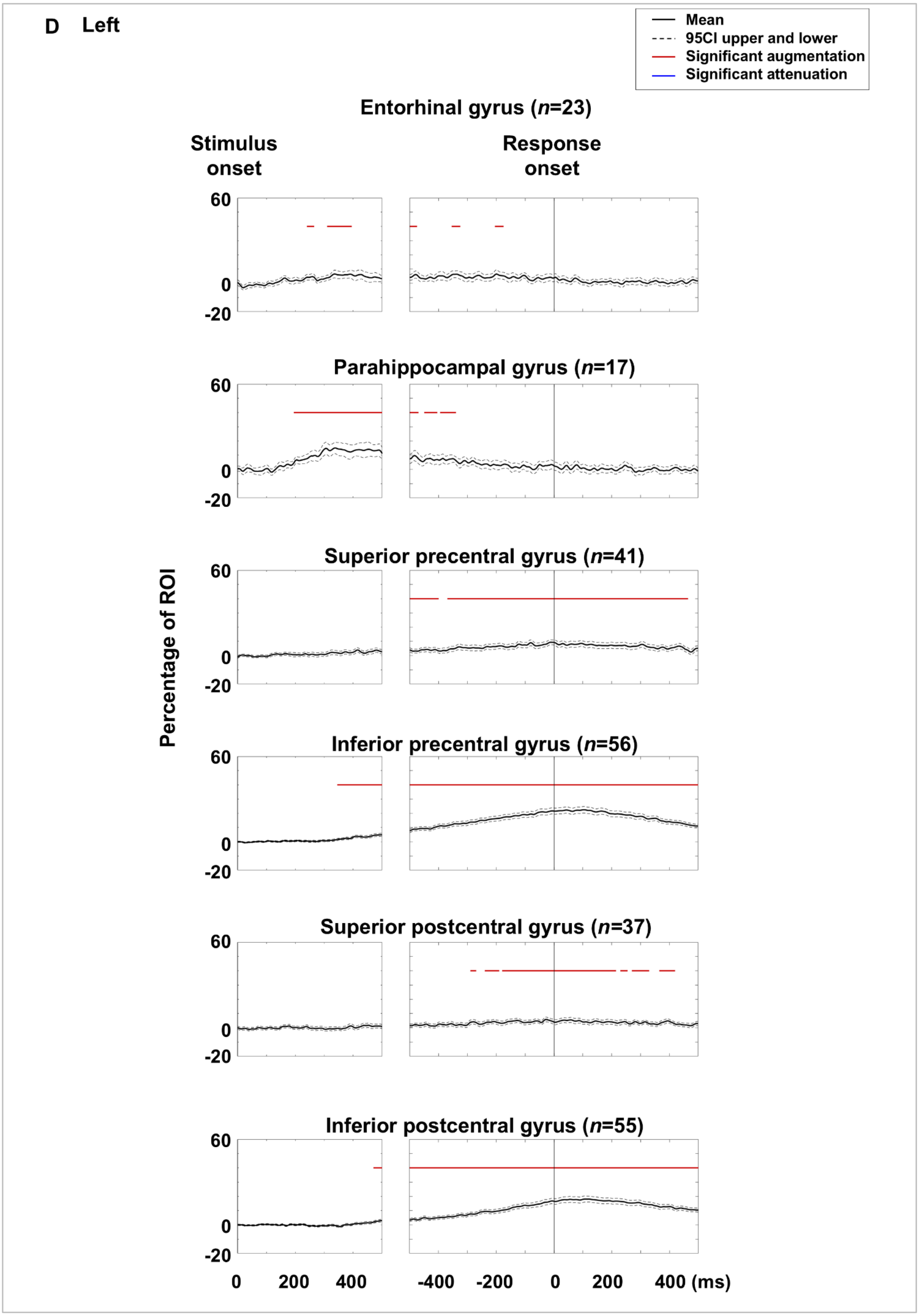
D Left
Mean
95CI upper and lower
Significant augmentation
Significant attenuation
Entorhinal gyrus (n=23)
Stimulus onset
Response onset
Parahippocampal gyrus (n=17)
Superior precentral gyrus (n=41)
Inferior precentral gyrus (n=56)
Superior postcentral gyrus (n=37)
Inferior postcentral gyrus (n=55)
Percentage of ROI
60
0
-20
0
200
400
-400
-200
0
200
400 (ms)

eFigure 11

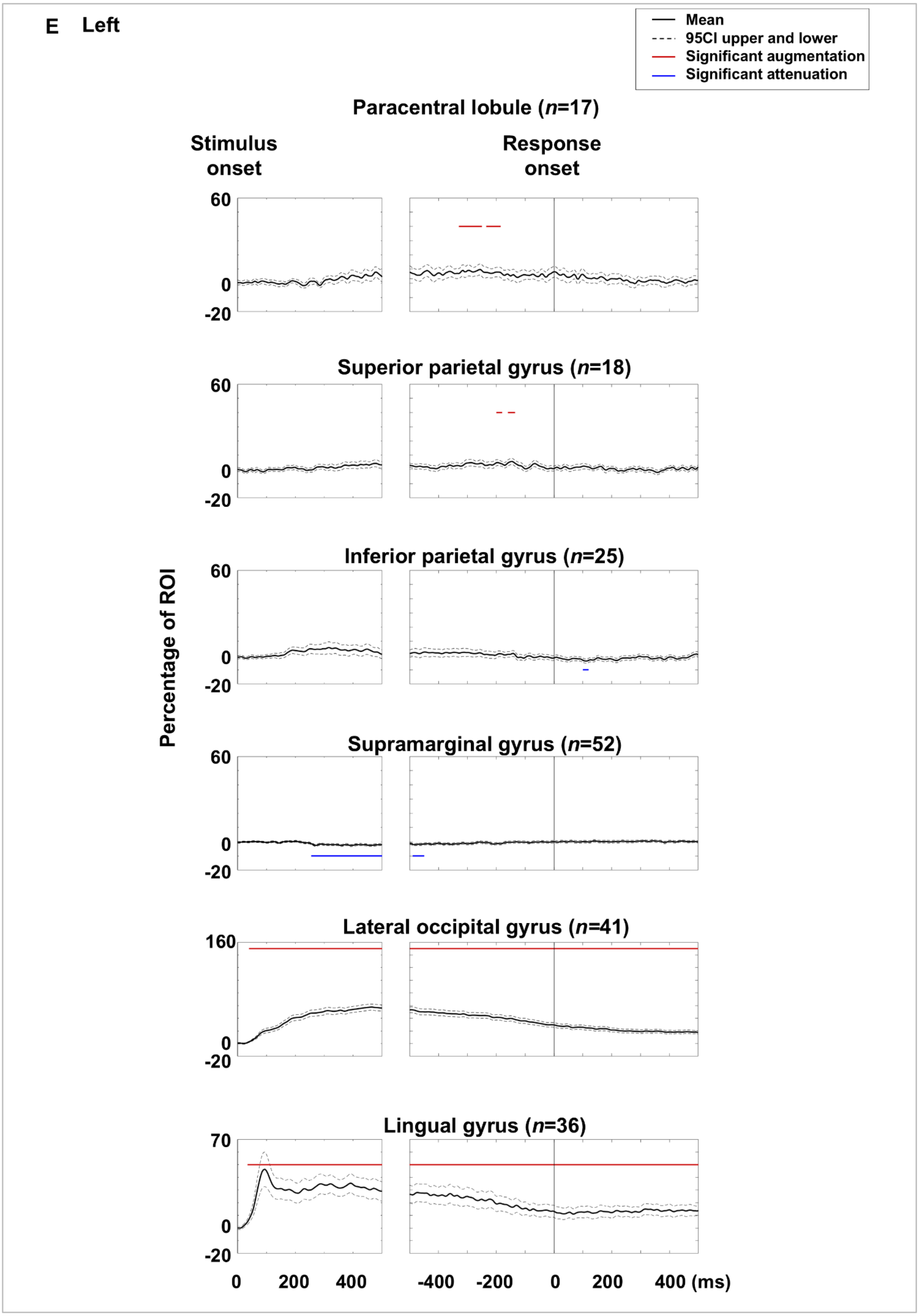
E Left
Mean
95CI upper and lower
Significant augmentation
Significant attenuation
Paracentral lobule (n=17)
Stimulus onset
Response onset
Superior parietal gyrus (n=18)
Inferior parietal gyrus (n=25)
Supramarginal gyrus (n=52)
Lateral occipital gyrus (n=41)
Lingual gyrus (n=36)
Percentage of ROI
60
0
-20
160
70
0 200 400
-400 -200 0 200 400 (ms)

eFigure 11

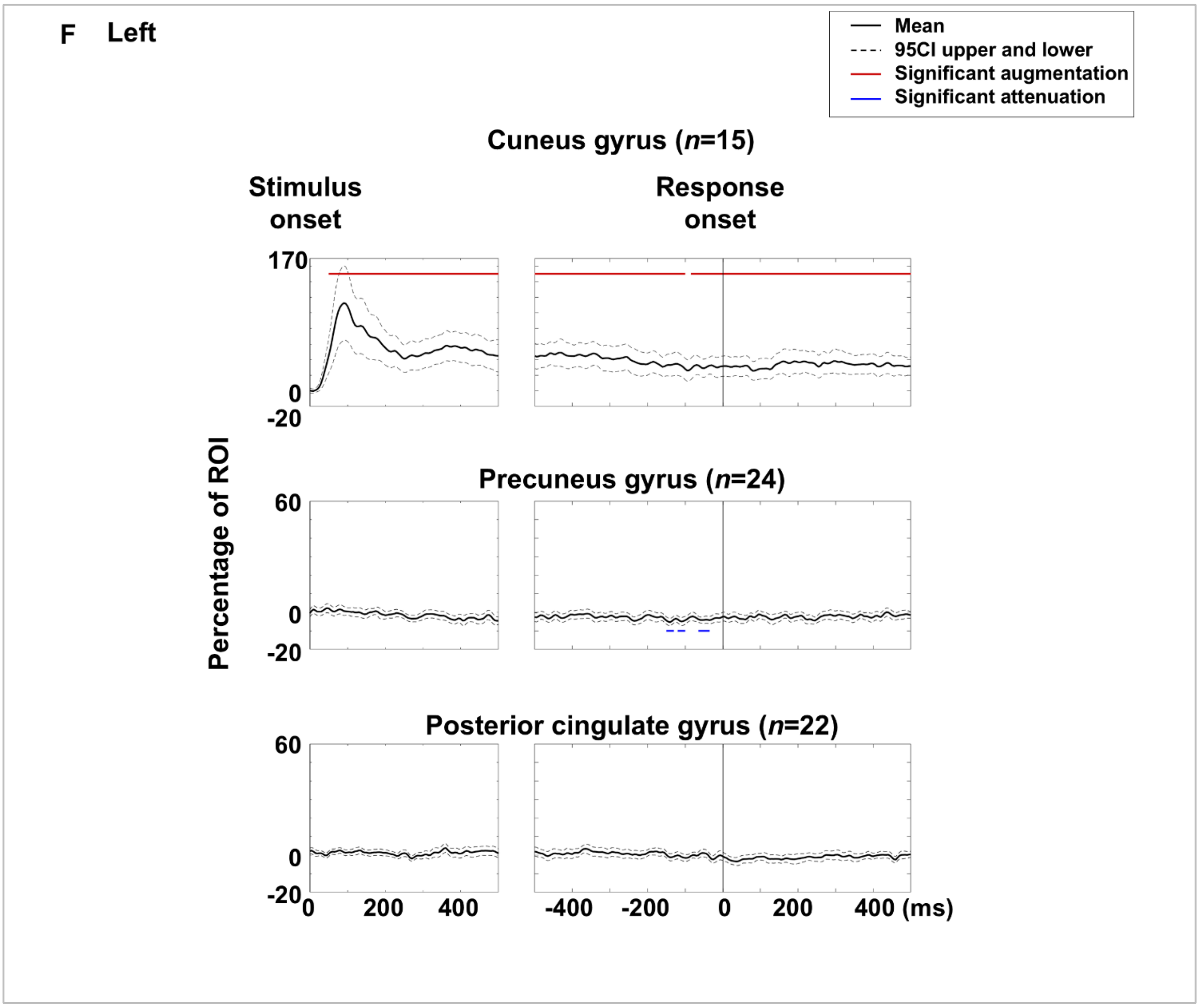
F Left
Mean
95CI upper and lower
Significant augmentation
Significant attenuation
Cuneus gyrus (n=15)
Stimulus onset
Response onset
170
0
-20
Percentage of ROI
Precuneus gyrus (n=24)
60
0
-20
Posterior cingulate gyrus (n=22)
60
0
-20
0
200
400
-400
-200
0
200
400 (ms)

eFigure 11

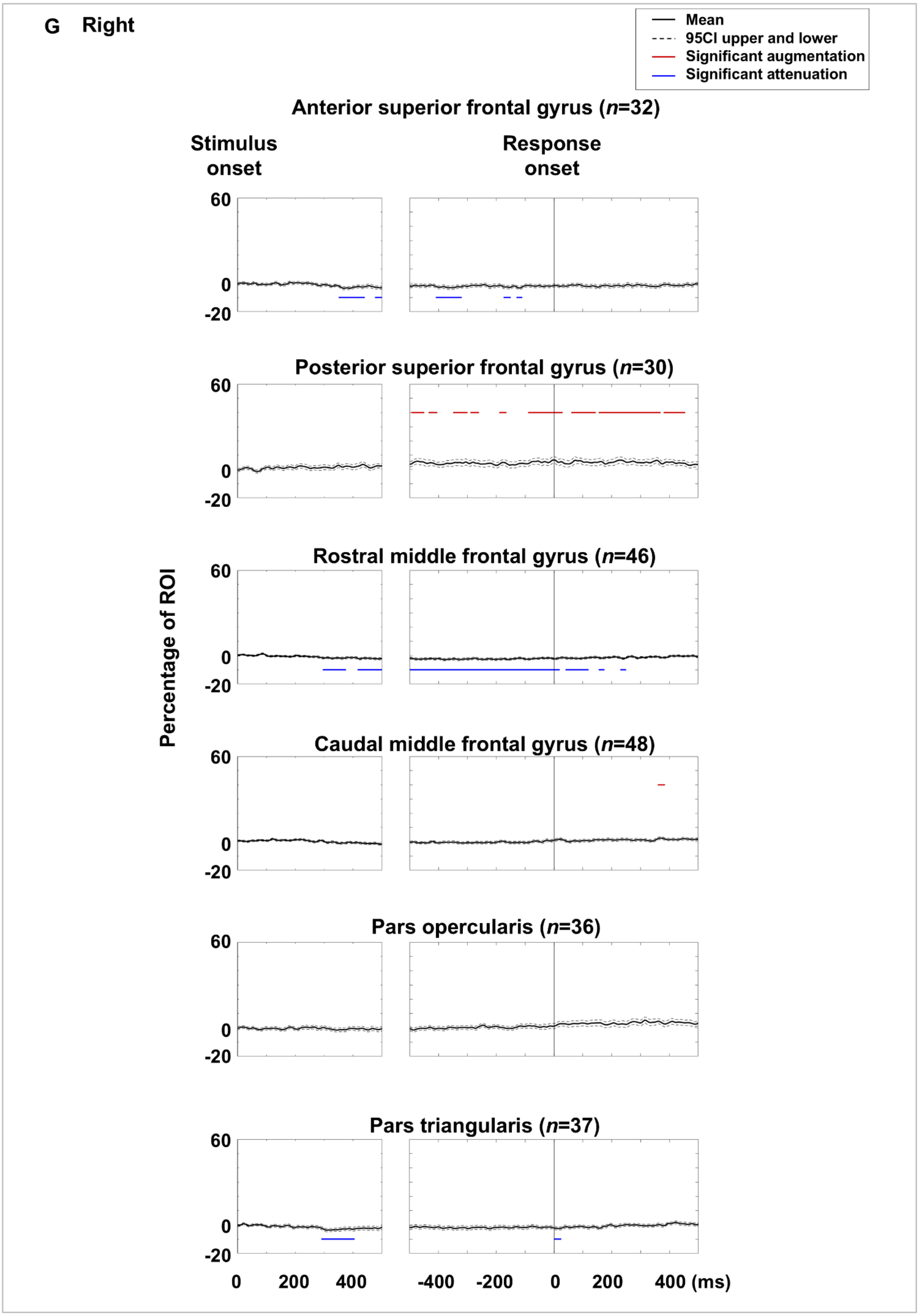
G Right
Mean
95CI upper and lower
Significant augmentation
Significant attenuation
Anterior superior frontal gyrus (n=32)
Stimulus onset
Response onset
Posterior superior frontal gyrus (n=30)
Rostral middle frontal gyrus (n=46)
Caudal middle frontal gyrus (n=48)
Pars opercularis (n=36)
Pars triangularis (n=37)
Percentage of ROI
60
0
-20
0 200 400
-400 -200 0 200 400 (ms)

eFigure 11

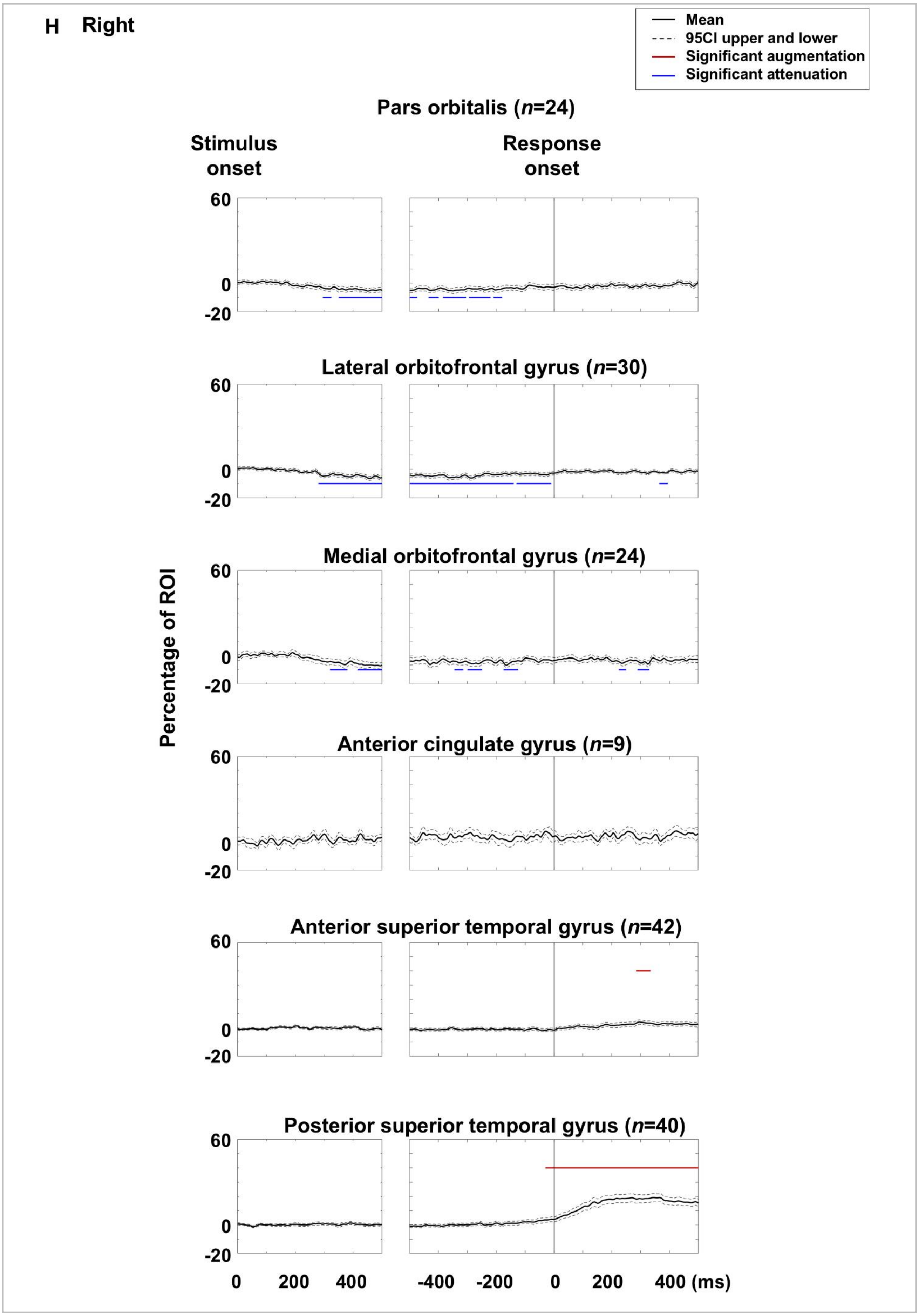
H Right
Mean
95CI upper and lower
Significant augmentation
Significant attenuation
Pars orbitalis (n=24)
Stimulus onset
Response onset
Lateral orbitofrontal gyrus (n=30)
Medial orbitofrontal gyrus (n=24)
Anterior cingulate gyrus (n=9)
Anterior superior temporal gyrus (n=42)
Posterior superior temporal gyrus (n=40)
Percentage of ROI
60
0
-20
0 200 400
-400 -200 0 200 400 (ms)

eFigure 11

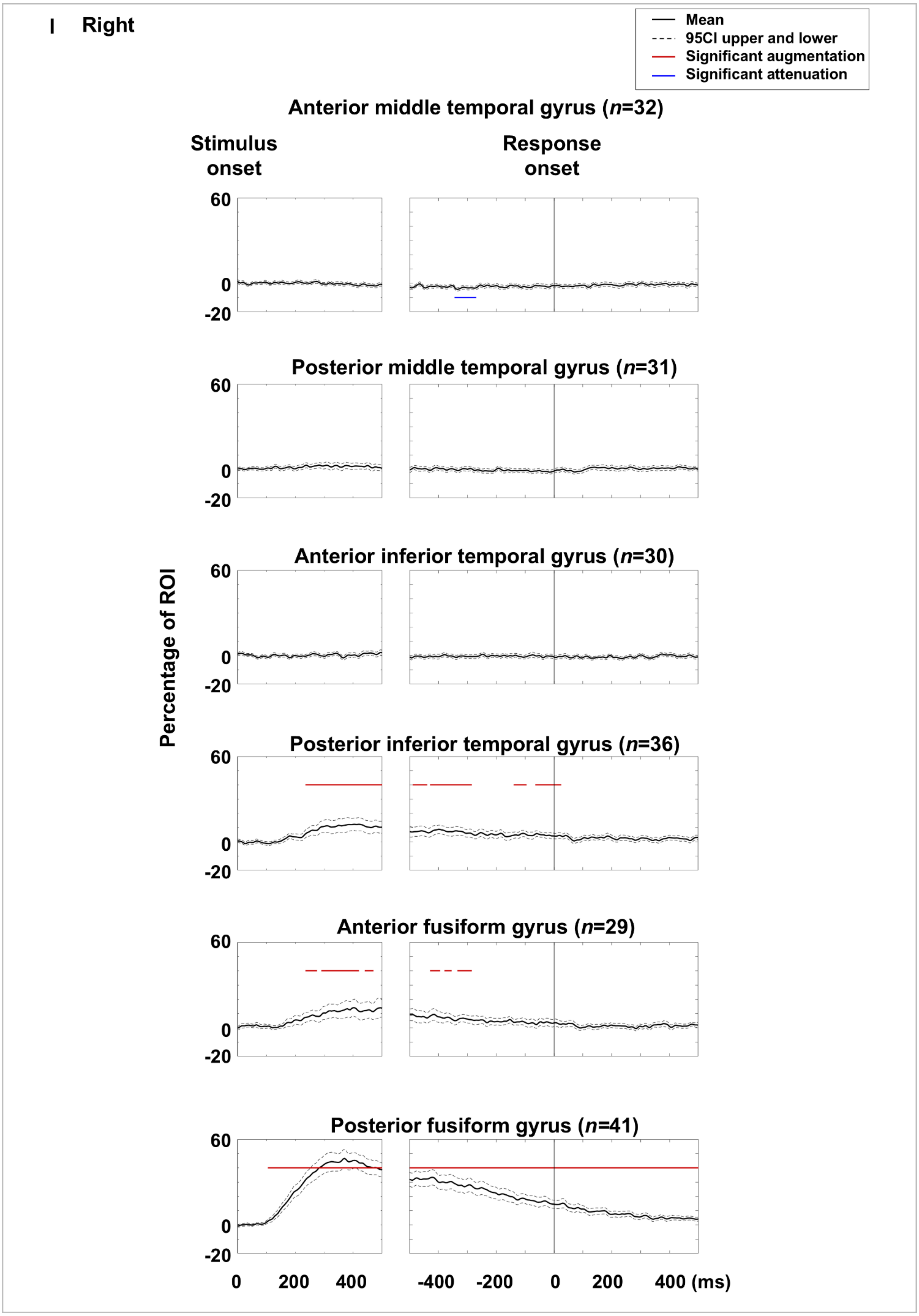
I Right
Mean
95CI upper and lower
Significant augmentation
Significant attenuation
Anterior middle temporal gyrus (n=32)
Stimulus onset
Response onset
Posterior middle temporal gyrus (n=31)
Anterior inferior temporal gyrus (n=30)
Posterior inferior temporal gyrus (n=36)
Anterior fusiform gyrus (n=29)
Posterior fusiform gyrus (n=41)
Percentage of ROI
60
0
-20
0
200
400
-400
-200
0
200
400 (ms)

eFigure 11

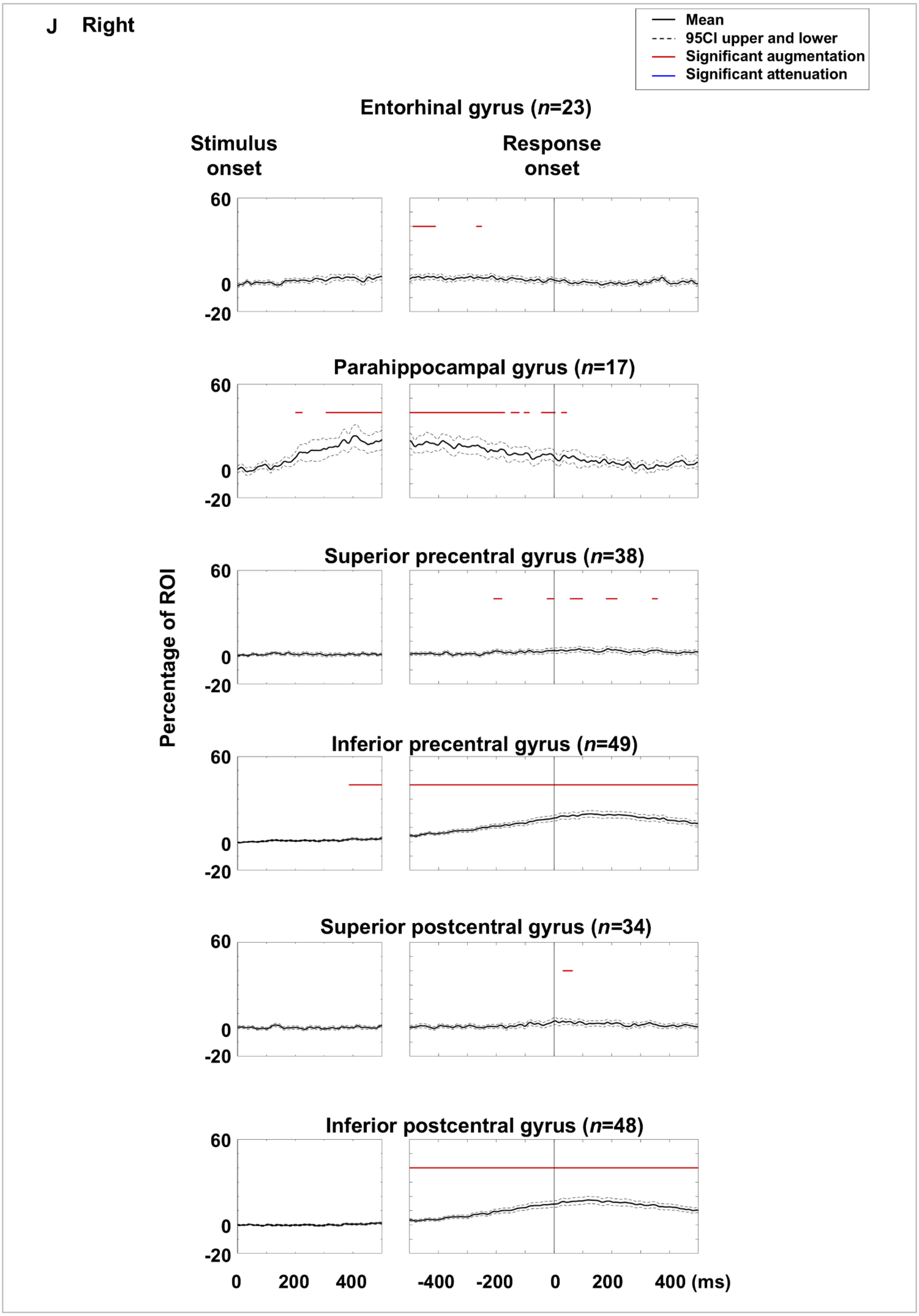

J Right
Mean
95CI upper and lower
Significant augmentation
Significant attenuation
Entorhinal gyrus (n=23)
Stimulus onset
Response onset
Parahippocampal gyrus (n=17)
Superior precentral gyrus (n=38)
Inferior precentral gyrus (n=49)
Superior postcentral gyrus (n=34)
Inferior postcentral gyrus (n=48)
Percentage of ROI
60
0
-20
0 200 400
-400 -200 0 200 400 (ms)


eFigure 11

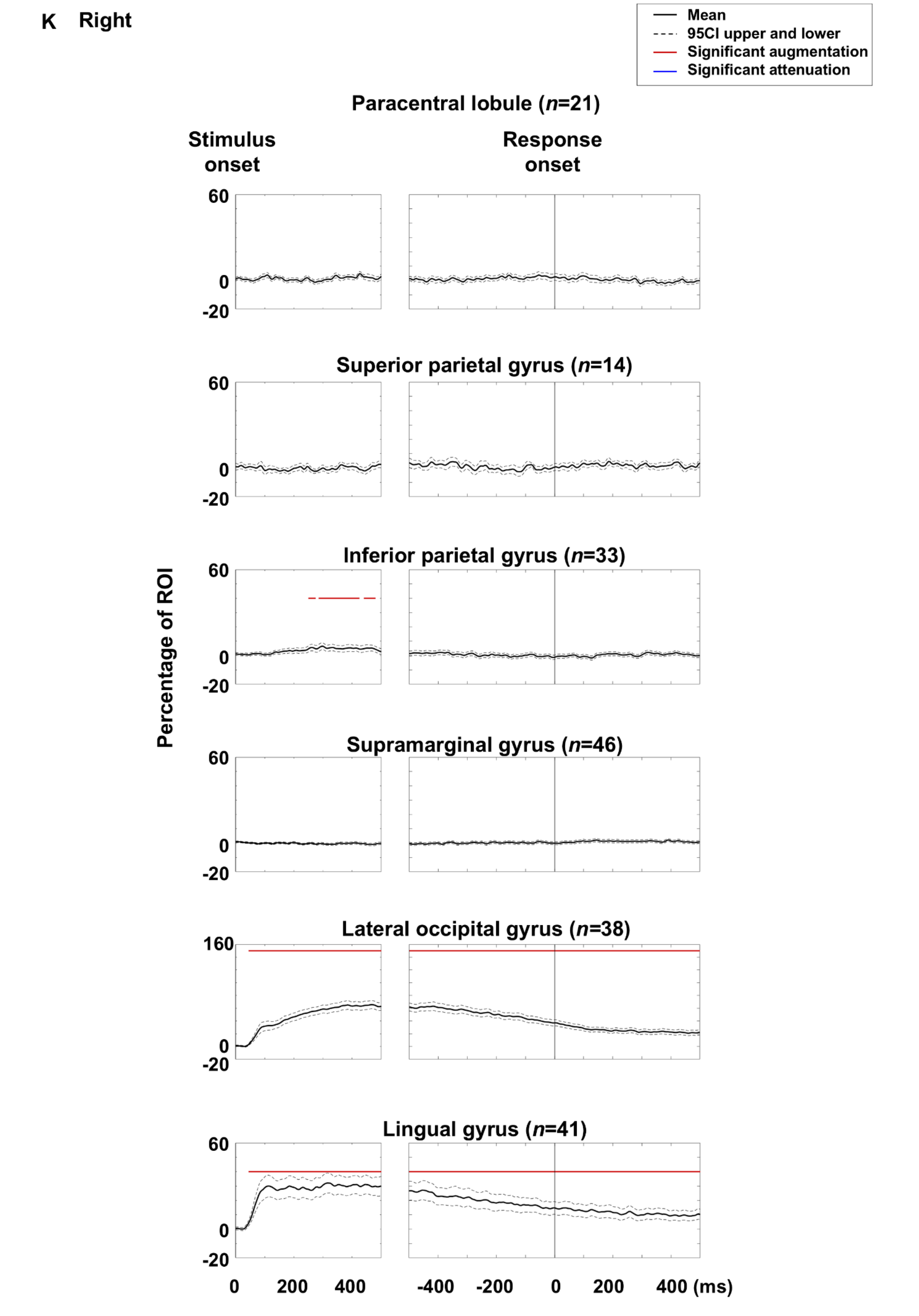
K Right
Mean
95CI upper and lower
Significant augmentation
Significant attenuation
Paracentral lobule (n=21)
Stimulus onset
Response onset
Superior parietal gyrus (n=14)
Inferior parietal gyrus (n=33)
Supramarginal gyrus (n=46)
Lateral occipital gyrus (n=38)
Lingual gyrus (n=41)
Percentage of ROI
60
0
-20
160
0
200
400
-400
-200
400 (ms)

eFigure 11

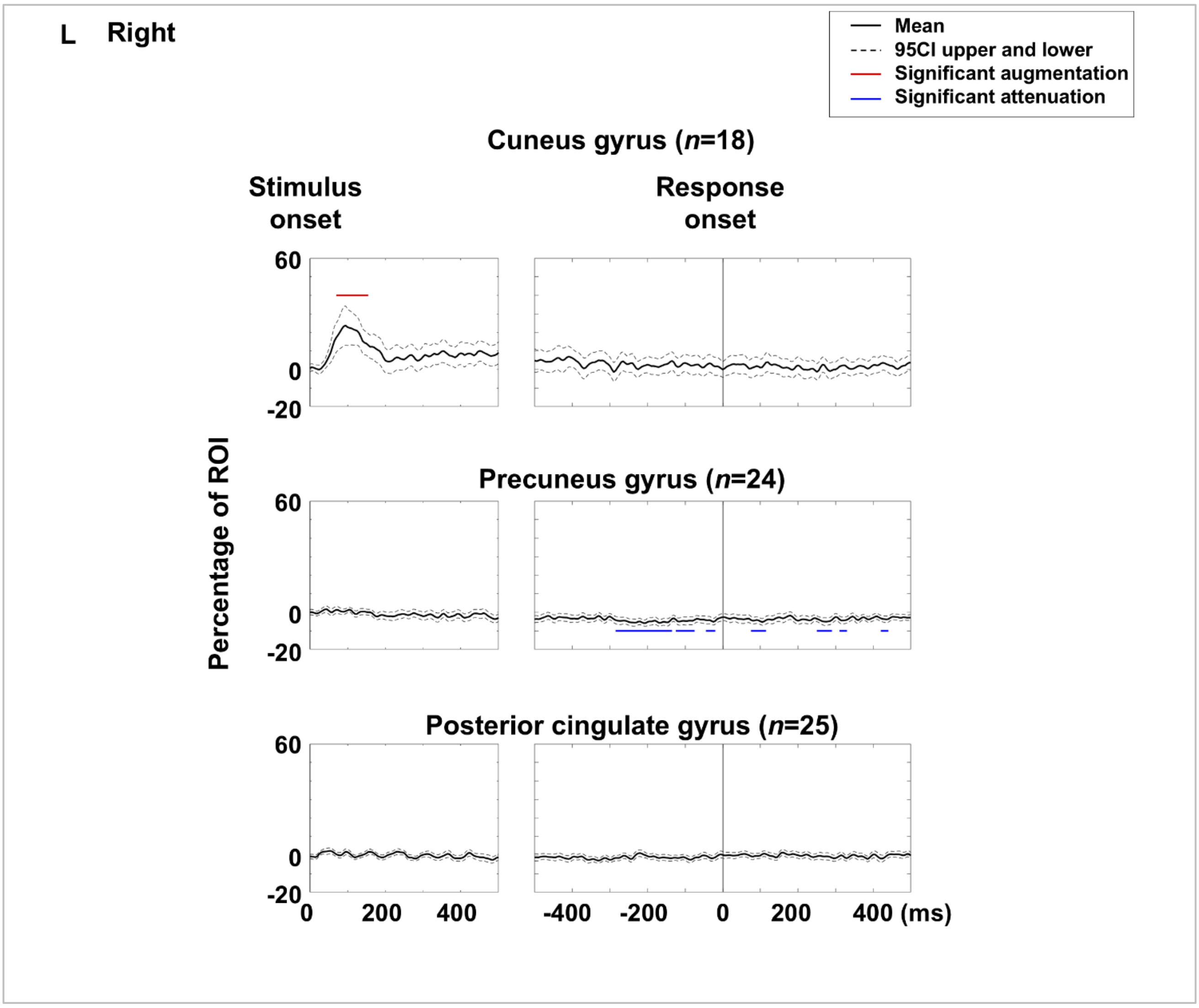


**eFigure 11. Picture naming-related high-gamma amplitude dynamics at regions of interest (ROIs).** Mean high-gamma amplitude (% change relative to baseline) at each ROI is shown with 95% confidence intervals. **A–F** Left hemispheric ROIs. **G-L** Right hemispheric ROIs.

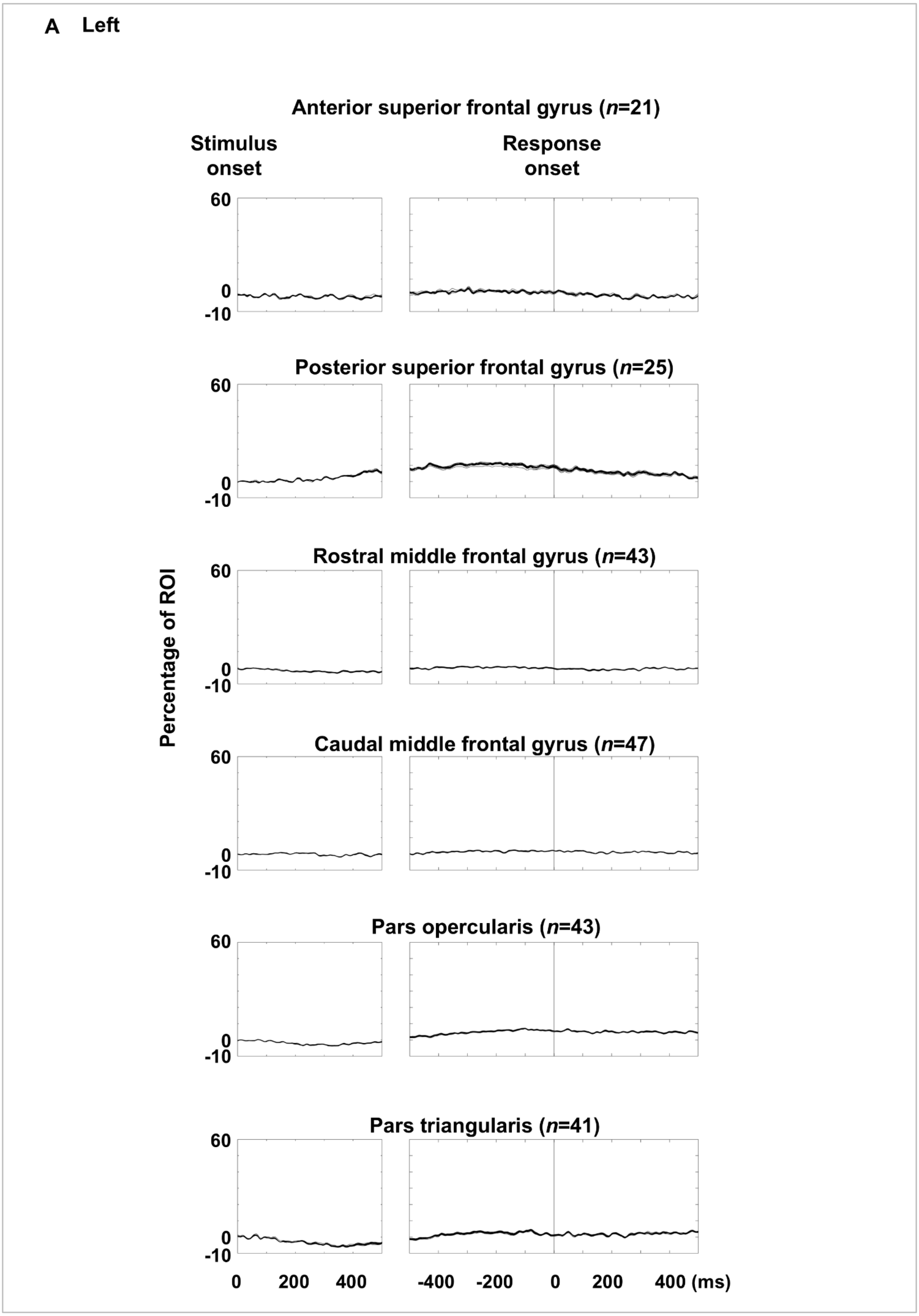
A Left
Anterior superior frontal gyrus (n=21)
Stimulus onset
Response onset
60
0
-10
Posterior superior frontal gyrus (n=25)
Rostral middle frontal gyrus (n=43)
Caudal middle frontal gyrus (n=47)
Pars opercularis (n=43)
Pars triangularis (n=41)
Percentage of ROI
0 200 400
-400 -200 0 200 400 (ms)

eFigure 12

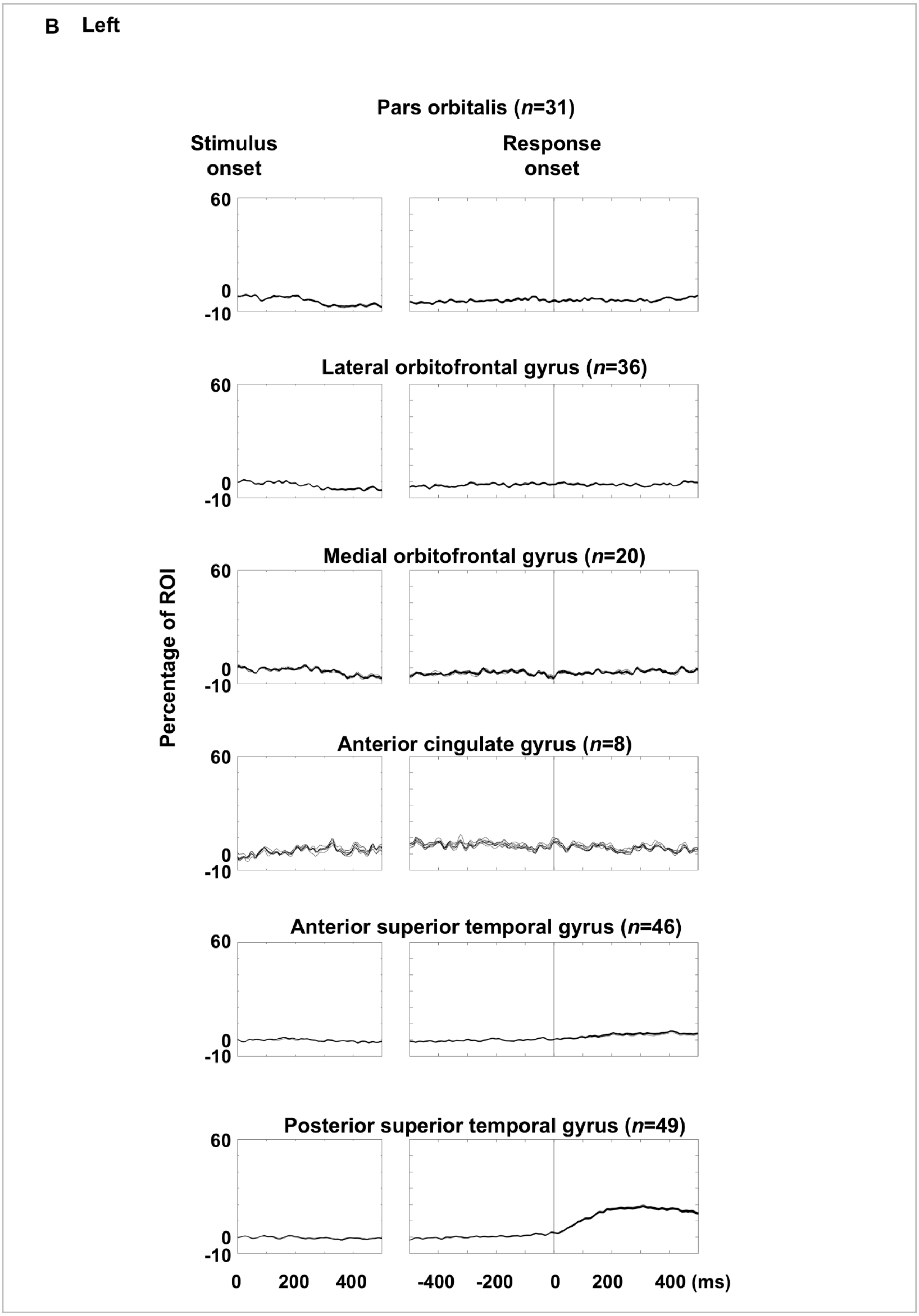
B Left
Pars orbitalis (n=31)
Stimulus onset
Response onset
Lateral orbitofrontal gyrus (n=36)
Medial orbitofrontal gyrus (n=20)
Anterior cingulate gyrus (n=8)
Anterior superior temporal gyrus (n=46)
Posterior superior temporal gyrus (n=49)
Percentage of ROI
60
0
-10
0 200 400
-400 -200 0 200 400 (ms)

eFigure 12

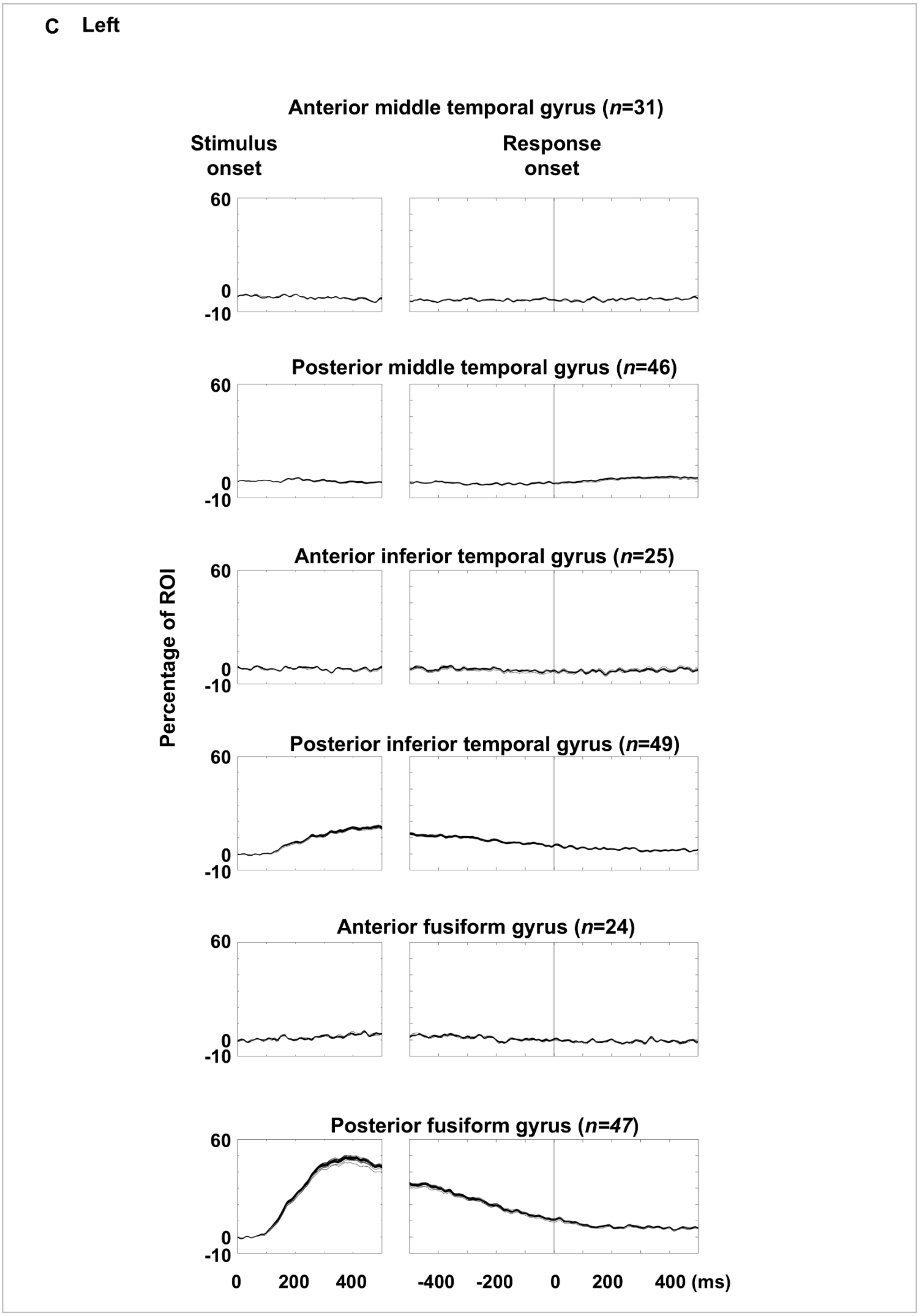
C Left
Anterior middle temporal gyrus (n=31)
Stimulus onset
Response onset
60
0
-10
Posterior middle temporal gyrus (n=46)
Anterior inferior temporal gyrus (n=25)
Percentage of ROI
Posterior inferior temporal gyrus (n=49)
Anterior fusiform gyrus (n=24)
Posterior fusiform gyrus (n=47)
0 200 400
-400 -200 0 200 400 (ms)

eFigure 12

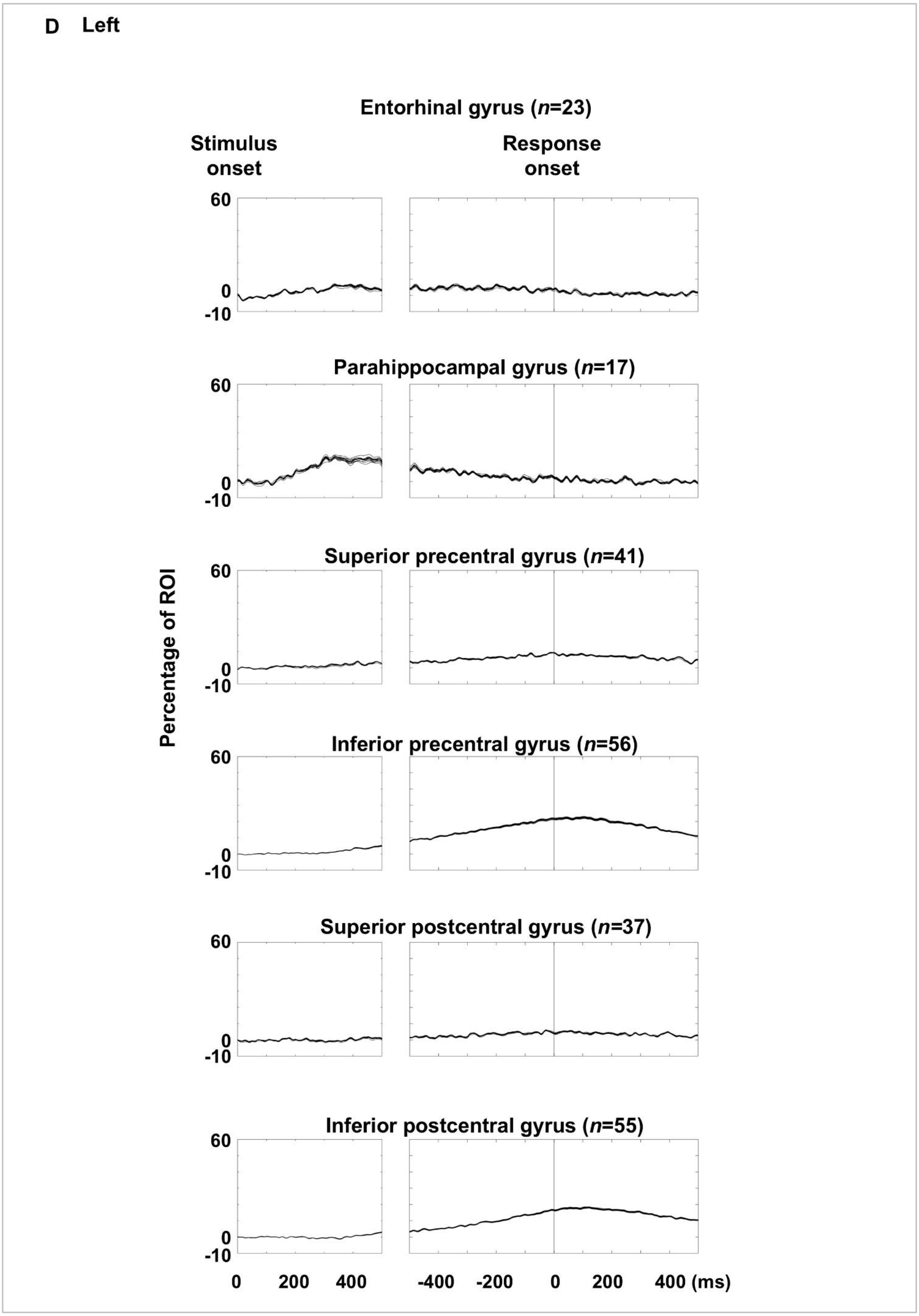
D Left
Entorhinal gyrus (n=23)
Stimulus onset
Response onset
Parahippocampal gyrus (n=17)
Superior precentral gyrus (n=41)
Inferior precentral gyrus (n=56)
Superior postcentral gyrus (n=37)
Inferior postcentral gyrus (n=55)
Percentage of ROI
60
0
-10
0 200 400
-400 -200 0 200 400 (ms)

eFigure 12

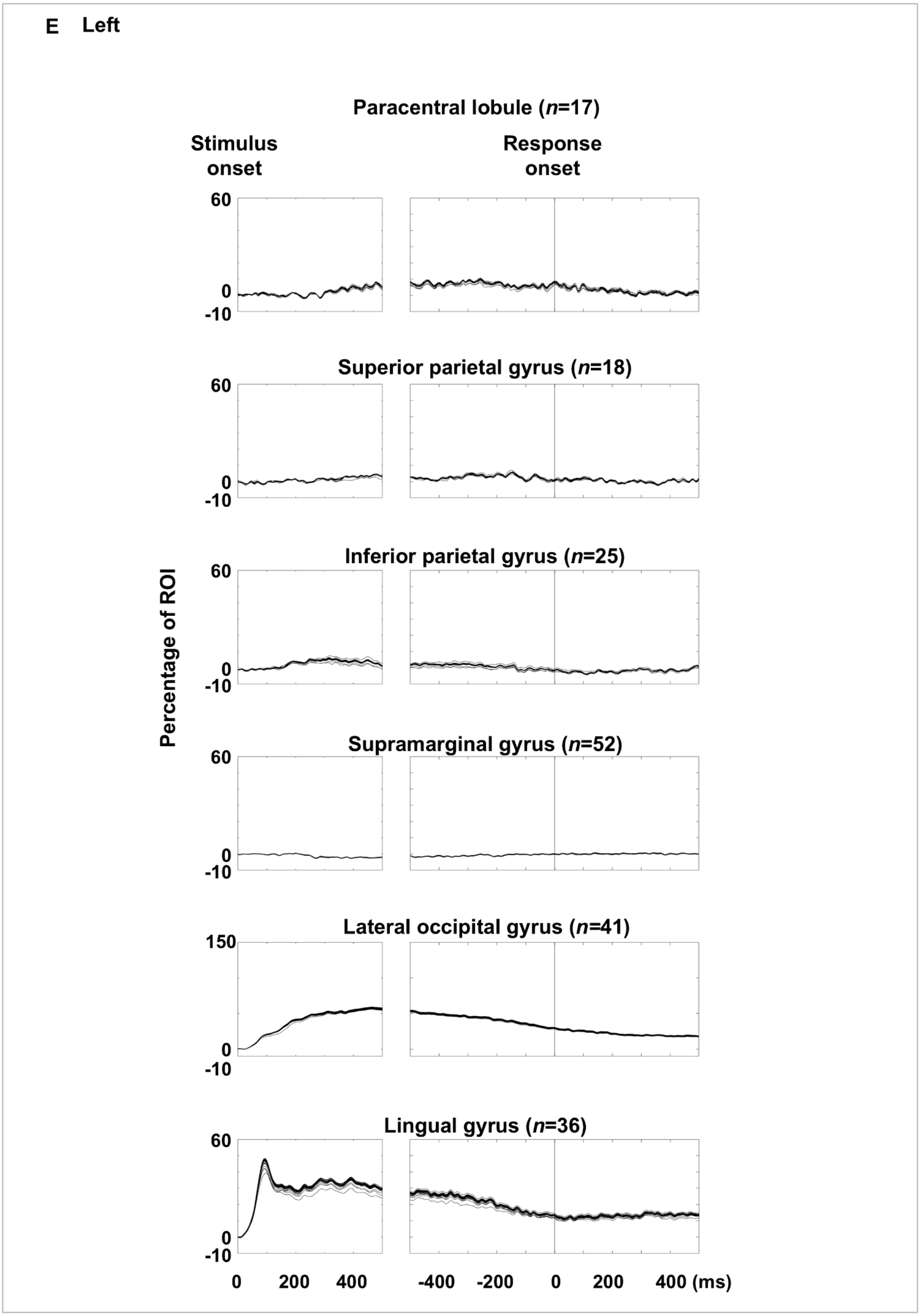
E Left
Paracentral lobule (n=17)
Stimulus onset
Response onset
60
0
-10
Superior parietal gyrus (n=18)
60
0
-10
Inferior parietal gyrus (n=25)
60
0
-10
Supramarginal gyrus (n=52)
60
0
-10
Lateral occipital gyrus (n=41)
150
0
-10
Lingual gyrus (n=36)
60
0
-10
Percentage of ROI
0 200 400
-400 -200 0 200 400 (ms)

eFigure 12

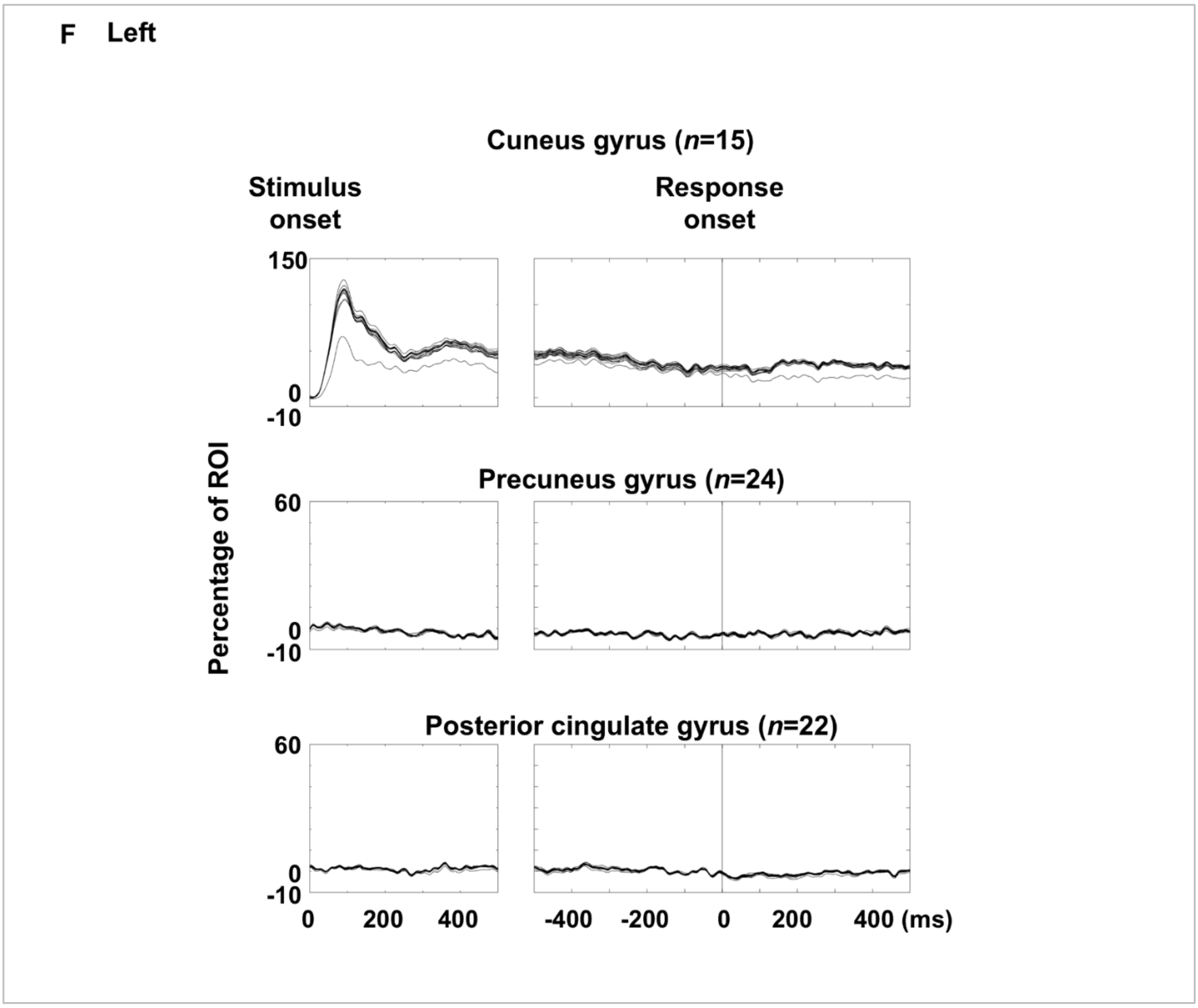

F Left
Cuneus gyrus (n=15)
Stimulus onset
Response onset
150
0
-10
Percentage of ROI
Precuneus gyrus (n=24)
60
0
-10
Posterior cingulate gyrus (n=22)
60
0
-10
0
200
400
-400
-200
0
200
400 (ms)


eFigure 12

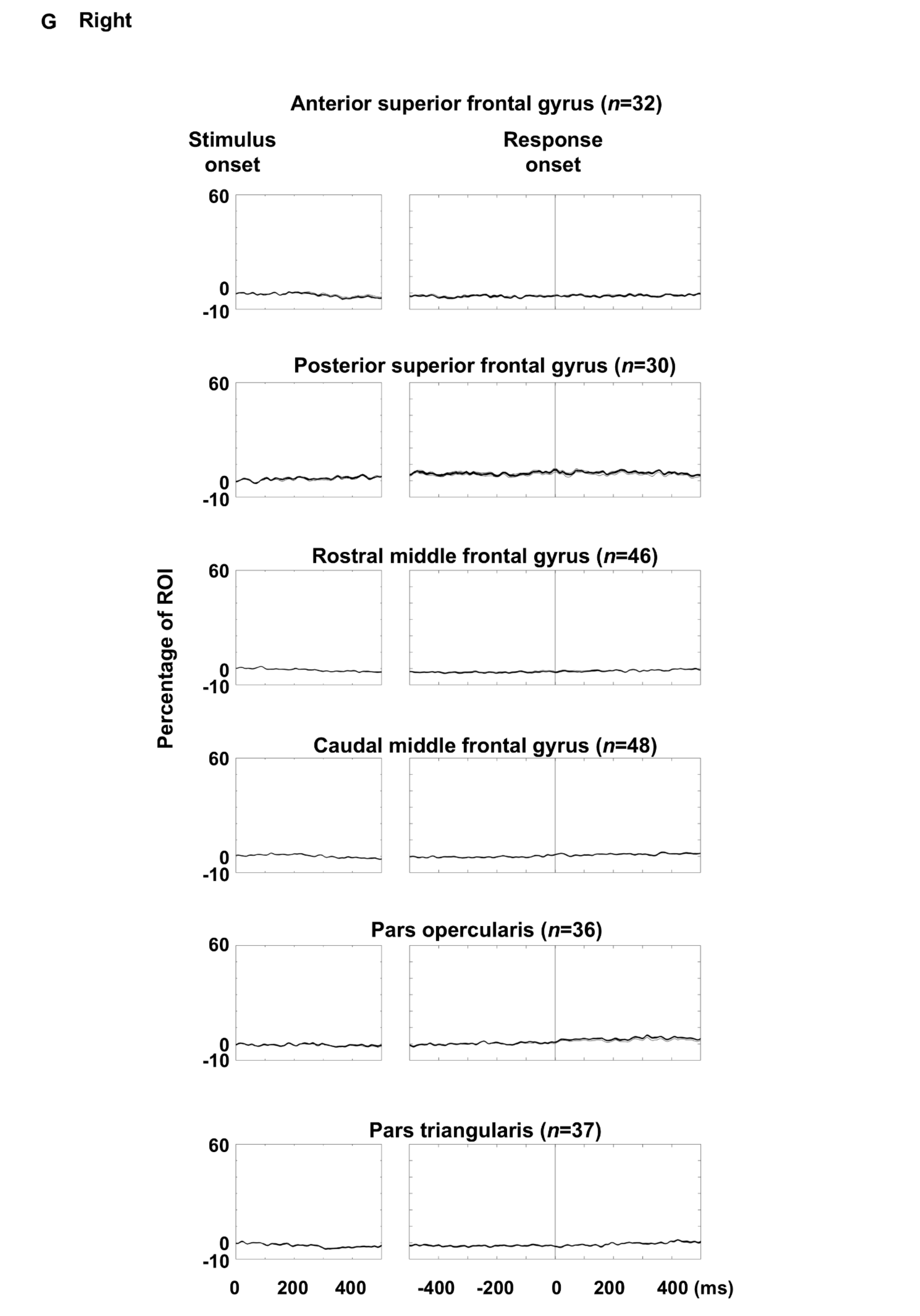
G Right
Anterior superior frontal gyrus (n=32)
Stimulus onset
Response onset
Posterior superior frontal gyrus (n=30)
Rostral middle frontal gyrus (n=46)
Caudal middle frontal gyrus (n=48)
Pars opercularis (n=36)
Pars triangularis (n=37)
Percentage of ROI
60
0
-10
0 200 400
-400 -200 0 200 400 (ms)

eFigure 12

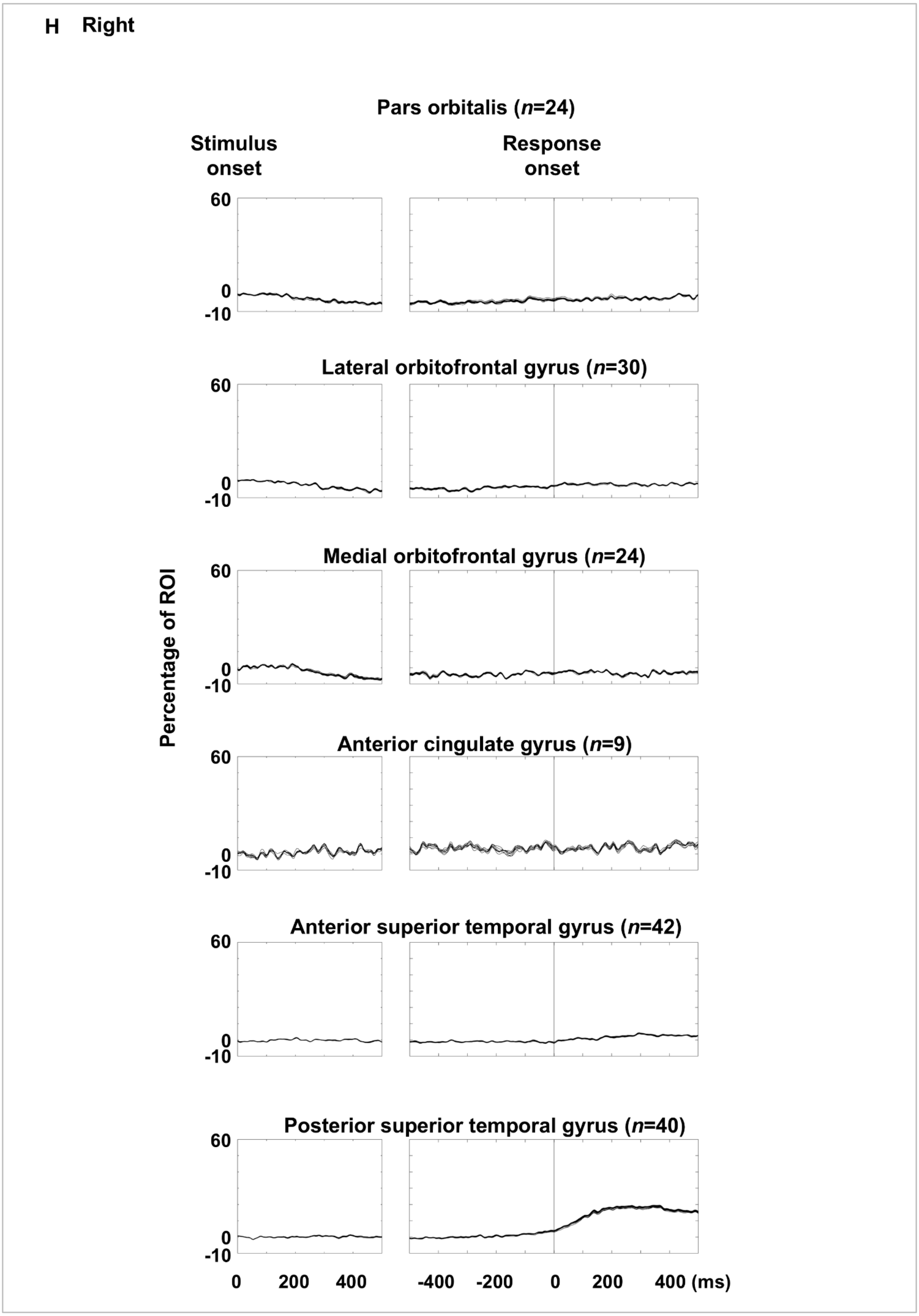
H Right
Pars orbitalis (n=24)
Stimulus onset
Response onset
60
0
-10
Lateral orbitofrontal gyrus (n=30)
Medial orbitofrontal gyrus (n=24)
Percentage of ROI
Anterior cingulate gyrus (n=9)
Anterior superior temporal gyrus (n=42)
Posterior superior temporal gyrus (n=40)
0 200 400
-400 -200 0 200 400 (ms)

eFigure 12

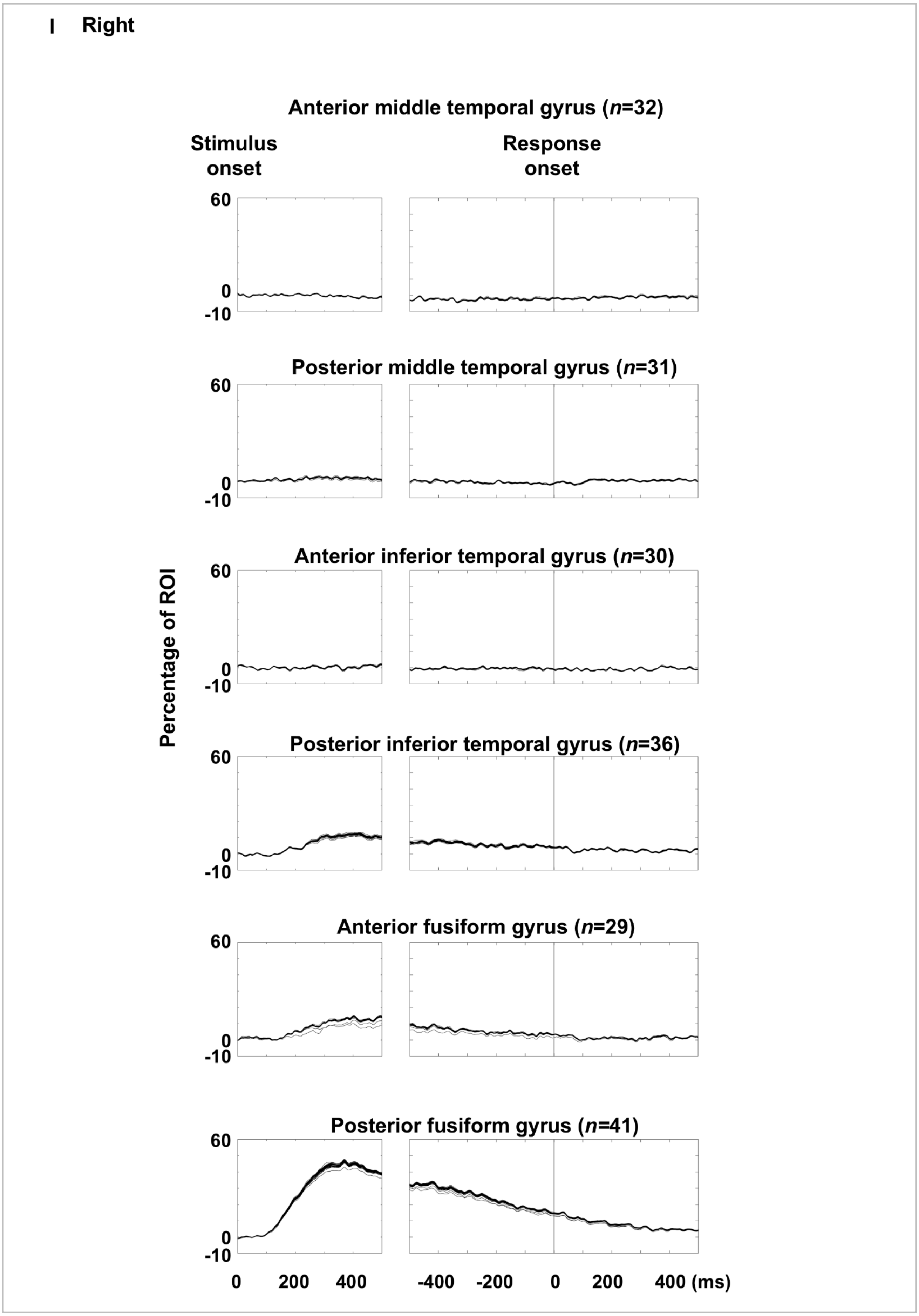
I Right
Anterior middle temporal gyrus (n=32)
Stimulus onset
Response onset
60
0
-10
Posterior middle temporal gyrus (n=31)
Anterior inferior temporal gyrus (n=30)
Posterior inferior temporal gyrus (n=36)
Anterior fusiform gyrus (n=29)
Posterior fusiform gyrus (n=41)
Percentage of ROI
0
200
400
-400
-200
0
200
400 (ms)

eFigure 12

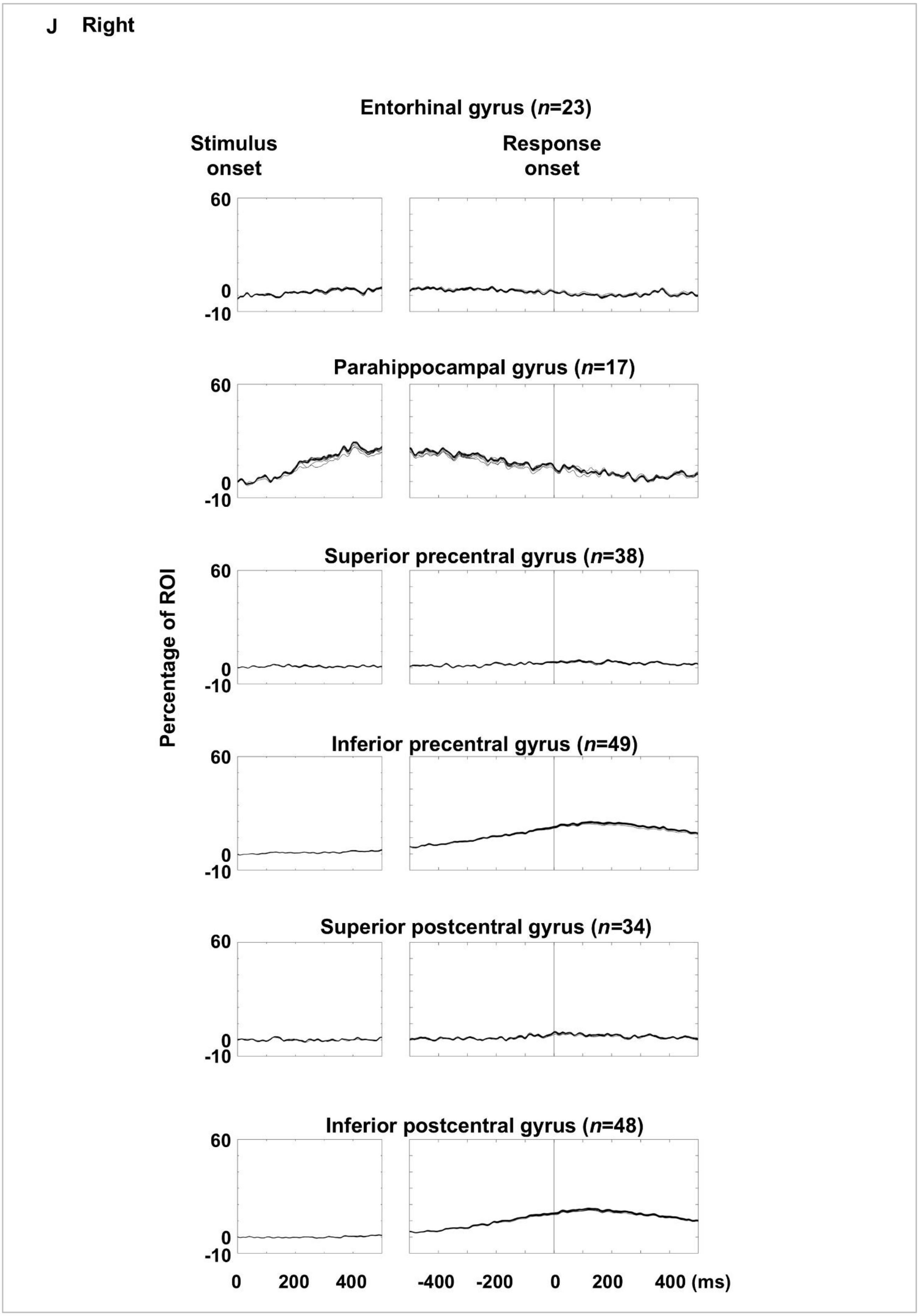
J Right
Entorhinal gyrus (n=23)
Stimulus onset
Response onset
Parahippocampal gyrus (n=17)
Superior precentral gyrus (n=38)
Inferior precentral gyrus (n=49)
Superior postcentral gyrus (n=34)
Inferior postcentral gyrus (n=48)
Percentage of ROI
60
0
-10
0
200
400
-400
-200
0
200
400 (ms)

eFigure 12

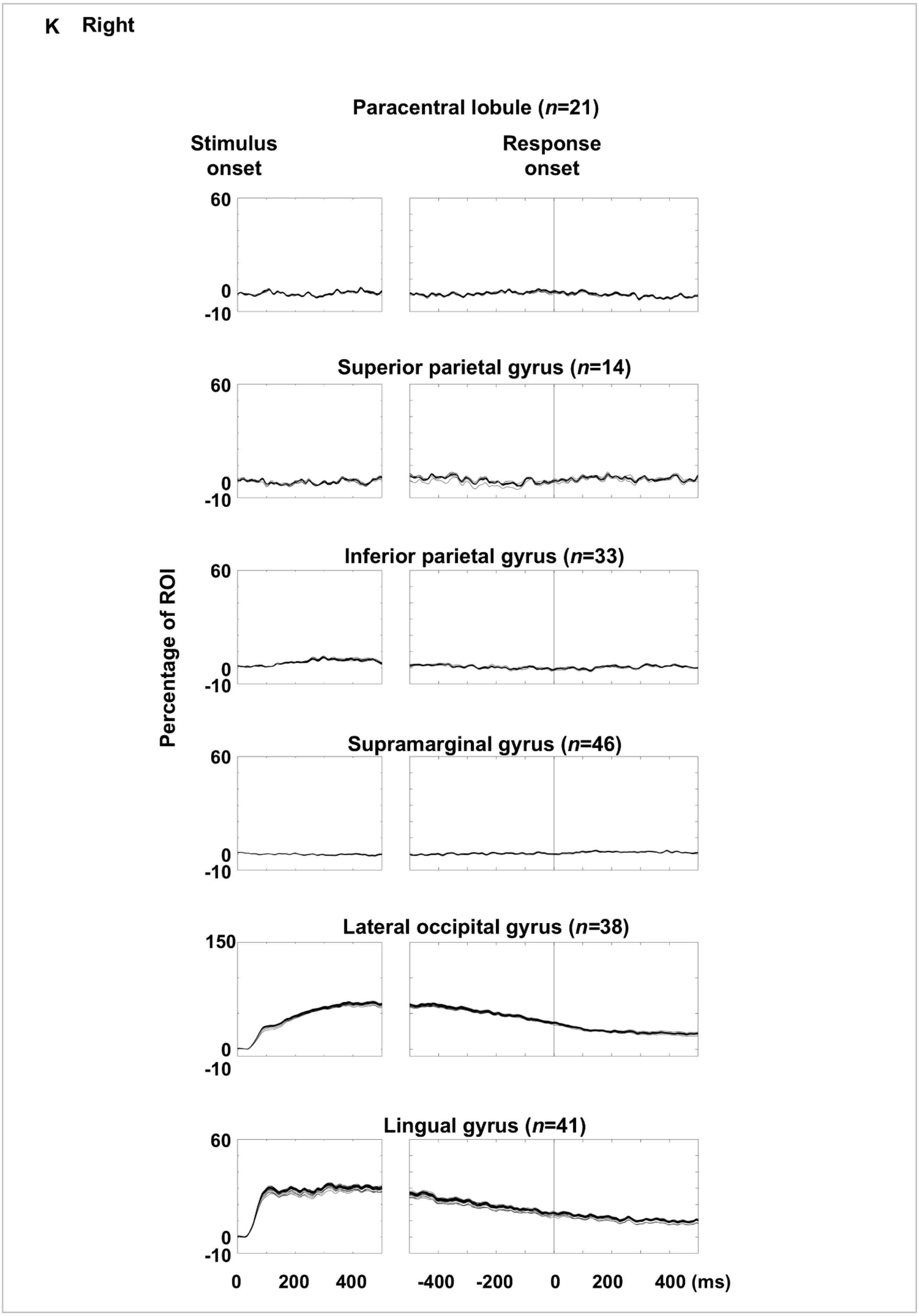
K Right
Paracentral lobule (n=21)
Stimulus onset
Response onset
60
0
-10
Superior parietal gyrus (n=14)
Inferior parietal gyrus (n=33)
Percentage of ROI
Supramarginal gyrus (n=46)
Lateral occipital gyrus (n=38)
150
Lingual gyrus (n=41)
0 200 400
-400 -200 0 200 400 (ms)

eFigure 12

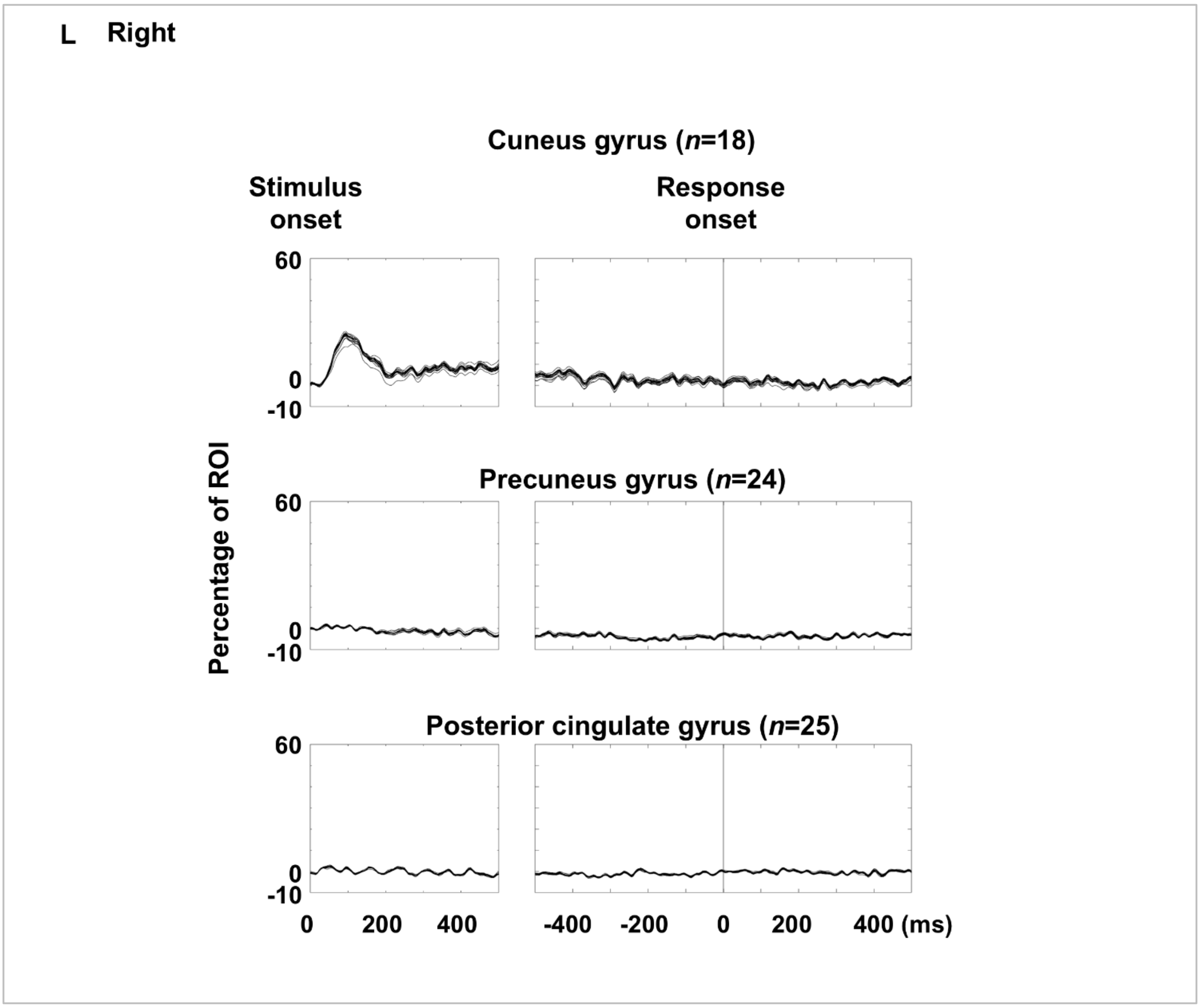


**eFigure 12. Picture naming-related high-gamma amplitude dynamics at regions of interest (ROIs): leave-one-patient-out analysis.** Each plot shows the dynamics of high-gamma amplitude at each ROI (% change relative to baseline) averaged across all patients except one excluded patient. **A–F** Left hemispheric ROIs. **G-L** Right hemispheric ROIs. To demonstrate that high-gamma dynamics are robust and not driven disproportionately by individual patients, we computed Spearman’s correlation coefficient (rho value) repeatedly, each time excluding a different patient's data (leave-one-patient-out approach). The mean Spearman’s rho value at each ROI ranged from 0.955 to 0.999. One-sample *t*-tests confirmed mean rho values significantly greater than zero at all 66 ROIs (p-value range: $3.61 \times 10^{-211}$ to $1.50 \times 10^{-12}$).
